# Rapid, large-scale species discovery in hyperdiverse taxa using 1D MinION sequencing

**DOI:** 10.1101/622365

**Authors:** Amrita Srivathsan, Emily Hartop, Jayanthi Puniamoorthy, Wan Ting Lee, Sujatha Narayanan Kutty, Olavi Kurina, Rudolf Meier

**Affiliations:** Department of Biological Sciences, National University of Singapore, 14 Science Drive 4; Zoology Department, Stockholms Universitet, Stockholm, Sweden; Estonian University of Life Sciences, Kreutzwaldi 5D, Tartu, Estonia; Tropical Marine Science Institute, National University of Singapore, Singapore; Station Linné, Öland, Sweden; Naturhistoriska Riksmuseet, Stockholm, Sweden

**Keywords:** NGS barcoding, DNA barcoding, Nanopore sequencing, MinION, large-scale species discovery

## Abstract

**Background:** More than 80% of all animal species remain unknown to science. Most of these species live in the tropics and belong to animal taxa that combine small body size with high specimen abundance and large species richness. For such clades, using morphology for species discovery is slow because large numbers of specimens must be sorted using detailed microscopic investigations. Fortunately, species discovery could be greatly accelerated if DNA sequences could be used for sorting specimens to species. Morphological verification of such “molecular Operational Taxonomic Units” (mOTUs) could then be based on dissection of a small subset of specimens. However, this approach requires cost-effective and low-tech DNA barcoding techniques because well equipped, well-funded molecular laboratories are not readily available in many biodiverse countries.

**Results:** We here document how MinION sequencing can be used for large-scale species discovery in a specimen- and species-rich taxon like the hyper-diverse fly family Phoridae (Diptera). We sequenced 7,059 specimens collected in a single Malaise trap in Kibale National Park, Uganda over the short period of eight weeks. We discovered >650 species which exceeded the number of phorid species currently described for the entire Afrotropical region. The barcodes were obtained using an improved low-cost MinION pipeline that increased the barcoding capacity sevenfold from 500 to 3,500 barcodes per flowcell. This was achieved by adopting 1D sequencing, re-sequencing weak amplicons on a used flowcell, and improving demultiplexing. Comparison with Illumina data revealed that the MinION barcodes were very accurate (99.99% accuracy, 0.46% Ns) and thus yielded very similar species units (match ratio: 0.991). Morphological examination of 100 mOTUs also confirmed good congruence with morphology (93% of mOTUs; >99% of specimens) and revealed that 90% of the putative species belong to a neglected, megadiverse genus (*Megaselia*). We demonstrate for one *Megaselia* species how the molecular data can guide the description of a new species (*Megaselia sepsioides* sp. nov.).

**Conclusions:** We document that one field site in Africa can be home to an estimated 1000 species of phorids and speculate that the Afrotropical diversity could exceed 100,000 species. We furthermore conclude that low-cost MinION sequencers are very suitable for reliable, rapid, and large-scale species discovery in hyperdiverse taxa. MinION sequencing could quickly reveal the extent of the unknown diversity and is especially suitable for biodiverse countries with limited access to capital-intensive sequencing facilities.

## INTRODUCTION

In 2011 the former president of the Royal Society, Robert May, wrote that “[w]e are astonishingly ignorant about how many species are alive on earth today, and even more ignorant about how many we can lose [and] yet still maintain ecosystem services that humanity ultimately depends upon.” [1] Little has changed since 2011 and >80% of all extant animal species remain unknown to science [2]. Most of these unknown species belong to hyper-diverse and species-rich invertebrate clades. They are ubiquitous, contain most of the multicellular animal species, and often occur in great abundance. However, research on the species diversity of such clades is underdeveloped because it requires the examination of large numbers of specimens. These specimens have to be grouped into species before they can be either identified (if they belong to a known species) or described (if they are unknown to science).

In invertebrates, species discovery often starts with obtaining specimens via bulk sampling methods. In insects, one of the most widely used methods is Malaise trapping. Such traps routinely collect thousands, or even tens of thousands, of specimens per site and week; i.e., sorting all specimens to species-level virtually never happens and the world’s natural history museums store billions of unsorted specimens. Species-level sorting is usually restricted to a few taxa with small to moderate numbers of specimens. It is accomplished in two stages. The first is grouping specimens into easily identifiable major taxa (e.g., major groups of beetles, flies, wasps). This type of pre-sorting is usually accomplished by parataxonomists with basic training in morphology (e.g., students). The main challenge is the second sorting stage; i.e., sorting to species-level. This work is best carried out by taxonomic experts whose techniques are, however, mostly effective for taxa that have fairly small numbers of specimens and species. In contrast, large, hyperdiverse and abundant taxa are ill-suited because they require dissection and microscopic study of many specimens. An alternative to species-level sorting by taxonomists is a hybrid approach that combines rapid pre-sorting to “morpho-species” by parataxonomists with subsequent verification of morpho-species via DNA barcodes that are obtained for a few specimens for each morpho-species [3]. DNA barcodes are only obtained for few specimens because it would be too time-consuming and expensive to generate them for all specimens using the traditional DNA barcoding methods that require formal DNA extractions and Sanger sequencing [4]. Unfortunately, this widely used hybrid approach has three problems. Firstly, species-level sorting by parataxonomists is very imprecise [5, 6]. Secondly, small-scale DNA barcoding tends to overlook morphologically cryptic species. Thirdly, the hybrid approach requires a lot of manpower for morpho-species sorting.

An alternative approach to species discovery is the ‘reverse workflow’ of Wang et al. (2018) [4]. Here, every specimen in a sample is DNA barcoded with minimal or no damage to the specimen [4, 7, 8] using simplified DNA extraction protocols and Illumina sequencing [9]. After barcoding, the specimens are grouped into molecular Operational Taxonomic Units (mOTUs) that in most cases represent species. The confirmation of these mOTUs as species comes last. Taxonomic experts use morphology to study a subset of the specimens that were pre-sorted to putative species based on DNA sequences. The selection of the specimens can be guided by the genetic distance between individuals [3]. This “reverse workflow” has the advantage that species-level sorting relies on DNA sequencing which can potentially be automated. It also associates morphologically dissimilar males, females, and immature specimens that belong to the same species [7]. However, barcoding all specimens in a sample is unrealistically expensive with traditional Sanger sequencing. The implementation of the reverse workflow thus requires more cost-effective sequencing solutions that are now provided by High Throughput Sequencing platforms (e.g., Illumina, Nanopore, PacBio: [4, 8, 10–13]). For example, tens of thousands of specimens can be barcoded on a single lane of Illumina HiSeq with the total cost of a barcode being as low as 0.17 USD (including PCR cost, see discussion in Wang et al., 2018 [4]). However, due to read length restrictions, barcodes obtained with Illumina are <400 bp and new solutions for obtaining full-length barcodes based on PacBio [10] or MinION [14] sequencing have only recently emerged.

Unfortunately, barcoding with Illumina and PacBio sequencing has some downsides. Firstly, both technologies are only cost-effective if >10,000 specimens are simultaneously barcoded because the cost of flowcells is high. Secondly, sequencing must usually be outsourced; i.e., amplicon pools have to be shipped to sequencing facilities. This is not a major concern in developed countries, but it is often a problem for species discovery research in countries that lack capital-intensive, high-throughput sequencing facilities or have restrictive regulations with regard to the export of genetic material. It would thus be desirable to have alternative sequencing techniques that are fast, scalable, cost-effective, and require low initial investment. Such solutions would be particularly useful if barcoding could be accomplished under field conditions and/or by citizen scientists [15–18].

Oxford Nanopore’s MinION has the potential to be such a solution. It is a low-cost, portable device and delivers real-time sequencing. However, it unfortunately still generates error-prone data (ca. 10-15% [19]) at a fairly high cost per base pair. Therefore, its use and reliability for large-scale specimen barcoding remains poorly explored. A first step toward the use of MinION for barcoding was the recent demonstration that 500 DNA barcodes can be obtained on one flowcell of MinION using 1D^2^ sequencing [14]. The study increased the throughput of one MinION flowcell by one order of magnitude compared to existing protocols. However, the scale was arguably still not sufficient for large-scale species discovery where thousands of specimens have to be barcoded. Furthermore, the experiment used 1D^2^ sequencing, which requires complicated and time-consuming library preparation techniques and access to computer servers for base-calling. Here, we test whether the more straightforward, but less accurate, 1D sequencing can be used for large-scale species discovery.

Improved species discovery techniques are particularly needed for hyperdiverse clades of invertebrates that have many species in the tropics. A good example are insects whose diversity is concentrated in four hyper-diverse insect orders: Coleoptera (beetles), Diptera (midges and flies), Hymenoptera (bees, wasps, and ants), and Lepidoptera (moths and butterflies). Species estimates for all Insecta vary between 3 and 13 million (reviewed by Stork, 2018 [20]) with only ca. 1,000,000 currently described [21]. Historically, Coleoptera has been considered the most species-rich order of insects which is said to have led the evolutionary biologist J. B. S. Haldane to remark that the creator must have had an “inordinate fondness for beetles.” [22]. However, it now appears that the impression that Coleoptera is the most species-rich order may have been due to an inordinate fondness of taxonomists for beetles. Recent studies suggest that Diptera and Hymenoptera may be more species-rich. For example, Forbes et al. [23] proposed that Hymenoptera contained more species than either Diptera or Coleoptera based on parasite host ratios for Microhymenoptera. Similarly, a large barcoding study of Canadian insects found that Hymenoptera and Diptera together accounted for two thirds of the 46,937 molecular Operational Units found (in the form of BINs or Barcode Index Numbers [24]). The study predicted that one dipteran family alone, gall midges (Cecidomyiidae), may have 16,000 species in Canada. Once extrapolated to a worldwide scale, the authors estimated that 1.8 million of the 10 million predicted insect species could be cecidomyiids [25]; i.e., a single family of Diptera would far surpass the number of described beetle species. Other studies similarly hint at the extraordinary richness of Diptera. For example, the Zurqui All Diptera Biodiversity Inventory (ZADBI) of a single site in Costa Rica was heavily reliant on specimens collected with two Malaise traps over one year [26]. Only 41,001 specimens (a small fraction of the hundreds of thousands collected) were studied by taxonomic experts [27]. These specimens belonged to 4,332 species of Diptera, of which 800 were Cecidomyiidae and 404 Phoridae [27], the fly family of focus here. Phoridae, or scuttle flies, is a family of true flies with approximately 4,300 described species [28]. Currently, only 466 species of phorids have been described for the Afrotropical Region [28] while Henry Disney, a world expert on the family, has recorded 75 species of phorids in his suburban garden in Cambridge alone [29]. Similarly, the BioSCAN project in Los Angeles recorded up to 82 species in city backyards [29]. These numbers make it very likely that the Afrotropical fauna is very large and currently vastly understudied. But not all phorid taxa are equally poorly sampled. The main obstacle to understanding phorid diversity is *Megaselia* Rondani which contains >1,600 of the 4,300 described species. This makes *Megaselia* “one of the largest, most biologically diverse and taxonomically difficult genera in the entire animal kingdom” [30]. In groups like *Megaselia*, the obstacles to completing species discovery with traditional methods appear insurmountable. Extremely large numbers of specimens are routinely collected which can belong to very large numbers of species. This makes sorting such samples into species-level units using traditional workflows very labour-intensive. Rare and new species are often hidden among very large numbers of common and described species. The rare species cannot be found without the microscopic study of thousands of specimens for which prodigious notes have to be taken. Detailed drawings must be prepared (for *Megaselia* drawings of male genitalia are essential) – often based on dissections and slide mounts. This traditional workflow thus discourages all but the most tenacious taxonomists from taking up the study of hyper-diverse genera within insects.

Here, we test whether 1D MinION sequencing can help to reveal phorid diversity more comprehensively by relegating the sorting to species level to sequencing. MinION sequencing is here applied to ca. 30% of the phorid specimens that were collected in a single Malaise trap in Kibale National Park, Uganda. We describe how we processed ∼8,700 specimens, obtained ∼7,000 accurate barcodes, and found >650 putative species. All this was accomplished using a workflow that would take less than a month.

## RESULTS

### 1. MinION based DNA barcoding

The experiment was designed to obtain full-length COI barcodes via tagged amplicon sequencing for two sets of specimens. A total of 8,699 phorid flies were processed (Set 1: 4,275; Set 2: 4,519; 95 specimens were duplicated in both sets) (Fig. 1). In order to assess amplification success rates, a subset of PCR products for each of the ninety-two 96-well plates were verified with agarose gels. Amplification success rates were estimated to be 86% and 74% for the two sets of specimens (80.7% overall); i.e., we estimated that >3,600 and >3,300 DNA barcodes should be obtainable via MinION sequencing given that gels tend to underestimate amplification success rates for weak amplicons that cannot be reliably visualized with commercial dyes (Table 1). The PCR products for each set were pooled and sequenced using MinION (set 1: 7,035,075; set 2: 7,179,121 1D nanopore reads). Both sets were sequenced in two MinION runs. The first run for each set was based on the pooled PCR products for all specimens in the set. It generated 3,069,048 and 4,853,363 reads, respectively. The results of the first run were used to estimate coverage for each PCR product. Products with weak coverage (<=50x) were re-pooled and re-sequenced (set 1: 2,172 amplicons; set 2: 2,211 amplicons). This added 3,966,027 and 2,325,758 reads to each set and improved the coverage of many low-coverage barcodes (Fig. 2).

**Figure 1:**
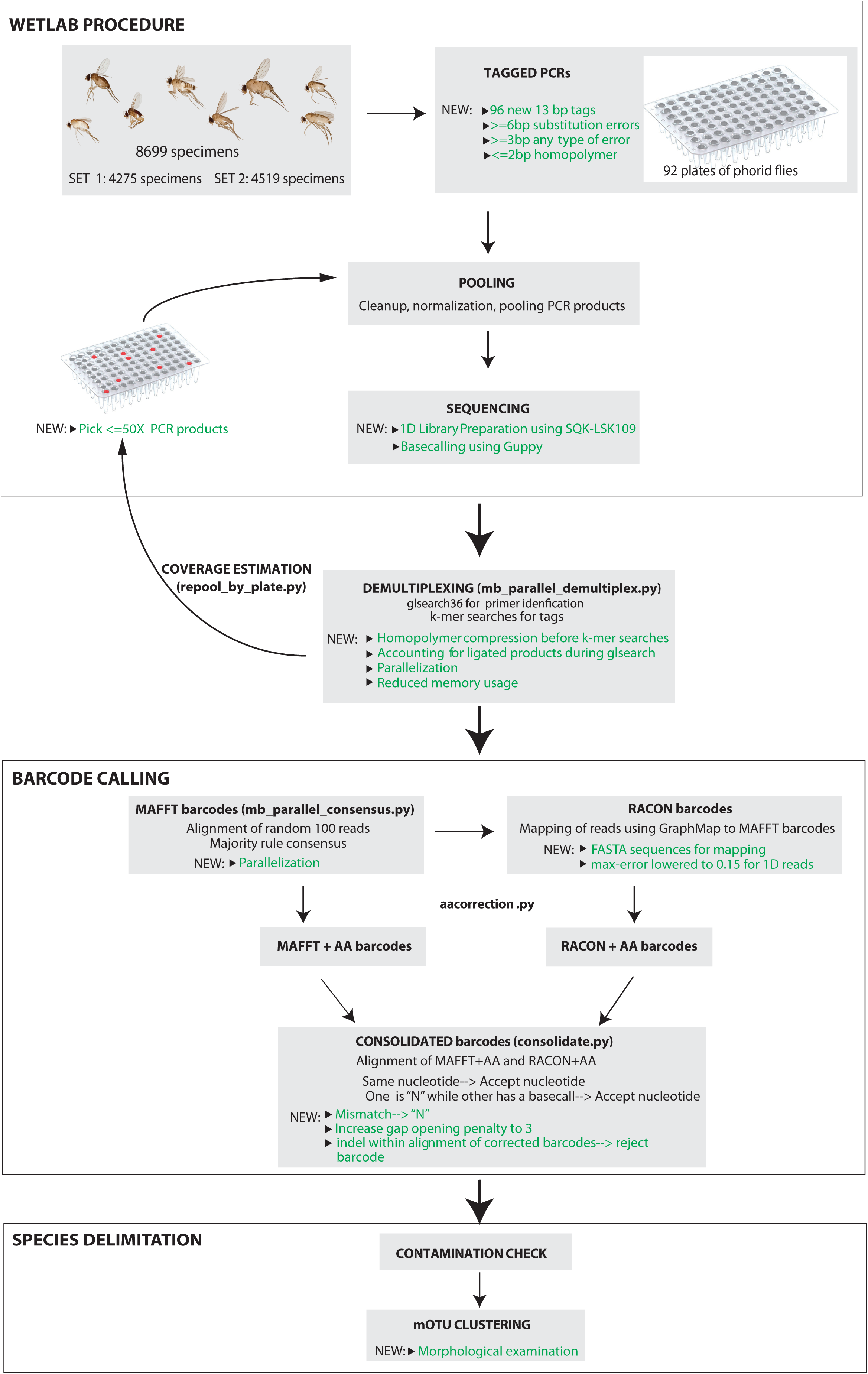
Flowchart for generating MinION barcodes from experimental set-up to final barcodes. The novel steps introduced in this study are highlighted in green and the scripts available in *miniBarcoder* for analyses are further indicated.

**Figure 2:**
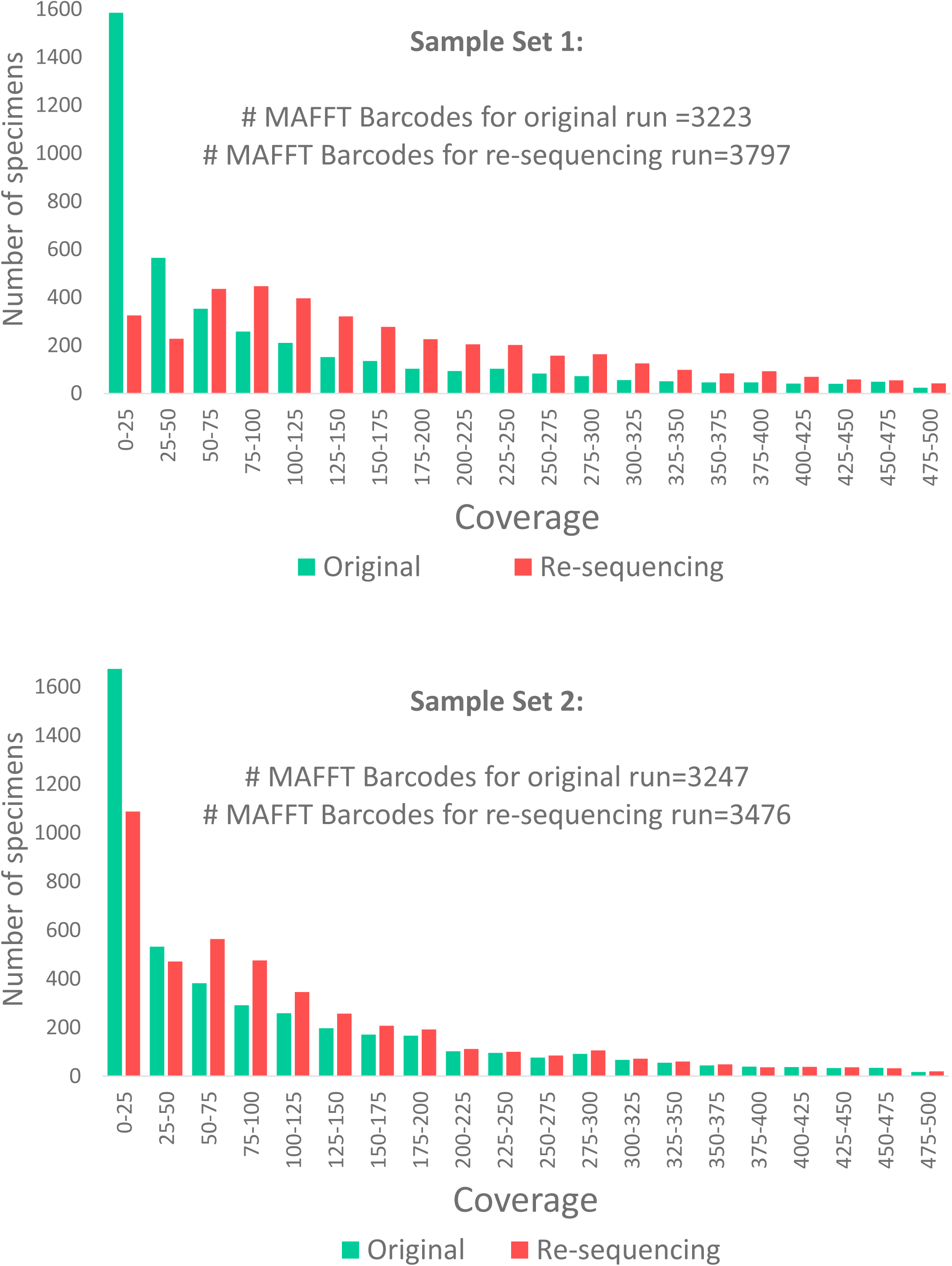
Effect of re-pooling on coverage of barcodes for both sets of specimens. Barcodes with coverage <50X were re-pooled and hence the coverage of these barcodes increases.

**Table 1:** Number of reads and barcodes generated via MinION sequencing.

|  | Set 1: Two flowcells | Set 2: One flowcell | Combined (set 1 & 2)* |
| --- | --- | --- | --- |
| # Specimens | 4,275 | 4,519 | 8,699 |
| Resequencing (re-pooled) | 2,172 | 2,211 |  |
| # reads/# reads >600 bp | 7,035,075/3,703,712 | 7,179,121/2,652,657 | NA |
| Initial sequencing (all) | 3,069,048/1,942,212 | 4,853,363/2,250,591 |  |
| Resequencing (re-pooled) | 3,966,027/1,761,500 | 2,325,758/402,066 |  |
| # demultiplexed reads | 898,979 (24.3%) | 647,152 (24.4%) |  |
| Initial sequencing (all) | 562,434 (29%) | 561,383 (24.9%) |  |
| Resequencing (re-pooled | 336,545 (19%) | 85,769 (21.3%) |  |
| Combined results of original and resequencing runs |  |  |  |
| # specimens with >=5X coverage | 4,227 (98.9%) | 4,287 (94.9%) | 8,428 (96.9%) |
| # MAFFT barcodes <1% N's | 3,797 (88.8%) | 3,476 (76.9%) | 7,221 (83%) |
| # MAFFT + AA barcodes | 3,771 (88.2%) | 3,459 (76.5%) | 7,178 (82.5%) |
| # RACON barcodes | 3,797 (88.8%) | 3,476 (76.9%) | 7,221 (83%) |
| # RACON +AA barcodes | 3,788 (88.6%) | 3,461 (76.6%) | 7,194 (82.7%) |
| # Consolidated barcodes | 3,762 (88%) | 3,446 (76.3%) | 7,155 (82.3%) |
| # Consolidated barcodes (non-phorids removed) | 3,727 (87.2%) | 3,426 (75.8%) | 7,059 (81.1%) |
| # mOTUS (2/3/4%) |  |  | 705/663/613 |

\* one plate was accidentally sequenced in both runs, duplicates removed for combined set.

The combined data were processed using an improved version of a bioinformatics pipeline introduced in Srivathsan et al. [14]. The improvements led to a higher demultiplexing rate (14% increase for set 1: 898,979 vs. 787,239 reads; 9% increase for set 2: 647,152 vs. 593,131 reads) and faster demultiplexing (10X using 4 cores: demultiplexing in 9 min vs 87 min for one of the datasets).

### 2. Assessment of demultiplexing accuracy

We indirectly assessed the accuracy of the demultiplexing pipeline by testing whether reads would be incorrectly demultiplexed into bins belonging to unused tag combinations. This happened for a very small proportion of reads (0.23%: 2,054 of 900,698 reads in set 1; 0.44%: 2,837 of 649,587 reads in set 2). Note that such low error rates are unlikely to yield poor quality barcodes given that the average coverage per amplicon was 210X (set 1) and 143X (set 2). Surprisingly, 37% and 69% of these incorrectly demultiplexed reads were due to one tag: GTCCAACTTCAGT although the edit distances between all tag-pairs were high (>=5 bp); i.e., it is currently unclear whether the underperforming tag was due to a primer synthesis issue, systematic sequencing bias, or a wet lab problem (Additional File 1: Fig 1). Out of caution, we provided four additional tag sequences that can be used as replacements (Additional File 2).

### 3. Barcode calling

Demultiplexing all data and calling preliminary barcodes generated 3,797 and 3,476 preliminary “MAFFT barcodes” with >=5X coverage and <1% ambiguous bases. These barcodes were subjected to correction using RACON [31] which yielded the same number of “RACON barcodes”. We overall obtained 7,221 MAFFT and RACON barcodes. These preliminary barcodes still contained indel and substitution errors that were corrected with an amino-acid correction pipeline that was first implemented in Srivathsan et al. [14]. It yielded 7,178 AA-corrected MAFFT barcodes (“MAFFT+AA”) and 7,194 AA-corrected RACON barcodes (“RACON+AA”). This pipeline rejects barcodes that have five or more consecutive indel errors so that there are fewer corrected than uncorrected barcodes. Finally, the two sets of corrected barcodes were consolidated. This yielded a set of 7,155 consolidated, final barcodes. During this process, the consolidated barcodes where the alignment of MAFFT+AA and RACON+AA barcodes required the insertion of indels were rejected because AA-corrected barcodes are expected to be indel-free. The overall barcoding success rate was thus 82.3% (7,155 barcodes for 8,699 specimens). This was close to the expected 80.7% success rate based on gel electrophoresis; i.e., MinION sequencing consistently produced sequence data for successfully amplified products.

A subsequent contamination check via BLAST revealed that of the 7,155 barcodes, 96 barcodes were unlikely to be phorid flies (<1.5%). These included 53 barcodes with matches to *Wolbachia, Rickettsia,* nematodes, human, and occasionally insects from other families (e.g. *Drosophila, Hemipyrellia*) and another 43 that did not belong to Phoridae and were incorrectly pre-sorted by parataxonomists. After removal of these, we retained 7,059 confirmed phorid barcodes. Lastly, we inspected the reads obtained for the 92 negative PCR controls (1 per microplate). Five negatives yielded MAFFT barcodes. Four of these had a >97% match to non-phorids (two humans, one fish, one mollusc) and were eliminated. One low coverage (13X) negative survived all filters and matched phorid COI. It was removed after ascertaining that it did not impact the accuracy of the barcodes in the plate. This could be tested by comparing the MinION barcodes for the plate with Illumina barcodes obtained from different PCR products for the same DNA extraction plate (see below).

### 4. Comparison of MinION barcodes with Illumina barcodes

Illumina barcodes were obtained for 6,251 of the 7,059 specimens with MinION barcodes using a different set of primers that amplified a 313 bp subset of the full-length barcodes; i.e. comparison with MinION sequencing is based on 48% of the MinION sequence. The comparisons showed that the uncorrected MAFFT and RACON barcodes had an accuracy of 99.61% and 99.51% (Table 2). Correction of these barcodes with the amino-acid correction pipeline improved the accuracy considerably (>99.9% in all cases). The barcodes were corrected after optimizing a parameter that is here called “namino” because it specifies the length of the AA motifs that is used for correction. Overall, namino=2 was found to optimize overall accuracy while minimizing the number of inaccurate barcodes. We found that MAFFT+AA barcodes were more accurate than RACON+AA barcodes, but MAFFT+AA barcodes contained a much higher number of ambiguous nucleotides (Fig. 3). When RACON+AA and MAFFT+AA barcodes were consolidated, the resulting “consolidated barcodes” were found to be highly accurate (99.99%) and containing few ambiguous bases (median = 0.3%, average=0.46%). These accuracy rates were obtained after excluding the <0.5% specimens that had >3% divergence with corresponding Illumina barcodes. Such barcode discrepancies are likely due to wet-lab errors (e.g., amplification of residual contaminating signals, see details in methods). Note that such errors are regularly observed in large-scale barcoding projects. For examples, a recent study by Hebert et al. [10] using PacBio Sequel for DNA barcoding found that 1.5-1.6% of the specimens had high abundances of non-target sequences.

**Figure 3:**
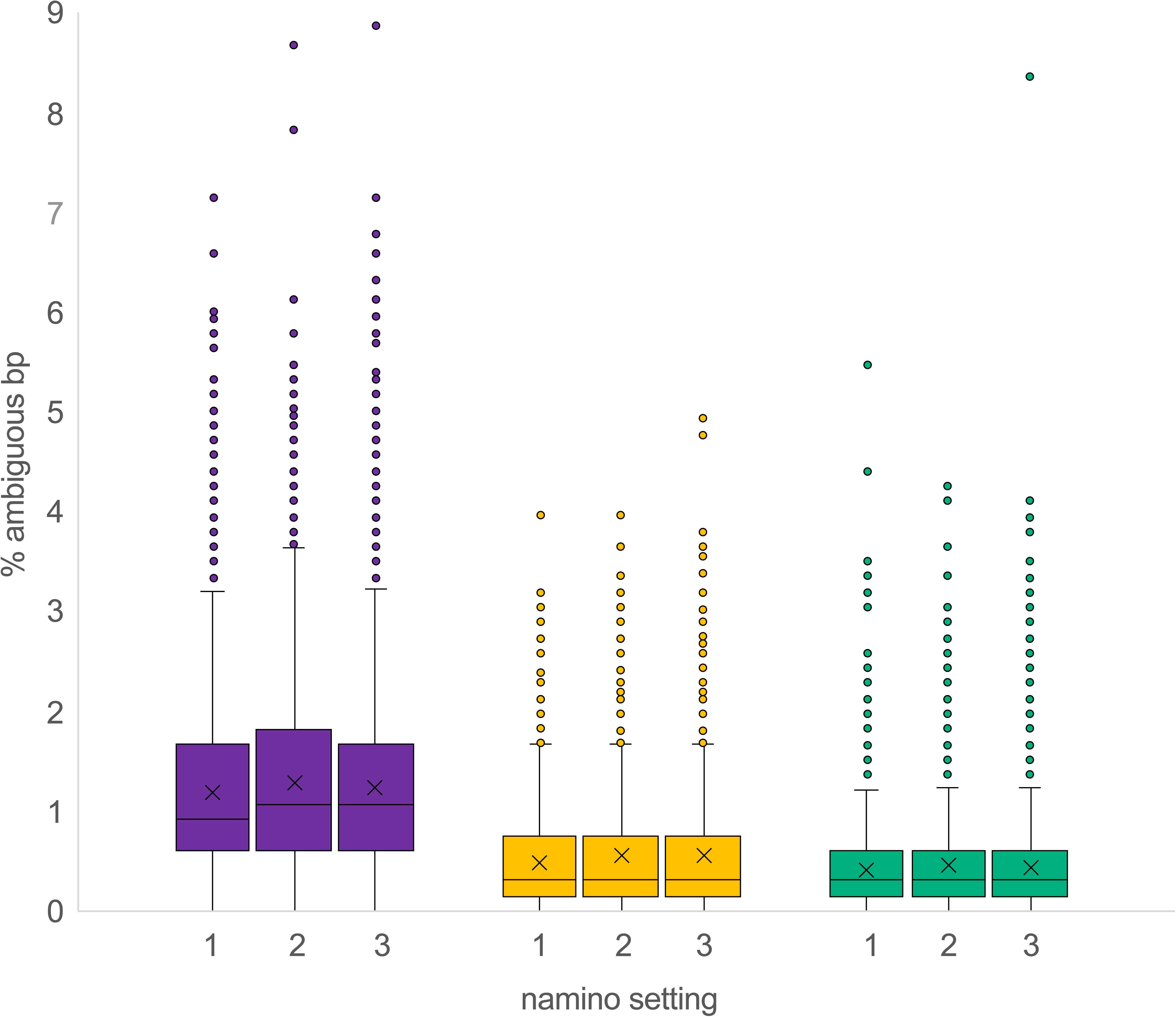
Ambiguities in MAFFT+AA (Purple), RACON+AA (Yellow) and Consolidated barcodes (Green) with varying namino parameters (1,2 and 3). One outlier value for Racon+3AA barcode was excluded from the plot. The plot shows that the consolidated barcodes have few ambiguities remaining.

**Table 2.** Accuracy of MinION as assessed by Illumina barcodes. The MinION barcodes were trimmed to the 313 bp that were sequenced using Illumina. The overall optimal strategy is “Consolidated (namino=2)”. Optimal congruence values are highlighted in bold.

| Dataset | # compared with Illumina/ # 3% mOTUs | Accuracy Mean (stdev) | %Ns | # barcodes with errors/# >3% errors | mOTU richness deviation between MinION and Illumina barcodes |  |  |
| --- | --- | --- | --- | --- | --- | --- | --- |
|  |  |  |  |  | 2% | 3% | 4% |
| MAFFT | 6,291/641 | 99.6136 (0.37) | 0.18 (0.2) | 4,473/30 | -3 (-0.44%) | **-2 (-0.31%)** | -11 (-1.85%) |
| RACON | 6,291/645 | 99.5097 (0.48) | <0.001 (0.01) | 4,494/36 | 12 (1.72%) | **2 (0.31%)** | **-3 (-0.5%)** |
| MAFFT + AA (namino=1) | 6,269/635 | 99.9689 (0.19) | 1.19 (0.84) | 273/27 | -6 (-0.89%) | -6 (-0.94%) | -6 (-1.02%) |
| MAFFT + AA (namino=2) | 6,269/635 | 99.9802 (0.15) | 1.28 (0.92) | 216/28 | -8 (-1.19%) | -6 (-0.94%) | -14 (-2.37%) |
| MAFFT + AA (namino=3) | 6,269/633 | 99.9685 (0.18) | 1.25 (0.91) | 310/27 | -8 (-1.19%) | -8 (-1.26%) | -17 (-2.9%) |
| RACON + AA (namino=1) | 6,273/639 | 99.9636 (0.19) | 0.48 (0.47) | 392/26 | -4 (-0.59%) | -4 (-0.63%) | -13 (-2.19%) |
| RACON + AA (namino=2) | 6,273/635 | 99.9736 (0.16) | 0.55 (0.55) | 288/26 | -5 (-0.74%) | -8 (-1.26%) | -14 (-2.36%) |
| RACON + AA (namino=3) | 6,273/636 | 99.9684 (0.17) | 0.57 (0.6) | 345/28 | **-1 (-0.15%)** | -7 (-1.1%) | -13 (-2.19%) |
| Consolidated (namino=1) | 6,229/639 | 99.9849 (0.12) | 0.41 (0.44) | 191/26 | -2 (-0.29%) | **-2 (-0.31%)** | **-3 (-0.5%)** |
| **Consolidated (namino=2)** | 6,251/638 | **99.9859 (0.11)** | **0.46 (0.5)** | **184/25** | -2 (-0.29%) | -3 (-0.47%) | -5 (-0.83%) |
| Consolidated (namino=3) | 6,245/639 | 99.9795 (0.12) | 0.44 (0.49) | 285/26 | -4 (-0.59%) | **-2 (-0.31%)** | -4 (-0.67%) |

### 5. Comparison of MinION and Illumina barcodes at mOTU-level

Given that the barcodes were obtained for the purpose of species richness estimates, we compared the mOTU richness estimated based on the different barcode sets against those obtained with Illumina barcodes. For this purpose, we trimmed the MinION barcode sets to the 313 bp fragment that was sequenced using Illumina. mOTU richness was very similar (Table 2). However, comparison of mOTU numbers alone does not imply that the same specimens were grouped into mOTUs obtained with the MinION and Illumina barcodes. One also has to assess whether the contents of the mOTUs are identical. We thus calculated the match ratio for the datasets (3% clustering threshold). We found that all five barcode sets (MAFFT, RACON, MAFFT+AA, RACON+AA, and consolidated barcodes, namino=2) had high match ratios (>0.95). The consolidated and RACON barcodes performed best with match ratios of >0.98 (consolidated barcodes: 0.991, RACON: 0.981). However, upon closer inspection the multiple sequence alignment (MSA) for the RACON barcodes contained indels while the consolidated barcodes are insertion-free and contain next to no deletions. Here, single bp deletions were found in the first twenty bps of the barcode for 3/7059 specimens. The largest number of indels was found in the MSA of uncorrected RACON barcodes which indicated that the RACON barcodes retained a fair number of indel errors; i.e., RACON barcodes may not be of sufficient quality for submission to sequence databases. We thus recommend the usage of consolidated barcodes. This recommendation is based on maximizing per-base accuracy (see below), yielding high-quality alignments, and revealing very similar mOTU diversity and composition (high match ratio) when compared to Illumina barcodes.

Given the different length of MinION and Illumina barcodes, we also compared the mOTUs obtained by full-length MinION barcodes (658 bp) with the mOTUs obtained with Illumina barcodes for those specimens for which both types of data were available. The match ratio was again high (0.951). For incongruent clusters, we analysed at which distance threshold they would become congruent. We found that all clusters were congruent within the 1.9-3.7% range; i.e., the remaining 345 bp are not showing a major deviation from the signal obtained from the 313 bp fragment (Additional File 3). We next characterized if there was an increase in error in the 345 bp stretch of the MinION sequence that could not be directly compared to Illumina sequence: if this were the case, we would expect that spurious base calls would increase genetic distances for specimens. However, we found the opposite: in 18 of 21 cases, the threshold was lowered, i.e., the 345 additional nucleotides reduced the minimum distance in the cluster (Additional File 3).

### 6. Species richness estimation

After these quality checks, we proceeded to characterize the diversity of phorid flies based on the MinION barcodes of highest accuracy based on comparison with Illumina; i.e., the consolidated barcodes (namino=2). We obtained a mean of 660 mOTUs when the thresholds were varied from 2-4% (2%: 705, 3%: 663, 4%: 613). These thresholds are widely used in the literature, but also supported by empirical data from GenBank. GenBank has 12,072 phorid sequences with species-level identifications belonging to 106 species. The intraspecific variability is overwhelmingly <3% (>95% of pairwise distances) and the match ratios between mOTUs and species identifications from GenBank are maximized for clustering thresholds of 2 - 3% (Additional File 1: Fig S2, S3). In addition to clustering the barcodes based on *a priori* thresholds, we also used species delimitation based on Poisson Tree Processes (PTP) to estimate the number of species for the phorids from the trap. It yielded even higher richness estimate of 747 putative species than the threshold-based methods. Lastly, we used species accumulation and Chao 1 curves (mOTUs at 3%) to estimate the full phorid diversity of the Ugandan site. We find that the curves have yet to reach a plateau, but the shape of the curves suggests an estimated diversity of ∼1,000 species of Phoridae at a single field site in Uganda, collected by one Malaise trap (Fig. 4).

**Figure 4:**
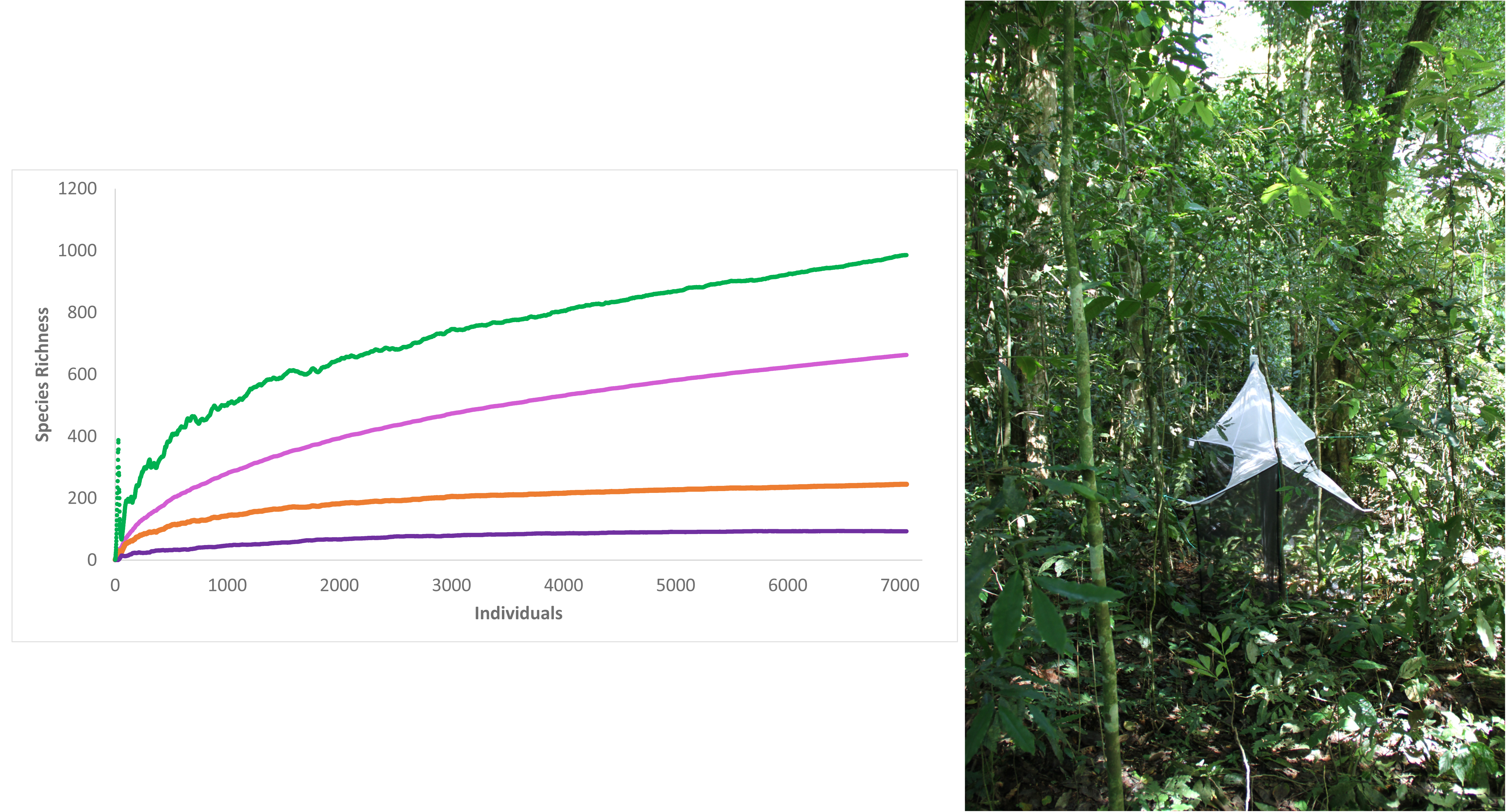
The Malaise trap that revealed the estimated >1000 mOTUs as shown by the species richness estimation curve. Green: Chao1 Mean, Pink: S (Mean), Orange: Singleton Mean, Purple: Doubleton mean.

### 7. Paralogy check

We found that the Illumina were translatable which is not expected for sequences obtained for old NuMTs. In addition, the congruence between the mOTUs estimated based on sequences for two different amplicons of different lengths and different primer specificity is very high. This would not be expected if NuMTs were regularly amplifying. We also scrutinized the read sets for Illumina amplicons for the presence of secondary phorid signal. We found such signal in 7% (30) of the 406 mOTUs with multiple specimens. Such signal can be caused by paralogs or low-level lab contamination when small amounts of template from one well contaminates the PCR reaction in another well. We suspect that much of the secondary signal is caused by the latter, but it is arguably more important that the level of secondary signal is sufficiently low that it could not significantly lower the overall species richness estimate of the site even if all secondary signal was caused by paralogy (Additional File 4).

### 8. Congruence with morphology

We conducted a morphological check of 100 randomly selected clusters (>1,500 specimens). We found that 6 of the 100 clusters contained, among other specimens, a single misplaced specimen. There was one cluster of four specimens that appeared to consist of a mixture of three morpho-species. This implies that 9 of the >1,500 examined barcoded specimens were misplaced due to lab contamination. This morphological check took ca. 30 hours. mOTUs based on barcodes are expected to lump those species that recently speciated and split species with well-differentiated populations [32]. This means that taxonomists working with mOTUs should check for signs of lumping and splitting in closely related taxa. This requires morphological examination of a subset of specimens whose selection is guided by genetic information. This is aided by keeping closely related mOTUs physically together. In the case of phorids this can be done by slide mounting representative specimens from the sub-clusters. This is here illustrated by describing one species based on a complex cluster.

#### New Species Description

During the morphological work, a distinctive new species of *Megaselia* was found. A mOTU-specific haplotype network was constructed and informed on which specimens should be studied based on morphology. The new species is here described. To continue reducing redundancy and ambiguity in species descriptions, the description of this species has excluded the character table from the method previously established for *Megaselia* [33–35] and uses a molecular and photographic description. Photographs are a key element in descriptions for large, diverse groups [36], where verbose descriptions require much time while remaining insufficiently diagnostic. Most characters that would have been in table form are clearly visible in the photographs provided.

*Megaselia sepsioides* Hartop sp. n.

`urn:lsid:zoobank.org`:pub:ED268DF2-A886-4C31-A4FB-6271C382DECE

DNA barcode for UGC0005996 (GenBank accession: MN403533)

actttatattttatttttggagcttgagctggaatagtaggtacttccttaagaatcataattcgtgctgaattaggacacccaggagcacttattggtgatgaccaaatttataatgtgattgttactgcacatgcttttattataattttttttatagtaatacctattataataggaggttttggtaattgacttgtacctttaatattaggagccccagatatggcattccctcgaatgaataatataagtttttgaatattacctccttctttaactcttttattagccagaagtatagtagaaaatggagctggaactggttgaacagtttatcctcctttatcttctagaatcgctcatagtggagcttctgttgatttagcaattttctctcttcatttagctggaatttcatctattttaggagctgtaaattttattacaacaattattaatatacgatcatcaggtattacatttgaccgaatacctctatttgtttgatctgtaggtattacagctttattgctactcttatcacttcctgttttagctggtgctattacaatactattaacagaccgaaattttaatacttcattttttgacccagcaggaggaggagatccaattttataccaacatttattc Fig. 5, 6, 7

**Figure 5:**
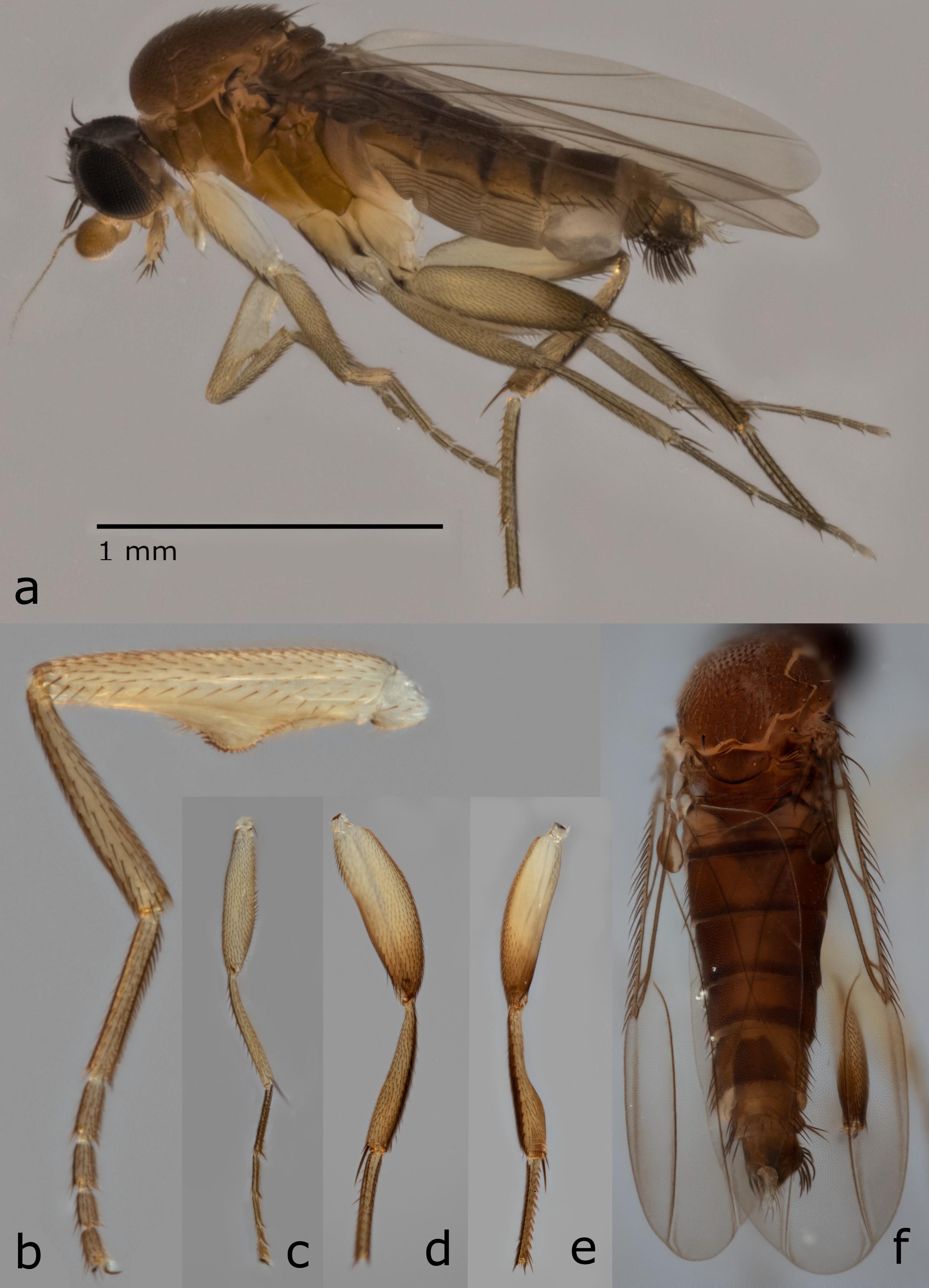
Lateral habitus (a) and diagnostic features of *Megaselia sepsioides* spec. nov. (a, inset) terminalia, (b) posterior view of foreleg, (c) anterior view of midleg (d,e) anterior and postero-dorsal views of hindleg, (e) dorsal view of thorax and abdomen.

**Figure 6:**
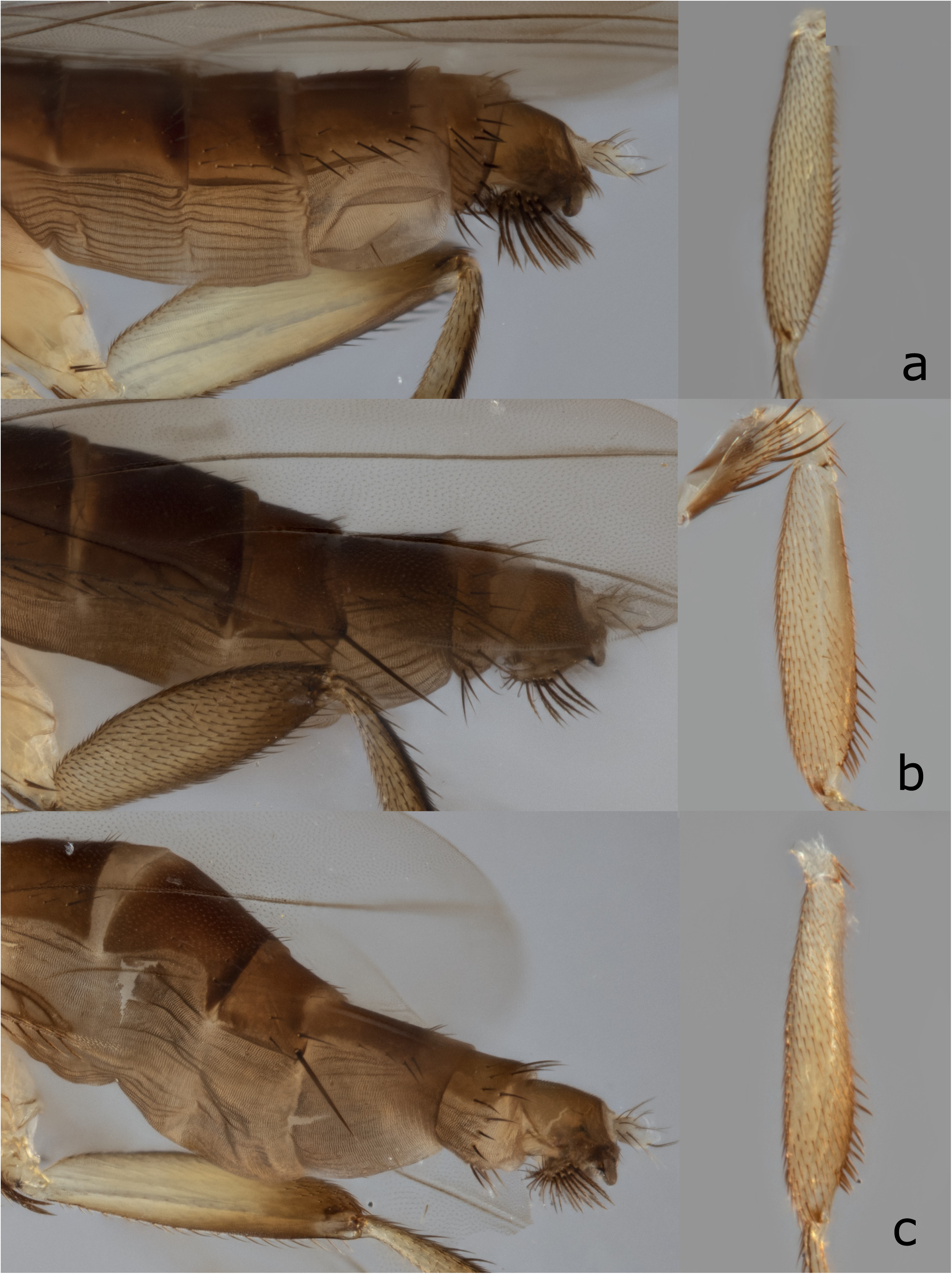
Haplotype variation of *Megaselia sepsioides* spec. nov. (a) UGC0005996, (b) UGC0012244, (c) UGC0012899. UGC numbers refer to specimen IDs.

**Figure 7:**
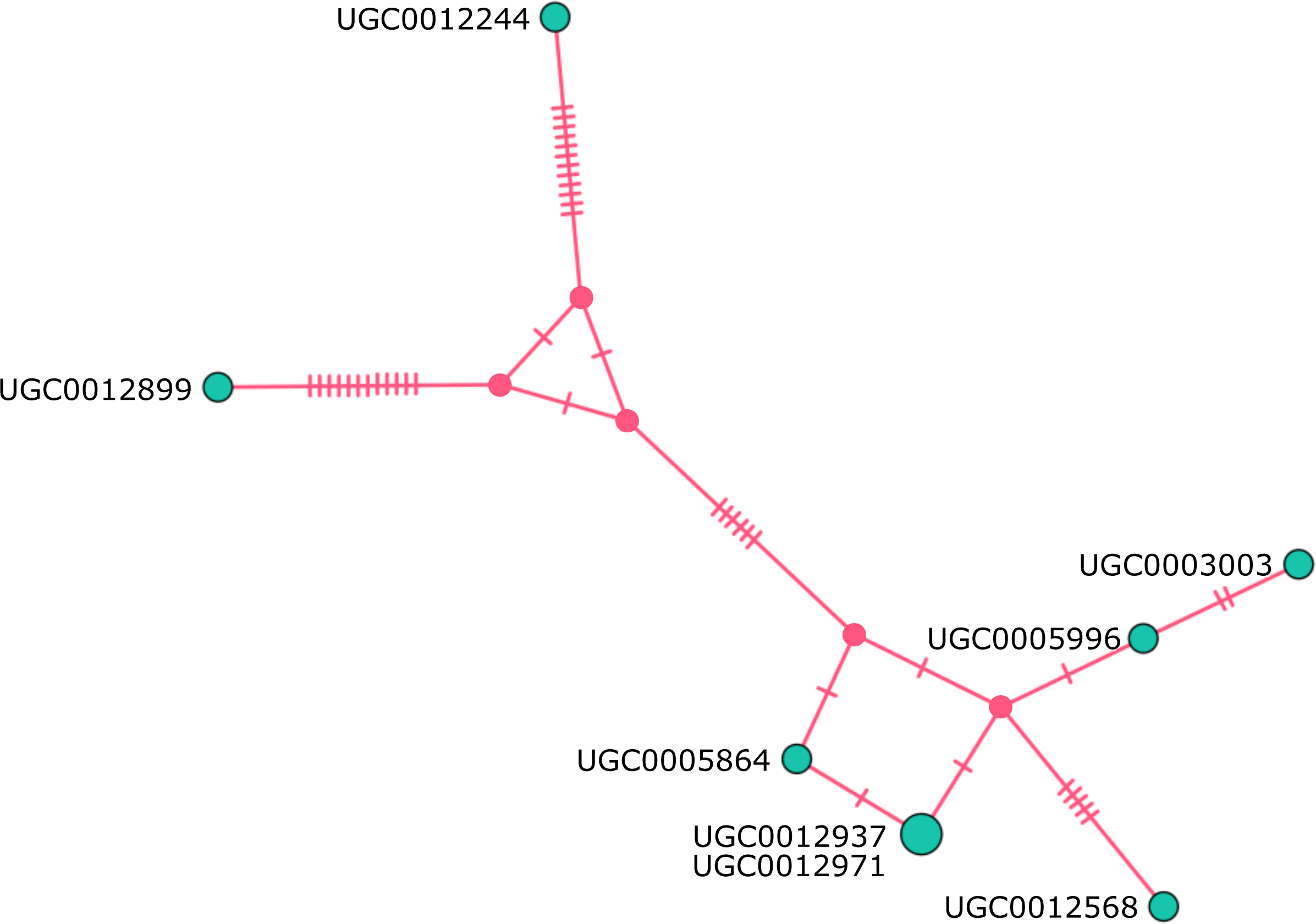
Haplotype network for *Megaselia sepsioides* spec. nov. UGC numbers refer to specimen IDs.

#### Diagnosis

Well characterized by the following combination of characters: with unique semi-circular expansion with modified peg-like setae on the forefemur (Fig. 5, b), hind tibia strongly constricted (Fig. 5, d and e), and abdomen narrow and elongate. Three haplotypes were examined; variations in setation were observed between the main cluster and two haplotypes (Fig. 6, 7). Only single specimens of the two distinct haplotypes are available; more specimens will be necessary to determine if these are eventually removed as distinct species or fall within a continuum of intraspecific variation.

#### Material Examined

Holotype. ♂, UGANDA: Kamwenge, Kibale National Park (00°33′54.2″N 30°21′31.3″E,1530 m), iii-xii.2010, Olavi Kurina & Swaibu Katusabe (LKCNHM UGC0005996).

Paratypes. 7 ♂, UGANDA: Kamwenge, Kibale National Park (00°33′54.2″N 30°21′31.3″E,1530 m), iii-xii.2010, Olavi Kurina & Swaibu Katusabe (LKCNHM: UGC0012899, UGC0012244, UGC0012568, UGC0003003, UGC0005864, UGC0012937, UGC0012971).

#### Distribution

Known from a single site in Kibale National Park, Uganda.

#### Biology

Unknown.

#### Etymology

Name suggested by Yuchen Ang for the sepsid-like (Diptera: Sepsidae) foreleg modification.

## DISCUSSION

### Remarkably high diversity of Phoridae in Kibale National Park

The full extent of the world’s biodiversity is poorly understood because many hyperdiverse taxa are data-deficient. One such hyperdiverse clade is phorid flies. We here reveal that even a modest amount of sampling (one Malaise trap placed in Kibale National Park, Uganda) can lead to the discovery of >650 putative species. This diversity constitutes 150% of the described phorid diversity of the entire Afrotropical region (466: [28]). Note that the ca. 7,000 barcoded specimens covered in our study only represent 8 one-week samples obtained between March 2010 and February 2011. There are an additional 44 weekly samples that remain un-sequenced. We thus expect the diversity from this single site to eventually exceed 1,000 species. This prediction is supported by a formal species-richness estimation based on the available data (Fig. 4). Such extreme diversity from a single site raises the question of whether these numbers are biologically plausible and/or whether they could be caused by unreliable data or species delimitation methods.

We would argue that they are both biological plausible and analytically sound. If a single garden in a temperate city like Cambridge (UK) can have 75 species and the urban backyards of Los Angeles 82, observing 10-15 times of this diversity at a site in a tropical National Park does not appear unrealistic. Our proposition that there are >1,000 species at a single site in Kibale National Park is further supported by the results of the Zurqui survey in Costa Rica which revealed 404 species of Phoridae without completing the species discovery process [26]. Furthermore, we are confident that our species richness estimate is not an artefact of poor data quality because the high species richness estimate is supported by both Illumina and MinION barcodes generated independently using different primer pairs. It is furthermore stable to modifications of sequence clustering thresholds which are widely used across Metazoa and here shown to be appropriate for phorid flies based on the available Genbank data [37]. Lastly, we checked 100 randomly chosen mOTUs for congruence between molecular and morphological evidence. We find that 93% of the clusters and >99% of specimens are congruently placed (six of the seven cases of incongruence involved single specimens). This is in line with congruence levels that we observed previously for ants and odonates [4, 7].

The high species richness found in one study site inspired us to speculate about the species diversity of phorids in the Afrotropical region. This is what Terry Erwin did when he famously estimated a tropical arthropod fauna of 30 million species based on his explorations of beetle diversity in Panama [38]. Such extrapolations are arguably useful because they raise new questions and inspire follow-up research. Speculation is inevitable given that it remains remarkably difficult to estimate the species richness of diverse taxa [39]. This is particularly so for the undersampled Afrotropical region which comprises roughly 2,000 squares of 100 km^2^ size. In our study, we only sampled a tiny area within one of these squares and observed >650 species which likely represent a species community that exceeds >1,000 species. Note that Malaise traps only sample some phorid species because many specialist species (e.g., termite inquilines) rarely enter such traps. In addition, the estimated 1,000 species that can be caught in a single Malaise trap are also only a subset of the species occurring in the remaining habitats of the same 100 km^2^ grid cell. Overall, it seems thus likely that the 100 km^2^ will be home to several thousand species of phorids. If we assume that on average each of the two-thousand 100 km^2^ grid cells of the Afrotropical Region has “only” 100 endemic phorid species, the endemic phorids alone would contribute 200,000 species of phorids to the Afrotropical fauna without even considering the contributions by the remaining species with a wider distribution. What is even more remarkable is that most of the diversity would belong to a single genus. We find that 90% of the newly discovered species and specimens in our sample belong to the genus *Megaselia* as currently circumscribed. Unless broken up, this genus could eventually have >100,000 Afrotropical species. All these estimates would only be lower if the vast majority of phorid species had very wide distributions and/or the average number of species in 100 km^2^ grid cells would be more than one order of magnitude lower than observed here. However, we consider this unlikely given that many areas of the Afrotropical region are biodiverse and cover a wide variety of climates and habitats which increases beta diversity.

Unfortunately, most of this diversity would have remained overlooked using the traditional taxonomic workflow because it is not well suited for taxa with high species diversity and specimen abundances. This means that the phorid specimens from the Kibale National Park Malaise trap would have remained in the unsorted residues for decades or centuries. Indeed, there are thousands of vials labelled “Phoridae” shelved in all major museums worldwide. Arguably, it is these unprocessed samples that make it so important to develop new rapid species-level sorting methods. We here favour sorting with “NGS barcodes” [4] because it would allow biologists to work through the Malaise trap residues in Natural History museums. These residues will almost certainly contain several times the number of species that have already been described. We predict that there will be two stages to species discovery with NGS barcodes. The first is species-level sorting which can yield fairly accurate estimates of species diversity and abundance [4, 8]. Many biodiversity-related questions can already be addressed based on these data. The second phase is the refinement of mOTUs based on morphological testing with subsequent species identification (described species) or species description (new species). Given the large number of new species, this will require optimized “turbo-taxonomic” methods. Fortunately, new approaches to large-scale species description are being developed and there are now a number of publications that describe ∼100 or more new species [36, 40–42].

### MinION sequencing and the “reverse workflow”

MinION barcodes can be obtained without having to invest heavily into sequencing facilities and laboratories fully equipped for barcoding specimens with MinION can be mobile and operate under difficult conditions in the field or field stations [15–18]. The technology is thus likely to become important for the “democratization” of biodiversity research because the data are generated quickly enough that they can be integrated into high school and citizen scientist initiatives. Based on our data, we would also argue that MinION is now suitable for wide-spread implementation of the “reverse-workflow” where all specimens are sequenced before mOTUs are assessed for consistency with morphology. Reverse-workflow differs from the traditional workflow in that it relies on DNA sequences for sorting all specimens into putative species while the traditional workflow starts with species-level sorting based on morphology; only some morpho-species are subsequently examined with a limited amount of barcoding. We would argue that the reverse-workflow is more suitable for handling species- and specimen-rich clades because it requires less time than high-quality sorting based on morphology which often involves genitalia preparations and slide-mounts. For example, even if we assume that an expert can sort and identify 50 specimens of unknown phorids per day, the reverse workflow pipeline would increase the species-level sorting rate by >10 times (based on the extraction and PCR of six microplates per day). In addition, the molecular sorting can be carried out by lab personnel trained in amplicon sequencing while accurate morpho-species sorting requires highly trained taxonomic experts. Yet, even highly trained experts are usually not able to match morphologically disparate males and females belonging to the same species (often one sex is ignored in morphological sorting) while the matching of sexes (and immatures) is an automatic and desirable by-product of applying the reverse workflow [7]. All these benefits can be reaped rapidly. One lab member can amplify the barcode of 600-1,000 specimens per day (2-2.5 weeks for 8,000 specimens). Obtaining DNA sequences requires ca. one week because it involves two cycles of pooling and sequencing on two MinION flowcells followed by two cycles of re-pooling and sequencing of weak amplicons. The bioinformatics work requires less than one week.

One key element of the reverse workflow is that the vials with specimens that have haplotype distances <5% are physically kept together. This helps when assessing congruence between mOTUs and morphology. Indeed, graphical representations of haplotype relationships (e.g., haplotype networks) can guide the morphological re-examination as illustrated in our description *of Megaselia sepsioides* (Fig. 7). Here, eight specimens belonging to seven haplotypes were dissected and the most dissimilar haplotypes were studied in order to test whether the data support the presence of one or two species. Variations in setation were observed (Fig. 6) but they were consistent with intraspecific variation. Note that the morphological examination of clusters was straightforward because the use of QuickExtract™ for DNA extractions ensures morphologically intact specimens.

### Large-scale species discovery using MinION 1D reads

Our results suggest that MinION’s 1D sequencing yields data of sufficient quality for producing DNA barcodes that can be used for large-scale species discovery. Through the development of new primer-tags, re-pooling of low coverage amplicons, and an improved bioinformatics workflow, we here increased the barcoding capacity of a MinION flowcell by 700% from 500 specimens (Srivathsan et al., [14]: 1D^2^ sequencing) to ∼3,500 specimens. This is achieved without a drop in accuracy because the error correction pipeline is effective at eliminating most of the errors in the 1D reads (ca. 10%). Indeed, even the initial estimates of barcodes (“MAFFT” & “RACON”) have very high accuracy (>99.5%) when compared to Illumina data, while the accuracy of consolidated barcodes is even higher (>99.9%). Note that the accuracy of the barcodes obtained in this study are even higher than what was obtained with 1D^2^ sequencing in Srivathsan et al. (99.2%) [14]. We suspect that this partially due to improvements in MinION sequencing chemistry and base-calling, but our upgraded bioinformatics pipeline also helps because it increases coverage for the amplicons. This is welcome news because 1D library preparations are much simpler than the library preps for 1D^2^. In addition, 1D^2^ reads are currently less suitable for amplicon sequencing [43]. Overall, the accuracy of our MinION barcodes are also comparable with what can be obtained with PacBio Sequel ([10]: Additional File 1: Figure S5). This platform also allows for obtaining full-length barcodes, but the cost of purchasing and maintaining a PacBio sequencer is very high.

Using the workflow described here, MinION barcodes can be generated rapidly and at a low sequencing cost of <0.35 USD per barcode. Molecular cost of PCR is 0.16 USD per reaction [4] while QuickExtract™ reagent costs 0.06 per specimen. These properties make MinION a valuable tool for species discovery whenever a few thousand specimens (<5,000) must be sorted to species. Even larger-scale barcoding projects are probably still best tackled with Illumina short-read or PacBio’s Sequel sequencing [4, 10, 11] because the barcoding cost is even lower. However, both require access to expensive sequencing instruments, sequencing is thus usually outsourced, and the users usually have to wait for several weeks in order to obtain the data. This is not the case for barcoding with MinION, where most of the data are collected within 10 hours of starting a sequencing run. Another advantage of the MinION pipeline is that it only requires basic molecular lab equipment including thermocyclers, a magnetic rack, a Qubit, a custom-built computational device for base-calling ONT data (“MinIT”), and a laptop (total cost of lab <USD 10,000). Arguably, the biggest operational issue is access to a sufficiently large number of thermocyclers given that a study of the scale described here involves amplifying PCR products in 92 microplates (=92 PCR runs).

Our new workflow for large-scale species discovery is based on sequencing the amplicons in two sequencing runs. The second sequencing run can re-use the flowcell that was used for the first run. Two runs are desirable because they improve overall barcoding success rates. The first run is used to identify those PCR products with “weak” signal (=low coverage). These weak products are then re-sequenced in the second run. This dual-run strategy overcomes the challenges related to sequencing large numbers of PCR products: the quality and quantity of DNA extracts are poorly controlled and PCR efficiency varies considerably. Pooling of products ideally requires normalization, but this is not practical when thousands of specimens are handled. Instead, one can use the real-time sequencing provided by MinION to determine coverage and then boost the coverage of low-coverage products by preparing and re-sequencing a separate library that contains only the low coverage samples. Given that library preparations only require <200 ng of DNA, even a pool of weak amplicons will contain sufficient DNA. This ability to re-pool within days of obtaining the first sequencing results is a key advantage of MinION. The same strategy could be pursued with Illumina and PacBio but it would take a long time to obtain all the results because one would have to wait for the completion of two consecutive runs.

## METHODS

### 1. Sampling

Samples were collected by a single Townes-type Malaise trap [44], in the Kibale National Park, close to Kanyawara Biological Station in the evergreen primeval forest at an altitude of 1513 m (00°33′54.2″N 30°21′31.3″E) (Fig. 4). Kibale National Park is characterized as a fragment of submontane equatorial woodland [45]. Temperatures in Kibale range from 16°C to 23°C (annual mean daily minimum and maximum, respectively) [46]. The Malaise trap was checked every week when the collecting bottle with the material was replaced by a resident parataxonomist ([47]: Mr. Swaibu Katusabe). Subsequently, the material was collected and transferred in accordance with approvals from the Uganda Wildlife Authority (UWA/FOD/33/02) and Uganda National Council for Science and Technology (NS 290/ September 8, 2011), respectively. The material was thereafter sorted to higher-level taxa. Target taxa belonging to Diptera were sorted to family and we here used the phorid fraction. The sampling was done over several months between 2010 and 2011. For the study carried out here, we only barcoded ca. 30% of the phorid specimens. The flies were stored in ethyl alcohol at -20-25°C until extraction.

### 2. DNA extraction

DNA was extracted using whole flies. The fly first taken out from the vial and washed in Milli-Q^®^ water prior to being placed in a well of a 96 well PCR plate. DNA extraction was done using 10 ul of QuickExtract™ (Lucigen) in a 96 well plate format and the whole fly was used to extract DNA. The reagent allows for rapid DNA extraction by incubation (no centrifugation or columns are required). The solution with the fly was incubated at 65°C for 15 min followed by 98°C for 2 min. No homogenization was carried to ensure that the intact specimen was available for morphological examination.

### 3. MinION based DNA barcoding

#### I. Polymerase Chain Reactions (PCRs)

Each plate with 96 QuickExtract™ extracts (95 specimens and 1 control, with exception of one plate with no negative and one partial plate) was subjected to PCR in order to amplify the 658 bp fragment of COI using LCO1490 5’ GGTCAACAAATCATAAAGATATTGG 3’ and HCO2198 5’ TAAACTTCAGGGTGACCAAAAAATCA 3’ [48]. This primer pair has had high PCR success rates for flies in our previous study [14] and hence was chosen for phorid flies. Each PCR product was amplified using primers that included a 13 bp tag. For this study, 96 thirteen-bp tags were newly generated in order to allow for upscaling of barcoding; these tags allow for multiplexing >9200 products in a single flowcell of MinION through unique tag combinations (96×96 combinations). To obtain these 96 tags, we first generated one thousand tags that differed by at least 6 bp using BarcodeGenerator [49] However, tag distances of >6 bp are not sufficiently distinct because they do not take into account MinION’s propensity for creating errors in homopolymer stretches and other indel errors. We thus excluded tags with homopolymeric stretches that were >2 bp long. We next used a custom script to identify tags that differed from each other by indel errors. Such tags were eliminated recursively to ensure that the final sets of tags differed from each other by >=3bp errors of any type (any combination of insertions/deletions/substitutions). This procedure yielded a tag set with the edit-distance distribution shown in Additional File 1: Fig S1 [minimum edit distance (as calculated by *stringdist* module in Python) of 5 nucleotides (38.5%) which is much higher than Nanopore error rates]. Lastly, we excluded tags that ended with “GG” because LCO1490 starts with this motif. Note that longer tags would allow for higher demultiplexing rates, but our preliminary results on PCR success rates suggested that the use of long tags reduced amplification success (one plate: 7% drop).

The PCR conditions for all amplifications were as follows, reaction mix: 10 µl Mastermix (CWBio), 0.16 µl of 25mM MgCl_2_, 2 µl of 1 mg/ml BSA, 1 µl each of 10 µM primers, and 1ul of DNA. The PCR conditions were 5 min initial denaturation at 94°C followed by 35 cycles of denaturation at 94°C (30 sec), annealing at 45°C (1 min), extension at 72°C (1 min), followed by final extension of 72°C (5 min). For each plate, a subset of 7-12 products were run on a 2% agarose gel to ensure that PCRs were successful. Of the 96 plates studied, 4 plates were excluded from further analyses as they had <50% amplification success and one plate was inadvertently duplicated across the two runs.

#### II. MinION sequencing

We developed an optimized strategy for nanopore sequencing during the study. For the initial experiment (set 1), we sequenced amplicons for 4,275 phorid flies. For this, all plates were grouped by amplicon strength as judged by the intensity of products on agarose gels and pooled accordingly (5 strong pools + 2 weak pools). The pools were cleaned using either 1X Ampure beads (Beckman Coulter) or 1.1X Sera-Mag beads (GE Healthcare Life Sciences) in PEG and quantified prior to library preparation. The flowcell sequenced for 48 hours and yielded barcodes for ∼3200 products, but we noticed lack of data for products for which amplification bands could be observed on the agarose gel. We thus re-pooled products with low coverage (<=50X), prepared a new library and sequenced them on a new flowcell. The experiment was successful. However, in order reduce sequencing cost and improve initial success rates, we pursued a different strategy for the second set of specimens (4,519 specimens). In the first sequencing run, we stopped the sequencing after 24 hours. The flowcell was then washed using ONT’s flowcell wash kit and prepared for reuse. The results from the first 24 hours of sequencing were then used to identify amplicons with weak coverage. They were re-pooled, a second library was prepared, and sequenced on the pre-used and washed flowcell.

Pooling of weak products with <=50X coverage was done as follows: We located (1) specimens <=10X coverage (set 1: 1,054, set 2: 1,054) and (2) specimens with coverage between 10X and 50X (set 1: 1,118, set 2: 1,065). Lastly, we also created a (3) third pool of specimens with “problematic” products that were defined as those that were found to be of low accuracy during comparisons with Illumina barcodes and those that had high levels of ambiguous bases (>1% ambiguous bases during preliminary barcode calling). Very few amplicons belonged to this category (set 1: 68, set 2: 92). In order to efficiently re-pool hundreds of specimens across plates we wrote a script that generates visual maps of the microplates that illustrate the wells where the weak products are found (available in github repository for *miniBarcoder*).

Library preparation and sequencing: We used the SQK-LSK109 ligation sequencing kit (Oxford Nanopore Technologies) for library preparation and sequencing. Our first experiment on set 1 used 1 ug of starting DNA while all other libraries used 200 ng pooled product. Library preparation was carried out as per manufacturer’s instructions with one exception: the various clean-up procedures at the end-prep and ligation stages used 1X Ampure beads (Beckmann Coulter) instead of 0.4 X as suggested in the instructions because the amplicons in our experiments were short (∼735 bp with primers and tags). The sequencing was carried out using MinION sequencer with varying MinKNOW versions between August 2018 - January 2019. Fast5 files generated were uploaded onto a computer server and base-calling was carried out using Guppy 2.3.5+53a111f. No quality filtering criteria were used. Our initial work with Albacore suggested that quality filtering improved demultiplexing rate but overall more reads could be demultiplexed without the filtering criterion.

#### III. Data analyses for MinION barcoding

We attempted to demultiplex the data for set 1 using *minibar* [50], however it was found to demultiplex only 1,039 barcodes for the 4275 specimens (command used: python ../minibar/minibar.py -F -C -e 2 minibar_demfile_1 phorid_run1ab.fa_overlen599 > out). This success rate was so low that we discontinued the use of *minibar*. Instead, we analysed the data using an improved version of *miniBarcoder* [14]. This pipeline starts with a primer search with *glsearch36,* followed by identifying the “sequence tags” in the flanking nucleotide sequences, before the reads are demultiplexed based on tags. For the latter, errors of up to 2 bp are allowed. These “erroneous” tags are generated by “mutating” the original tags 96 tags to account for all possible insertions/deletions/substitutions. The sequence tags are matched with this set of input tags + tag mutants. This speeds up demultiplexing as it does not have to align each tag. When comparing the performance of *miniBarcoder* with *minibar*, we found that using 4 cores, *miniBarcoder* could demultiplex data in <10 minutes while *minibar* required 74 minutes in one. Both pipelines demultiplexed similar numbers of reads: 898,979 reads using *miniBarcoder*, while 940,568 HH reads of *minibar* (56,648 reads in multiple samples). The demultiplexed reads were aligned using MAFFT v7 (--op 0) (here v7) [51]. In order to improve speed, we only used a random subset of 100 reads from each demultiplexed file for alignment. Based on these alignments, a majority rule consensus was called to obtain what we call “MAFFT barcodes”.

Other studies usually incorporate a step of clustering of data at low thresholds (e.g. 70% by Maestri et al. 2019 [18]) in order to account for the read errors produced by MinION. The subsequent analysis is then carried out on the cluster that has the largest number of reads. We deviate from this approach because it requires high coverage. In our barcoding pipeline, we assess congruence for each base-pair by firstly eliminating the MAFFT gap-opening penalty (-- op 0); this allows for all data to be used when calling the consensus [14]: this gap opening penalty essentially treats indel and substitutions similarly and staggers the alignment. Any base that is found in <50% of the position is called as an ambiguity; i.e., the majority-rule criterion is applied for each site instead of filtering at the read level which is based on averages across all bases. This site-specific approach maximizes data use by allowing barcodes to be called at much lower coverage levels than with other pipelines (5-20X coverage compared to >200x coverage in Maestri et al. 2019 [18]), while also identifying contaminated specimen barcodes.

In our pipeline, MAFFT barcodes are improved by mapping all the reads back to the barcode using GraphMap (here v0.5.2) [52] and calling the consensus using RACON (here, v1.3.1) [31]. This yields what we call “RACON barcodes”. Both MAFFT and RACON barcodes are subject to further correction based on publicly available barcodes in GenBank. These corrections are advisable in order to fix the remaining indel errors. The correction takes advantage of the fact that COI sequences are translatable; i.e., an amino acid-based error correction pipeline can be used (details can be found in Srivathsan et al. [14]). Applying this pipeline to MAFFT and RACON barcodes, respectively, yields MAFFT+AA and RACON+AA barcodes. Lastly, these barcodes can be consolidated into “consolidated barcodes”.

The version of the pipeline in Srivathsan et al. [14] was modified as follows:

a. *Tackling 1D reads for massively multiplexed data:* We developed ways for correcting for the increased number of errors in 1D reads by identifying objective ways for quality assessments based on the MinION data and publicly available data (GenBank): (1) The GraphMap max error was increased from 0.05 to 0.15 to account for error rates of 1D reads. (2) We modified the approach for calculating consolidated barcodes. These consolidated barcodes are generated by aligning translatable MAFFT and RACON barcodes. We then use the strict consensus of MAFFT+AA and RACON+AA barcodes in order to resolve conflicts between MAFFT and RACON barcodes if there are substitution conflicts. In Srivathsan et al. [14] we accepted MAFFT+AA barcodes in cases of conflict, but for the 1D data we found that MAFFT+AA barcodes had more ambiguities than RACON+AA barcodes which could be resolved via calculating a strict consensus. We increased the gap-open penalty from default 1.53 to 3, as the MAFFT+AA and RACON+AA barcodes are translatable and their alignments should not contain indels. Lastly if the alignment still contains indels despite the increase in gap opening penalty the barcode is rejected. (3) We assessed how different window sizes can impact the amino-acid correction pipeline by varying the “namino” parameter (number of AA to be inspected in each side of an indel). (4) During the amino-acid correction, we introduced a final sequence translation check to ensure that a translatable product was obtained: if stop codons were still present these would be replaced by ambiguities. These affected four barcodes in our study.
b. *Demultiplexing rate*: (1) We introduced a “homopolymer compression” of the putative tag sequences in order to improve demultiplexing rates. After primer searches, the old pipeline used to identify the flanking 13 bps that were likely to be tag sequence. Instead we now use a 20 bp flanking sequence and then compress any homopolymer >3bp before searching for 13 bp tags. (2) We now analyze very long reads that can be formed when two amplicons ligate during library preparation. Long reads are split into size classes of <1,300, >1,300 and <2,000. These settings were set based on 658 bp barcode of COI: the total product size including tags and primers is 735 bp, and hence, a sequence with two products ligated to each other is expected to be 1,470 bp long. The sequences were split in a manner that ensured that the tags of first product are not affecting the tag found in the second, i.e., primer search for second product was conducted after the first 650 bp of the sequence. Currently, this option is only available for reads that consist of two ligated products.
c. *Processing speed and memory requirements*: (1) For primer identification we now limit the searches to the first and last 100 bp of each read which allowed for improving speed of the primer search. (2) We parallelized the process of demultiplexing and MAFFT barcode calling using multiprocessing in python. This allows for fast demultiplexing whenever computational power is limited. (3) We optimized the pipeline with regard to memory consumption and ensured that all processes are applied to batches of <=20,000 sequences. The latter modification is needed given the rapid increase in MinION output and batch processing is scalable with increased output.

### 4. Assessment of demultiplexing accuracy

Demultiplexing accuracy was assessed by trying to demultiplex reads into bins representing unused tag combinations: 96×96 tags combinations were designed but set 1 used only 45 tags with LCO1490. This allowed for an assessment if *miniBarcoder* would erroneously assign reads for the unused remaining 51X96 combinations. Set 2 used 48 tags thus allowing us to assess if the demultiplexing pipeline erroneously found the remaining 48X96. Lastly one plate used only 55 tags associated with HCO2198. Overall, we could assess demultiplexing accuracy associated with 80/96 tags.

### 5. Illumina Based NGS Barcoding for validation

In order to validate MinION barcodes and optimize error correction strategies, we used reference COI barcodes obtained via Illumina sequencing for the same QuickExtract™ DNA extractions. Illumina sequencing was carried out for a 313 bp fragment of the same COI barcoding region using m1COlintF: 5’-GGWACWGGWTGAACWGTWTAYCCYCC-3’ [53] and modified jgHCO2198: 50-TANACYTCNGGRTGNCCRAARAAYCA-3 [54]. We recently conducted an extensive analysis to understand if a 313 bp minibarcode is able to provide similar identifications and species delimitations as 658 bp barcodes and found that minibarcodes performing equally well as full-length barcodes when examining >5,000 species [55].

Tagged primers were used for the PCRs as specified in Wang et al. (2018) [4]. The PCR mix was as follows: 4 µl Mastermix from CWBio, 1µl of 1 mg/ml BSA, 1µl of 10 µM of each primer, and 1ul of DNA. PCR conditions: 5 min initial denaturation at 94°C followed by 35 cycles of denaturation at 94°C (1 min), 47°C (2 min), 72°C (1 min), followed by final extension of 72°C (5 min). The PCR products were pooled and sequenced along with thousands of other specimens in a lane of HiSeq 2500 (250 bp PE sequencing). The data processing followed Wang et al. (2018) [4]: paired end reads were merged using PEAR (v 0.9.6) [56], reads were demultiplexed by an in-house pipeline that looks for perfect tags while allowing for 2 bp mismatches in primer sequence. For each sample, demultiplexed reads are merged to identify the most dominant sequence with >50X dataset coverage and 10X coverage for the sequence, and a barcode is accepted only if this sequence is 5X as common as the next most common sequence. Demultiplexed reads with >10x coverage were also per sample were retained for analyses of contaminations and paralogs.

### 6. Assessment of MinION barcodes and mOTU delimitations

Both MinION and Illumina barcodes were subject to contamination check. For MinION barcodes we used preliminary MAFFT barcodes given that this is the largest barcode set. Barcodes were matched to GenBank using BLASTN and taxonomic classifications were assigned using readsidentifier [57]. Any barcode with >95% match to a non-phorid sequence was excluded from the dataset. Furthermore, if any barcode best matched to bacteria, it was also excluded. Lastly, in cases where the best BLAST matches were below 95% and no phorid sequence was in the top 100 BLAST hits, we retrieved the specimen and examined in morphologically to determine if a non-phorid had been sequenced (=pre-sorting error).

MinION barcodes were assessed by scripts provided in the *miniBarcoder* package (assess_corrected_barcodes.py and assess_uncorrected_barcodes.py). For uncorrected barcodes, this was done by aligning barcodes to reference Illumina barcodes using dnadiff [58]. For corrected barcodes (+AA), we used MAFFT to obtain pairwise alignments. This allowed us to compare per base accuracy. Here we excluded those barcodes that had >3% difference between MinION and Illumina barcodes. These are likely to be caused by wetlab error whenever a different product as the Illumina and MinION barcodes were amplified using different primers (<0.5% of specimens) as discussed here: for consolidated barcodes 25/6,251 specimens are involved in such >3% distances in the comparisons of consolidated barcodes with Illumina barcodes. If these distances were a result of consensus errors, sequences are unlikely to create consensus that are identical to other specimens. 21/25 specimens have identical sequences to some other specimen and two are within 0.6% of another. By matching MinION barcodes to Illumina’s read set for each sample (>=10 counts), we also find that many of the signals being picked up by HCO-LCO are in the Illumina data at low frequencies (in 15/25 cases, including one of the two cases that remain unaccounted) that was generated using different primers (Additional File 5). We do not include these for consensus quality, but instead consider them as misplaced sequences.

We were furthermore interested in understanding how mOTU delimitation is affected by error correction. Barcodes were aligned using MAFFT and MinION barcodes were further trimmed to the 313 bp region of the Illumina barcode. mOTU delimitation was done at 2, 3 and 4% using SpeciesIdentifier (objective clustering) [59]. mOTU richness was estimated and we furthermore calculated match ratio between two sets of clusters [60]. Match ratio is given by 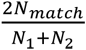. For consolidated barcodes, we also calculated the match ratio between mOTU estimates by full length barcodes and Illumina barcodes (313 bp). If the mOTUs were inconsistent we also identified the distance threshold for which they became consistent.

### 7. Paralog check

Copies of mitochondrial sequences in nuclear genomes (NuMTs, paralogs) can lead to inflated species richness estimation if the mitochondrial gene amplifies in one specimens and the NuMT copy in another specimen that is conspecific. If the sequence copies are sufficiently dissimilar, the two specimens will be placed in two mOTUs although they were conspecific. In our study, multiple checks are used to avoid inflated diversity estimates based on NuMTs. Firstly, we amplify COI sequences of different lengths using two different primer pairs and then check whether the resulting mOTUs are congruent. Congruence would not be expected if the primers have similar affinity to the mitochondrial and nuclear copy of the gene. Secondly, the Illumina read sets were obtained for an amplicon generated with highly degenerate primers. This increases the chance for co-amplification of paralogs. For each demultiplexed set, we obtained the unique sequences that had >=10X coverage. All secondary were matched against the barcode sequences using BLASTN, e-value 1e-5, perc_identity 99%, hit length >=300. If secondary signals for a specimen belonged to a different specimen these could be low level contaminations or paralogs. However, if secondary sequences from multiple specimens of one cluster match to a different cluster, the likelihood that they are paralogs is higher. Note, however, that this technique does not allow for distinguishing between low-level contamination and paralogs.

### 8. Species richness estimation

The most accurate set of barcodes were used for assessing the overall diversity of the barcoded phorids. Distance based mOTU delimitation was based on SpeciesIdentifier [59] while tree-based mOTU delimitation was conducted using Poisson Tree Processes (PTP) model [61]. The number of species was estimated using three different thresholds (2%, 3%, 4%). To understand if these thresholds are not misleading for Phoridae, we examined the publicly available data in GenBank for this family. We downloaded 79,506 phorid COI sequences to from NCBI using the following search terms: txid36164[Organism] AND (COI[Gene Name] OR Cox1[Gene Name] OR COXI[Gene Name] OR “cytochrome oxidase subunit 1”[Gene Name] OR “cytochrome c oxidase subunit 1”[Gene Name] OR “cytochrome oxidase subunit I”[Gene Name]). The data were filtered to exclude unidentified barcode records (containing sp. or identified as Phoridae BOLD) and sequences that had <500 bp overlap with the 658 bp COI 5’ barcoding region. Four additional sequences were excluded as they resulted in a frameshift in the sequence alignment conducted using MAFFT. The final alignment contained 12,072 DNA barcodes corresponding to 106 species, 74 of which had multiple barcodes. The barcodes corresponding to these 74 species were used to assess the distribution of pairwise intraspecific genetic distances. All 12,072 barcodes were clustered at various distance thresholds (1-10%) and the congruence with morphology was assessed using match ratios. Congruence was highest for a 3% threshold which was then also used to for estimating the species richness in the Ugandan sample of phorids. The species richness estimation was carried out with EstimateS9 [62] using the classical formula of Chao1 given that the coefficient of variation of the abundance or incidence distribution was >0.5.

For tree based species delimitation, the PTP model [61] was applied on a maximum likelihood phylogeny built using the aligned haplotypes of consolidated barcode dataset and the GTRGAMMA model in RaXML v 8.4.2 [63]. Twenty independent searches were conducted to obtain the best scoring phylogeny. PTP model was then applied on the resulting best scoring ML phylogeny (mPTP --single --ML) [64].

### 9. Morphological examination

For morphological examination of the clustered specimens we used 100 randomly selected non-singleton mOTUs delimited at 5% but also kept track of sub-clusters within the large mOTUs that were splitting at 1-5%. This allowed for examination of closely related, but potentially distinct, species. We were mostly interested in understanding if MinION barcodes were placing specimens into mOTUs incorrectly and hence examined if the specimens were consistent morphologically in each of these 5% clusters. The choice of 5% cluster may seem initially inconsistent with the choice of 3% for mOTU delimitation for other analyses, but examination of all specimens within 5% clusters allows for comparing multiple closely related 3% (or 1-5%) mOTUs. For the strictest evaluation, this would often require genitalia preparations, but for reasons of scope, this was here only carried out for one species. For this species we illustrate how the haplotype network obtained with the median joining method in PopART (Fig. 7) [65] guides the morphological examination of specimens for the new species, *Megaselia sepsioides* sp. nov.. Specimens with the most dissimilar haplotypes were dissected in order to rule out the presence of multiple closely related species. Differences in setation were observed between the two distant haplotypes (UGC0012899 and UGC0012244) and the main cluster (UGC0003003, UGC0005864, UGC0005996, UGC0012568, UGC0012937, and UGC0012971) and are illustrated in Fig. 6. Specimen examination was done with a Leica m80 and Leica M205 C stereo microscopes and the images were obtained with a Dun Inc. Microscope Macrophotography system (Canon 7D chassis with 10X Mitutoyo lens). Photography stacking was done with Zerene stacker. Specimens were cleared with clove oil and mounted on slides in Canada balsam following the protocol of Disney [66]. The type series is housed in the Lee Kong Chian Natural History Museum, Singapore.

## List of abbreviations

mOTU: molecular Operational Taxonomic Units
NGS: Next Generation Sequencing
NuMTs: Nuclear mitochondrial DNA sequences
BIN: Barcode Index Number
PTP: Poisson Tree Processes
MSA: Multiple Sequence Alignment

## DECLARATIONS

### Ethics approval and consent to participation

The specimens were collected with valid permits (see acknowledgements).

### Consent for publication

Not applicable

### Availability of data and materials

#### Source code

- Project name: *miniBarcoder*
- Project home page: https://github.com/asrivathsan/miniBarcoder [67]
- Operating system(s): Linux, MacOSX
- Programming language: Python
- Other requirements: MAFFT BARCODE: MAFFT, glsearch36, RACON BARCODE: GraphMap, Racon, and amino acid correction: BioPython, BLAST and MAFFT.
- License: GNU GPL
- New scripts included: mb_parallel_consensus.py mb_parallel_demultiplex.py, repool_by_plate.py
- Updated scripts: consolidate.py, aacorrection.py

#### Others

Nanopore raw reads are available in NCBI SRA, project PRJNA563237 (SRR10054347-SRR10054350) [68] and the consolidated barcodes are available in FigShare [69] and GenBank (Accession Nos. MN403320-MN410421). The Additional File 6 contains the information for demultiplexing the data.

### Author’s Contributions

R.M. and A.S. conceived the workflow and the analytical approach. O.K. conducted/organized the sampling and Diptera sorting and commented on the manuscript. Molecular work was conducted by J.P., W.T.L., S.N.K., E.H. and A.S. Pipeline development and data analyses was conducted by A.S. Morphological examination, and species description was conducted by E.H. Figures were prepared by E.H., S.N.K., A.S. Manuscript was written by R.M., A.S. and E.H..

## Acknowledgements

The material in Kibale National Park in Uganda was collected and transferred in accordance with approvals from the Uganda Wildlife Authority (UWA/FOD/33/02) and Uganda National Council for Science and Technology (NS 290/ September 8, 2011), respectively. Yuchen Ang is thanked for his help photographing the new species and for his suggestion of sepsioides as the specific epithet. Henry Disney and Brian Brown are thanked for their consultation on the new species of *Megaselia*. We thank Arina Adom for help in curating the Illumina barcodes and Sabrina Tang for help with photography.

## ADDITIONAL FILES

Additional File 1.docx: Fig S1: Pairwise edit distances between tags. Fig S2 and S3: Analyses of GenBank phorid data

Additional File 2.xlsx: Tag sequences used in this study

Additional File 3.xlsx: Details on incongruent clusters

Additional File 4.xlsx: Analyses of paralogs/contaminations

Additional File 5.xlsx: Details on specimens with >3% divergence from Illumina barcodes for final consolidated barcode sets

Additional File 6.xlsx: Demultiplexing information for the two sample sets

