## Supplementary Information for "Rapid, large-scale species discovery in hyperdiverse taxa using 1D MinION sequencing" (from another version of this preprint: version 4, 2019-09-06)

Additional File 1

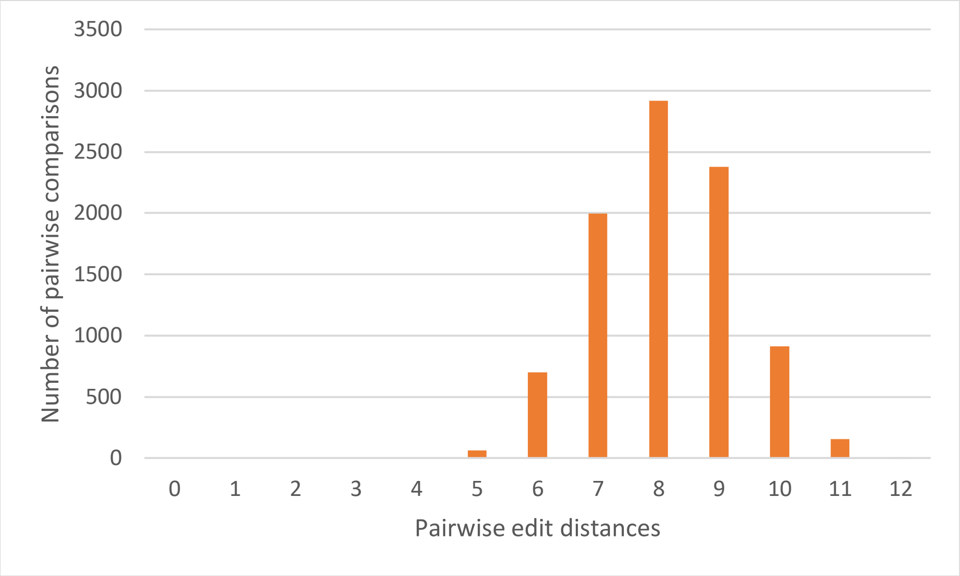

Figure S1: Pairwise Levenshtein distances between designed tags.

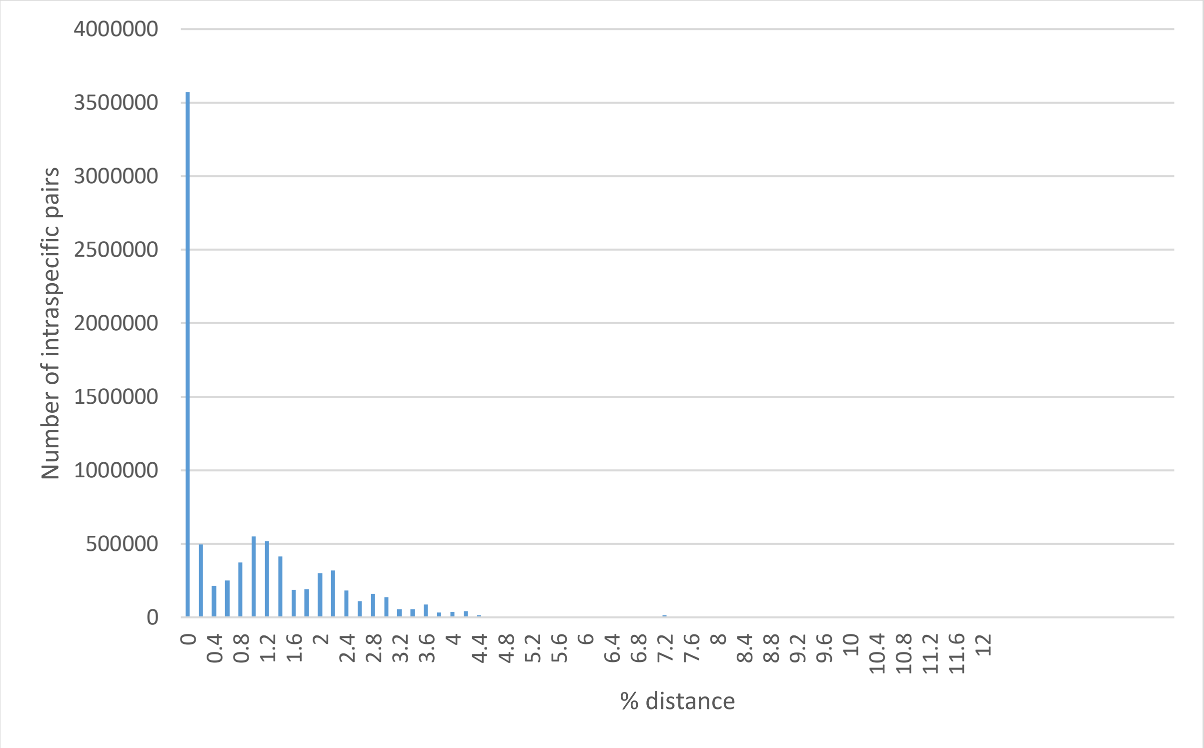

Figure S2: Distribution of intraspecific pairwise distances for publically available barcodes in GenBank.

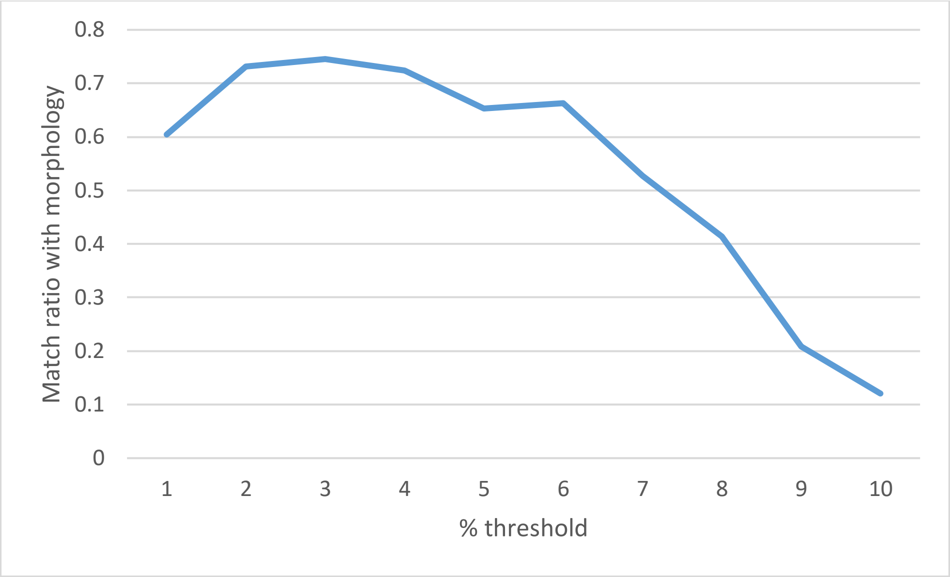

Figure S3: Match ratio between mOTUs at various thresholds and morphology for publically available barcodes from GenBank. Overall, highest match ratio is found for 3%.

| **Additional File 2: Tags used in the experiment with four alternatives.** |
|---|
| ATCCGGTCGGAGA |
| CTGAGGTGATCAG |
| AGTGTCCTGCTAG |
| ATAAGCAATTCGA |
| ATCTAACAGGACC |
| CATTATATAGCCA |
| CCTGATTACGTAA |
| CTAATACACACCG |
| TTATGACCGTTGC |
| TTCTCGGCCGCAT |
| TTGGCGGTAAGAA |
| ACTTGTGCACCTG |
| CATGCCTTGATAC |
| CTGACGTTCGGTA |
| GAGGAGAGACACA |
| GCCGGTCCAAGTG |
| TTGCGTCTCACGC |
| TTATTCGCACACT |
| TGCACAGGCGGCA |
| AACATGTGGTAAG |
| ACGCGCTCTTATA |
| CAATGTTAATGGT |
| CTAGGATATAGTT |
| CTGTAGGCGAGGA |
| CTGTCGAGGCGAC |
| GACTAATGAGGAA |
| TATCCGTAATCAT |
| TCCTGGAGGATAA |
| TCTACTGTTGTGC |
| TGTATATTCAGCG |
| ACATCAGTAGTTC |
| AGCGCTGGACCAA |
| CAAGATCGGTACC |
| CCAGCCTGACGGT |
| CCGAAGCCGGTTC |
| CCGAGCAGCTGTT |
| CCTCCGCACAATC |
| CTCTAACCTATTA |
| GCACCTTGACATA |
| TAAGCGTGCTGAA |
| TTCGGCACAGGAG |
| AGACTCGATTGAG |
| ATCCTTCTGTTCT |
| CGTTACTTAGGTC |
| CTGGTTGGCTATC |
| GCACCAACGATTG |
| TCTTATGCATAAC |
| TTAGGTGTGGACC |
| TTCTTCTGGAACA |
| AGCAAGTGGCTCT |
| TCAAGTCACTAAT |

| |
|---|
| ATGAAGACTACTG |
| CACCGGTAGAACC |
| CAGAACAACGCAT |
| CCTGCCGAGAACG |
| GTGATGGAGCGGT |
| TACGAACTATAGC |
| TACGGACCGCTCA |
| TGGACGCTAGATC |
| AGTCTTCACCTCC |
| AGTGCTTATCCGT |
| ATCGCTGCCTGGC |
| ATGCTTGCGACAG |
| CAATCCTCAAGAG |
| CATGCACACCTCT |
| CGGAGAATATTAT |
| GAGCGAAGGTAGT |
| GGCGACTGTTCCT |
| GTCAGTTAACGAC |
| TCCACAGATACAG |
| TCGAGTTGTAACT |
| TCGGTCTTAGACG |
| TCTAAGGAGTTAT |
| TCTATCTCTCGCT |
| TGTGAAGTTGCCA |
| TTACCATTCCTCA |
| AGATTCTACACAA |
| ATGCGATTAATTG |
| CCGTAGTGTTGAT |
| CCTGTACCATTGT |
| GGCTGTTACAACA |
| GGTTCAATGACGC |
| GGTTCCTGCACTT |
| GTCCAACTTCAGT |
| GTCCAGGCCTTCT |
| GTGACGCCGTATT |
| TGCAGCAACCAGC |
| TGGACATGGCATT |
| TGGTGGTCATTCA |
| TTACCAGAGCCTG |
| TTCCGCGGCATAG |
| ACAACCATTGGCA |
| AGCCTAGCCAACT |
| ATAATCGAGGCTT |
| CTGAGTCAGAAGT |
| GATAGGCGCCTTG |
| **GCGACAAGCACCG** |
| **GCTCCTACTGCCA** |
| **GTAGCCTATGGCG** |
| **GTTAGACAATACT** |

**Additional File 3: Incongruent clusters at 3% (MinION full length vs Illumina, and MinION 313 vs Illumina), but congruent at a different varying thresholds**

| % threshold values provided | | | Members of the clusters |
|---|---|---|---|
| Minion full | Minion 313 | Illumina 313 | |
| 2.4 | 3.1 | 3.1 | ('UGC0008455', 'UGC0010557') |
| 2.4 | 3.4 | 3.4 | ('UGC0004136', 'UGC0005900', 'UGC0006285', 'UGC0006329', 'UGC0008947', 'UGC0013192') |
| 2.8 | 3.1 | 3.4 | ('UGC0002514', 'UGC0003417', 'UGC0004496', 'UGC0004839', 'UGC0005028', 'UGC0005839', 'UGC0006379', 'UGC0006971', 'UGC0006975', 'UGC0008324', 'UGC0010270', 'UGC0010761', 'UGC0010835', 'UGC0012183', 'UGC0012555', 'UGC0012874', 'UGC0012969') |
| 3 | 3.2 | 3.4 | ('UGC0002121', 'UGC0002140', 'UGC0002159', 'UGC0002369', 'UGC0002545', 'UGC0003010', 'UGC0003306', 'UGC0003545', 'UGC0003593', 'UGC0004140', 'UGC0004177', 'UGC0004816', 'UGC0004841', 'UGC0005009', 'UGC0005058', 'UGC0005063', 'UGC0005361', 'UGC0005374', 'UGC0005557', 'UGC0005567', 'UGC0005570', 'UGC0005576', 'UGC0005665', 'UGC0005687', 'UGC0005710', 'UGC0005718', 'UGC0005722', 'UGC0005724', 'UGC0005729', 'UGC0005735', 'UGC0005756', 'UGC0005941', 'UGC0006107', 'UGC0006317', 'UGC0006357', 'UGC0006407', 'UGC0006472', 'UGC0006544', 'UGC0006568', 'UGC0006600', 'UGC0006743', 'UGC0006781', 'UGC0006890', 'UGC0007043', 'UGC0007051', 'UGC0007084', 'UGC0007097', 'UGC0007220', 'UGC0007262', 'UGC0007280', 'UGC0007283', 'UGC0007291', 'UGC0007306', 'UGC0007319', 'UGC0007349', 'UGC0007352', 'UGC0007388', 'UGC0007460', 'UGC0007491', 'UGC0007499', 'UGC0007504', 'UGC0007517', 'UGC0007668', 'UGC0007679', 'UGC0007687', 'UGC0008478', 'UGC0008644', 'UGC0008730', 'UGC0008894', 'UGC0009027', 'UGC0009030', 'UGC0009118', 'UGC0009128', 'UGC0009182', 'UGC0009207', 'UGC0009283', 'UGC0009284', 'UGC0009411', 'UGC0009417', 'UGC0009719', 'UGC0009723', 'UGC0009983', 'UGC0010018', 'UGC0010074', 'UGC0010077', 'UGC0010158', 'UGC0010176', 'UGC0010181', 'UGC0010226', 'UGC0010244', 'UGC0010245', 'UGC0010251', 'UGC0010258', 'UGC0010494', 'UGC0010555', 'UGC0010760', 'UGC0010822', 'UGC0010856', 'UGC0010878', 'UGC0010956', 'UGC0010966', 'UGC0010997', 'UGC0011019', 'UGC0011135', 'UGC0011154', 'UGC0011181', 'UGC0011189', 'UGC0011210', 'UGC0011865', 'UGC0011936', 'UGC0011953', 'UGC0012060', 'UGC0012111', 'UGC0012147', 'UGC0012190', 'UGC0012257', 'UGC0012312', 'UGC0012342', 'UGC0012353', 'UGC0012785', 'UGC0012958', 'UGC0012990', 'UGC0013065', 'UGC0013284', 'UGC0013289', 'UGC0013298') |
| 2.5 | 3.1 | 3.1 | ('UGC0003030', 'UGC0003597') |
| 3 | 3.1 | 3.1 | ('UGC0003285', 'UGC0005403', 'UGC0005587', 'UGC0006877', 'UGC0006920', 'UGC0007760', 'UGC0010161', 'UGC0010166', 'UGC0010175', 'UGC0010500', 'UGC0010992', 'UGC0012006') |
| 3.2 | 2.8 | 2.8 | ('UGC0006047', 'UGC0007640', 'UGC0008337', 'UGC0010332', 'UGC0010534', 'UGC0012001', 'UGC0012162', 'UGC0012759') |
| 3 | 3.1 | 3.1 | ('UGC0004163', 'UGC0005340', 'UGC0005404', 'UGC0005528', 'UGC0006645', 'UGC0006835', 'UGC0007056', 'UGC0007158', 'UGC0007188', 'UGC0007248', 'UGC0012611') |
| 2.7 | 3.4 | 3.4 | ('UGC0002106', 'UGC0002123', 'UGC0002143', 'UGC0002146', 'UGC0002160', 'UGC0002181', 'UGC0003245', 'UGC0003264', 'UGC0003270', 'UGC0010250', 'UGC0012558', 'UGC0012564', 'UGC0012916', 'UGC0013113', 'UGC0013123', 'UGC0013139', 'UGC0013177', 'UGC0013185', 'UGC0013187', 'UGC0013315', 'UGC0013322') |
| 3 | 3.7 | 3.7 | ('UGC0003003', 'UGC0005864', 'UGC0005996', 'UGC0012244', 'UGC0012568', 'UGC0012899', 'UGC0012937', 'UGC0012971') |
| 2.3 | 2.8 | 3.1 | ('UGC0002996', 'UGC0003542', 'UGC0004684', 'UGC0007648', 'UGC0008440', 'UGC0012437') |
| 3 | 3.1 | 3.4 | ('UGC0002099', 'UGC0002100', 'UGC0002130', 'UGC0002163', 'UGC0002168', 'UGC0002195', 'UGC0002328', 'UGC0002513', 'UGC0003014', 'UGC0003237', 'UGC0003314', 'UGC0003360', 'UGC0003535', 'UGC0003537', 'UGC0003725', 'UGC0004101', 'UGC0004125', 'UGC0004139', 'UGC0004160', 'UGC0004164', 'UGC0004468', 'UGC0004482', 'UGC0004483', 'UGC0004497', 'UGC0004523', 'UGC0004537', 'UGC0004565', 'UGC0004577', 'UGC0004598', 'UGC0004620', 'UGC0004631', 'UGC0004649', 'UGC0004651', 'UGC0004660', 'UGC0004687', 'UGC0004688', 'UGC0004706', 'UGC0004713', 'UGC0004732', 'UGC0005004', 'UGC0005375', 'UGC0005775', 'UGC0005871', 'UGC0005901', 'UGC0005903', 'UGC0005916', 'UGC0005989', 'UGC0006003', 'UGC0006034', 'UGC0006053', 'UGC0006069', 'UGC0006312', 'UGC0006326', 'UGC0006360', 'UGC0006405', 'UGC0006419', 'UGC0006427', 'UGC0006440', 'UGC0006502', 'UGC0006531', 'UGC0006581', 'UGC0006638', 'UGC0006646', 'UGC0006666', 'UGC0006746', 'UGC0006823', 'UGC0006850', 'UGC0007053', 'UGC0007132', 'UGC0007161', 'UGC0007233', 'UGC0007240', 'UGC0007244', 'UGC0007266', 'UGC0007295', 'UGC0007426', 'UGC0007646', 'UGC0007663', 'UGC0007700', 'UGC0007708', 'UGC0007756', 'UGC0007900', 'UGC0007927', 'UGC0007937', 'UGC0008172', 'UGC0008180', 'UGC0008250', 'UGC0008302', 'UGC0008314', 'UGC0008318', 'UGC0008333', 'UGC0008336', 'UGC0008386', 'UGC0008422', 'UGC0008522', 'UGC0008605', 'UGC0008661', 'UGC0008669', 'UGC0008682', 'UGC0008797', 'UGC0008879', 'UGC0008881', 'UGC0008890', 'UGC0008895', 'UGC0008901', 'UGC0008907', 'UGC0009050', 'UGC0009069', 'UGC0009109', 'UGC0009111', 'UGC0009191', 'UGC0009249', 'UGC0009266', 'UGC0009270', 'UGC0009325', 'UGC0009442', 'UGC0009594', 'UGC0009774', 'UGC0009998', 'UGC0010213', 'UGC0010273', 'UGC0010287', 'UGC0010496', 'UGC0010560', 'UGC0010807', 'UGC0010816', 'UGC0010912', 'UGC0010928', 'UGC0010974', 'UGC0011016', 'UGC0011020', 'UGC0011424', 'UGC0011547', 'UGC0011562', 'UGC0011567', 'UGC0011654', 'UGC0011784', 'UGC0011895', 'UGC0011921', 'UGC0012056', 'UGC0012110', 'UGC0012182', 'UGC0012186', 'UGC0012260', 'UGC0012308', 'UGC0012397', 'UGC0012626', 'UGC0012629', 'UGC0012792', 'UGC0012801', 'UGC0012806', 'UGC0012815', 'UGC0012930', 'UGC0012939', 'UGC0012948', 'UGC0012980', 'UGC0013006', 'UGC0013015', 'UGC0013047', 'UGC0013076', 'UGC0013091', 'UGC0013105', 'UGC0013202', 'UGC0013203', 'UGC0013233', 'UGC0013239', 'UGC0013263', 'UGC0013352') |
| 2.8 | 3.1 | 3.1 | ('UGC0006321', 'UGC0012728') |
| 3 | 3.1 | 3.4 | ('UGC0002107', 'UGC0002177', 'UGC0002209', 'UGC0002308', 'UGC0002315', 'UGC0002364', 'UGC0002561', 'UGC0003016', 'UGC0003026', 'UGC0003242', 'UGC0003527', 'UGC0003549', 'UGC0003716', 'UGC0003783', 'UGC0004114', 'UGC0004150', 'UGC0004478', 'UGC0004489', 'UGC0004502', 'UGC0004503', 'UGC0004512', 'UGC0004518', 'UGC0004519', 'UGC0004545', 'UGC0004549', 'UGC0004569', 'UGC0004681', 'UGC0004683', 'UGC0004704', 'UGC0004751', 'UGC0004752', 'UGC0004790', 'UGC0005023', 'UGC0005556', 'UGC0005605', 'UGC0005695', 'UGC0005804', 'UGC0005819', 'UGC0005860', 'UGC0005870', 'UGC0005875', 'UGC0005891', 'UGC0005936', 'UGC0005943', 'UGC0005954', 'UGC0005963', 'UGC0005973', 'UGC0006031', 'UGC0006081', 'UGC0006085', 'UGC0006156', 'UGC0006157', 'UGC0006158', 'UGC0006309', 'UGC0006311', 'UGC0006313', 'UGC0006331', 'UGC0006361', 'UGC0006465', 'UGC0006491', 'UGC0006528', 'UGC0006540', 'UGC0006660', 'UGC0006684', 'UGC0006686', 'UGC0006783', 'UGC0006793', 'UGC0006804', 'UGC0006930', 'UGC0006939', 'UGC0006953', 'UGC0007023', 'UGC0007070', 'UGC0007123', 'UGC0007135', 'UGC0007239', 'UGC0007249', 'UGC0007282', 'UGC0007288', 'UGC0007378', 'UGC0007395', 'UGC0007400', 'UGC0007422', 'UGC0007539', 'UGC0007546', 'UGC0007550', 'UGC0007552', 'UGC0007605', 'UGC0007682', 'UGC0007709', 'UGC0007892', 'UGC0007913', 'UGC0007924', 'UGC0008009', 'UGC0008025', 'UGC0008202', 'UGC0008246', 'UGC0008399', 'UGC0008466', 'UGC0008544', 'UGC0008623', 'UGC0008680', 'UGC0008715', 'UGC0008741', 'UGC0008802', 'UGC0008823', 'UGC0008834', 'UGC0008838', 'UGC0008839', 'UGC0008965', 'UGC0009003', 'UGC0009201', 'UGC0009220', 'UGC0009277', 'UGC0009295', 'UGC0009317', 'UGC0009420', 'UGC0009483', 'UGC0009502', 'UGC0009523', 'UGC0009557', 'UGC0009584', 'UGC0009625', 'UGC0009628', 'UGC0009673', 'UGC0009675', 'UGC0009682', 'UGC0009690', 'UGC0009709', 'UGC0009750', 'UGC0009994', 'UGC0010105', 'UGC0010209', 'UGC0010540', 'UGC0010587', 'UGC0010845', 'UGC0010886', 'UGC0011164', 'UGC0011406', 'UGC0011477', 'UGC0011493', 'UGC0011498', 'UGC0011503', 'UGC0011527', 'UGC0011532', 'UGC0011535', 'UGC0011560', 'UGC0011578', 'UGC0011581', 'UGC0011607', 'UGC0011631', 'UGC0011634', 'UGC0011643', 'UGC0011661', 'UGC0011783', 'UGC0011793', 'UGC0011795', 'UGC0011801', 'UGC0011803', 'UGC0011808', 'UGC0011809', 'UGC0011827', 'UGC0011839', 'UGC0011845', 'UGC0011882', 'UGC0011908', 'UGC0011914', 'UGC0011959', 'UGC0011967', 'UGC0011970', 'UGC0011994', 'UGC0012037', 'UGC0012255', 'UGC0012324', 'UGC0012604', 'UGC0012750', 'UGC0012754', 'UGC0012777', 'UGC0012791', 'UGC0013104', 'UGC0013216', 'UGC0013223', 'UGC0013258', 'UGC0013394') |
| 3 | 3.1 | 3.1 | ('UGC0003248', 'UGC0003576', 'UGC0004092', 'UGC0004175', 'UGC0004180', 'UGC0004485', 'UGC0004769', 'UGC0005572', 'UGC0006775', 'UGC0006818', 'UGC0007027', 'UGC0007080', 'UGC0007402', 'UGC0007488', 'UGC0008282', 'UGC0008645', 'UGC0008694', 'UGC0009153', 'UGC0009219', 'UGC0009659', 'UGC0009738', 'UGC0009985', 'UGC0010168', 'UGC0010254', 'UGC0010552', 'UGC0012422', 'UGC0013206', 'UGC0013319') |
| 3.5 | 1.9 | 2.2 | ('UGC0004169', 'UGC0003344') |
| 2.5 | 3.7 | 3.7 | ('UGC0004814', 'UGC0005948', 'UGC0006162', 'UGC0006741', 'UGC0008285', 'UGC0009006', 'UGC0009565', 'UGC0010597', 'UGC0013168') |
| 2.4 | 3.4 | 3.4 | ('UGC0002144', 'UGC0002161', 'UGC0002165', 'UGC0002230', 'UGC0002286', 'UGC0002352', 'UGC0002538', 'UGC0003750', 'UGC0004494', 'UGC0004741', 'UGC0004745', 'UGC0005785', 'UGC0005791', 'UGC0005817', 'UGC0005842', 'UGC0005893', 'UGC0005978', 'UGC0006028', 'UGC0006090', 'UGC0006097', 'UGC0006588', 'UGC0007146', 'UGC0008267', 'UGC0008927', 'UGC0009444', 'UGC0010087', 'UGC0010171', 'UGC0010859', 'UGC0011125', 'UGC0011129', 'UGC0012279', 'UGC0012595', 'UGC0013007') |
| 2.3 | 2.8 | 3.1 | ('UGC0002317', 'UGC0002977', 'UGC0006566', 'UGC0007057', 'UGC0007162', 'UGC0010195', 'UGC0011649', 'UGC0012234') |
| 2.1 | 2.8 | 3.1 | ('UGC0003759', 'UGC0004541', 'UGC0006767', 'UGC0006923', 'UGC0013257') |
| 3.1 | 2.8 | 3.1 | ('UGC0002967', 'UGC0004755', 'UGC0005093', 'UGC0005113', 'UGC0009146', 'UGC0009150', 'UGC0011132') |

**Additional File 4: Analyses of paralogs/contaminations**

**Table 1: Pairs of clusters that have specimens sharing signals**

| Cluster pair | Number of specimens containing shared signals |
|---|---|
| Clust-UGC0002107_Clust-UGC0002177 | 12 |
| Clust-UGC0002177_Clust-UGC0002107 | 8 |
| Clust-UGC0002310_Clust-UGC0007917 | 3 |
| Clust-UGC0002158_Clust-UGC0002126 | 3 |
| Clust-UGC0002158_Clust-UGC0002106 | 3 |
| Clust-UGC0002179_Clust-UGC0002216 | 2 |
| Clust-UGC0002179_Clust-UGC0002283 | 2 |
| Clust-UGC0002978_Clust-UGC0002122 | 2 |
| Clust-UGC0002121_Clust-UGC0002122 | 2 |
| Clust-UGC0006346_Clust-UGC0003776 | 2 |
| Clust-UGC0005824_Clust-UGC0010028 | 2 |
| Clust-UGC0002348_Clust-UGC0002122 | 2 |
| Clust-UGC0002152_Clust-UGC0002122 | 2 |
| Clust-UGC0002142_Clust-UGC0002122 | 2 |
| Clust-UGC0002092_Clust-UGC0002164 | 2 |
| Clust-UGC0002112_Clust-UGC0002164 | 2 |
| Clust-UGC0002124_Clust-UGC0002122 | 2 |
| Clust-UGC0002344_Clust-UGC0002094 | 2 |
| Clust-UGC0002177_Clust-UGC0002122 | 2 |
| Clust-UGC0002330_Clust-UGC0002092 | 2 |
| Clust-UGC0002092_Clust-UGC0002106 | 2 |
| Clust-UGC0002158_Clust-UGC0002132 | 2 |
| Clust-UGC0002114_Clust-UGC0002147 | 2 |

**Table 2: details of specimens analysed**

| SpecimenID | Illumina_clusterID (Lowest UGC code of 3% cluster) | Illumina barcode coverage | Nuniques | Multsignals (yes/no=1/0) | Incongruent cl | Congruent |
|---|---|---|---|---|---|---|
| UGC0012823 | Clust-UGC0002091 | 559 | 1 | 0 | 0 | 0 |
| UGC0010150 | Clust-UGC0002091 | 9379 | 5 | 1 | 1 | 1 |
| UGC0005906 | Clust-UGC0002091 | 2098 | 1 | 0 | 0 | 0 |
| UGC0002091 | Clust-UGC0002091 | 2299 | 2 | 1 | 0 | 0 |
| UGC0003592 | Clust-UGC0002091 | 7899 | 3 | 1 | 0 | 1 |
| UGC0005527 | Clust-UGC0002091 | 79 | 1 | 0 | 0 | 0 |
| UGC0002178 | Clust-UGC0002091 | 1952 | 2 | 1 | 0 | 0 |
| UGC0013332 | Clust-UGC0002091 | 992 | 3 | 1 | 0 | 0 |
| UGC0006106 | Clust-UGC0002091 | 2764 | 2 | 1 | 0 | 0 |
| UGC0007560 | Clust-UGC0002091 | 3623 | 3 | 1 | 0 | 1 |
| UGC0009662 | Clust-UGC0002091 | 1940 | 3 | 1 | 0 | 1 |
| UGC0002284 | Clust-UGC0002091 | 1836 | 1 | 0 | 0 | 0 |
| UGC0011413 | Clust-UGC0002091 | 5408 | 4 | 1 | 0 | 1 |
| UGC0011813 | Clust-UGC0002091 | 1790 | 1 | 0 | 0 | 0 |
| UGC0006138 | Clust-UGC0002091 | 2122 | 1 | 0 | 0 | 0 |
| UGC0008374 | Clust-UGC0002091 | 727 | 1 | 0 | 0 | 0 |
| UGC0012152 | Clust-UGC0002092 | 3206 | 2 | 1 | 0 | 1 |
| UGC0006314 | Clust-UGC0002092 | 2442 | 2 | 1 | 0 | 1 |
| UGC0004811 | Clust-UGC0002092 | 13113 | 75 | 1 | 0 | 1 |
| UGC0005915 | Clust-UGC0002092 | 2706 | 4 | 1 | 0 | 1 |
| UGC0010998 | Clust-UGC0002092 | 1442 | 3 | 1 | 0 | 0 |
| UGC0009005 | Clust-UGC0002092 | 5101 | 5 | 1 | 1 | 1 |
| UGC0011150 | Clust-UGC0002092 | 226 | 1 | 0 | 0 | 0 |
| UGC0005951 | Clust-UGC0002092 | 1398 | 1 | 0 | 0 | 0 |
| UGC0011905 | Clust-UGC0002092 | 9079 | 17 | 1 | 0 | 1 |
| UGC0007064 | Clust-UGC0002092 | 2776 | 2 | 1 | 1 | 0 |
| UGC0006076 | Clust-UGC0002092 | 829 | 1 | 0 | 0 | 0 |
| UGC0007653 | Clust-UGC0002092 | 5747 | 8 | 1 | 0 | 1 |
| UGC0012808 | Clust-UGC0002092 | 1833 | 2 | 1 | 0 | 1 |
| UGC0009545 | Clust-UGC0002092 | 4305 | 4 | 1 | 0 | 1 |
| UGC0013248 | Clust-UGC0002092 | 605 | 1 | 0 | 0 | 0 |
| UGC0012864 | Clust-UGC0002092 | 115 | 1 | 0 | 0 | 0 |
| UGC0012027 | Clust-UGC0002092 | 734 | 1 | 0 | 0 | 0 |
| UGC0011883 | Clust-UGC0002092 | 947 | 2 | 1 | 0 | 1 |
| UGC0006033 | Clust-UGC0002092 | 2554 | 1 | 0 | 0 | 0 |
| UGC0012842 | Clust-UGC0002092 | 6997 | 12 | 1 | 1 | 1 |
| UGC0012843 | Clust-UGC0002092 | 2730 | 2 | 1 | 0 | 0 |
| UGC0009002 | Clust-UGC0002092 | 6309 | 15 | 1 | 1 | 1 |
| UGC0012670 | Clust-UGC0002092 | 4280 | 4 | 1 | 1 | 1 |
| UGC0006757 | Clust-UGC0002092 | 4403 | 3 | 1 | 0 | 0 |
| UGC0005588 | Clust-UGC0002092 | 2867 | 3 | 1 | 0 | 1 |
| UGC0007981 | Clust-UGC0002092 | 1431 | 1 | 0 | 0 | 0 |
| UGC0010805 | Clust-UGC0002092 | 2219 | 5 | 1 | 0 | 0 |
| UGC0011989 | Clust-UGC0002092 | 4665 | 2 | 1 | 0 | 0 |
| UGC0007427 | Clust-UGC0002092 | 910 | 1 | 0 | 0 | 0 |

| | | | | | | |
|---|---|---|---|---|---|---|
| UGC0002092 | Clust-UGC0002092 | 84 | 1 | 0 | 0 | 0 |
| UGC0012570 | Clust-UGC0002092 | 2113 | 3 | 1 | 0 | 1 |
| UGC0009204 | Clust-UGC0002092 | 4157 | 3 | 1 | 0 | 1 |
| UGC0007486 | Clust-UGC0002092 | 1929 | 2 | 1 | 0 | 0 |
| UGC0011826 | Clust-UGC0002092 | 3115 | 2 | 1 | 0 | 1 |
| UGC0008041 | Clust-UGC0002092 | 813 | 1 | 0 | 0 | 0 |
| UGC0008955 | Clust-UGC0002092 | 4672 | 4 | 1 | 0 | 1 |
| UGC0008951 | Clust-UGC0002092 | 2429 | 2 | 1 | 0 | 0 |
| UGC0012926 | Clust-UGC0002092 | 1971 | 3 | 1 | 0 | 1 |
| UGC0012925 | Clust-UGC0002092 | 3998 | 2 | 1 | 0 | 0 |
| UGC0007527 | Clust-UGC0002092 | 4136 | 1 | 0 | 0 | 0 |
| UGC0009626 | Clust-UGC0002092 | 1812 | 1 | 0 | 0 | 0 |
| UGC0006468 | Clust-UGC0002092 | 1416 | 1 | 0 | 0 | 0 |
| UGC0005848 | Clust-UGC0002092 | 3087 | 3 | 1 | 0 | 1 |
| UGC0005844 | Clust-UGC0002092 | 310 | 1 | 0 | 0 | 0 |
| UGC0005843 | Clust-UGC0002092 | 7351 | 11 | 1 | 0 | 1 |
| UGC0007304 | Clust-UGC0002092 | 877 | 1 | 0 | 0 | 0 |
| UGC0009361 | Clust-UGC0002092 | 2100 | 3 | 1 | 0 | 0 |
| UGC0009366 | Clust-UGC0002092 | 5991 | 7 | 1 | 0 | 1 |
| UGC0008994 | Clust-UGC0002092 | 1528 | 3 | 1 | 0 | 0 |
| UGC0006040 | Clust-UGC0002092 | 2270 | 1 | 0 | 0 | 0 |
| UGC0009345 | Clust-UGC0002092 | 4790 | 6 | 1 | 1 | 1 |
| UGC0007561 | Clust-UGC0002092 | 4789 | 5 | 1 | 0 | 1 |
| UGC0005800 | Clust-UGC0002092 | 7176 | 6 | 1 | 0 | 1 |
| UGC0005805 | Clust-UGC0002092 | 4975 | 6 | 1 | 0 | 1 |
| UGC0011187 | Clust-UGC0002092 | 3615 | 5 | 1 | 0 | 1 |
| UGC0006148 | Clust-UGC0002092 | 1746 | 2 | 1 | 0 | 1 |
| UGC0005669 | Clust-UGC0002092 | 1917 | 1 | 0 | 0 | 0 |
| UGC0005793 | Clust-UGC0002092 | 3810 | 1 | 0 | 0 | 0 |
| UGC0005799 | Clust-UGC0002092 | 1238 | 1 | 0 | 0 | 0 |
| UGC0009688 | Clust-UGC0002092 | 1238 | 1 | 0 | 0 | 0 |
| UGC0005961 | Clust-UGC0002092 | 4707 | 3 | 1 | 0 | 1 |
| UGC0009301 | Clust-UGC0002092 | 2575 | 1 | 0 | 0 | 0 |
| UGC0009519 | Clust-UGC0002092 | 1368 | 2 | 1 | 0 | 0 |
| UGC0009132 | Clust-UGC0002092 | 384 | 1 | 0 | 0 | 0 |
| UGC0009186 | Clust-UGC0002092 | 6708 | 7 | 1 | 0 | 1 |
| UGC0005986 | Clust-UGC0002092 | 5581 | 6 | 1 | 0 | 1 |
| UGC0012358 | Clust-UGC0002092 | 2044 | 3 | 1 | 0 | 0 |
| UGC0007971 | Clust-UGC0002092 | 574 | 1 | 0 | 0 | 0 |
| UGC0010986 | Clust-UGC0002092 | 1551 | 4 | 1 | 0 | 0 |
| UGC0011124 | Clust-UGC0002092 | 343 | 1 | 0 | 0 | 0 |
| UGC0011122 | Clust-UGC0002092 | 741 | 1 | 0 | 0 | 0 |
| UGC0008280 | Clust-UGC0002092 | 749 | 1 | 0 | 0 | 0 |
| UGC0006130 | Clust-UGC0002092 | 1457 | 1 | 0 | 0 | 0 |
| UGC0013218 | Clust-UGC0002092 | 245 | 1 | 0 | 0 | 0 |
| UGC0007623 | Clust-UGC0002092 | 4321 | 3 | 1 | 0 | 0 |
| UGC0006499 | Clust-UGC0002092 | 1137 | 1 | 0 | 0 | 0 |
| UGC0006728 | Clust-UGC0002092 | 347 | 1 | 0 | 0 | 0 |
| UGC0011897 | Clust-UGC0002092 | 1774 | 1 | 0 | 0 | 0 |
| UGC0012128 | Clust-UGC0002092 | 2806 | 4 | 1 | 0 | 1 |
| UGC0011004 | Clust-UGC0002092 | 1979 | 1 | 0 | 0 | 0 |
| UGC0009039 | Clust-UGC0002092 | 2176 | 2 | 1 | 0 | 0 |
| UGC0007035 | Clust-UGC0002092 | 3007 | 3 | 1 | 0 | 0 |
| UGC0006345 | Clust-UGC0002092 | 6797 | 7 | 1 | 0 | 1 |
| UGC0013252 | Clust-UGC0002092 | 1449 | 2 | 1 | 1 | 0 |
| UGC0006024 | Clust-UGC0002092 | 396 | 1 | 0 | 0 | 0 |
| UGC0005979 | Clust-UGC0002092 | 1598 | 1 | 0 | 0 | 0 |
| UGC0007419 | Clust-UGC0002092 | 4987 | 3 | 1 | 0 | 1 |
| UGC0004997 | Clust-UGC0002092 | 6634 | 13 | 1 | 0 | 1 |
| UGC0005921 | Clust-UGC0002092 | 4827 | 4 | 1 | 0 | 1 |
| UGC0006963 | Clust-UGC0002092 | 472 | 1 | 0 | 0 | 0 |
| UGC0006629 | Clust-UGC0002092 | 1575 | 2 | 1 | 0 | 0 |
| UGC0006789 | Clust-UGC0002092 | 7216 | 10 | 1 | 0 | 1 |
| UGC0007151 | Clust-UGC0002092 | 3453 | 3 | 1 | 0 | 1 |
| UGC0009486 | Clust-UGC0002092 | 2414 | 3 | 1 | 0 | 0 |
| UGC0010838 | Clust-UGC0002092 | 332 | 1 | 0 | 0 | 0 |
| UGC0008019 | Clust-UGC0002092 | 245 | 1 | 0 | 0 | 0 |
| UGC0005768 | Clust-UGC0002092 | 3752 | 4 | 1 | 0 | 1 |
| UGC0012395 | Clust-UGC0002092 | 10998 | 46 | 1 | 0 | 1 |
| UGC0011497 | Clust-UGC0002092 | 14403 | 93 | 1 | 0 | 1 |
| UGC0010791 | Clust-UGC0002092 | 499 | 1 | 0 | 0 | 0 |
| UGC0006080 | Clust-UGC0002092 | 2439 | 2 | 1 | 0 | 1 |
| UGC0010855 | Clust-UGC0002092 | 945 | 2 | 1 | 0 | 0 |
| UGC0012566 | Clust-UGC0002092 | 3512 | 4 | 1 | 0 | 1 |
| UGC0013087 | Clust-UGC0002092 | 8596 | 26 | 1 | 0 | 1 |
| UGC0007479 | Clust-UGC0002092 | 3260 | 5 | 1 | 0 | 1 |
| UGC0008675 | Clust-UGC0002092 | 161 | 1 | 0 | 0 | 0 |
| UGC0009092 | Clust-UGC0002092 | 1414 | 2 | 1 | 0 | 1 |
| UGC0013327 | Clust-UGC0002092 | 207 | 1 | 0 | 0 | 0 |

| | | | | | | |
|---|---|---|---|---|---|---|
| UGC0012748 | Clust-UGC0002092 | 2761 | 2 | 1 | 0 | 1 |
| UGC0006478 | Clust-UGC0002092 | 1367 | 1 | 0 | 0 | 0 |
| UGC0007703 | Clust-UGC0002092 | 3303 | 6 | 1 | 0 | 1 |
| UGC0008397 | Clust-UGC0002092 | 1399 | 1 | 0 | 0 | 0 |
| UGC0006140 | Clust-UGC0002092 | 1005 | 1 | 0 | 0 | 0 |
| UGC0009373 | Clust-UGC0002092 | 3340 | 3 | 1 | 0 | 0 |
| UGC0006398 | Clust-UGC0002092 | 6208 | 8 | 1 | 1 | 1 |
| UGC0012016 | Clust-UGC0002092 | 2558 | 2 | 1 | 0 | 1 |
| UGC0005857 | Clust-UGC0002092 | 4597 | 5 | 1 | 0 | 1 |
| UGC0002128 | Clust-UGC0002092 | 7760 | 13 | 1 | 0 | 1 |
| UGC0006118 | Clust-UGC0002092 | 2884 | 1 | 0 | 0 | 0 |
| UGC0005836 | Clust-UGC0002092 | 6075 | 5 | 1 | 0 | 1 |
| UGC0007969 | Clust-UGC0002092 | 176 | 1 | 0 | 0 | 0 |
| UGC0005367 | Clust-UGC0002092 | 6678 | 10 | 1 | 0 | 1 |
| UGC0012440 | Clust-UGC0002092 | 2115 | 1 | 0 | 0 | 0 |
| UGC0012443 | Clust-UGC0002092 | 4989 | 5 | 1 | 0 | 1 |
| UGC0008238 | Clust-UGC0002092 | 1479 | 1 | 0 | 0 | 0 |
| UGC0005811 | Clust-UGC0002092 | 3708 | 1 | 0 | 0 | 0 |
| UGC0011191 | Clust-UGC0002092 | 1702 | 2 | 1 | 0 | 0 |
| UGC0006372 | Clust-UGC0002092 | 5926 | 7 | 1 | 0 | 1 |
| UGC0011943 | Clust-UGC0002092 | 11282 | 38 | 1 | 0 | 1 |
| UGC0002093 | Clust-UGC0002093 | 2451 | 1 | 0 | 0 | 0 |
| UGC0009731 | Clust-UGC0002094 | 3346 | 2 | 1 | 0 | 1 |
| UGC0004818 | Clust-UGC0002094 | 6544 | 6 | 1 | 0 | 1 |
| UGC0011155 | Clust-UGC0002094 | 1041 | 3 | 1 | 1 | 0 |
| UGC0004743 | Clust-UGC0002094 | 284 | 1 | 0 | 0 | 0 |
| UGC0006580 | Clust-UGC0002094 | 2384 | 3 | 1 | 0 | 1 |
| UGC0012734 | Clust-UGC0002094 | 1867 | 1 | 0 | 0 | 0 |
| UGC0002094 | Clust-UGC0002094 | 2709 | 2 | 1 | 0 | 0 |
| UGC0006797 | Clust-UGC0002094 | 4752 | 2 | 1 | 0 | 1 |
| UGC0012578 | Clust-UGC0002094 | 8028 | 23 | 1 | 0 | 1 |
| UGC0004612 | Clust-UGC0002094 | 6894 | 14 | 1 | 0 | 1 |
| UGC0012907 | Clust-UGC0002094 | 14542 | 107 | 1 | 0 | 1 |
| UGC0009145 | Clust-UGC0002094 | 4278 | 3 | 1 | 0 | 1 |
| UGC0006386 | Clust-UGC0002094 | 8000 | 24 | 1 | 0 | 1 |
| UGC0009649 | Clust-UGC0002094 | 420 | 1 | 0 | 0 | 0 |
| UGC0006000 | Clust-UGC0002094 | 1356 | 1 | 0 | 0 | 0 |
| UGC0010214 | Clust-UGC0002094 | 517 | 2 | 1 | 0 | 1 |
| UGC0005929 | Clust-UGC0002094 | 5257 | 8 | 1 | 0 | 1 |
| UGC0004600 | Clust-UGC0002094 | 3888 | 1 | 0 | 0 | 0 |
| UGC0009237 | Clust-UGC0002094 | 208 | 1 | 0 | 0 | 0 |
| UGC0004682 | Clust-UGC0002094 | 19319 | 165 | 1 | 0 | 1 |
| UGC0007228 | Clust-UGC0002094 | 6661 | 6 | 1 | 0 | 1 |
| UGC0011194 | Clust-UGC0002094 | 1027 | 1 | 0 | 0 | 0 |
| UGC0002095 | Clust-UGC0002095 | 3314 | 6 | 1 | 0 | 1 |
| UGC0003286 | Clust-UGC0002095 | 9167 | 26 | 1 | 0 | 1 |
| UGC0002096 | Clust-UGC0002096 | 1289 | 3 | 1 | 0 | 0 |
| UGC0002524 | Clust-UGC0002096 | 11377 | 68 | 1 | 0 | 1 |
| UGC0003031 | Clust-UGC0002096 | 3149 | 1 | 0 | 0 | 0 |
| UGC0008274 | Clust-UGC0002097 | 147 | 1 | 0 | 0 | 0 |
| UGC0006991 | Clust-UGC0002097 | 4388 | 1 | 0 | 0 | 0 |
| UGC0004121 | Clust-UGC0002097 | 144 | 1 | 0 | 0 | 0 |
| UGC0005677 | Clust-UGC0002097 | 321 | 1 | 0 | 0 | 0 |
| UGC0005615 | Clust-UGC0002097 | 408 | 1 | 0 | 0 | 0 |
| UGC0003519 | Clust-UGC0002097 | 3975 | 1 | 0 | 0 | 0 |
| UGC0002097 | Clust-UGC0002097 | 2602 | 4 | 1 | 0 | 1 |
| UGC0003571 | Clust-UGC0002097 | 8575 | 4 | 1 | 0 | 1 |
| UGC0007717 | Clust-UGC0002097 | 4819 | 1 | 0 | 0 | 0 |
| UGC0012259 | Clust-UGC0002097 | 166 | 1 | 0 | 0 | 0 |
| UGC0003284 | Clust-UGC0002097 | 2534 | 4 | 1 | 0 | 1 |
| UGC0006571 | Clust-UGC0002097 | 1110 | 1 | 0 | 0 | 0 |
| UGC0007669 | Clust-UGC0002097 | 4068 | 3 | 1 | 0 | 1 |
| UGC0005015 | Clust-UGC0002097 | 10396 | 14 | 1 | 0 | 1 |
| UGC0007255 | Clust-UGC0002097 | 3594 | 1 | 0 | 0 | 0 |
| UGC0006061 | Clust-UGC0002097 | 662 | 1 | 0 | 0 | 0 |
| UGC0005411 | Clust-UGC0002097 | 3053 | 1 | 0 | 0 | 0 |
| UGC0008568 | Clust-UGC0002097 | 1689 | 3 | 1 | 0 | 1 |
| UGC0011830 | Clust-UGC0002097 | 2057 | 1 | 0 | 0 | 0 |
| UGC0006038 | Clust-UGC0002097 | 2797 | 2 | 1 | 0 | 1 |
| UGC0008296 | Clust-UGC0002098 | 665 | 1 | 0 | 0 | 0 |
| UGC0012618 | Clust-UGC0002098 | 1812 | 2 | 1 | 0 | 0 |
| UGC0008555 | Clust-UGC0002098 | 1923 | 2 | 1 | 0 | 0 |
| UGC0007980 | Clust-UGC0002098 | 552 | 1 | 0 | 0 | 0 |
| UGC0002098 | Clust-UGC0002098 | 524 | 1 | 0 | 0 | 0 |
| UGC0010528 | Clust-UGC0002098 | 501 | 1 | 0 | 0 | 0 |
| UGC0003366 | Clust-UGC0002098 | 4339 | 3 | 1 | 1 | 0 |
| UGC0008046 | Clust-UGC0002098 | 417 | 1 | 0 | 0 | 0 |
| UGC0013317 | Clust-UGC0002098 | 4932 | 4 | 1 | 0 | 0 |
| UGC0008321 | Clust-UGC0002098 | 598 | 2 | 1 | 0 | 0 |

| | | | | | | |
|---|---|---|---|---|---|---|
| UGC0011912 | Clust-UGC0002098 | 2439 | 3 | 1 | 0 | 0 |
| UGC0012890 | Clust-UGC0002098 | 4068 | 7 | 1 | 1 | 0 |
| UGC0011805 | Clust-UGC0002098 | 5739 | 4 | 1 | 0 | 0 |
| UGC0008924 | Clust-UGC0002098 | 77 | 1 | 0 | 0 | 0 |
| UGC0007556 | Clust-UGC0002098 | 6544 | 5 | 1 | 0 | 0 |
| UGC0002109 | Clust-UGC0002098 | 3155 | 4 | 1 | 0 | 0 |
| UGC0009583 | Clust-UGC0002098 | 90 | 1 | 0 | 0 | 0 |
| UGC0011567 | Clust-UGC0002099 | 4253 | 2 | 1 | 0 | 1 |
| UGC0011562 | Clust-UGC0002099 | 1879 | 2 | 1 | 0 | 1 |
| UGC0004164 | Clust-UGC0002099 | 3360 | 4 | 1 | 0 | 1 |
| UGC0004160 | Clust-UGC0002099 | 3635 | 4 | 1 | 0 | 1 |
| UGC0007927 | Clust-UGC0002099 | 6367 | 20 | 1 | 0 | 1 |
| UGC0008314 | Clust-UGC0002099 | 394 | 2 | 1 | 0 | 1 |
| UGC0008318 | Clust-UGC0002099 | 1041 | 1 | 0 | 0 | 0 |
| UGC0009191 | Clust-UGC0002099 | 3004 | 1 | 0 | 0 | 0 |
| UGC0005375 | Clust-UGC0002099 | 4806 | 7 | 1 | 0 | 1 |
| UGC0006312 | Clust-UGC0002099 | 3255 | 2 | 1 | 0 | 1 |
| UGC0011921 | Clust-UGC0002099 | 4105 | 1 | 0 | 0 | 0 |
| UGC0011547 | Clust-UGC0002099 | 1862 | 1 | 0 | 0 | 0 |
| UGC0004101 | Clust-UGC0002099 | 373 | 1 | 0 | 0 | 0 |
| UGC0005916 | Clust-UGC0002099 | 1743 | 1 | 0 | 0 | 0 |
| UGC0006053 | Clust-UGC0002099 | 2592 | 1 | 0 | 0 | 0 |
| UGC0007900 | Clust-UGC0002099 | 8029 | 40 | 1 | 0 | 1 |
| UGC0010287 | Clust-UGC0002099 | 2988 | 1 | 0 | 0 | 0 |
| UGC0004598 | Clust-UGC0002099 | 4221 | 2 | 1 | 1 | 0 |
| UGC0007240 | Clust-UGC0002099 | 5094 | 5 | 1 | 0 | 1 |
| UGC0007244 | Clust-UGC0002099 | 6695 | 9 | 1 | 0 | 1 |
| UGC0004125 | Clust-UGC0002099 | 1114 | 1 | 0 | 0 | 0 |
| UGC0012110 | Clust-UGC0002099 | 1893 | 1 | 0 | 0 | 0 |
| UGC0012806 | Clust-UGC0002099 | 2435 | 1 | 0 | 0 | 0 |
| UGC0012801 | Clust-UGC0002099 | 499 | 1 | 0 | 0 | 0 |
| UGC0009774 | Clust-UGC0002099 | 3489 | 3 | 1 | 0 | 1 |
| UGC0013203 | Clust-UGC0002099 | 1804 | 1 | 0 | 0 | 0 |
| UGC0013202 | Clust-UGC0002099 | 488 | 1 | 0 | 0 | 0 |
| UGC0012792 | Clust-UGC0002099 | 498 | 1 | 0 | 0 | 0 |
| UGC0008881 | Clust-UGC0002099 | 4294 | 4 | 1 | 0 | 1 |
| UGC0013015 | Clust-UGC0002099 | 7491 | 16 | 1 | 0 | 1 |
| UGC0011654 | Clust-UGC0002099 | 6622 | 11 | 1 | 0 | 1 |
| UGC0005901 | Clust-UGC0002099 | 4794 | 2 | 1 | 0 | 1 |
| UGC0005903 | Clust-UGC0002099 | 5457 | 6 | 1 | 0 | 1 |
| UGC0006034 | Clust-UGC0002099 | 1937 | 1 | 0 | 0 | 0 |
| UGC0013076 | Clust-UGC0002099 | 3936 | 2 | 1 | 0 | 0 |
| UGC0003725 | Clust-UGC0002099 | 326 | 1 | 0 | 0 | 0 |
| UGC0006581 | Clust-UGC0002099 | 4496 | 4 | 1 | 0 | 1 |
| UGC0010807 | Clust-UGC0002099 | 1816 | 5 | 1 | 0 | 0 |
| UGC0009249 | Clust-UGC0002099 | 9032 | 35 | 1 | 0 | 1 |
| UGC0009069 | Clust-UGC0002099 | 2433 | 1 | 0 | 0 | 0 |
| UGC0008682 | Clust-UGC0002099 | 864 | 1 | 0 | 0 | 0 |
| UGC0007426 | Clust-UGC0002099 | 2519 | 2 | 1 | 0 | 0 |
| UGC0010560 | Clust-UGC0002099 | 3825 | 1 | 0 | 0 | 0 |
| UGC0013263 | Clust-UGC0002099 | 2289 | 1 | 0 | 0 | 0 |
| UGC0007266 | Clust-UGC0002099 | 8783 | 46 | 1 | 0 | 1 |
| UGC0007161 | Clust-UGC0002099 | 4611 | 9 | 1 | 0 | 1 |
| UGC0011424 | Clust-UGC0002099 | 7105 | 14 | 1 | 0 | 1 |
| UGC0009266 | Clust-UGC0002099 | 1941 | 2 | 1 | 1 | 0 |
| UGC0004537 | Clust-UGC0002099 | 9858 | 35 | 1 | 0 | 1 |
| UGC0008522 | Clust-UGC0002099 | 1949 | 1 | 0 | 0 | 0 |
| UGC0002099 | Clust-UGC0002099 | 5822 | 3 | 1 | 0 | 1 |
| UGC0006638 | Clust-UGC0002099 | 1116 | 1 | 0 | 0 | 0 |
| UGC0013091 | Clust-UGC0002099 | 4251 | 3 | 1 | 0 | 1 |
| UGC0006405 | Clust-UGC0002099 | 6874 | 11 | 1 | 0 | 1 |
| UGC0009998 | Clust-UGC0002099 | 1239 | 1 | 0 | 0 | 0 |
| UGC0006326 | Clust-UGC0002099 | 5362 | 4 | 1 | 0 | 1 |
| UGC0011020 | Clust-UGC0002099 | 7630 | 16 | 1 | 0 | 1 |
| UGC0004577 | Clust-UGC0002099 | 9472 | 30 | 1 | 0 | 1 |
| UGC0008669 | Clust-UGC0002099 | 189 | 1 | 0 | 0 | 0 |
| UGC0008661 | Clust-UGC0002099 | 2157 | 1 | 0 | 0 | 0 |
| UGC0006427 | Clust-UGC0002099 | 328 | 1 | 0 | 0 | 0 |
| UGC0003360 | Clust-UGC0002099 | 6628 | 11 | 1 | 0 | 1 |
| UGC0006502 | Clust-UGC0002099 | 478 | 1 | 0 | 0 | 0 |
| UGC0002195 | Clust-UGC0002099 | 2141 | 3 | 1 | 0 | 1 |
| UGC0013352 | Clust-UGC0002099 | 2434 | 3 | 1 | 0 | 0 |
| UGC0006850 | Clust-UGC0002099 | 1311 | 1 | 0 | 0 | 0 |
| UGC0008386 | Clust-UGC0002099 | 2598 | 2 | 1 | 0 | 0 |
| UGC0004651 | Clust-UGC0002099 | 8677 | 19 | 1 | 0 | 1 |
| UGC0009594 | Clust-UGC0002099 | 1505 | 1 | 0 | 0 | 0 |
| UGC0010928 | Clust-UGC0002099 | 156 | 1 | 0 | 0 | 0 |
| UGC0012930 | Clust-UGC0002099 | 7668 | 28 | 1 | 0 | 1 |
| UGC0007756 | Clust-UGC0002099 | 5466 | 5 | 1 | 0 | 1 |

| | | | | | | |
|---|---|---|---|---|---|---|
| UGC0012186 | Clust-UGC0002099 | 4922 | 5 | 1 | 0 | 1 |
| UGC0012182 | Clust-UGC0002099 | 4397 | 5 | 1 | 0 | 1 |
| UGC0012980 | Clust-UGC0002099 | 1536 | 1 | 0 | 0 | 0 |
| UGC0006360 | Clust-UGC0002099 | 4636 | 1 | 0 | 0 | 0 |
| UGC0007937 | Clust-UGC0002099 | 1686 | 1 | 0 | 0 | 0 |
| UGC0008797 | Clust-UGC0002099 | 2062 | 1 | 0 | 0 | 0 |
| UGC0008302 | Clust-UGC0002099 | 1555 | 1 | 0 | 0 | 0 |
| UGC0008879 | Clust-UGC0002099 | 5148 | 4 | 1 | 0 | 1 |
| UGC0012948 | Clust-UGC0002099 | 2744 | 1 | 0 | 0 | 0 |
| UGC0002328 | Clust-UGC0002099 | 1334 | 2 | 1 | 1 | 0 |
| UGC0006531 | Clust-UGC0002099 | 1160 | 1 | 0 | 0 | 0 |
| UGC0007663 | Clust-UGC0002099 | 5280 | 6 | 1 | 0 | 1 |
| UGC0005989 | Clust-UGC0002099 | 7174 | 14 | 1 | 0 | 1 |
| UGC0007053 | Clust-UGC0002099 | 1749 | 1 | 0 | 0 | 0 |
| UGC0013105 | Clust-UGC0002099 | 1486 | 1 | 0 | 0 | 0 |
| UGC0004497 | Clust-UGC0002099 | 6084 | 8 | 1 | 0 | 1 |
| UGC0006440 | Clust-UGC0002099 | 2632 | 2 | 1 | 0 | 1 |
| UGC0007646 | Clust-UGC0002099 | 8338 | 29 | 1 | 0 | 1 |
| UGC0012626 | Clust-UGC0002099 | 6966 | 19 | 1 | 0 | 1 |
| UGC0012629 | Clust-UGC0002099 | 1518 | 2 | 1 | 0 | 0 |
| UGC0004139 | Clust-UGC0002099 | 1179 | 1 | 0 | 0 | 0 |
| UGC0003014 | Clust-UGC0002099 | 3532 | 2 | 1 | 0 | 1 |
| UGC0006069 | Clust-UGC0002099 | 2008 | 1 | 0 | 0 | 0 |
| UGC0010273 | Clust-UGC0002099 | 3587 | 1 | 0 | 0 | 0 |
| UGC0012815 | Clust-UGC0002099 | 651 | 1 | 0 | 0 | 0 |
| UGC0008895 | Clust-UGC0002099 | 3398 | 1 | 0 | 0 | 0 |
| UGC0008890 | Clust-UGC0002099 | 3266 | 2 | 1 | 0 | 0 |
| UGC0004732 | Clust-UGC0002099 | 1968 | 1 | 0 | 0 | 0 |
| UGC0013233 | Clust-UGC0002099 | 3435 | 2 | 1 | 0 | 1 |
| UGC0013239 | Clust-UGC0002099 | 1106 | 2 | 1 | 0 | 0 |
| UGC0002130 | Clust-UGC0002099 | 6232 | 4 | 1 | 0 | 1 |
| UGC0011895 | Clust-UGC0002099 | 5017 | 2 | 1 | 0 | 1 |
| UGC0006003 | Clust-UGC0002099 | 3307 | 1 | 0 | 0 | 0 |
| UGC0011784 | Clust-UGC0002099 | 2096 | 2 | 1 | 0 | 0 |
| UGC0013006 | Clust-UGC0002099 | 3509 | 2 | 1 | 1 | 0 |
| UGC0006746 | Clust-UGC0002099 | 7379 | 24 | 1 | 0 | 1 |
| UGC0007295 | Clust-UGC0002099 | 6232 | 10 | 1 | 0 | 1 |
| UGC0008172 | Clust-UGC0002099 | 1712 | 1 | 0 | 0 | 0 |
| UGC0004482 | Clust-UGC0002099 | 7823 | 20 | 1 | 0 | 1 |
| UGC0004713 | Clust-UGC0002099 | 14977 | 113 | 1 | 0 | 1 |
| UGC0006646 | Clust-UGC0002099 | 2282 | 2 | 1 | 0 | 1 |
| UGC0003314 | Clust-UGC0002099 | 4713 | 4 | 1 | 0 | 1 |
| UGC0010816 | Clust-UGC0002099 | 1151 | 4 | 1 | 0 | 0 |
| UGC0010213 | Clust-UGC0002099 | 531 | 2 | 1 | 1 | 0 |
| UGC0004523 | Clust-UGC0002099 | 8287 | 17 | 1 | 0 | 1 |
| UGC0013047 | Clust-UGC0002099 | 11118 | 64 | 1 | 1 | 1 |
| UGC0009442 | Clust-UGC0002099 | 851 | 1 | 0 | 0 | 0 |
| UGC0011016 | Clust-UGC0002099 | 5129 | 6 | 1 | 0 | 1 |
| UGC0010496 | Clust-UGC0002099 | 257 | 1 | 0 | 0 | 0 |
| UGC0009270 | Clust-UGC0002099 | 4202 | 4 | 1 | 0 | 1 |
| UGC0008901 | Clust-UGC0002099 | 2649 | 1 | 0 | 0 | 0 |
| UGC0008907 | Clust-UGC0002099 | 6236 | 7 | 1 | 0 | 1 |
| UGC0009050 | Clust-UGC0002099 | 3639 | 1 | 0 | 0 | 0 |
| UGC0006419 | Clust-UGC0002099 | 2992 | 2 | 1 | 0 | 1 |
| UGC0012397 | Clust-UGC0002099 | 779 | 1 | 0 | 0 | 0 |
| UGC0004483 | Clust-UGC0002099 | 3579 | 2 | 1 | 0 | 1 |
| UGC0007132 | Clust-UGC0002099 | 336 | 1 | 0 | 0 | 0 |
| UGC0004565 | Clust-UGC0002099 | 5265 | 3 | 1 | 0 | 1 |
| UGC0003537 | Clust-UGC0002099 | 6605 | 12 | 1 | 0 | 1 |
| UGC0003535 | Clust-UGC0002099 | 9760 | 33 | 1 | 0 | 1 |
| UGC0002513 | Clust-UGC0002099 | 3119 | 1 | 0 | 0 | 0 |
| UGC0012939 | Clust-UGC0002099 | 6183 | 13 | 1 | 0 | 1 |
| UGC0004620 | Clust-UGC0002099 | 5140 | 9 | 1 | 0 | 1 |
| UGC0002163 | Clust-UGC0002099 | 3814 | 3 | 1 | 0 | 1 |
| UGC0002168 | Clust-UGC0002099 | 4769 | 5 | 1 | 0 | 1 |
| UGC0012308 | Clust-UGC0002099 | 1618 | 2 | 1 | 0 | 1 |
| UGC0006666 | Clust-UGC0002099 | 4402 | 2 | 1 | 0 | 0 |
| UGC0008422 | Clust-UGC0002099 | 1520 | 1 | 0 | 0 | 0 |
| UGC0005871 | Clust-UGC0002099 | 1050 | 1 | 0 | 0 | 0 |
| UGC0007700 | Clust-UGC0002099 | 4352 | 1 | 0 | 0 | 0 |
| UGC0007708 | Clust-UGC0002099 | 6565 | 4 | 1 | 0 | 1 |
| UGC0005775 | Clust-UGC0002099 | 5502 | 5 | 1 | 0 | 1 |
| UGC0012260 | Clust-UGC0002099 | 411 | 1 | 0 | 0 | 0 |
| UGC0004649 | Clust-UGC0002099 | 9580 | 36 | 1 | 0 | 1 |
| UGC0009111 | Clust-UGC0002099 | 2054 | 1 | 0 | 0 | 0 |
| UGC0008180 | Clust-UGC0002099 | 1247 | 3 | 1 | 0 | 1 |
| UGC0007233 | Clust-UGC0002099 | 8384 | 24 | 1 | 0 | 1 |
| UGC0010912 | Clust-UGC0002099 | 1651 | 1 | 0 | 0 | 0 |
| UGC0004706 | Clust-UGC0002099 | 9696 | 57 | 1 | 0 | 1 |

| | | | | | | |
|---|---|---|---|---|---|---|
| UGC0004660 | Clust-UGC0002099 | 2892 | 2 | 1 | 0 | 1 |
| UGC0010974 | Clust-UGC0002099 | 2213 | 1 | 0 | 0 | 0 |
| UGC0004468 | Clust-UGC0002099 | 4845 | 7 | 1 | 0 | 1 |
| UGC0002100 | Clust-UGC0002099 | 4336 | 4 | 1 | 0 | 1 |
| UGC0006823 | Clust-UGC0002099 | 4614 | 3 | 1 | 0 | 1 |
| UGC0004687 | Clust-UGC0002099 | 6666 | 19 | 1 | 0 | 1 |
| UGC0004688 | Clust-UGC0002099 | 14490 | 107 | 1 | 0 | 1 |
| UGC0012056 | Clust-UGC0002099 | 1869 | 1 | 0 | 0 | 0 |
| UGC0008250 | Clust-UGC0002099 | 536 | 1 | 0 | 0 | 0 |
| UGC0008336 | Clust-UGC0002099 | 1606 | 1 | 0 | 0 | 0 |
| UGC0008333 | Clust-UGC0002099 | 1876 | 1 | 0 | 0 | 0 |
| UGC0004631 | Clust-UGC0002099 | 8051 | 24 | 1 | 0 | 1 |
| UGC0006529 | Clust-UGC0002101 | 4564 | 13 | 1 | 0 | 1 |
| UGC0012119 | Clust-UGC0002101 | 9398 | 41 | 1 | 0 | 1 |
| UGC0012795 | Clust-UGC0002101 | 1109 | 1 | 0 | 0 | 0 |
| UGC0007901 | Clust-UGC0002101 | 4908 | 7 | 1 | 0 | 1 |
| UGC0013356 | Clust-UGC0002101 | 1647 | 3 | 1 | 0 | 1 |
| UGC0005880 | Clust-UGC0002101 | 3262 | 4 | 1 | 1 | 1 |
| UGC0011417 | Clust-UGC0002101 | 4881 | 9 | 1 | 0 | 1 |
| UGC0008693 | Clust-UGC0002101 | 3526 | 8 | 1 | 0 | 1 |
| UGC0012559 | Clust-UGC0002101 | 6451 | 29 | 1 | 0 | 1 |
| UGC0007787 | Clust-UGC0002101 | 6733 | 33 | 1 | 0 | 1 |
| UGC0002101 | Clust-UGC0002101 | 663 | 1 | 0 | 0 | 0 |
| UGC0007453 | Clust-UGC0002101 | 7645 | 32 | 1 | 0 | 1 |
| UGC0008351 | Clust-UGC0002101 | 2386 | 7 | 1 | 0 | 1 |
| UGC0010165 | Clust-UGC0002102 | 4797 | 3 | 1 | 1 | 1 |
| UGC0005654 | Clust-UGC0002102 | 1411 | 1 | 0 | 0 | 0 |
| UGC0005090 | Clust-UGC0002102 | 2514 | 2 | 1 | 0 | 1 |
| UGC0005380 | Clust-UGC0002102 | 164 | 1 | 0 | 0 | 0 |
| UGC0007167 | Clust-UGC0002102 | 6531 | 1 | 0 | 0 | 0 |
| UGC0010488 | Clust-UGC0002102 | 4305 | 3 | 1 | 0 | 1 |
| UGC0004616 | Clust-UGC0002102 | 9917 | 1 | 0 | 0 | 0 |
| UGC0003566 | Clust-UGC0002102 | 7413 | 2 | 1 | 0 | 0 |
| UGC0004693 | Clust-UGC0002102 | 8249 | 3 | 1 | 0 | 1 |
| UGC0010478 | Clust-UGC0002102 | 4942 | 1 | 0 | 0 | 0 |
| UGC0004710 | Clust-UGC0002102 | 5728 | 1 | 0 | 0 | 0 |
| UGC0010537 | Clust-UGC0002102 | 2153 | 1 | 0 | 0 | 0 |
| UGC0009754 | Clust-UGC0002102 | 2901 | 1 | 0 | 0 | 0 |
| UGC0010122 | Clust-UGC0002102 | 3792 | 3 | 1 | 0 | 1 |
| UGC0002102 | Clust-UGC0002102 | 5486 | 5 | 1 | 0 | 1 |
| UGC0004594 | Clust-UGC0002104 | 3562 | 4 | 1 | 1 | 1 |
| UGC0006758 | Clust-UGC0002104 | 5332 | 7 | 1 | 0 | 1 |
| UGC0012866 | Clust-UGC0002104 | 8231 | 35 | 1 | 0 | 1 |
| UGC0011880 | Clust-UGC0002104 | 1446 | 3 | 1 | 0 | 1 |
| UGC0010011 | Clust-UGC0002104 | 512 | 3 | 1 | 0 | 0 |
| UGC0012909 | Clust-UGC0002104 | 847 | 2 | 1 | 0 | 0 |
| UGC0008433 | Clust-UGC0002104 | 145 | 1 | 0 | 0 | 0 |
| UGC0009715 | Clust-UGC0002104 | 2074 | 3 | 1 | 0 | 0 |
| UGC0007771 | Clust-UGC0002104 | 2986 | 4 | 1 | 0 | 0 |
| UGC0012237 | Clust-UGC0002104 | 6055 | 14 | 1 | 0 | 1 |
| UGC0003007 | Clust-UGC0002104 | 758 | 1 | 0 | 0 | 0 |
| UGC0013392 | Clust-UGC0002104 | 3501 | 1 | 0 | 0 | 0 |
| UGC0013390 | Clust-UGC0002104 | 1739 | 2 | 1 | 0 | 0 |
| UGC0008288 | Clust-UGC0002104 | 1927 | 2 | 1 | 0 | 0 |
| UGC0012838 | Clust-UGC0002104 | 7645 | 32 | 1 | 0 | 1 |
| UGC0007999 | Clust-UGC0002104 | 111 | 1 | 0 | 0 | 0 |
| UGC0006908 | Clust-UGC0002104 | 1440 | 1 | 0 | 0 | 0 |
| UGC0008036 | Clust-UGC0002104 | 256 | 1 | 0 | 0 | 0 |
| UGC0008039 | Clust-UGC0002104 | 4417 | 4 | 1 | 0 | 1 |
| UGC0002104 | Clust-UGC0002104 | 1512 | 2 | 1 | 0 | 0 |
| UGC0008819 | Clust-UGC0002104 | 2593 | 2 | 1 | 0 | 0 |
| UGC0008391 | Clust-UGC0002104 | 505 | 2 | 1 | 1 | 0 |
| UGC0006569 | Clust-UGC0002104 | 3567 | 1 | 0 | 0 | 0 |
| UGC0012829 | Clust-UGC0002105 | 6330 | 3 | 1 | 0 | 1 |
| UGC0005935 | Clust-UGC0002105 | 3923 | 3 | 1 | 0 | 1 |
| UGC0002947 | Clust-UGC0002105 | 2224 | 1 | 0 | 0 | 0 |
| UGC0010456 | Clust-UGC0002105 | 4974 | 2 | 1 | 1 | 0 |
| UGC0006430 | Clust-UGC0002105 | 4011 | 1 | 0 | 0 | 0 |
| UGC0002105 | Clust-UGC0002105 | 12785 | 25 | 1 | 0 | 1 |
| UGC0013177 | Clust-UGC0002106 | 127 | 1 | 0 | 0 | 0 |
| UGC0012558 | Clust-UGC0002106 | 7883 | 24 | 1 | 0 | 1 |
| UGC0002146 | Clust-UGC0002106 | 13316 | 73 | 1 | 0 | 1 |
| UGC0003264 | Clust-UGC0002106 | 3528 | 2 | 1 | 0 | 1 |
| UGC0013123 | Clust-UGC0002106 | 4832 | 5 | 1 | 0 | 1 |
| UGC0013187 | Clust-UGC0002106 | 13794 | 88 | 1 | 0 | 1 |
| UGC0010250 | Clust-UGC0002106 | 4138 | 7 | 1 | 1 | 0 |
| UGC0002106 | Clust-UGC0002106 | 14824 | 87 | 1 | 0 | 1 |
| UGC0003270 | Clust-UGC0002106 | 6745 | 17 | 1 | 0 | 1 |
| UGC0012564 | Clust-UGC0002106 | 8536 | 38 | 1 | 0 | 1 |

| | | | | | | |
|---|---|---|---|---|---|---|
| UGC0012916 | Clust-UGC0002106 | 4641 | 6 | 1 | 0 | 1 |
| UGC0002181 | Clust-UGC0002106 | 12191 | 69 | 1 | 0 | 1 |
| UGC0002160 | Clust-UGC0002106 | 13324 | 81 | 1 | 0 | 1 |
| UGC0013322 | Clust-UGC0002106 | 4950 | 13 | 1 | 0 | 1 |
| UGC0002143 | Clust-UGC0002106 | 12127 | 60 | 1 | 0 | 1 |
| UGC0003245 | Clust-UGC0002106 | 7993 | 27 | 1 | 0 | 1 |
| UGC0002123 | Clust-UGC0002106 | 2257 | 3 | 1 | 0 | 1 |
| UGC0013113 | Clust-UGC0002106 | 5883 | 11 | 1 | 0 | 1 |
| UGC0013139 | Clust-UGC0002106 | 1176 | 1 | 0 | 0 | 0 |
| UGC0011560 | Clust-UGC0002107 | 595 | 1 | 0 | 0 | 0 |
| UGC0002315 | Clust-UGC0002107 | 6307 | 6 | 1 | 0 | 1 |
| UGC0009690 | Clust-UGC0002107 | 1276 | 2 | 1 | 0 | 1 |
| UGC0009502 | Clust-UGC0002107 | 4578 | 2 | 1 | 0 | 1 |
| UGC0006331 | Clust-UGC0002107 | 7340 | 20 | 1 | 1 | 1 |
| UGC0011908 | Clust-UGC0002107 | 4789 | 12 | 1 | 1 | 1 |
| UGC0007249 | Clust-UGC0002107 | 4594 | 5 | 1 | 1 | 1 |
| UGC0009523 | Clust-UGC0002107 | 7763 | 32 | 1 | 0 | 1 |
| UGC0012324 | Clust-UGC0002107 | 3913 | 6 | 1 | 0 | 1 |
| UGC0007400 | Clust-UGC0002107 | 363 | 1 | 0 | 0 | 0 |
| UGC0005891 | Clust-UGC0002107 | 5325 | 4 | 1 | 0 | 1 |
| UGC0002561 | Clust-UGC0002107 | 433 | 1 | 0 | 0 | 0 |
| UGC0012791 | Clust-UGC0002107 | 2541 | 3 | 1 | 0 | 1 |
| UGC0009750 | Clust-UGC0002107 | 2791 | 2 | 1 | 0 | 1 |
| UGC0006660 | Clust-UGC0002107 | 2824 | 3 | 1 | 0 | 1 |
| UGC0006031 | Clust-UGC0002107 | 4325 | 4 | 1 | 0 | 1 |
| UGC0012255 | Clust-UGC0002107 | 9765 | 47 | 1 | 0 | 1 |
| UGC0006939 | Clust-UGC0002107 | 5799 | 10 | 1 | 0 | 1 |
| UGC0009003 | Clust-UGC0002107 | 4028 | 4 | 1 | 0 | 1 |
| UGC0011634 | Clust-UGC0002107 | 869 | 2 | 1 | 0 | 1 |
| UGC0006311 | Clust-UGC0002107 | 2907 | 3 | 1 | 0 | 1 |
| UGC0011406 | Clust-UGC0002107 | 3643 | 3 | 1 | 0 | 1 |
| UGC0010209 | Clust-UGC0002107 | 1879 | 1 | 0 | 0 | 0 |
| UGC0005936 | Clust-UGC0002107 | 2374 | 1 | 0 | 0 | 0 |
| UGC0007422 | Clust-UGC0002107 | 6688 | 15 | 1 | 0 | 1 |
| UGC0011498 | Clust-UGC0002107 | 780 | 2 | 1 | 0 | 1 |
| UGC0008009 | Clust-UGC0002107 | 1337 | 3 | 1 | 1 | 1 |
| UGC0005973 | Clust-UGC0002107 | 98 | 1 | 0 | 0 | 0 |
| UGC0009994 | Clust-UGC0002107 | 696 | 1 | 0 | 0 | 0 |
| UGC0011801 | Clust-UGC0002107 | 1727 | 2 | 1 | 0 | 0 |
| UGC0007123 | Clust-UGC0002107 | 647 | 1 | 0 | 0 | 0 |
| UGC0012777 | Clust-UGC0002107 | 981 | 1 | 0 | 0 | 0 |
| UGC0010886 | Clust-UGC0002107 | 2264 | 2 | 1 | 0 | 1 |
| UGC0005860 | Clust-UGC0002107 | 3896 | 5 | 1 | 0 | 1 |
| UGC0002209 | Clust-UGC0002107 | 1153 | 2 | 1 | 0 | 1 |
| UGC0006465 | Clust-UGC0002107 | 1723 | 1 | 0 | 0 | 0 |
| UGC0013104 | Clust-UGC0002107 | 999 | 1 | 0 | 0 | 0 |
| UGC0008839 | Clust-UGC0002107 | 3242 | 4 | 1 | 1 | 1 |
| UGC0011535 | Clust-UGC0002107 | 5836 | 6 | 1 | 0 | 1 |
| UGC0010587 | Clust-UGC0002107 | 4112 | 2 | 1 | 0 | 1 |
| UGC0006361 | Clust-UGC0002107 | 4580 | 3 | 1 | 0 | 1 |
| UGC0007892 | Clust-UGC0002107 | 10565 | 64 | 1 | 1 | 1 |
| UGC0007546 | Clust-UGC0002107 | 8567 | 29 | 1 | 0 | 1 |
| UGC0004150 | Clust-UGC0002107 | 1013 | 1 | 0 | 0 | 0 |
| UGC0002308 | Clust-UGC0002107 | 2325 | 6 | 1 | 1 | 1 |
| UGC0005963 | Clust-UGC0002107 | 1274 | 2 | 1 | 0 | 1 |
| UGC0009682 | Clust-UGC0002107 | 1681 | 4 | 1 | 1 | 1 |
| UGC0011164 | Clust-UGC0002107 | 3337 | 6 | 1 | 1 | 1 |
| UGC0011578 | Clust-UGC0002107 | 1705 | 2 | 1 | 0 | 1 |
| UGC0007913 | Clust-UGC0002107 | 2374 | 2 | 1 | 0 | 1 |
| UGC0011493 | Clust-UGC0002107 | 16060 | 174 | 1 | 0 | 1 |
| UGC0013394 | Clust-UGC0002107 | 187 | 1 | 0 | 0 | 0 |
| UGC0004114 | Clust-UGC0002107 | 1703 | 1 | 0 | 0 | 0 |
| UGC0005943 | Clust-UGC0002107 | 2738 | 2 | 1 | 0 | 1 |
| UGC0009709 | Clust-UGC0002107 | 897 | 2 | 1 | 0 | 1 |
| UGC0011914 | Clust-UGC0002107 | 685 | 1 | 0 | 0 | 0 |
| UGC0004751 | Clust-UGC0002107 | 496 | 2 | 1 | 0 | 1 |
| UGC0013216 | Clust-UGC0002107 | 332 | 1 | 0 | 0 | 0 |
| UGC0009557 | Clust-UGC0002107 | 2244 | 2 | 1 | 0 | 1 |
| UGC0002364 | Clust-UGC0002107 | 273 | 1 | 0 | 0 | 0 |
| UGC0003716 | Clust-UGC0002107 | 1443 | 1 | 0 | 0 | 0 |
| UGC0003016 | Clust-UGC0002107 | 744 | 2 | 1 | 0 | 1 |
| UGC0007605 | Clust-UGC0002107 | 4482 | 5 | 1 | 0 | 1 |
| UGC0011783 | Clust-UGC0002107 | 3463 | 3 | 1 | 1 | 1 |
| UGC0009295 | Clust-UGC0002107 | 639 | 1 | 0 | 0 | 0 |
| UGC0007378 | Clust-UGC0002107 | 602 | 2 | 1 | 1 | 0 |
| UGC0004502 | Clust-UGC0002107 | 6222 | 6 | 1 | 0 | 1 |
| UGC0005023 | Clust-UGC0002107 | 2076 | 3 | 1 | 0 | 1 |
| UGC0009420 | Clust-UGC0002107 | 3061 | 1 | 0 | 0 | 0 |
| UGC0011607 | Clust-UGC0002107 | 3415 | 5 | 1 | 1 | 1 |

| | | | | | | |
|---|---|---|---|---|---|---|
| UGC0003026 | Clust-UGC0002107 | 1394 | 2 | 1 | 0 | 1 |
| UGC0009277 | Clust-UGC0002107 | 4091 | 6 | 1 | 0 | 1 |
| UGC0007135 | Clust-UGC0002107 | 4909 | 6 | 1 | 0 | 1 |
| UGC0006085 | Clust-UGC0002107 | 3055 | 2 | 1 | 0 | 1 |
| UGC0006081 | Clust-UGC0002107 | 602 | 2 | 1 | 0 | 1 |
| UGC0007682 | Clust-UGC0002107 | 9251 | 52 | 1 | 1 | 1 |
| UGC0011809 | Clust-UGC0002107 | 4973 | 4 | 1 | 0 | 1 |
| UGC0011808 | Clust-UGC0002107 | 1488 | 1 | 0 | 0 | 0 |
| UGC0007552 | Clust-UGC0002107 | 7387 | 19 | 1 | 0 | 1 |
| UGC0011581 | Clust-UGC0002107 | 10819 | 80 | 1 | 0 | 1 |
| UGC0007539 | Clust-UGC0002107 | 8890 | 35 | 1 | 0 | 1 |
| UGC0008965 | Clust-UGC0002107 | 1134 | 2 | 1 | 0 | 1 |
| UGC0007709 | Clust-UGC0002107 | 6298 | 12 | 1 | 0 | 1 |
| UGC0009584 | Clust-UGC0002107 | 2633 | 5 | 1 | 1 | 1 |
| UGC0011527 | Clust-UGC0002107 | 6396 | 10 | 1 | 0 | 1 |
| UGC0002107 | Clust-UGC0002107 | 4206 | 6 | 1 | 0 | 1 |
| UGC0008823 | Clust-UGC0002107 | 1073 | 1 | 0 | 0 | 0 |
| UGC0009483 | Clust-UGC0002107 | 5484 | 10 | 1 | 0 | 1 |
| UGC0011967 | Clust-UGC0002107 | 437 | 2 | 1 | 0 | 1 |
| UGC0006686 | Clust-UGC0002107 | 8280 | 18 | 1 | 0 | 1 |
| UGC0011503 | Clust-UGC0002107 | 1167 | 1 | 0 | 0 | 0 |
| UGC0010105 | Clust-UGC0002107 | 3434 | 2 | 1 | 0 | 1 |
| UGC0005819 | Clust-UGC0002107 | 6983 | 21 | 1 | 0 | 1 |
| UGC0009675 | Clust-UGC0002107 | 5859 | 14 | 1 | 0 | 1 |
| UGC0009673 | Clust-UGC0002107 | 3222 | 2 | 1 | 0 | 1 |
| UGC0006540 | Clust-UGC0002107 | 1693 | 1 | 0 | 0 | 0 |
| UGC0003783 | Clust-UGC0002107 | 368 | 1 | 0 | 0 | 0 |
| UGC0006158 | Clust-UGC0002107 | 340 | 2 | 1 | 0 | 1 |
| UGC0006157 | Clust-UGC0002107 | 1407 | 2 | 1 | 0 | 1 |
| UGC0006156 | Clust-UGC0002107 | 1724 | 2 | 1 | 0 | 1 |
| UGC0005689 | Clust-UGC0002108 | 2234 | 1 | 0 | 0 | 0 |
| UGC0002108 | Clust-UGC0002108 | 1032 | 1 | 0 | 0 | 0 |
| UGC0012804 | Clust-UGC0002110 | 1045 | 1 | 0 | 0 | 0 |
| UGC0011404 | Clust-UGC0002110 | 4840 | 5 | 1 | 0 | 1 |
| UGC0006778 | Clust-UGC0002110 | 7098 | 6 | 1 | 0 | 0 |
| UGC0005956 | Clust-UGC0002110 | 2907 | 2 | 1 | 0 | 1 |
| UGC0003329 | Clust-UGC0002110 | 3312 | 2 | 1 | 0 | 1 |
| UGC0010842 | Clust-UGC0002110 | 2246 | 1 | 0 | 0 | 0 |
| UGC0005750 | Clust-UGC0002110 | 4233 | 1 | 0 | 0 | 0 |
| UGC0011487 | Clust-UGC0002110 | 8630 | 7 | 1 | 0 | 1 |
| UGC0002351 | Clust-UGC0002110 | 3669 | 3 | 1 | 1 | 0 |
| UGC0005341 | Clust-UGC0002110 | 2449 | 2 | 1 | 0 | 1 |
| UGC0010083 | Clust-UGC0002110 | 7699 | 1 | 0 | 0 | 0 |
| UGC0003543 | Clust-UGC0002110 | 8789 | 3 | 1 | 0 | 1 |
| UGC0008551 | Clust-UGC0002110 | 4524 | 3 | 1 | 0 | 1 |
| UGC0002522 | Clust-UGC0002110 | 2946 | 1 | 0 | 0 | 0 |
| UGC0010577 | Clust-UGC0002110 | 2965 | 1 | 0 | 0 | 0 |
| UGC0005021 | Clust-UGC0002110 | 8958 | 7 | 1 | 0 | 1 |
| UGC0012726 | Clust-UGC0002110 | 3638 | 3 | 1 | 0 | 1 |
| UGC0011011 | Clust-UGC0002110 | 9009 | 4 | 1 | 0 | 1 |
| UGC0008656 | Clust-UGC0002110 | 2818 | 1 | 0 | 0 | 0 |
| UGC0002975 | Clust-UGC0002110 | 2441 | 2 | 1 | 0 | 1 |
| UGC0002110 | Clust-UGC0002110 | 9837 | 4 | 1 | 0 | 1 |
| UGC0006847 | Clust-UGC0002110 | 1319 | 1 | 0 | 0 | 0 |
| UGC0006862 | Clust-UGC0002110 | 2926 | 3 | 1 | 1 | 0 |
| UGC0008406 | Clust-UGC0002110 | 2880 | 3 | 1 | 1 | 1 |
| UGC0008789 | Clust-UGC0002110 | 2319 | 2 | 1 | 0 | 1 |
| UGC0002293 | Clust-UGC0002111 | 8538 | 22 | 1 | 0 | 1 |
| UGC0010869 | Clust-UGC0002111 | 1112 | 1 | 0 | 0 | 0 |
| UGC0006466 | Clust-UGC0002111 | 3489 | 2 | 1 | 0 | 0 |
| UGC0002949 | Clust-UGC0002111 | 1301 | 1 | 0 | 0 | 0 |
| UGC0008191 | Clust-UGC0002111 | 3777 | 4 | 1 | 0 | 1 |
| UGC0012184 | Clust-UGC0002111 | 6748 | 13 | 1 | 0 | 1 |
| UGC0002111 | Clust-UGC0002111 | 6848 | 13 | 1 | 0 | 1 |
| UGC0007973 | Clust-UGC0002111 | 847 | 1 | 0 | 0 | 0 |
| UGC0006494 | Clust-UGC0002111 | 1971 | 1 | 0 | 0 | 0 |
| UGC0006004 | Clust-UGC0002111 | 5277 | 1 | 0 | 0 | 0 |
| UGC0005595 | Clust-UGC0002111 | 2021 | 1 | 0 | 0 | 0 |
| UGC0002373 | Clust-UGC0002111 | 584 | 1 | 0 | 0 | 0 |
| UGC0012778 | Clust-UGC0002111 | 3333 | 7 | 1 | 0 | 1 |
| UGC0007111 | Clust-UGC0002111 | 1860 | 2 | 1 | 1 | 0 |
| UGC0012655 | Clust-UGC0002111 | 6868 | 13 | 1 | 0 | 1 |
| UGC0013301 | Clust-UGC0002111 | 128 | 1 | 0 | 0 | 0 |
| UGC0003738 | Clust-UGC0002111 | 648 | 1 | 0 | 0 | 0 |
| UGC0008339 | Clust-UGC0002111 | 1505 | 1 | 0 | 0 | 0 |
| UGC0002311 | Clust-UGC0002112 | 5425 | 5 | 1 | 0 | 1 |
| UGC0013159 | Clust-UGC0002112 | 1273 | 2 | 1 | 0 | 1 |
| UGC0007048 | Clust-UGC0002112 | 1785 | 2 | 1 | 0 | 1 |
| UGC0003574 | Clust-UGC0002112 | 9466 | 22 | 1 | 0 | 1 |

| | | | | | | |
|---|---|---|---|---|---|---|
| UGC0007069 | Clust-UGC0002112 | 3304 | 1 | 0 | 0 | 0 |
| UGC0007405 | Clust-UGC0002112 | 4328 | 5 | 1 | 0 | 1 |
| UGC0007267 | Clust-UGC0002112 | 6417 | 14 | 1 | 0 | 1 |
| UGC0008526 | Clust-UGC0002112 | 2247 | 1 | 0 | 0 | 0 |
| UGC0007148 | Clust-UGC0002112 | 679 | 1 | 0 | 0 | 0 |
| UGC0012579 | Clust-UGC0002112 | 2270 | 2 | 1 | 0 | 0 |
| UGC0005868 | Clust-UGC0002112 | 2900 | 5 | 1 | 0 | 1 |
| UGC0012215 | Clust-UGC0002112 | 5678 | 5 | 1 | 0 | 1 |
| UGC0008209 | Clust-UGC0002112 | 2008 | 3 | 1 | 1 | 1 |
| UGC0012235 | Clust-UGC0002112 | 5019 | 4 | 1 | 0 | 1 |
| UGC0002112 | Clust-UGC0002112 | 6293 | 6 | 1 | 0 | 1 |
| UGC0007238 | Clust-UGC0002112 | 3622 | 2 | 1 | 0 | 1 |
| UGC0002350 | Clust-UGC0002112 | 2181 | 1 | 0 | 0 | 0 |
| UGC0008854 | Clust-UGC0002112 | 1996 | 1 | 0 | 0 | 0 |
| UGC0007583 | Clust-UGC0002112 | 8991 | 32 | 1 | 0 | 1 |
| UGC0007587 | Clust-UGC0002112 | 3581 | 2 | 1 | 0 | 1 |
| UGC0007645 | Clust-UGC0002112 | 7142 | 19 | 1 | 0 | 1 |
| UGC0004991 | Clust-UGC0002112 | 8884 | 29 | 1 | 0 | 1 |
| UGC0012912 | Clust-UGC0002112 | 11218 | 62 | 1 | 1 | 1 |
| UGC0009158 | Clust-UGC0002112 | 969 | 1 | 0 | 0 | 0 |
| UGC0007119 | Clust-UGC0002112 | 1262 | 1 | 0 | 0 | 0 |
| UGC0005701 | Clust-UGC0002112 | 8119 | 28 | 1 | 0 | 1 |
| UGC0007518 | Clust-UGC0002112 | 7110 | 10 | 1 | 0 | 1 |
| UGC0009350 | Clust-UGC0002112 | 3548 | 4 | 1 | 1 | 1 |
| UGC0003730 | Clust-UGC0002112 | 981 | 1 | 0 | 0 | 0 |
| UGC0009193 | Clust-UGC0002114 | 2771 | 2 | 1 | 0 | 1 |
| UGC0008485 | Clust-UGC0002114 | 1176 | 1 | 0 | 0 | 0 |
| UGC0007907 | Clust-UGC0002114 | 6442 | 23 | 1 | 0 | 1 |
| UGC0008844 | Clust-UGC0002114 | 2484 | 2 | 1 | 0 | 1 |
| UGC0005659 | Clust-UGC0002114 | 3337 | 4 | 1 | 0 | 1 |
| UGC0005653 | Clust-UGC0002114 | 1211 | 1 | 0 | 0 | 0 |
| UGC0011137 | Clust-UGC0002114 | 1807 | 1 | 0 | 0 | 0 |
| UGC0010249 | Clust-UGC0002114 | 3264 | 3 | 1 | 0 | 1 |
| UGC0005611 | Clust-UGC0002114 | 4511 | 2 | 1 | 0 | 1 |
| UGC0005618 | Clust-UGC0002114 | 5605 | 8 | 1 | 0 | 1 |
| UGC0012922 | Clust-UGC0002114 | 2256 | 2 | 1 | 0 | 1 |
| UGC0008519 | Clust-UGC0002114 | 450 | 1 | 0 | 0 | 0 |
| UGC0005051 | Clust-UGC0002114 | 1642 | 1 | 0 | 0 | 0 |
| UGC0005636 | Clust-UGC0002114 | 5235 | 3 | 1 | 0 | 1 |
| UGC0008520 | Clust-UGC0002114 | 212 | 1 | 0 | 0 | 0 |
| UGC0006898 | Clust-UGC0002114 | 1254 | 3 | 1 | 0 | 1 |
| UGC0008531 | Clust-UGC0002114 | 658 | 1 | 0 | 0 | 0 |
| UGC0004964 | Clust-UGC0002114 | 6988 | 11 | 1 | 0 | 1 |
| UGC0002501 | Clust-UGC0002114 | 1163 | 1 | 0 | 0 | 0 |
| UGC0006422 | Clust-UGC0002114 | 1495 | 1 | 0 | 0 | 0 |
| UGC0006054 | Clust-UGC0002114 | 322 | 1 | 0 | 0 | 0 |
| UGC0008047 | Clust-UGC0002114 | 2831 | 1 | 0 | 0 | 0 |
| UGC0013333 | Clust-UGC0002114 | 5862 | 9 | 1 | 2 | 1 |
| UGC0008413 | Clust-UGC0002114 | 1986 | 1 | 0 | 0 | 0 |
| UGC0008361 | Clust-UGC0002114 | 874 | 1 | 0 | 0 | 0 |
| UGC0002153 | Clust-UGC0002114 | 5210 | 3 | 1 | 0 | 1 |
| UGC0002154 | Clust-UGC0002114 | 3140 | 3 | 1 | 0 | 1 |
| UGC0004472 | Clust-UGC0002114 | 4714 | 8 | 1 | 0 | 1 |
| UGC0002135 | Clust-UGC0002114 | 2241 | 4 | 1 | 0 | 1 |
| UGC0005128 | Clust-UGC0002114 | 2189 | 2 | 1 | 0 | 1 |
| UGC0007950 | Clust-UGC0002114 | 1895 | 2 | 1 | 0 | 1 |
| UGC0005559 | Clust-UGC0002114 | 2372 | 1 | 0 | 0 | 0 |
| UGC0008227 | Clust-UGC0002114 | 108 | 1 | 0 | 0 | 0 |
| UGC0007934 | Clust-UGC0002114 | 3327 | 3 | 1 | 0 | 1 |
| UGC0007939 | Clust-UGC0002114 | 1942 | 2 | 1 | 0 | 1 |
| UGC0008301 | Clust-UGC0002114 | 1217 | 1 | 0 | 0 | 0 |
| UGC0007217 | Clust-UGC0002114 | 2148 | 2 | 1 | 0 | 1 |
| UGC0008266 | Clust-UGC0002114 | 260 | 1 | 0 | 0 | 0 |
| UGC0009135 | Clust-UGC0002114 | 4494 | 7 | 1 | 0 | 1 |
| UGC0009137 | Clust-UGC0002114 | 4517 | 5 | 1 | 0 | 1 |
| UGC0007664 | Clust-UGC0002114 | 3128 | 2 | 1 | 0 | 1 |
| UGC0003547 | Clust-UGC0002114 | 11009 | 48 | 1 | 0 | 1 |
| UGC0010455 | Clust-UGC0002114 | 3740 | 1 | 0 | 0 | 0 |
| UGC0010082 | Clust-UGC0002114 | 4093 | 2 | 1 | 0 | 1 |
| UGC0004734 | Clust-UGC0002114 | 2036 | 2 | 1 | 0 | 1 |
| UGC0009579 | Clust-UGC0002114 | 3192 | 2 | 1 | 0 | 0 |
| UGC0007353 | Clust-UGC0002114 | 2901 | 1 | 0 | 0 | 0 |
| UGC0007607 | Clust-UGC0002114 | 3084 | 2 | 1 | 0 | 1 |
| UGC0012871 | Clust-UGC0002114 | 2068 | 2 | 1 | 0 | 0 |
| UGC0012878 | Clust-UGC0002114 | 9976 | 41 | 1 | 0 | 1 |
| UGC0012663 | Clust-UGC0002114 | 5977 | 14 | 1 | 0 | 1 |
| UGC0008505 | Clust-UGC0002114 | 1564 | 1 | 0 | 0 | 0 |
| UGC0008528 | Clust-UGC0002114 | 210 | 1 | 0 | 0 | 0 |
| UGC0010551 | Clust-UGC0002114 | 3596 | 1 | 0 | 0 | 0 |

| | | | | | | |
|---|---|---|---|---|---|---|
| UGC0005555 | Clust-UGC0002114 | 2438 | 2 | 1 | 0 | 1 |
| UGC0004542 | Clust-UGC0002114 | 5806 | 6 | 1 | 0 | 1 |
| UGC0007457 | Clust-UGC0002114 | 4351 | 3 | 1 | 0 | 1 |
| UGC0012919 | Clust-UGC0002114 | 363 | 1 | 0 | 0 | 0 |
| UGC0003583 | Clust-UGC0002114 | 8663 | 30 | 1 | 0 | 1 |
| UGC0010188 | Clust-UGC0002114 | 2327 | 1 | 0 | 0 | 0 |
| UGC0009091 | Clust-UGC0002114 | 1754 | 3 | 1 | 0 | 1 |
| UGC0008459 | Clust-UGC0002114 | 1065 | 1 | 0 | 0 | 0 |
| UGC0002114 | Clust-UGC0002114 | 3463 | 2 | 1 | 0 | 0 |
| UGC0003556 | Clust-UGC0002114 | 10461 | 42 | 1 | 0 | 1 |
| UGC0013326 | Clust-UGC0002114 | 5439 | 8 | 1 | 0 | 1 |
| UGC0013329 | Clust-UGC0002114 | 6400 | 8 | 1 | 2 | 1 |
| UGC0013302 | Clust-UGC0002114 | 1957 | 1 | 0 | 0 | 0 |
| UGC0005112 | Clust-UGC0002114 | 1917 | 1 | 0 | 0 | 0 |
| UGC0004091 | Clust-UGC0002114 | 64 | 1 | 0 | 0 | 0 |
| UGC0008215 | Clust-UGC0002114 | 2995 | 2 | 1 | 0 | 1 |
| UGC0007573 | Clust-UGC0002114 | 1879 | 2 | 1 | 0 | 1 |
| UGC0011504 | Clust-UGC0002114 | 2785 | 1 | 0 | 0 | 0 |
| UGC0009679 | Clust-UGC0002114 | 1334 | 1 | 0 | 0 | 0 |
| UGC0008253 | Clust-UGC0002114 | 1596 | 1 | 0 | 0 | 0 |
| UGC0008330 | Clust-UGC0002114 | 687 | 1 | 0 | 0 | 0 |
| UGC0008803 | Clust-UGC0002114 | 2515 | 1 | 0 | 0 | 0 |
| UGC0007774 | Clust-UGC0002114 | 2895 | 5 | 1 | 0 | 0 |
| UGC0006318 | Clust-UGC0002115 | 7716 | 19 | 1 | 0 | 1 |
| UGC0007260 | Clust-UGC0002115 | 6336 | 10 | 1 | 0 | 1 |
| UGC0006508 | Clust-UGC0002115 | 3546 | 2 | 1 | 0 | 1 |
| UGC0006057 | Clust-UGC0002115 | 4588 | 5 | 1 | 0 | 1 |
| UGC0007903 | Clust-UGC0002115 | 5420 | 2 | 1 | 0 | 1 |
| UGC0006338 | Clust-UGC0002115 | 2482 | 1 | 0 | 0 | 0 |
| UGC0007002 | Clust-UGC0002115 | 7011 | 23 | 1 | 0 | 1 |
| UGC0013249 | Clust-UGC0002115 | 6754 | 10 | 1 | 0 | 1 |
| UGC0006760 | Clust-UGC0002115 | 8104 | 22 | 1 | 0 | 1 |
| UGC0007180 | Clust-UGC0002115 | 2575 | 3 | 1 | 0 | 1 |
| UGC0007368 | Clust-UGC0002115 | 4979 | 4 | 1 | 0 | 1 |
| UGC0007458 | Clust-UGC0002115 | 4430 | 5 | 1 | 0 | 1 |
| UGC0009262 | Clust-UGC0002115 | 10818 | 69 | 1 | 0 | 1 |
| UGC0010016 | Clust-UGC0002115 | 2175 | 1 | 0 | 0 | 0 |
| UGC0009203 | Clust-UGC0002115 | 4451 | 2 | 1 | 0 | 1 |
| UGC0006903 | Clust-UGC0002115 | 4777 | 1 | 0 | 0 | 0 |
| UGC0006424 | Clust-UGC0002115 | 7728 | 23 | 1 | 0 | 1 |
| UGC0007101 | Clust-UGC0002115 | 2617 | 3 | 1 | 0 | 1 |
| UGC0007503 | Clust-UGC0002115 | 7059 | 15 | 1 | 0 | 1 |
| UGC0009349 | Clust-UGC0002115 | 7919 | 18 | 1 | 0 | 1 |
| UGC0013295 | Clust-UGC0002115 | 2060 | 3 | 1 | 1 | 1 |
| UGC0005776 | Clust-UGC0002115 | 8371 | 28 | 1 | 0 | 1 |
| UGC0009492 | Clust-UGC0002115 | 297 | 1 | 0 | 0 | 0 |
| UGC0004155 | Clust-UGC0002115 | 5499 | 5 | 1 | 0 | 1 |
| UGC0008870 | Clust-UGC0002115 | 4284 | 3 | 1 | 0 | 1 |
| UGC0009728 | Clust-UGC0002115 | 4678 | 5 | 1 | 0 | 1 |
| UGC0006517 | Clust-UGC0002115 | 1508 | 1 | 0 | 0 | 0 |
| UGC0012294 | Clust-UGC0002115 | 3472 | 3 | 1 | 0 | 1 |
| UGC0009707 | Clust-UGC0002115 | 3798 | 3 | 1 | 0 | 1 |
| UGC0009551 | Clust-UGC0002115 | 686 | 1 | 0 | 0 | 0 |
| UGC0009766 | Clust-UGC0002115 | 4599 | 6 | 1 | 0 | 1 |
| UGC0007451 | Clust-UGC0002115 | 2759 | 1 | 0 | 0 | 0 |
| UGC0003037 | Clust-UGC0002115 | 2994 | 2 | 1 | 0 | 1 |
| UGC0006837 | Clust-UGC0002115 | 3580 | 4 | 1 | 0 | 1 |
| UGC0009290 | Clust-UGC0002115 | 3247 | 3 | 1 | 1 | 1 |
| UGC0005838 | Clust-UGC0002115 | 6861 | 12 | 1 | 0 | 1 |
| UGC0007193 | Clust-UGC0002115 | 2503 | 2 | 1 | 0 | 1 |
| UGC0007197 | Clust-UGC0002115 | 4015 | 5 | 1 | 0 | 1 |
| UGC0007370 | Clust-UGC0002115 | 2294 | 1 | 0 | 0 | 0 |
| UGC0007411 | Clust-UGC0002115 | 6084 | 8 | 1 | 0 | 1 |
| UGC0005895 | Clust-UGC0002115 | 4242 | 3 | 1 | 0 | 1 |
| UGC0009785 | Clust-UGC0002115 | 4957 | 7 | 1 | 0 | 1 |
| UGC0007683 | Clust-UGC0002115 | 7382 | 23 | 1 | 0 | 1 |
| UGC0009098 | Clust-UGC0002115 | 848 | 1 | 0 | 0 | 0 |
| UGC0007497 | Clust-UGC0002115 | 10013 | 32 | 1 | 0 | 2 |
| UGC0002115 | Clust-UGC0002115 | 3194 | 3 | 1 | 0 | 1 |
| UGC0009392 | Clust-UGC0002115 | 7585 | 19 | 1 | 0 | 1 |
| UGC0007392 | Clust-UGC0002115 | 3362 | 3 | 1 | 0 | 1 |
| UGC0011144 | Clust-UGC0002115 | 2401 | 2 | 1 | 0 | 1 |
| UGC0009610 | Clust-UGC0002115 | 2163 | 1 | 0 | 0 | 0 |
| UGC0009639 | Clust-UGC0002115 | 1597 | 2 | 1 | 0 | 1 |
| UGC0007963 | Clust-UGC0002115 | 3056 | 5 | 1 | 0 | 1 |
| UGC0008236 | Clust-UGC0002115 | 3031 | 2 | 1 | 0 | 1 |
| UGC0006685 | Clust-UGC0002115 | 4508 | 1 | 0 | 0 | 0 |
| UGC0006747 | Clust-UGC0002115 | 3247 | 2 | 1 | 0 | 1 |
| UGC0008805 | Clust-UGC0002115 | 3371 | 3 | 1 | 0 | 1 |

| | | | | | | |
|---|---|---|---|---|---|---|
| UGC0006554 | Clust-UGC0002116 | 2331 | 1 | 0 | 0 | 0 |
| UGC0004783 | Clust-UGC0002116 | 6137 | 9 | 1 | 1 | 1 |
| UGC0007044 | Clust-UGC0002116 | 2978 | 1 | 0 | 0 | 0 |
| UGC0012112 | Clust-UGC0002116 | 3081 | 2 | 1 | 0 | 1 |
| UGC0006710 | Clust-UGC0002116 | 1124 | 1 | 0 | 0 | 0 |
| UGC0002527 | Clust-UGC0002116 | 3097 | 1 | 0 | 0 | 0 |
| UGC0005932 | Clust-UGC0002116 | 5459 | 7 | 1 | 0 | 1 |
| UGC0006914 | Clust-UGC0002116 | 1708 | 1 | 0 | 0 | 0 |
| UGC0007307 | Clust-UGC0002116 | 6693 | 11 | 1 | 0 | 1 |
| UGC0012552 | Clust-UGC0002116 | 4710 | 4 | 1 | 0 | 1 |
| UGC0005778 | Clust-UGC0002116 | 9599 | 43 | 1 | 0 | 1 |
| UGC0007443 | Clust-UGC0002116 | 894 | 1 | 0 | 0 | 0 |
| UGC0006635 | Clust-UGC0002116 | 735 | 1 | 0 | 0 | 0 |
| UGC0003305 | Clust-UGC0002116 | 2485 | 2 | 1 | 0 | 0 |
| UGC0006986 | Clust-UGC0002116 | 5994 | 12 | 1 | 0 | 1 |
| UGC0005699 | Clust-UGC0002116 | 374 | 1 | 0 | 0 | 0 |
| UGC0006618 | Clust-UGC0002116 | 878 | 1 | 0 | 0 | 0 |
| UGC0002964 | Clust-UGC0002116 | 2453 | 2 | 1 | 0 | 1 |
| UGC0004576 | Clust-UGC0002116 | 5961 | 4 | 1 | 0 | 1 |
| UGC0008049 | Clust-UGC0002116 | 6055 | 16 | 1 | 0 | 1 |
| UGC0007714 | Clust-UGC0002116 | 3498 | 2 | 1 | 0 | 0 |
| UGC0009165 | Clust-UGC0002116 | 511 | 1 | 0 | 0 | 0 |
| UGC0007522 | Clust-UGC0002116 | 7811 | 23 | 1 | 0 | 1 |
| UGC0002543 | Clust-UGC0002116 | 2271 | 1 | 0 | 0 | 0 |
| UGC0011179 | Clust-UGC0002116 | 4725 | 6 | 1 | 0 | 1 |
| UGC0006787 | Clust-UGC0002116 | 5998 | 9 | 1 | 0 | 1 |
| UGC0003607 | Clust-UGC0002116 | 8023 | 25 | 1 | 0 | 1 |
| UGC0003602 | Clust-UGC0002116 | 11944 | 65 | 1 | 0 | 1 |
| UGC0002138 | Clust-UGC0002116 | 2382 | 3 | 1 | 0 | 1 |
| UGC0010112 | Clust-UGC0002116 | 11593 | 64 | 1 | 0 | 1 |
| UGC0002116 | Clust-UGC0002116 | 2729 | 2 | 1 | 0 | 1 |
| UGC0007896 | Clust-UGC0002116 | 11306 | 83 | 1 | 0 | 1 |
| UGC0007932 | Clust-UGC0002116 | 7903 | 30 | 1 | 0 | 1 |
| UGC0007275 | Clust-UGC0002116 | 5370 | 4 | 1 | 0 | 1 |
| UGC0002987 | Clust-UGC0002116 | 2371 | 1 | 0 | 0 | 0 |
| UGC0004802 | Clust-UGC0002116 | 12213 | 80 | 1 | 0 | 1 |
| UGC0008289 | Clust-UGC0002116 | 1534 | 1 | 0 | 0 | 0 |
| UGC0010274 | Clust-UGC0002116 | 12396 | 57 | 1 | 1 | 1 |
| UGC0013238 | Clust-UGC0002116 | 2317 | 1 | 0 | 0 | 0 |
| UGC0003261 | Clust-UGC0002116 | 3910 | 2 | 1 | 0 | 1 |
| UGC0005042 | Clust-UGC0002116 | 2069 | 2 | 1 | 0 | 1 |
| UGC0005971 | Clust-UGC0002116 | 2213 | 1 | 0 | 0 | 0 |
| UGC0007194 | Clust-UGC0002116 | 2835 | 3 | 1 | 0 | 1 |
| UGC0007413 | Clust-UGC0002116 | 1958 | 1 | 0 | 0 | 0 |
| UGC0007173 | Clust-UGC0002116 | 343 | 1 | 0 | 0 | 0 |
| UGC0005927 | Clust-UGC0002116 | 2744 | 1 | 0 | 0 | 0 |
| UGC0005742 | Clust-UGC0002116 | 1182 | 1 | 0 | 0 | 0 |
| UGC0005516 | Clust-UGC0002116 | 1629 | 1 | 0 | 0 | 0 |
| UGC0008452 | Clust-UGC0002116 | 1085 | 1 | 0 | 0 | 0 |
| UGC0002117 | Clust-UGC0002116 | 2440 | 1 | 0 | 0 | 0 |
| UGC0005532 | Clust-UGC0002116 | 190 | 1 | 0 | 0 | 0 |
| UGC0008574 | Clust-UGC0002116 | 3858 | 2 | 1 | 0 | 1 |
| UGC0006135 | Clust-UGC0002116 | 1901 | 1 | 0 | 0 | 0 |
| UGC0008181 | Clust-UGC0002116 | 5577 | 8 | 1 | 0 | 1 |
| UGC0005852 | Clust-UGC0002116 | 3292 | 2 | 1 | 0 | 1 |
| UGC0002375 | Clust-UGC0002116 | 959 | 1 | 0 | 0 | 0 |
| UGC0006807 | Clust-UGC0002116 | 4306 | 1 | 0 | 0 | 0 |
| UGC0004663 | Clust-UGC0002116 | 3250 | 2 | 1 | 0 | 1 |
| UGC0006541 | Clust-UGC0002116 | 3704 | 2 | 1 | 0 | 1 |
| UGC0005359 | Clust-UGC0002118 | 3119 | 6 | 1 | 0 | 1 |
| UGC0002563 | Clust-UGC0002118 | 135 | 1 | 0 | 0 | 0 |
| UGC0002171 | Clust-UGC0002118 | 4937 | 5 | 1 | 0 | 1 |
| UGC0006161 | Clust-UGC0002118 | 828 | 1 | 0 | 0 | 0 |
| UGC0002118 | Clust-UGC0002118 | 7130 | 12 | 1 | 0 | 1 |
| UGC0004500 | Clust-UGC0002118 | 4272 | 4 | 1 | 0 | 1 |
| UGC0011627 | Clust-UGC0002118 | 1197 | 4 | 1 | 0 | 0 |
| UGC0002120 | Clust-UGC0002120 | 3471 | 5 | 1 | 0 | 1 |
| UGC0009411 | Clust-UGC0002121 | 5500 | 5 | 1 | 0 | 1 |
| UGC0005374 | Clust-UGC0002121 | 7950 | 22 | 1 | 0 | 1 |
| UGC0007043 | Clust-UGC0002121 | 687 | 1 | 0 | 0 | 0 |
| UGC0012111 | Clust-UGC0002121 | 4410 | 2 | 1 | 0 | 1 |
| UGC0011135 | Clust-UGC0002121 | 1689 | 1 | 0 | 0 | 0 |
| UGC0010244 | Clust-UGC0002121 | 749 | 1 | 0 | 0 | 0 |
| UGC0010245 | Clust-UGC0002121 | 909 | 1 | 0 | 0 | 0 |
| UGC0009284 | Clust-UGC0002121 | 7333 | 16 | 1 | 0 | 1 |
| UGC0009283 | Clust-UGC0002121 | 3072 | 2 | 1 | 0 | 1 |
| UGC0009027 | Clust-UGC0002121 | 1297 | 1 | 0 | 0 | 0 |
| UGC0007280 | Clust-UGC0002121 | 5794 | 8 | 1 | 0 | 1 |
| UGC0007499 | Clust-UGC0002121 | 4613 | 2 | 1 | 0 | 1 |

| | | | | | | |
|---|---|---|---|---|---|---|
| UGC0012257 | Clust-UGC0002121 | 2857 | 2 | 1 | 0 | 1 |
| UGC0010226 | Clust-UGC0002121 | 3238 | 4 | 1 | 0 | 0 |
| UGC0010074 | Clust-UGC0002121 | 1754 | 1 | 0 | 0 | 0 |
| UGC0005058 | Clust-UGC0002121 | 2343 | 1 | 0 | 0 | 0 |
| UGC0010822 | Clust-UGC0002121 | 3583 | 8 | 1 | 0 | 1 |
| UGC0010018 | Clust-UGC0002121 | 198 | 1 | 0 | 0 | 0 |
| UGC0004816 | Clust-UGC0002121 | 7635 | 11 | 1 | 0 | 1 |
| UGC0009207 | Clust-UGC0002121 | 3214 | 1 | 0 | 0 | 0 |
| UGC0007460 | Clust-UGC0002121 | 2570 | 1 | 0 | 0 | 0 |
| UGC0005361 | Clust-UGC0002121 | 1784 | 1 | 0 | 0 | 0 |
| UGC0005567 | Clust-UGC0002121 | 2459 | 2 | 1 | 0 | 1 |
| UGC0003593 | Clust-UGC0002121 | 5915 | 7 | 1 | 0 | 1 |
| UGC0005735 | Clust-UGC0002121 | 2356 | 2 | 1 | 0 | 1 |
| UGC0002140 | Clust-UGC0002121 | 5596 | 4 | 1 | 0 | 1 |
| UGC0009417 | Clust-UGC0002121 | 5531 | 15 | 1 | 0 | 1 |
| UGC0006317 | Clust-UGC0002121 | 6860 | 9 | 1 | 0 | 1 |
| UGC0007262 | Clust-UGC0002121 | 7085 | 17 | 1 | 1 | 1 |
| UGC0005718 | Clust-UGC0002121 | 4103 | 5 | 1 | 0 | 1 |
| UGC0005710 | Clust-UGC0002121 | 8092 | 18 | 1 | 0 | 1 |
| UGC0010077 | Clust-UGC0002121 | 13503 | 64 | 1 | 1 | 1 |
| UGC0007388 | Clust-UGC0002121 | 277 | 1 | 0 | 0 | 0 |
| UGC0002545 | Clust-UGC0002121 | 817 | 1 | 0 | 0 | 0 |
| UGC0007306 | Clust-UGC0002121 | 6950 | 15 | 1 | 0 | 1 |
| UGC0009719 | Clust-UGC0002121 | 4424 | 5 | 1 | 0 | 1 |
| UGC0006107 | Clust-UGC0002121 | 2381 | 2 | 1 | 0 | 1 |
| UGC0012060 | Clust-UGC0002121 | 3124 | 2 | 1 | 0 | 1 |
| UGC0007504 | Clust-UGC0002121 | 3129 | 2 | 1 | 0 | 1 |
| UGC0003010 | Clust-UGC0002121 | 3510 | 1 | 0 | 0 | 0 |
| UGC0012958 | Clust-UGC0002121 | 5740 | 11 | 1 | 0 | 1 |
| UGC0009128 | Clust-UGC0002121 | 8872 | 24 | 1 | 0 | 1 |
| UGC0013298 | Clust-UGC0002121 | 743 | 2 | 1 | 1 | 0 |
| UGC0010966 | Clust-UGC0002121 | 1583 | 4 | 1 | 0 | 0 |
| UGC0011189 | Clust-UGC0002121 | 8524 | 26 | 1 | 0 | 1 |
| UGC0011181 | Clust-UGC0002121 | 12734 | 90 | 1 | 0 | 1 |
| UGC0007097 | Clust-UGC0002121 | 3312 | 3 | 1 | 0 | 1 |
| UGC0010176 | Clust-UGC0002121 | 338 | 1 | 0 | 0 | 0 |
| UGC0004841 | Clust-UGC0002121 | 7784 | 13 | 1 | 0 | 1 |
| UGC0010158 | Clust-UGC0002121 | 7629 | 15 | 1 | 0 | 1 |
| UGC0004177 | Clust-UGC0002121 | 2084 | 3 | 1 | 0 | 1 |
| UGC0007668 | Clust-UGC0002121 | 5607 | 7 | 1 | 0 | 1 |
| UGC0009723 | Clust-UGC0002121 | 3498 | 3 | 1 | 0 | 1 |
| UGC0011936 | Clust-UGC0002121 | 4669 | 5 | 1 | 0 | 1 |
| UGC0007051 | Clust-UGC0002121 | 833 | 1 | 0 | 0 | 0 |
| UGC0002369 | Clust-UGC0002121 | 307 | 1 | 0 | 0 | 0 |
| UGC0003545 | Clust-UGC0002121 | 6907 | 7 | 1 | 0 | 1 |
| UGC0008894 | Clust-UGC0002121 | 5880 | 6 | 1 | 0 | 1 |
| UGC0012312 | Clust-UGC0002121 | 1192 | 1 | 0 | 0 | 0 |
| UGC0005063 | Clust-UGC0002121 | 4076 | 3 | 1 | 0 | 1 |
| UGC0007352 | Clust-UGC0002121 | 3476 | 1 | 0 | 0 | 0 |
| UGC0012785 | Clust-UGC0002121 | 818 | 1 | 0 | 0 | 0 |
| UGC0010258 | Clust-UGC0002121 | 480 | 1 | 0 | 0 | 0 |
| UGC0010251 | Clust-UGC0002121 | 750 | 2 | 1 | 0 | 1 |
| UGC0006743 | Clust-UGC0002121 | 610 | 1 | 0 | 0 | 0 |
| UGC0007291 | Clust-UGC0002121 | 9370 | 36 | 1 | 0 | 1 |
| UGC0005756 | Clust-UGC0002121 | 8433 | 22 | 1 | 0 | 1 |
| UGC0008730 | Clust-UGC0002121 | 2673 | 3 | 1 | 0 | 1 |
| UGC0008644 | Clust-UGC0002121 | 1559 | 1 | 0 | 0 | 0 |
| UGC0007319 | Clust-UGC0002121 | 3080 | 1 | 0 | 0 | 0 |
| UGC0006781 | Clust-UGC0002121 | 7118 | 15 | 1 | 0 | 1 |
| UGC0010555 | Clust-UGC0002121 | 2886 | 3 | 1 | 0 | 1 |
| UGC0005557 | Clust-UGC0002121 | 1800 | 1 | 0 | 0 | 0 |
| UGC0005009 | Clust-UGC0002121 | 7356 | 9 | 1 | 0 | 1 |
| UGC0012190 | Clust-UGC0002121 | 2872 | 1 | 0 | 0 | 0 |
| UGC0010760 | Clust-UGC0002121 | 2314 | 4 | 1 | 0 | 0 |
| UGC0010494 | Clust-UGC0002121 | 2624 | 3 | 1 | 0 | 1 |
| UGC0005576 | Clust-UGC0002121 | 2943 | 1 | 0 | 0 | 0 |
| UGC0005570 | Clust-UGC0002121 | 2113 | 2 | 1 | 1 | 0 |
| UGC0005687 | Clust-UGC0002121 | 4737 | 5 | 1 | 0 | 1 |
| UGC0007687 | Clust-UGC0002121 | 6661 | 8 | 1 | 0 | 1 |
| UGC0010856 | Clust-UGC0002121 | 3300 | 2 | 1 | 0 | 1 |
| UGC0003306 | Clust-UGC0002121 | 6074 | 9 | 1 | 0 | 1 |
| UGC0009983 | Clust-UGC0002121 | 5481 | 9 | 1 | 0 | 1 |
| UGC0010878 | Clust-UGC0002121 | 4234 | 4 | 1 | 0 | 1 |
| UGC0006600 | Clust-UGC0002121 | 1866 | 2 | 1 | 0 | 1 |
| UGC0005724 | Clust-UGC0002121 | 7510 | 20 | 1 | 1 | 1 |
| UGC0005722 | Clust-UGC0002121 | 5804 | 7 | 1 | 0 | 1 |
| UGC0005729 | Clust-UGC0002121 | 6116 | 5 | 1 | 0 | 1 |
| UGC0009118 | Clust-UGC0002121 | 1182 | 1 | 0 | 0 | 0 |
| UGC0007517 | Clust-UGC0002121 | 6355 | 4 | 1 | 0 | 1 |

| | | | | | | |
|---|---|---|---|---|---|---|
| UGC0002121 | Clust-UGC0002121 | 5720 | 9 | 1 | 0 | 1 |
| UGC0006568 | Clust-UGC0002121 | 4135 | 6 | 1 | 0 | 1 |
| UGC0006357 | Clust-UGC0002121 | 8018 | 11 | 1 | 1 | 1 |
| UGC0007084 | Clust-UGC0002121 | 3841 | 1 | 0 | 0 | 0 |
| UGC0007220 | Clust-UGC0002121 | 2771 | 2 | 1 | 0 | 1 |
| UGC0006544 | Clust-UGC0002121 | 1226 | 1 | 0 | 0 | 0 |
| UGC0002312 | Clust-UGC0002122 | 2960 | 3 | 1 | 0 | 1 |
| UGC0009142 | Clust-UGC0002122 | 71 | 1 | 0 | 0 | 0 |
| UGC0007928 | Clust-UGC0002122 | 8426 | 30 | 1 | 0 | 1 |
| UGC0006523 | Clust-UGC0002122 | 118 | 1 | 0 | 0 | 0 |
| UGC0008310 | Clust-UGC0002122 | 944 | 2 | 1 | 0 | 1 |
| UGC0009732 | Clust-UGC0002122 | 2854 | 2 | 1 | 0 | 1 |
| UGC0006310 | Clust-UGC0002122 | 4957 | 4 | 1 | 0 | 1 |
| UGC0007040 | Clust-UGC0002122 | 3018 | 2 | 1 | 0 | 1 |
| UGC0010147 | Clust-UGC0002122 | 92 | 1 | 0 | 0 | 0 |
| UGC0006671 | Clust-UGC0002122 | 1703 | 1 | 0 | 0 | 0 |
| UGC0007902 | Clust-UGC0002122 | 8552 | 26 | 1 | 0 | 1 |
| UGC0005950 | Clust-UGC0002122 | 994 | 2 | 1 | 0 | 1 |
| UGC0004591 | Clust-UGC0002122 | 3609 | 3 | 1 | 1 | 0 |
| UGC0004597 | Clust-UGC0002122 | 5705 | 6 | 1 | 0 | 1 |
| UGC0007066 | Clust-UGC0002122 | 2640 | 2 | 1 | 0 | 1 |
| UGC0007067 | Clust-UGC0002122 | 1787 | 1 | 0 | 0 | 0 |
| UGC0002995 | Clust-UGC0002122 | 2082 | 2 | 1 | 0 | 1 |
| UGC0006127 | Clust-UGC0002122 | 1506 | 1 | 0 | 0 | 0 |
| UGC0003760 | Clust-UGC0002122 | 356 | 1 | 0 | 0 | 0 |
| UGC0006713 | Clust-UGC0002122 | 3039 | 2 | 1 | 0 | 1 |
| UGC0005399 | Clust-UGC0002122 | 3847 | 2 | 1 | 0 | 1 |
| UGC0007633 | Clust-UGC0002122 | 7356 | 16 | 1 | 0 | 1 |
| UGC0008880 | Clust-UGC0002122 | 2159 | 1 | 0 | 0 | 0 |
| UGC0012653 | Clust-UGC0002122 | 2881 | 3 | 1 | 0 | 1 |
| UGC0010223 | Clust-UGC0002122 | 360 | 2 | 1 | 0 | 1 |
| UGC0011792 | Clust-UGC0002122 | 3187 | 2 | 1 | 0 | 0 |
| UGC0006937 | Clust-UGC0002122 | 4949 | 7 | 1 | 1 | 1 |
| UGC0011635 | Clust-UGC0002122 | 4704 | 3 | 1 | 0 | 1 |
| UGC0005585 | Clust-UGC0002122 | 2558 | 1 | 0 | 0 | 0 |
| UGC0004982 | Clust-UGC0002122 | 247 | 1 | 0 | 0 | 0 |
| UGC0005639 | Clust-UGC0002122 | 2544 | 1 | 0 | 0 | 0 |
| UGC0007187 | Clust-UGC0002122 | 1890 | 2 | 1 | 0 | 1 |
| UGC0007362 | Clust-UGC0002122 | 2030 | 1 | 0 | 0 | 0 |
| UGC0002523 | Clust-UGC0002122 | 718 | 1 | 0 | 0 | 0 |
| UGC0002526 | Clust-UGC0002122 | 862 | 1 | 0 | 0 | 0 |
| UGC0008534 | Clust-UGC0002122 | 1733 | 1 | 0 | 0 | 0 |
| UGC0009160 | Clust-UGC0002122 | 703 | 1 | 0 | 0 | 0 |
| UGC0008407 | Clust-UGC0002122 | 1558 | 1 | 0 | 0 | 0 |
| UGC0011425 | Clust-UGC0002122 | 9283 | 42 | 1 | 0 | 1 |
| UGC0008024 | Clust-UGC0002122 | 2956 | 2 | 1 | 0 | 1 |
| UGC0009048 | Clust-UGC0002122 | 534 | 1 | 0 | 0 | 0 |
| UGC0006650 | Clust-UGC0002122 | 1877 | 2 | 1 | 0 | 1 |
| UGC0011481 | Clust-UGC0002122 | 3163 | 4 | 1 | 0 | 1 |
| UGC0007440 | Clust-UGC0002122 | 1151 | 1 | 0 | 0 | 0 |
| UGC0010542 | Clust-UGC0002122 | 3626 | 1 | 0 | 0 | 0 |
| UGC0004815 | Clust-UGC0002122 | 4743 | 5 | 1 | 0 | 1 |
| UGC0005540 | Clust-UGC0002122 | 1741 | 2 | 1 | 0 | 1 |
| UGC0006091 | Clust-UGC0002122 | 2292 | 1 | 0 | 0 | 0 |
| UGC0006636 | Clust-UGC0002122 | 1343 | 2 | 1 | 0 | 1 |
| UGC0006952 | Clust-UGC0002122 | 5037 | 2 | 1 | 0 | 1 |
| UGC0006984 | Clust-UGC0002122 | 2753 | 3 | 1 | 1 | 1 |
| UGC0006985 | Clust-UGC0002122 | 3577 | 1 | 0 | 0 | 0 |
| UGC0003596 | Clust-UGC0002122 | 4942 | 3 | 1 | 0 | 1 |
| UGC0008453 | Clust-UGC0002122 | 276 | 1 | 0 | 0 | 0 |
| UGC0002969 | Clust-UGC0002122 | 611 | 1 | 0 | 0 | 0 |
| UGC0006426 | Clust-UGC0002122 | 2798 | 3 | 1 | 0 | 1 |
| UGC0010508 | Clust-UGC0002122 | 3490 | 2 | 1 | 0 | 1 |
| UGC0002192 | Clust-UGC0002122 | 170 | 1 | 0 | 0 | 0 |
| UGC0013350 | Clust-UGC0002122 | 3659 | 4 | 1 | 1 | 1 |
| UGC0013355 | Clust-UGC0002122 | 3513 | 2 | 1 | 0 | 1 |
| UGC0006852 | Clust-UGC0002122 | 1886 | 1 | 0 | 0 | 0 |
| UGC0008431 | Clust-UGC0002122 | 2292 | 2 | 1 | 0 | 1 |
| UGC0008404 | Clust-UGC0002122 | 5089 | 4 | 1 | 0 | 1 |
| UGC0006641 | Clust-UGC0002122 | 1259 | 2 | 1 | 0 | 1 |
| UGC0006384 | Clust-UGC0002122 | 6462 | 11 | 1 | 0 | 1 |
| UGC0007732 | Clust-UGC0002122 | 5122 | 5 | 1 | 0 | 1 |
| UGC0008362 | Clust-UGC0002122 | 1342 | 1 | 0 | 0 | 0 |
| UGC0008360 | Clust-UGC0002122 | 854 | 1 | 0 | 0 | 0 |
| UGC0004675 | Clust-UGC0002122 | 6721 | 15 | 1 | 0 | 1 |
| UGC0007502 | Clust-UGC0002122 | 3412 | 2 | 1 | 0 | 1 |
| UGC0008193 | Clust-UGC0002122 | 6026 | 8 | 1 | 0 | 1 |
| UGC0009347 | Clust-UGC0002122 | 4315 | 2 | 1 | 0 | 1 |
| UGC0004477 | Clust-UGC0002122 | 6044 | 8 | 1 | 0 | 1 |

| | | | | | | |
|---|---|---|---|---|---|---|
| UGC0004475 | Clust-UGC0002122 | 5493 | 8 | 1 | 0 | 1 |
| UGC0005120 | Clust-UGC0002122 | 1951 | 2 | 1 | 0 | 1 |
| UGC0007564 | Clust-UGC0002122 | 5919 | 6 | 1 | 0 | 1 |
| UGC0005808 | Clust-UGC0002122 | 5607 | 5 | 1 | 0 | 1 |
| UGC0006579 | Clust-UGC0002122 | 1019 | 2 | 1 | 0 | 1 |
| UGC0008225 | Clust-UGC0002122 | 500 | 1 | 0 | 0 | 0 |
| UGC0010588 | Clust-UGC0002122 | 2226 | 2 | 1 | 0 | 1 |
| UGC0009322 | Clust-UGC0002122 | 5154 | 4 | 1 | 0 | 1 |
| UGC0002309 | Clust-UGC0002122 | 5138 | 3 | 1 | 0 | 1 |
| UGC0007219 | Clust-UGC0002122 | 1364 | 1 | 0 | 0 | 0 |
| UGC0010629 | Clust-UGC0002122 | 1544 | 2 | 1 | 0 | 1 |
| UGC0002326 | Clust-UGC0002122 | 297 | 1 | 0 | 0 | 0 |
| UGC0008729 | Clust-UGC0002122 | 1201 | 1 | 0 | 0 | 0 |
| UGC0003552 | Clust-UGC0002122 | 8903 | 27 | 1 | 0 | 1 |
| UGC0006536 | Clust-UGC0002122 | 1148 | 1 | 0 | 0 | 0 |
| UGC0007919 | Clust-UGC0002122 | 7201 | 12 | 1 | 0 | 1 |
| UGC0008268 | Clust-UGC0002122 | 1248 | 2 | 1 | 0 | 1 |
| UGC0010543 | Clust-UGC0002122 | 2412 | 1 | 0 | 0 | 0 |
| UGC0005987 | Clust-UGC0002122 | 3363 | 2 | 1 | 0 | 1 |
| UGC0007058 | Clust-UGC0002122 | 2684 | 1 | 0 | 0 | 0 |
| UGC0007581 | Clust-UGC0002122 | 5799 | 4 | 1 | 0 | 1 |
| UGC0006706 | Clust-UGC0002122 | 521 | 1 | 0 | 0 | 0 |
| UGC0004808 | Clust-UGC0002122 | 12659 | 80 | 1 | 0 | 1 |
| UGC0009538 | Clust-UGC0002122 | 516 | 1 | 0 | 0 | 0 |
| UGC0005641 | Clust-UGC0002122 | 4035 | 2 | 1 | 0 | 1 |
| UGC0012291 | Clust-UGC0002122 | 2654 | 1 | 0 | 0 | 0 |
| UGC0007641 | Clust-UGC0002122 | 8686 | 35 | 1 | 0 | 1 |
| UGC0011916 | Clust-UGC0002122 | 15313 | 114 | 1 | 1 | 1 |
| UGC0012623 | Clust-UGC0002122 | 6615 | 7 | 1 | 0 | 1 |
| UGC0008541 | Clust-UGC0002122 | 638 | 1 | 0 | 0 | 0 |
| UGC0006289 | Clust-UGC0002122 | 2757 | 3 | 1 | 0 | 0 |
| UGC0006286 | Clust-UGC0002122 | 4358 | 2 | 1 | 0 | 0 |
| UGC0006284 | Clust-UGC0002122 | 4020 | 4 | 1 | 0 | 1 |
| UGC0011453 | Clust-UGC0002122 | 4064 | 4 | 1 | 0 | 1 |
| UGC0005883 | Clust-UGC0002122 | 4252 | 2 | 1 | 0 | 1 |
| UGC0008475 | Clust-UGC0002122 | 900 | 1 | 0 | 0 | 0 |
| UGC0011668 | Clust-UGC0002122 | 2055 | 2 | 1 | 0 | 0 |
| UGC0006001 | Clust-UGC0002122 | 3163 | 2 | 1 | 0 | 1 |
| UGC0011471 | Clust-UGC0002122 | 1188 | 2 | 1 | 0 | 0 |
| UGC0011478 | Clust-UGC0002122 | 2724 | 2 | 1 | 0 | 0 |
| UGC0010257 | Clust-UGC0002122 | 682 | 1 | 0 | 0 | 0 |
| UGC0013003 | Clust-UGC0002122 | 3643 | 4 | 1 | 0 | 1 |
| UGC0006926 | Clust-UGC0002122 | 2159 | 2 | 1 | 0 | 1 |
| UGC0008557 | Clust-UGC0002122 | 3544 | 3 | 1 | 0 | 1 |
| UGC0008171 | Clust-UGC0002122 | 264 | 1 | 0 | 0 | 0 |
| UGC0005621 | Clust-UGC0002122 | 3526 | 1 | 0 | 0 | 0 |
| UGC0005977 | Clust-UGC0002122 | 818 | 1 | 0 | 0 | 0 |
| UGC0008696 | Clust-UGC0002122 | 484 | 1 | 0 | 0 | 0 |
| UGC0011625 | Clust-UGC0002122 | 3706 | 1 | 0 | 0 | 0 |
| UGC0011629 | Clust-UGC0002122 | 2834 | 4 | 1 | 0 | 0 |
| UGC0007137 | Clust-UGC0002122 | 3627 | 1 | 0 | 0 | 0 |
| UGC0008646 | Clust-UGC0002122 | 226 | 1 | 0 | 0 | 0 |
| UGC0008648 | Clust-UGC0002122 | 1650 | 2 | 1 | 0 | 1 |
| UGC0010573 | Clust-UGC0002122 | 1941 | 2 | 1 | 0 | 1 |
| UGC0008736 | Clust-UGC0002122 | 1415 | 1 | 0 | 0 | 0 |
| UGC0004996 | Clust-UGC0002122 | 1836 | 2 | 1 | 0 | 1 |
| UGC0006596 | Clust-UGC0002122 | 1922 | 2 | 1 | 0 | 1 |
| UGC0007179 | Clust-UGC0002122 | 3510 | 1 | 0 | 0 | 0 |
| UGC0007311 | Clust-UGC0002122 | 1579 | 1 | 0 | 0 | 0 |
| UGC0008032 | Clust-UGC0002122 | 1110 | 1 | 0 | 0 | 0 |
| UGC0008030 | Clust-UGC0002122 | 3464 | 1 | 0 | 0 | 0 |
| UGC0004529 | Clust-UGC0002122 | 3947 | 2 | 1 | 0 | 1 |
| UGC0006786 | Clust-UGC0002122 | 4893 | 4 | 1 | 0 | 1 |
| UGC0007785 | Clust-UGC0002122 | 2533 | 2 | 1 | 0 | 1 |
| UGC0005008 | Clust-UGC0002122 | 2452 | 4 | 1 | 0 | 1 |
| UGC0005558 | Clust-UGC0002122 | 931 | 1 | 0 | 0 | 0 |
| UGC0012646 | Clust-UGC0002122 | 4281 | 4 | 1 | 0 | 1 |
| UGC0010495 | Clust-UGC0002122 | 5863 | 8 | 1 | 0 | 1 |
| UGC0006358 | Clust-UGC0002122 | 6875 | 13 | 1 | 0 | 1 |
| UGC0012970 | Clust-UGC0002122 | 9025 | 33 | 1 | 0 | 1 |
| UGC0010536 | Clust-UGC0002122 | 5456 | 4 | 1 | 0 | 1 |
| UGC0004952 | Clust-UGC0002122 | 3787 | 3 | 1 | 0 | 1 |
| UGC0004487 | Clust-UGC0002122 | 7329 | 14 | 1 | 0 | 1 |
| UGC0004954 | Clust-UGC0002122 | 3728 | 3 | 1 | 0 | 1 |
| UGC0004958 | Clust-UGC0002122 | 913 | 1 | 0 | 0 | 0 |
| UGC0005578 | Clust-UGC0002122 | 1866 | 2 | 1 | 0 | 1 |
| UGC0006888 | Clust-UGC0002122 | 4734 | 3 | 1 | 0 | 1 |
| UGC0007136 | Clust-UGC0002122 | 2935 | 1 | 0 | 0 | 0 |
| UGC0008469 | Clust-UGC0002122 | 1822 | 2 | 1 | 0 | 1 |

| | | | | | | |
|---|---|---|---|---|---|---|
| UGC0004561 | Clust-UGC0002122 | 3270 | 2 | 1 | 0 | 1 |
| UGC0005962 | Clust-UGC0002122 | 4541 | 3 | 1 | 0 | 1 |
| UGC0002496 | Clust-UGC0002122 | 267 | 1 | 0 | 0 | 0 |
| UGC0006432 | Clust-UGC0002122 | 3357 | 4 | 1 | 0 | 1 |
| UGC0007221 | Clust-UGC0002122 | 5282 | 4 | 1 | 0 | 1 |
| UGC0010519 | Clust-UGC0002122 | 3778 | 3 | 1 | 0 | 1 |
| UGC0010510 | Clust-UGC0002122 | 1716 | 1 | 0 | 0 | 0 |
| UGC0008679 | Clust-UGC0002122 | 2011 | 1 | 0 | 0 | 0 |
| UGC0004977 | Clust-UGC0002122 | 4068 | 2 | 1 | 0 | 1 |
| UGC0013340 | Clust-UGC0002122 | 854 | 2 | 1 | 0 | 1 |
| UGC0012765 | Clust-UGC0002122 | 1811 | 1 | 0 | 0 | 0 |
| UGC0006601 | Clust-UGC0002122 | 1698 | 2 | 1 | 0 | 1 |
| UGC0008943 | Clust-UGC0002122 | 5080 | 4 | 1 | 0 | 1 |
| UGC0005753 | Clust-UGC0002122 | 5600 | 6 | 1 | 0 | 1 |
| UGC0005538 | Clust-UGC0002122 | 1746 | 2 | 1 | 0 | 1 |
| UGC0006848 | Clust-UGC0002122 | 2695 | 1 | 0 | 0 | 0 |
| UGC0007533 | Clust-UGC0002122 | 4694 | 5 | 1 | 0 | 1 |
| UGC0007530 | Clust-UGC0002122 | 5919 | 3 | 1 | 0 | 1 |
| UGC0008633 | Clust-UGC0002122 | 2697 | 2 | 1 | 0 | 1 |
| UGC0006737 | Clust-UGC0002122 | 3728 | 3 | 1 | 0 | 1 |
| UGC0005877 | Clust-UGC0002122 | 2675 | 2 | 1 | 0 | 1 |
| UGC0003792 | Clust-UGC0002122 | 815 | 1 | 0 | 0 | 0 |
| UGC0003791 | Clust-UGC0002122 | 863 | 2 | 1 | 0 | 1 |
| UGC0008395 | Clust-UGC0002122 | 457 | 1 | 0 | 0 | 0 |
| UGC0007768 | Clust-UGC0002122 | 4006 | 3 | 1 | 0 | 1 |
| UGC0002495 | Clust-UGC0002122 | 387 | 1 | 0 | 0 | 0 |
| UGC0005110 | Clust-UGC0002122 | 1782 | 1 | 0 | 0 | 0 |
| UGC0012014 | Clust-UGC0002122 | 814 | 1 | 0 | 0 | 0 |
| UGC0007510 | Clust-UGC0002122 | 6379 | 7 | 1 | 0 | 1 |
| UGC0008186 | Clust-UGC0002122 | 914 | 1 | 0 | 0 | 0 |
| UGC0012206 | Clust-UGC0002122 | 2294 | 3 | 1 | 0 | 0 |
| UGC0008210 | Clust-UGC0002122 | 829 | 1 | 0 | 0 | 0 |
| UGC0003240 | Clust-UGC0002122 | 4877 | 3 | 1 | 0 | 1 |
| UGC0002125 | Clust-UGC0002122 | 2785 | 2 | 1 | 0 | 0 |
| UGC0002122 | Clust-UGC0002122 | 4102 | 3 | 1 | 0 | 1 |
| UGC0004665 | Clust-UGC0002122 | 10390 | 49 | 1 | 0 | 1 |
| UGC0003364 | Clust-UGC0002122 | 9016 | 29 | 1 | 0 | 1 |
| UGC0012694 | Clust-UGC0002122 | 4683 | 4 | 1 | 0 | 1 |
| UGC0007748 | Clust-UGC0002122 | 3201 | 2 | 1 | 0 | 1 |
| UGC0008232 | Clust-UGC0002122 | 923 | 1 | 0 | 0 | 0 |
| UGC0004467 | Clust-UGC0002122 | 6235 | 17 | 1 | 0 | 1 |
| UGC0006171 | Clust-UGC0002122 | 2057 | 1 | 0 | 0 | 0 |
| UGC0007086 | Clust-UGC0002122 | 2972 | 2 | 1 | 0 | 1 |
| UGC0005814 | Clust-UGC0002122 | 4402 | 2 | 1 | 0 | 1 |
| UGC0008256 | Clust-UGC0002122 | 611 | 1 | 0 | 0 | 0 |
| UGC0005720 | Clust-UGC0002122 | 5060 | 3 | 1 | 0 | 1 |
| UGC0008613 | Clust-UGC0002124 | 5058 | 3 | 1 | 0 | 1 |
| UGC0012176 | Clust-UGC0002124 | 7053 | 3 | 1 | 0 | 1 |
| UGC0007490 | Clust-UGC0002124 | 8988 | 9 | 1 | 0 | 1 |
| UGC0008290 | Clust-UGC0002124 | 1875 | 1 | 0 | 0 | 0 |
| UGC0002993 | Clust-UGC0002124 | 1295 | 1 | 0 | 0 | 0 |
| UGC0006293 | Clust-UGC0002124 | 7101 | 5 | 1 | 0 | 1 |
| UGC0011138 | Clust-UGC0002124 | 537 | 2 | 1 | 0 | 0 |
| UGC0005405 | Clust-UGC0002124 | 4935 | 2 | 1 | 0 | 1 |
| UGC0006691 | Clust-UGC0002124 | 1591 | 4 | 1 | 0 | 0 |
| UGC0002374 | Clust-UGC0002124 | 3493 | 1 | 0 | 0 | 0 |
| UGC0006276 | Clust-UGC0002124 | 6412 | 4 | 1 | 0 | 1 |
| UGC0006768 | Clust-UGC0002124 | 6789 | 2 | 1 | 0 | 0 |
| UGC0007363 | Clust-UGC0002124 | 3888 | 1 | 0 | 0 | 0 |
| UGC0007366 | Clust-UGC0002124 | 6493 | 3 | 1 | 0 | 1 |
| UGC0006915 | Clust-UGC0002124 | 766 | 1 | 0 | 0 | 0 |
| UGC0005387 | Clust-UGC0002124 | 3639 | 2 | 1 | 0 | 1 |
| UGC0005032 | Clust-UGC0002124 | 12167 | 12 | 1 | 0 | 1 |
| UGC0012732 | Clust-UGC0002124 | 1429 | 1 | 0 | 0 | 0 |
| UGC0012883 | Clust-UGC0002124 | 10996 | 9 | 1 | 0 | 1 |
| UGC0009453 | Clust-UGC0002124 | 5582 | 2 | 1 | 0 | 1 |
| UGC0007321 | Clust-UGC0002124 | 750 | 1 | 0 | 0 | 0 |
| UGC0002520 | Clust-UGC0002124 | 3806 | 2 | 1 | 0 | 1 |
| UGC0007463 | Clust-UGC0002124 | 3507 | 1 | 0 | 0 | 0 |
| UGC0006619 | Clust-UGC0002124 | 265 | 1 | 0 | 0 | 0 |
| UGC0008065 | Clust-UGC0002124 | 5918 | 3 | 1 | 0 | 1 |
| UGC0002149 | Clust-UGC0002124 | 13032 | 16 | 1 | 0 | 1 |
| UGC0002548 | Clust-UGC0002124 | 3922 | 1 | 0 | 0 | 0 |
| UGC0009604 | Clust-UGC0002124 | 4746 | 3 | 1 | 0 | 1 |
| UGC0010775 | Clust-UGC0002124 | 63 | 1 | 0 | 0 | 0 |
| UGC0006105 | Clust-UGC0002124 | 2460 | 1 | 0 | 0 | 0 |
| UGC0012950 | Clust-UGC0002124 | 3776 | 2 | 1 | 0 | 1 |
| UGC0009341 | Clust-UGC0002124 | 9082 | 3 | 1 | 0 | 1 |
| UGC0006349 | Clust-UGC0002124 | 7126 | 5 | 1 | 0 | 1 |

| | | | | | | |
|---|---|---|---|---|---|---|
| UGC0012988 | Clust-UGC0002124 | 1685 | 1 | 0 | 0 | 0 |
| UGC0012981 | Clust-UGC0002124 | 4680 | 1 | 0 | 0 | 0 |
| UGC0006369 | Clust-UGC0002124 | 11826 | 18 | 1 | 0 | 1 |
| UGC0004842 | Clust-UGC0002124 | 6152 | 2 | 1 | 0 | 1 |
| UGC0013144 | Clust-UGC0002124 | 162 | 1 | 0 | 0 | 0 |
| UGC0009684 | Clust-UGC0002124 | 1991 | 1 | 0 | 0 | 0 |
| UGC0010949 | Clust-UGC0002124 | 6921 | 5 | 1 | 0 | 1 |
| UGC0005969 | Clust-UGC0002124 | 4003 | 1 | 0 | 0 | 0 |
| UGC0010153 | Clust-UGC0002124 | 5672 | 2 | 1 | 0 | 1 |
| UGC0004795 | Clust-UGC0002124 | 5703 | 3 | 1 | 0 | 1 |
| UGC0010451 | Clust-UGC0002124 | 5378 | 3 | 1 | 1 | 1 |
| UGC0010452 | Clust-UGC0002124 | 5737 | 2 | 1 | 0 | 1 |
| UGC0003755 | Clust-UGC0002124 | 281 | 1 | 0 | 0 | 0 |
| UGC0011558 | Clust-UGC0002124 | 8250 | 5 | 1 | 1 | 1 |
| UGC0008899 | Clust-UGC0002124 | 2667 | 1 | 0 | 0 | 0 |
| UGC0013234 | Clust-UGC0002124 | 4271 | 1 | 0 | 0 | 0 |
| UGC0008713 | Clust-UGC0002124 | 234 | 1 | 0 | 0 | 0 |
| UGC0008617 | Clust-UGC0002124 | 4589 | 3 | 1 | 0 | 0 |
| UGC0005757 | Clust-UGC0002124 | 6740 | 3 | 1 | 0 | 0 |
| UGC0009461 | Clust-UGC0002124 | 7178 | 3 | 1 | 0 | 1 |
| UGC0011439 | Clust-UGC0002124 | 5368 | 4 | 1 | 0 | 0 |
| UGC0010812 | Clust-UGC0002124 | 1735 | 5 | 1 | 0 | 0 |
| UGC0009481 | Clust-UGC0002124 | 5067 | 3 | 1 | 0 | 1 |
| UGC0011010 | Clust-UGC0002124 | 3894 | 2 | 1 | 0 | 1 |
| UGC0011817 | Clust-UGC0002124 | 4826 | 5 | 1 | 0 | 1 |
| UGC0007138 | Clust-UGC0002124 | 3102 | 2 | 1 | 0 | 1 |
| UGC0006628 | Clust-UGC0002124 | 1200 | 1 | 0 | 0 | 0 |
| UGC0006622 | Clust-UGC0002124 | 4262 | 2 | 1 | 0 | 1 |
| UGC0009214 | Clust-UGC0002124 | 2064 | 2 | 1 | 0 | 0 |
| UGC0009981 | Clust-UGC0002124 | 3986 | 3 | 1 | 0 | 1 |
| UGC0004978 | Clust-UGC0002124 | 9504 | 6 | 1 | 0 | 1 |
| UGC0004602 | Clust-UGC0002124 | 3044 | 1 | 0 | 0 | 0 |
| UGC0010891 | Clust-UGC0002124 | 3429 | 1 | 0 | 0 | 0 |
| UGC0007944 | Clust-UGC0002124 | 6253 | 3 | 1 | 0 | 1 |
| UGC0012011 | Clust-UGC0002124 | 7468 | 2 | 1 | 0 | 1 |
| UGC0002124 | Clust-UGC0002124 | 3056 | 2 | 1 | 0 | 0 |
| UGC0006802 | Clust-UGC0002124 | 4639 | 1 | 0 | 0 | 0 |
| UGC0004661 | Clust-UGC0002124 | 3461 | 2 | 1 | 0 | 1 |
| UGC0007572 | Clust-UGC0002124 | 7174 | 4 | 1 | 0 | 1 |
| UGC0006450 | Clust-UGC0002126 | 10724 | 61 | 1 | 1 | 1 |
| UGC0011171 | Clust-UGC0002126 | 3372 | 4 | 1 | 0 | 1 |
| UGC0012753 | Clust-UGC0002126 | 1562 | 2 | 1 | 0 | 1 |
| UGC0005918 | Clust-UGC0002126 | 4447 | 7 | 1 | 0 | 1 |
| UGC0002295 | Clust-UGC0002126 | 9737 | 36 | 1 | 0 | 1 |
| UGC0002290 | Clust-UGC0002126 | 6110 | 3 | 1 | 0 | 1 |
| UGC0012613 | Clust-UGC0002126 | 11580 | 74 | 1 | 0 | 1 |
| UGC0012114 | Clust-UGC0002126 | 3277 | 3 | 1 | 0 | 1 |
| UGC0006278 | Clust-UGC0002126 | 5500 | 9 | 1 | 0 | 1 |
| UGC0012035 | Clust-UGC0002126 | 959 | 1 | 0 | 0 | 0 |
| UGC0010804 | Clust-UGC0002126 | 8597 | 26 | 1 | 0 | 1 |
| UGC0003746 | Clust-UGC0002126 | 576 | 1 | 0 | 0 | 0 |
| UGC0005774 | Clust-UGC0002126 | 10754 | 49 | 1 | 0 | 1 |
| UGC0010549 | Clust-UGC0002126 | 7982 | 16 | 1 | 0 | 1 |
| UGC0010520 | Clust-UGC0002126 | 5836 | 2 | 1 | 0 | 1 |
| UGC0008501 | Clust-UGC0002126 | 4180 | 2 | 1 | 0 | 1 |
| UGC0010864 | Clust-UGC0002126 | 161 | 1 | 0 | 0 | 0 |
| UGC0009086 | Clust-UGC0002126 | 1016 | 2 | 1 | 0 | 1 |
| UGC0009080 | Clust-UGC0002126 | 1483 | 2 | 1 | 0 | 1 |
| UGC0012024 | Clust-UGC0002126 | 2942 | 2 | 1 | 0 | 1 |
| UGC0005923 | Clust-UGC0002126 | 1075 | 2 | 1 | 0 | 1 |
| UGC0012008 | Clust-UGC0002126 | 2815 | 4 | 1 | 0 | 1 |
| UGC0006380 | Clust-UGC0002126 | 7482 | 24 | 1 | 0 | 1 |
| UGC0006817 | Clust-UGC0002126 | 10838 | 54 | 1 | 0 | 1 |
| UGC0006348 | Clust-UGC0002126 | 12054 | 73 | 1 | 0 | 1 |
| UGC0011182 | Clust-UGC0002126 | 511 | 2 | 1 | 0 | 0 |
| UGC0006363 | Clust-UGC0002126 | 7108 | 21 | 1 | 0 | 1 |
| UGC0006143 | Clust-UGC0002126 | 1163 | 1 | 0 | 0 | 0 |
| UGC0007895 | Clust-UGC0002126 | 10005 | 57 | 1 | 0 | 1 |
| UGC0006044 | Clust-UGC0002126 | 6172 | 7 | 1 | 0 | 1 |
| UGC0006288 | Clust-UGC0002126 | 1844 | 2 | 1 | 0 | 1 |
| UGC0005885 | Clust-UGC0002126 | 4540 | 3 | 1 | 0 | 1 |
| UGC0012127 | Clust-UGC0002126 | 7290 | 19 | 1 | 1 | 1 |
| UGC0013000 | Clust-UGC0002126 | 2117 | 1 | 0 | 0 | 0 |
| UGC0008550 | Clust-UGC0002126 | 1521 | 2 | 1 | 0 | 1 |
| UGC0010857 | Clust-UGC0002126 | 7502 | 19 | 1 | 0 | 1 |
| UGC0011996 | Clust-UGC0002126 | 1342 | 1 | 0 | 0 | 0 |
| UGC0011999 | Clust-UGC0002126 | 3049 | 4 | 1 | 0 | 1 |
| UGC0005926 | Clust-UGC0002126 | 3892 | 5 | 1 | 0 | 1 |
| UGC0008821 | Clust-UGC0002126 | 4635 | 2 | 1 | 0 | 1 |

| | | | | | | |
|---|---|---|---|---|---|---|
| UGC0009425 | Clust-UGC0002126 | 3853 | 1 | 0 | 0 | 0 |
| UGC0002242 | Clust-UGC0002126 | 15120 | 116 | 1 | 0 | 1 |
| UGC0006087 | Clust-UGC0002126 | 6766 | 10 | 1 | 0 | 1 |
| UGC0010851 | Clust-UGC0002126 | 5532 | 8 | 1 | 0 | 1 |
| UGC0009213 | Clust-UGC0002126 | 2334 | 4 | 1 | 0 | 0 |
| UGC0006458 | Clust-UGC0002126 | 5776 | 7 | 1 | 0 | 1 |
| UGC0006843 | Clust-UGC0002126 | 7025 | 9 | 1 | 0 | 1 |
| UGC0006399 | Clust-UGC0002126 | 7410 | 15 | 1 | 1 | 1 |
| UGC0009630 | Clust-UGC0002126 | 3585 | 1 | 0 | 0 | 0 |
| UGC0012208 | Clust-UGC0002126 | 11645 | 66 | 1 | 0 | 1 |
| UGC0002126 | Clust-UGC0002126 | 8680 | 18 | 1 | 0 | 1 |
| UGC0006806 | Clust-UGC0002126 | 6661 | 7 | 1 | 0 | 1 |
| UGC0005830 | Clust-UGC0002126 | 3382 | 1 | 0 | 0 | 0 |
| UGC0012442 | Clust-UGC0002126 | 6191 | 4 | 1 | 0 | 1 |
| UGC0008355 | Clust-UGC0002126 | 2134 | 2 | 1 | 0 | 1 |
| UGC0011961 | Clust-UGC0002126 | 4007 | 2 | 1 | 0 | 1 |
| UGC0006155 | Clust-UGC0002126 | 2027 | 2 | 1 | 0 | 1 |
| UGC0010636 | Clust-UGC0002129 | 1167 | 1 | 0 | 0 | 0 |
| UGC0013178 | Clust-UGC0002129 | 1345 | 1 | 0 | 0 | 0 |
| UGC0003009 | Clust-UGC0002129 | 908 | 1 | 0 | 0 | 0 |
| UGC0004826 | Clust-UGC0002129 | 7491 | 8 | 1 | 0 | 1 |
| UGC0008743 | Clust-UGC0002129 | 1081 | 1 | 0 | 0 | 0 |
| UGC0006093 | Clust-UGC0002129 | 1951 | 1 | 0 | 0 | 0 |
| UGC0006164 | Clust-UGC0002129 | 1510 | 1 | 0 | 0 | 0 |
| UGC0005959 | Clust-UGC0002129 | 2049 | 2 | 1 | 0 | 1 |
| UGC0005656 | Clust-UGC0002129 | 636 | 1 | 0 | 0 | 0 |
| UGC0006307 | Clust-UGC0002129 | 4642 | 3 | 1 | 0 | 1 |
| UGC0005642 | Clust-UGC0002129 | 2857 | 1 | 0 | 0 | 0 |
| UGC0011457 | Clust-UGC0002129 | 11106 | 24 | 1 | 0 | 1 |
| UGC0010232 | Clust-UGC0002129 | 542 | 1 | 0 | 0 | 0 |
| UGC0009016 | Clust-UGC0002129 | 1388 | 2 | 1 | 1 | 0 |
| UGC0004520 | Clust-UGC0002129 | 2799 | 1 | 0 | 0 | 0 |
| UGC0008586 | Clust-UGC0002129 | 3702 | 2 | 1 | 0 | 1 |
| UGC0009093 | Clust-UGC0002129 | 294 | 1 | 0 | 0 | 0 |
| UGC0008458 | Clust-UGC0002129 | 1207 | 1 | 0 | 0 | 0 |
| UGC0009178 | Clust-UGC0002129 | 3053 | 2 | 1 | 0 | 1 |
| UGC0003022 | Clust-UGC0002129 | 1086 | 1 | 0 | 0 | 0 |
| UGC0007237 | Clust-UGC0002129 | 4541 | 4 | 1 | 0 | 1 |
| UGC0002129 | Clust-UGC0002129 | 4271 | 3 | 1 | 0 | 1 |
| UGC0006356 | Clust-UGC0002129 | 4475 | 4 | 1 | 0 | 1 |
| UGC0006154 | Clust-UGC0002129 | 2362 | 1 | 0 | 0 | 0 |
| UGC0013240 | Clust-UGC0002131 | 1341 | 1 | 0 | 0 | 0 |
| UGC0010505 | Clust-UGC0002131 | 2416 | 7 | 1 | 1 | 1 |
| UGC0005862 | Clust-UGC0002131 | 1605 | 3 | 1 | 0 | 1 |
| UGC0005106 | Clust-UGC0002131 | 1808 | 2 | 1 | 0 | 0 |
| UGC0002131 | Clust-UGC0002131 | 3617 | 9 | 1 | 0 | 1 |
| UGC0005121 | Clust-UGC0002131 | 997 | 2 | 1 | 0 | 0 |
| UGC0005809 | Clust-UGC0002131 | 6873 | 13 | 1 | 0 | 1 |
| UGC0005600 | Clust-UGC0002131 | 614 | 3 | 1 | 0 | 1 |
| UGC0012221 | Clust-UGC0002131 | 7455 | 16 | 1 | 0 | 1 |
| UGC0008649 | Clust-UGC0002131 | 809 | 2 | 1 | 0 | 0 |
| UGC0004993 | Clust-UGC0002131 | 4842 | 5 | 1 | 0 | 0 |
| UGC0008659 | Clust-UGC0002131 | 2550 | 6 | 1 | 0 | 1 |
| UGC0008477 | Clust-UGC0002131 | 892 | 1 | 0 | 0 | 0 |
| UGC0007723 | Clust-UGC0002131 | 4327 | 11 | 1 | 0 | 1 |
| UGC0006820 | Clust-UGC0002131 | 547 | 1 | 0 | 0 | 0 |
| UGC0011507 | Clust-UGC0002131 | 6950 | 7 | 1 | 0 | 1 |
| UGC0008275 | Clust-UGC0002132 | 1208 | 1 | 0 | 0 | 0 |
| UGC0008864 | Clust-UGC0002132 | 1839 | 1 | 0 | 0 | 0 |
| UGC0009198 | Clust-UGC0002132 | 5906 | 2 | 1 | 0 | 1 |
| UGC0008483 | Clust-UGC0002132 | 1086 | 1 | 0 | 0 | 0 |
| UGC0005997 | Clust-UGC0002132 | 177 | 1 | 0 | 0 | 0 |
| UGC0002332 | Clust-UGC0002132 | 2185 | 1 | 0 | 0 | 0 |
| UGC0013175 | Clust-UGC0002132 | 248 | 1 | 0 | 0 | 0 |
| UGC0013170 | Clust-UGC0002132 | 1999 | 1 | 0 | 0 | 0 |
| UGC0007671 | Clust-UGC0002132 | 5919 | 3 | 1 | 0 | 1 |
| UGC0004129 | Clust-UGC0002132 | 238 | 1 | 0 | 0 | 0 |
| UGC0004120 | Clust-UGC0002132 | 1676 | 1 | 0 | 0 | 0 |
| UGC0012633 | Clust-UGC0002132 | 3049 | 2 | 1 | 0 | 0 |
| UGC0008603 | Clust-UGC0002132 | 5074 | 2 | 1 | 0 | 0 |
| UGC0012868 | Clust-UGC0002132 | 8195 | 6 | 1 | 0 | 1 |
| UGC0008884 | Clust-UGC0002132 | 3752 | 1 | 0 | 0 | 0 |
| UGC0008573 | Clust-UGC0002132 | 4435 | 2 | 1 | 0 | 1 |
| UGC0013387 | Clust-UGC0002132 | 1850 | 2 | 1 | 0 | 0 |
| UGC0007401 | Clust-UGC0002132 | 930 | 1 | 0 | 0 | 0 |
| UGC0011986 | Clust-UGC0002132 | 1584 | 1 | 0 | 0 | 0 |
| UGC0009241 | Clust-UGC0002132 | 12064 | 22 | 1 | 0 | 1 |
| UGC0007420 | Clust-UGC0002132 | 173 | 1 | 0 | 0 | 0 |
| UGC0008026 | Clust-UGC0002132 | 6974 | 6 | 1 | 0 | 1 |

| | | | | | | |
|---|---|---|---|---|---|---|
| UGC0012574 | Clust-UGC0002132 | 8329 | 6 | 1 | 0 | 1 |
| UGC0009993 | Clust-UGC0002132 | 2287 | 1 | 0 | 0 | 0 |
| UGC0013371 | Clust-UGC0002132 | 3072 | 2 | 1 | 0 | 0 |
| UGC0013374 | Clust-UGC0002132 | 1888 | 2 | 1 | 0 | 0 |
| UGC0012596 | Clust-UGC0002132 | 5861 | 2 | 1 | 0 | 0 |
| UGC0009221 | Clust-UGC0002132 | 9605 | 9 | 1 | 0 | 1 |
| UGC0003369 | Clust-UGC0002132 | 5145 | 5 | 1 | 0 | 1 |
| UGC0012901 | Clust-UGC0002132 | 5986 | 3 | 1 | 0 | 0 |
| UGC0006469 | Clust-UGC0002132 | 4621 | 2 | 1 | 0 | 1 |
| UGC0013353 | Clust-UGC0002132 | 5113 | 3 | 1 | 0 | 1 |
| UGC0013351 | Clust-UGC0002132 | 312 | 2 | 1 | 0 | 0 |
| UGC0008040 | Clust-UGC0002132 | 2147 | 1 | 0 | 0 | 0 |
| UGC0008604 | Clust-UGC0002132 | 7081 | 1 | 0 | 0 | 0 |
| UGC0009601 | Clust-UGC0002132 | 1637 | 1 | 0 | 0 | 0 |
| UGC0008601 | Clust-UGC0002132 | 7722 | 2 | 1 | 0 | 1 |
| UGC0009478 | Clust-UGC0002132 | 251 | 1 | 0 | 0 | 0 |
| UGC0008626 | Clust-UGC0002132 | 4062 | 2 | 1 | 0 | 1 |
| UGC0008846 | Clust-UGC0002132 | 233 | 1 | 0 | 0 | 0 |
| UGC0008368 | Clust-UGC0002132 | 222 | 1 | 0 | 0 | 0 |
| UGC0009713 | Clust-UGC0002132 | 3364 | 1 | 0 | 0 | 0 |
| UGC0002155 | Clust-UGC0002132 | 3104 | 2 | 1 | 0 | 0 |
| UGC0009102 | Clust-UGC0002132 | 1698 | 1 | 0 | 0 | 0 |
| UGC0012682 | Clust-UGC0002132 | 9908 | 12 | 1 | 2 | 1 |
| UGC0005826 | Clust-UGC0002132 | 5983 | 6 | 1 | 1 | 1 |
| UGC0008346 | Clust-UGC0002132 | 684 | 1 | 0 | 0 | 0 |
| UGC0012610 | Clust-UGC0002132 | 6235 | 2 | 1 | 0 | 0 |
| UGC0002137 | Clust-UGC0002132 | 1954 | 2 | 1 | 0 | 0 |
| UGC0007954 | Clust-UGC0002132 | 858 | 1 | 0 | 0 | 0 |
| UGC0007951 | Clust-UGC0002132 | 3526 | 3 | 1 | 0 | 1 |
| UGC0006577 | Clust-UGC0002132 | 3108 | 1 | 0 | 0 | 0 |
| UGC0008222 | Clust-UGC0002132 | 856 | 1 | 0 | 0 | 0 |
| UGC0008326 | Clust-UGC0002132 | 1109 | 2 | 1 | 0 | 0 |
| UGC0009491 | Clust-UGC0002132 | 4591 | 3 | 1 | 0 | 0 |
| UGC0007894 | Clust-UGC0002132 | 3601 | 4 | 1 | 0 | 0 |
| UGC0007891 | Clust-UGC0002132 | 3398 | 2 | 1 | 0 | 0 |
| UGC0010173 | Clust-UGC0002132 | 2112 | 1 | 0 | 0 | 0 |
| UGC0008300 | Clust-UGC0002132 | 1043 | 1 | 0 | 0 | 0 |
| UGC0009307 | Clust-UGC0002132 | 278 | 1 | 0 | 0 | 0 |
| UGC0008491 | Clust-UGC0002132 | 1637 | 1 | 0 | 0 | 0 |
| UGC0013167 | Clust-UGC0002132 | 306 | 1 | 0 | 0 | 0 |
| UGC0007912 | Clust-UGC0002132 | 7171 | 5 | 1 | 0 | 1 |
| UGC0007916 | Clust-UGC0002132 | 694 | 1 | 0 | 0 | 0 |
| UGC0009187 | Clust-UGC0002132 | 7025 | 7 | 1 | 0 | 1 |
| UGC0012603 | Clust-UGC0002132 | 8511 | 7 | 1 | 0 | 1 |
| UGC0012834 | Clust-UGC0002132 | 4407 | 3 | 1 | 0 | 1 |
| UGC0012336 | Clust-UGC0002132 | 633 | 1 | 0 | 0 | 0 |
| UGC0004750 | Clust-UGC0002132 | 322 | 1 | 0 | 0 | 0 |
| UGC0013215 | Clust-UGC0002132 | 7323 | 3 | 1 | 0 | 1 |
| UGC0004134 | Clust-UGC0002132 | 1462 | 1 | 0 | 0 | 0 |
| UGC0007011 | Clust-UGC0002132 | 2550 | 2 | 1 | 0 | 1 |
| UGC0002132 | Clust-UGC0002132 | 3533 | 3 | 1 | 0 | 1 |
| UGC0012870 | Clust-UGC0002132 | 8799 | 4 | 1 | 0 | 1 |
| UGC0008173 | Clust-UGC0002132 | 1181 | 1 | 0 | 0 | 0 |
| UGC0007991 | Clust-UGC0002132 | 287 | 1 | 0 | 0 | 0 |
| UGC0011990 | Clust-UGC0002132 | 3381 | 1 | 0 | 0 | 0 |
| UGC0008922 | Clust-UGC0002132 | 1280 | 1 | 0 | 0 | 0 |
| UGC0009251 | Clust-UGC0002132 | 10451 | 12 | 1 | 0 | 1 |
| UGC0009075 | Clust-UGC0002132 | 3328 | 1 | 0 | 0 | 0 |
| UGC0009072 | Clust-UGC0002132 | 2999 | 1 | 0 | 0 | 0 |
| UGC0007788 | Clust-UGC0002132 | 3077 | 3 | 1 | 0 | 1 |
| UGC0010834 | Clust-UGC0002132 | 1380 | 1 | 0 | 0 | 0 |
| UGC0012896 | Clust-UGC0002132 | 230 | 1 | 0 | 0 | 0 |
| UGC0008903 | Clust-UGC0002132 | 850 | 1 | 0 | 0 | 0 |
| UGC0012976 | Clust-UGC0002132 | 6549 | 3 | 1 | 0 | 1 |
| UGC0013362 | Clust-UGC0002132 | 554 | 2 | 1 | 0 | 0 |
| UGC0013364 | Clust-UGC0002132 | 2292 | 3 | 1 | 0 | 1 |
| UGC0006084 | Clust-UGC0002132 | 5288 | 3 | 1 | 0 | 1 |
| UGC0004564 | Clust-UGC0002132 | 8193 | 10 | 1 | 0 | 1 |
| UGC0006438 | Clust-UGC0002132 | 5111 | 1 | 0 | 0 | 0 |
| UGC0008581 | Clust-UGC0002132 | 4943 | 3 | 1 | 0 | 1 |
| UGC0009984 | Clust-UGC0002132 | 1319 | 1 | 0 | 0 | 0 |
| UGC0009980 | Clust-UGC0002132 | 752 | 1 | 0 | 0 | 0 |
| UGC0009407 | Clust-UGC0002132 | 462 | 1 | 0 | 0 | 0 |
| UGC0012589 | Clust-UGC0002132 | 5419 | 3 | 1 | 0 | 1 |
| UGC0003795 | Clust-UGC0002132 | 827 | 1 | 0 | 0 | 0 |
| UGC0009235 | Clust-UGC0002132 | 6818 | 4 | 1 | 0 | 1 |
| UGC0008942 | Clust-UGC0002132 | 6665 | 3 | 1 | 0 | 1 |
| UGC0009094 | Clust-UGC0002132 | 3187 | 2 | 1 | 0 | 1 |
| UGC0004623 | Clust-UGC0002132 | 7291 | 4 | 1 | 1 | 1 |

| | | | | | | |
|---|---|---|---|---|---|---|
| UGC0004627 | Clust-UGC0002132 | 4136 | 4 | 1 | 0 | 1 |
| UGC0008424 | Clust-UGC0002132 | 417 | 1 | 0 | 0 | 0 |
| UGC0007955 | Clust-UGC0002132 | 2206 | 1 | 0 | 0 | 0 |
| UGC0007947 | Clust-UGC0002132 | 2435 | 1 | 0 | 0 | 0 |
| UGC0009115 | Clust-UGC0002132 | 1127 | 1 | 0 | 0 | 0 |
| UGC0008188 | Clust-UGC0002132 | 1027 | 1 | 0 | 0 | 0 |
| UGC0012205 | Clust-UGC0002132 | 5590 | 2 | 1 | 0 | 0 |
| UGC0005834 | Clust-UGC0002132 | 3670 | 1 | 0 | 0 | 0 |
| UGC0013114 | Clust-UGC0002132 | 1289 | 1 | 0 | 0 | 0 |
| UGC0003731 | Clust-UGC0002132 | 493 | 1 | 0 | 0 | 0 |
| UGC0010971 | Clust-UGC0002132 | 3624 | 3 | 1 | 0 | 1 |
| UGC0009330 | Clust-UGC0002132 | 3492 | 3 | 1 | 0 | 1 |
| UGC0003418 | Clust-UGC0002132 | 1065 | 1 | 0 | 0 | 0 |
| UGC0013136 | Clust-UGC0002132 | 267 | 1 | 0 | 0 | 0 |
| UGC0007766 | Clust-UGC0002132 | 3604 | 5 | 1 | 0 | 0 |
| UGC0008338 | Clust-UGC0002132 | 383 | 1 | 0 | 0 | 0 |
| UGC0008332 | Clust-UGC0002132 | 682 | 1 | 0 | 0 | 0 |
| UGC0010167 | Clust-UGC0002133 | 2473 | 1 | 0 | 0 | 0 |
| UGC0006064 | Clust-UGC0002133 | 1428 | 1 | 0 | 0 | 0 |
| UGC0012172 | Clust-UGC0002133 | 4102 | 2 | 1 | 0 | 1 |
| UGC0010282 | Clust-UGC0002133 | 2602 | 3 | 1 | 0 | 1 |
| UGC0003764 | Clust-UGC0002133 | 229 | 1 | 0 | 0 | 0 |
| UGC0003238 | Clust-UGC0002133 | 2835 | 4 | 1 | 0 | 1 |
| UGC0006897 | Clust-UGC0002133 | 2270 | 2 | 1 | 0 | 1 |
| UGC0012964 | Clust-UGC0002133 | 5672 | 9 | 1 | 0 | 1 |
| UGC0007108 | Clust-UGC0002133 | 472 | 1 | 0 | 0 | 0 |
| UGC0006679 | Clust-UGC0002133 | 1023 | 2 | 1 | 0 | 1 |
| UGC0006335 | Clust-UGC0002133 | 4713 | 5 | 1 | 0 | 1 |
| UGC0009687 | Clust-UGC0002133 | 660 | 1 | 0 | 0 | 0 |
| UGC0009189 | Clust-UGC0002133 | 124 | 1 | 0 | 0 | 0 |
| UGC0008859 | Clust-UGC0002133 | 2285 | 3 | 1 | 0 | 1 |
| UGC0010988 | Clust-UGC0002133 | 1281 | 3 | 1 | 0 | 1 |
| UGC0003546 | Clust-UGC0002133 | 3635 | 4 | 1 | 0 | 1 |
| UGC0002133 | Clust-UGC0002133 | 1456 | 5 | 1 | 0 | 1 |
| UGC0010003 | Clust-UGC0002133 | 360 | 1 | 0 | 0 | 0 |
| UGC0007335 | Clust-UGC0002133 | 1657 | 1 | 0 | 0 | 0 |
| UGC0010492 | Clust-UGC0002133 | 5222 | 8 | 1 | 1 | 1 |
| UGC0003276 | Clust-UGC0002133 | 8357 | 20 | 1 | 0 | 1 |
| UGC0010858 | Clust-UGC0002133 | 1284 | 1 | 0 | 0 | 0 |
| UGC0008678 | Clust-UGC0002133 | 350 | 1 | 0 | 0 | 0 |
| UGC0006801 | Clust-UGC0002133 | 4152 | 4 | 1 | 0 | 1 |
| UGC0008231 | Clust-UGC0002133 | 887 | 3 | 1 | 0 | 1 |
| UGC0002134 | Clust-UGC0002134 | 1050 | 1 | 0 | 0 | 0 |
| UGC0002136 | Clust-UGC0002136 | 5935 | 22 | 1 | 0 | 1 |
| UGC0011563 | Clust-UGC0002139 | 12463 | 86 | 1 | 0 | 1 |
| UGC0013171 | Clust-UGC0002139 | 2367 | 2 | 1 | 0 | 0 |
| UGC0004593 | Clust-UGC0002139 | 10060 | 40 | 1 | 0 | 1 |
| UGC0005356 | Clust-UGC0002139 | 2462 | 2 | 1 | 0 | 1 |
| UGC0011902 | Clust-UGC0002139 | 3047 | 1 | 0 | 0 | 0 |
| UGC0003767 | Clust-UGC0002139 | 421 | 1 | 0 | 0 | 0 |
| UGC0012632 | Clust-UGC0002139 | 2264 | 2 | 1 | 0 | 0 |
| UGC0010095 | Clust-UGC0002139 | 4104 | 4 | 1 | 0 | 1 |
| UGC0009544 | Clust-UGC0002139 | 2725 | 4 | 1 | 0 | 0 |
| UGC0008702 | Clust-UGC0002139 | 534 | 1 | 0 | 0 | 0 |
| UGC0011449 | Clust-UGC0002139 | 10843 | 32 | 1 | 0 | 1 |
| UGC0007635 | Clust-UGC0002139 | 3139 | 1 | 0 | 0 | 0 |
| UGC0012797 | Clust-UGC0002139 | 2884 | 3 | 1 | 0 | 1 |
| UGC0005617 | Clust-UGC0002139 | 4223 | 1 | 0 | 0 | 0 |
| UGC0007341 | Clust-UGC0002139 | 993 | 2 | 1 | 0 | 0 |
| UGC0008524 | Clust-UGC0002139 | 1357 | 1 | 0 | 0 | 0 |
| UGC0002324 | Clust-UGC0002139 | 3318 | 2 | 1 | 0 | 1 |
| UGC0008936 | Clust-UGC0002139 | 178 | 1 | 0 | 0 | 0 |
| UGC0013052 | Clust-UGC0002139 | 5924 | 7 | 1 | 2 | 1 |
| UGC0009066 | Clust-UGC0002139 | 5453 | 6 | 1 | 0 | 1 |
| UGC0007790 | Clust-UGC0002139 | 4330 | 3 | 1 | 0 | 1 |
| UGC0004535 | Clust-UGC0002139 | 4924 | 7 | 1 | 0 | 1 |
| UGC0005958 | Clust-UGC0002139 | 62 | 1 | 0 | 0 | 0 |
| UGC0004945 | Clust-UGC0002139 | 4411 | 2 | 1 | 0 | 1 |
| UGC0005541 | Clust-UGC0002139 | 736 | 1 | 0 | 0 | 0 |
| UGC0005547 | Clust-UGC0002139 | 427 | 1 | 0 | 0 | 0 |
| UGC0006894 | Clust-UGC0002139 | 6284 | 8 | 1 | 0 | 1 |
| UGC0013375 | Clust-UGC0002139 | 129 | 2 | 1 | 0 | 0 |
| UGC0012770 | Clust-UGC0002139 | 2795 | 2 | 1 | 0 | 1 |
| UGC0010196 | Clust-UGC0002139 | 497 | 2 | 1 | 1 | 0 |
| UGC0005736 | Clust-UGC0002139 | 4041 | 8 | 1 | 0 | 1 |
| UGC0008938 | Clust-UGC0002139 | 7124 | 12 | 1 | 0 | 1 |
| UGC0011820 | Clust-UGC0002139 | 4641 | 3 | 1 | 0 | 0 |
| UGC0008430 | Clust-UGC0002139 | 1356 | 1 | 0 | 0 | 0 |
| UGC0008045 | Clust-UGC0002139 | 129 | 1 | 0 | 0 | 0 |

| | | | | | | |
|---|---|---|---|---|---|---|
| UGC0009167 | Clust-UGC0002139 | 3584 | 3 | 1 | 0 | 1 |
| UGC0008363 | Clust-UGC0002139 | 2055 | 1 | 0 | 0 | 0 |
| UGC0004677 | Clust-UGC0002139 | 19425 | 164 | 1 | 0 | 1 |
| UGC0010351 | Clust-UGC0002139 | 62 | 1 | 0 | 0 | 0 |
| UGC0007978 | Clust-UGC0002139 | 938 | 1 | 0 | 0 | 0 |
| UGC0008207 | Clust-UGC0002139 | 2522 | 2 | 1 | 0 | 1 |
| UGC0008340 | Clust-UGC0002139 | 797 | 1 | 0 | 0 | 0 |
| UGC0002139 | Clust-UGC0002139 | 4496 | 3 | 1 | 0 | 1 |
| UGC0010960 | Clust-UGC0002139 | 374 | 2 | 1 | 0 | 0 |
| UGC0012983 | Clust-UGC0002139 | 6107 | 9 | 1 | 0 | 1 |
| UGC0010580 | Clust-UGC0002139 | 2044 | 2 | 1 | 0 | 1 |
| UGC0008796 | Clust-UGC0002139 | 3544 | 1 | 0 | 0 | 0 |
| UGC0008876 | Clust-UGC0002139 | 6824 | 14 | 1 | 0 | 1 |
| UGC0011956 | Clust-UGC0002139 | 296 | 1 | 0 | 0 | 0 |
| UGC0008497 | Clust-UGC0002139 | 2483 | 3 | 1 | 0 | 0 |
| UGC0007445 | Clust-UGC0002139 | 922 | 1 | 0 | 0 | 0 |
| UGC0006519 | Clust-UGC0002139 | 2001 | 1 | 0 | 0 | 0 |
| UGC0010086 | Clust-UGC0002139 | 3132 | 1 | 0 | 0 | 0 |
| UGC0005663 | Clust-UGC0002139 | 3957 | 3 | 1 | 0 | 1 |
| UGC0005415 | Clust-UGC0002139 | 2743 | 1 | 0 | 0 | 0 |
| UGC0005602 | Clust-UGC0002139 | 3165 | 1 | 0 | 0 | 0 |
| UGC0007037 | Clust-UGC0002139 | 2526 | 1 | 0 | 0 | 0 |
| UGC0008529 | Clust-UGC0002139 | 712 | 1 | 0 | 0 | 0 |
| UGC0008647 | Clust-UGC0002139 | 1129 | 1 | 0 | 0 | 0 |
| UGC0010579 | Clust-UGC0002139 | 1573 | 3 | 1 | 1 | 0 |
| UGC0011438 | Clust-UGC0002139 | 2384 | 3 | 1 | 0 | 0 |
| UGC0010219 | Clust-UGC0002139 | 599 | 2 | 1 | 0 | 0 |
| UGC0005680 | Clust-UGC0002139 | 266 | 1 | 0 | 0 | 0 |
| UGC0005519 | Clust-UGC0002139 | 768 | 1 | 0 | 0 | 0 |
| UGC0006608 | Clust-UGC0002139 | 2295 | 1 | 0 | 0 | 0 |
| UGC0012911 | Clust-UGC0002139 | 4464 | 4 | 1 | 0 | 1 |
| UGC0009618 | Clust-UGC0002139 | 1259 | 1 | 0 | 0 | 0 |
| UGC0007707 | Clust-UGC0002139 | 8560 | 19 | 1 | 0 | 1 |
| UGC0011685 | Clust-UGC0002139 | 3670 | 3 | 1 | 0 | 1 |
| UGC0003609 | Clust-UGC0002139 | 2454 | 1 | 0 | 0 | 0 |
| UGC0007724 | Clust-UGC0002139 | 8190 | 30 | 1 | 0 | 1 |
| UGC0008667 | Clust-UGC0002139 | 102 | 1 | 0 | 0 | 0 |
| UGC0012202 | Clust-UGC0002139 | 6152 | 12 | 1 | 0 | 1 |
| UGC0005327 | Clust-UGC0002139 | 2742 | 1 | 0 | 0 | 0 |
| UGC0008216 | Clust-UGC0002139 | 8824 | 38 | 1 | 0 | 1 |
| UGC0008372 | Clust-UGC0002139 | 668 | 1 | 0 | 0 | 0 |
| UGC0012195 | Clust-UGC0002139 | 9171 | 38 | 1 | 0 | 1 |
| UGC0010594 | Clust-UGC0002139 | 2463 | 3 | 1 | 0 | 1 |
| UGC0009339 | Clust-UGC0002139 | 5345 | 7 | 1 | 0 | 1 |
| UGC0006828 | Clust-UGC0002139 | 3946 | 3 | 1 | 0 | 1 |
| UGC0008504 | Clust-UGC0002139 | 415 | 1 | 0 | 0 | 0 |
| UGC0006373 | Clust-UGC0002139 | 5700 | 7 | 1 | 0 | 1 |
| UGC0007207 | Clust-UGC0002142 | 5343 | 2 | 1 | 0 | 0 |
| UGC0006550 | Clust-UGC0002142 | 2869 | 1 | 0 | 0 | 0 |
| UGC0004166 | Clust-UGC0002142 | 1203 | 1 | 0 | 0 | 0 |
| UGC0004162 | Clust-UGC0002142 | 2460 | 1 | 0 | 0 | 0 |
| UGC0008270 | Clust-UGC0002142 | 857 | 1 | 0 | 0 | 0 |
| UGC0005373 | Clust-UGC0002142 | 3688 | 2 | 1 | 0 | 1 |
| UGC0008299 | Clust-UGC0002142 | 3490 | 1 | 0 | 0 | 0 |
| UGC0002294 | Clust-UGC0002142 | 6456 | 5 | 1 | 0 | 0 |
| UGC0012323 | Clust-UGC0002142 | 1219 | 1 | 0 | 0 | 0 |
| UGC0011674 | Clust-UGC0002142 | 4380 | 2 | 1 | 0 | 1 |
| UGC0012796 | Clust-UGC0002142 | 1933 | 1 | 0 | 0 | 0 |
| UGC0008889 | Clust-UGC0002142 | 5396 | 2 | 1 | 0 | 1 |
| UGC0007286 | Clust-UGC0002142 | 5805 | 2 | 1 | 0 | 1 |
| UGC0006739 | Clust-UGC0002142 | 559 | 1 | 0 | 0 | 0 |
| UGC0007345 | Clust-UGC0002142 | 4778 | 2 | 1 | 0 | 0 |
| UGC0008516 | Clust-UGC0002142 | 1895 | 1 | 0 | 0 | 0 |
| UGC0008518 | Clust-UGC0002142 | 1607 | 1 | 0 | 0 | 0 |
| UGC0007403 | Clust-UGC0002142 | 4531 | 1 | 0 | 0 | 0 |
| UGC0013242 | Clust-UGC0002142 | 4287 | 3 | 1 | 2 | 0 |
| UGC0003232 | Clust-UGC0002142 | 3224 | 4 | 1 | 0 | 1 |
| UGC0007367 | Clust-UGC0002142 | 2413 | 1 | 0 | 0 | 0 |
| UGC0013053 | Clust-UGC0002142 | 7769 | 8 | 1 | 0 | 1 |
| UGC0010826 | Clust-UGC0002142 | 12507 | 25 | 1 | 0 | 1 |
| UGC0010827 | Clust-UGC0002142 | 6814 | 1 | 0 | 0 | 0 |
| UGC0012553 | Clust-UGC0002142 | 3681 | 3 | 1 | 0 | 1 |
| UGC0007143 | Clust-UGC0002142 | 4016 | 2 | 1 | 0 | 1 |
| UGC0007326 | Clust-UGC0002142 | 906 | 1 | 0 | 0 | 0 |
| UGC0007462 | Clust-UGC0002142 | 4302 | 1 | 0 | 0 | 0 |
| UGC0009435 | Clust-UGC0002142 | 96 | 1 | 0 | 0 | 0 |
| UGC0007129 | Clust-UGC0002142 | 759 | 1 | 0 | 0 | 0 |
| UGC0007693 | Clust-UGC0002142 | 4043 | 2 | 1 | 0 | 0 |
| UGC0006614 | Clust-UGC0002142 | 358 | 1 | 0 | 0 | 0 |

| | | | | | | |
|---|---|---|---|---|---|---|
| UGC0004573 | Clust-UGC0002142 | 10199 | 11 | 1 | 0 | 1 |
| UGC0008382 | Clust-UGC0002142 | 457 | 1 | 0 | 0 | 0 |
| UGC0004659 | Clust-UGC0002142 | 1661 | 1 | 0 | 0 | 0 |
| UGC0013335 | Clust-UGC0002142 | 421 | 1 | 0 | 0 | 0 |
| UGC0009164 | Clust-UGC0002142 | 3254 | 5 | 1 | 0 | 1 |
| UGC0011208 | Clust-UGC0002142 | 7970 | 4 | 1 | 0 | 1 |
| UGC0007526 | Clust-UGC0002142 | 8726 | 3 | 1 | 0 | 1 |
| UGC0003254 | Clust-UGC0002142 | 10171 | 12 | 1 | 0 | 1 |
| UGC0003283 | Clust-UGC0002142 | 456 | 1 | 0 | 0 | 0 |
| UGC0007758 | Clust-UGC0002142 | 6214 | 8 | 1 | 0 | 1 |
| UGC0007759 | Clust-UGC0002142 | 2554 | 1 | 0 | 0 | 0 |
| UGC0006830 | Clust-UGC0002142 | 8175 | 7 | 1 | 1 | 1 |
| UGC0003265 | Clust-UGC0002142 | 9870 | 8 | 1 | 0 | 1 |
| UGC0012047 | Clust-UGC0002142 | 2754 | 1 | 0 | 0 | 0 |
| UGC0012049 | Clust-UGC0002142 | 3832 | 1 | 0 | 0 | 0 |
| UGC0006144 | Clust-UGC0002142 | 1391 | 1 | 0 | 0 | 0 |
| UGC0007899 | Clust-UGC0002142 | 7469 | 5 | 1 | 0 | 1 |
| UGC0013145 | Clust-UGC0002142 | 1656 | 2 | 1 | 0 | 0 |
| UGC0008877 | Clust-UGC0002142 | 4596 | 1 | 0 | 0 | 0 |
| UGC0007211 | Clust-UGC0002142 | 1945 | 1 | 0 | 0 | 0 |
| UGC0007210 | Clust-UGC0002142 | 2616 | 2 | 1 | 1 | 0 |
| UGC0012608 | Clust-UGC0002142 | 6769 | 3 | 1 | 0 | 1 |
| UGC0013184 | Clust-UGC0002142 | 4778 | 3 | 1 | 1 | 1 |
| UGC0007016 | Clust-UGC0002142 | 6442 | 3 | 1 | 1 | 0 |
| UGC0011899 | Clust-UGC0002142 | 4504 | 3 | 1 | 1 | 1 |
| UGC0007290 | Clust-UGC0002142 | 8928 | 5 | 1 | 0 | 1 |
| UGC0003268 | Clust-UGC0002142 | 8109 | 3 | 1 | 0 | 0 |
| UGC0007965 | Clust-UGC0002142 | 2228 | 1 | 0 | 0 | 0 |
| UGC0006696 | Clust-UGC0002142 | 5684 | 2 | 1 | 0 | 0 |
| UGC0007196 | Clust-UGC0002142 | 2242 | 2 | 1 | 0 | 0 |
| UGC0013069 | Clust-UGC0002142 | 16801 | 71 | 1 | 0 | 1 |
| UGC0013088 | Clust-UGC0002142 | 2563 | 2 | 1 | 1 | 0 |
| UGC0007417 | Clust-UGC0002142 | 4163 | 4 | 1 | 0 | 1 |
| UGC0010757 | Clust-UGC0002142 | 2086 | 1 | 0 | 0 | 0 |
| UGC0008752 | Clust-UGC0002142 | 1538 | 1 | 0 | 0 | 0 |
| UGC0007314 | Clust-UGC0002142 | 6617 | 3 | 1 | 1 | 1 |
| UGC0008929 | Clust-UGC0002142 | 3758 | 1 | 0 | 0 | 0 |
| UGC0013049 | Clust-UGC0002142 | 10074 | 7 | 1 | 0 | 1 |
| UGC0009982 | Clust-UGC0002142 | 5874 | 4 | 1 | 0 | 1 |
| UGC0008611 | Clust-UGC0002142 | 6475 | 3 | 1 | 0 | 1 |
| UGC0007393 | Clust-UGC0002142 | 1708 | 1 | 0 | 0 | 0 |
| UGC0012601 | Clust-UGC0002142 | 8567 | 5 | 1 | 0 | 1 |
| UGC0003249 | Clust-UGC0002142 | 7817 | 6 | 1 | 0 | 1 |
| UGC0007940 | Clust-UGC0002142 | 2357 | 2 | 1 | 0 | 0 |
| UGC0002142 | Clust-UGC0002142 | 5476 | 2 | 1 | 0 | 0 |
| UGC0003606 | Clust-UGC0002142 | 11631 | 13 | 1 | 0 | 1 |
| UGC0007720 | Clust-UGC0002142 | 7082 | 7 | 1 | 0 | 1 |
| UGC0010918 | Clust-UGC0002142 | 821 | 1 | 0 | 0 | 0 |
| UGC0008370 | Clust-UGC0002142 | 1728 | 1 | 0 | 0 | 0 |
| UGC0003362 | Clust-UGC0002142 | 323 | 1 | 0 | 0 | 0 |
| UGC0007575 | Clust-UGC0002142 | 5446 | 1 | 0 | 0 | 0 |
| UGC0003263 | Clust-UGC0002142 | 10564 | 11 | 1 | 0 | 1 |
| UGC0011963 | Clust-UGC0002142 | 743 | 1 | 0 | 0 | 0 |
| UGC0007088 | Clust-UGC0002142 | 3112 | 2 | 1 | 0 | 0 |
| UGC0004148 | Clust-UGC0002142 | 2100 | 1 | 0 | 0 | 0 |
| UGC0005785 | Clust-UGC0002144 | 5400 | 7 | 1 | 1 | 1 |
| UGC0002352 | Clust-UGC0002144 | 497 | 1 | 0 | 0 | 0 |
| UGC0006588 | Clust-UGC0002144 | 880 | 1 | 0 | 0 | 0 |
| UGC0004494 | Clust-UGC0002144 | 3933 | 4 | 1 | 0 | 1 |
| UGC0007146 | Clust-UGC0002144 | 2968 | 3 | 1 | 0 | 1 |
| UGC0012595 | Clust-UGC0002144 | 2607 | 3 | 1 | 0 | 1 |
| UGC0002144 | Clust-UGC0002144 | 1607 | 3 | 1 | 0 | 1 |
| UGC0005842 | Clust-UGC0002144 | 4735 | 2 | 1 | 0 | 1 |
| UGC0005791 | Clust-UGC0002144 | 5996 | 8 | 1 | 0 | 1 |
| UGC0008267 | Clust-UGC0002144 | 275 | 1 | 0 | 0 | 0 |
| UGC0003750 | Clust-UGC0002144 | 54 | 1 | 0 | 0 | 0 |
| UGC0002286 | Clust-UGC0002144 | 494 | 1 | 0 | 0 | 0 |
| UGC0010859 | Clust-UGC0002144 | 841 | 1 | 0 | 0 | 0 |
| UGC0008927 | Clust-UGC0002144 | 1315 | 1 | 0 | 0 | 0 |
| UGC0002161 | Clust-UGC0002144 | 3904 | 3 | 1 | 0 | 1 |
| UGC0004789 | Clust-UGC0002147 | 10372 | 139 | 1 | 0 | 1 |
| UGC0012368 | Clust-UGC0002147 | 19740 | 132 | 1 | 0 | 1 |
| UGC0013176 | Clust-UGC0002147 | 7286 | 17 | 1 | 0 | 1 |
| UGC0011679 | Clust-UGC0002147 | 12876 | 105 | 1 | 0 | 1 |
| UGC0011655 | Clust-UGC0002147 | 16764 | 161 | 1 | 0 | 1 |
| UGC0002234 | Clust-UGC0002147 | 12027 | 69 | 1 | 0 | 1 |
| UGC0002238 | Clust-UGC0002147 | 17008 | 162 | 1 | 0 | 1 |
| UGC0002215 | Clust-UGC0002147 | 217 | 1 | 0 | 0 | 0 |
| UGC0002212 | Clust-UGC0002147 | 22187 | 186 | 1 | 0 | 1 |

| | | | | | | |
|---|---|---|---|---|---|---|
| UGC0002251 | Clust-UGC0002147 | 8500 | 88 | 1 | 0 | 1 |
| UGC0003341 | Clust-UGC0002147 | 18724 | 152 | 1 | 0 | 1 |
| UGC0002147 | Clust-UGC0002147 | 22610 | 183 | 1 | 0 | 1 |
| UGC0012020 | Clust-UGC0002147 | 5414 | 21 | 1 | 1 | 1 |
| UGC0013101 | Clust-UGC0002147 | 15050 | 95 | 1 | 0 | 1 |
| UGC0013312 | Clust-UGC0002147 | 11637 | 81 | 1 | 0 | 1 |
| UGC0010969 | Clust-UGC0002147 | 3958 | 10 | 1 | 0 | 1 |
| UGC0002270 | Clust-UGC0002147 | 1474 | 2 | 1 | 0 | 0 |
| UGC0002259 | Clust-UGC0002147 | 8067 | 23 | 1 | 0 | 1 |
| UGC0002302 | Clust-UGC0002147 | 11410 | 30 | 1 | 0 | 1 |
| UGC0004798 | Clust-UGC0002147 | 19000 | 208 | 1 | 0 | 1 |
| UGC0012359 | Clust-UGC0002147 | 15453 | 156 | 1 | 0 | 1 |
| UGC0004775 | Clust-UGC0002147 | 6536 | 35 | 1 | 0 | 1 |
| UGC0013186 | Clust-UGC0002147 | 7706 | 7 | 1 | 0 | 1 |
| UGC0002280 | Clust-UGC0002147 | 16156 | 155 | 1 | 0 | 1 |
| UGC0002253 | Clust-UGC0002147 | 13379 | 53 | 1 | 0 | 1 |
| UGC0002220 | Clust-UGC0002147 | 30539 | 254 | 1 | 0 | 1 |
| UGC0002222 | Clust-UGC0002147 | 23501 | 210 | 1 | 0 | 1 |
| UGC0002228 | Clust-UGC0002147 | 23392 | 221 | 1 | 0 | 1 |
| UGC0011998 | Clust-UGC0002147 | 9104 | 56 | 1 | 0 | 1 |
| UGC0002208 | Clust-UGC0002147 | 3111 | 15 | 1 | 0 | 1 |
| UGC0002240 | Clust-UGC0002147 | 19324 | 188 | 1 | 0 | 1 |
| UGC0002249 | Clust-UGC0002147 | 7167 | 48 | 1 | 0 | 1 |
| UGC0002246 | Clust-UGC0002147 | 24115 | 206 | 1 | 1 | 1 |
| UGC0002243 | Clust-UGC0002147 | 14689 | 151 | 1 | 0 | 1 |
| UGC0013085 | Clust-UGC0002147 | 8761 | 27 | 1 | 0 | 1 |
| UGC0003378 | Clust-UGC0002147 | 25307 | 225 | 1 | 0 | 1 |
| UGC0002261 | Clust-UGC0002147 | 13058 | 100 | 1 | 0 | 1 |
| UGC0002262 | Clust-UGC0002147 | 12737 | 137 | 1 | 0 | 1 |
| UGC0013323 | Clust-UGC0002147 | 9289 | 61 | 1 | 0 | 1 |
| UGC0013328 | Clust-UGC0002147 | 9295 | 44 | 1 | 0 | 1 |
| UGC0010931 | Clust-UGC0002147 | 7829 | 13 | 1 | 0 | 1 |
| UGC0008377 | Clust-UGC0002147 | 1574 | 2 | 1 | 0 | 1 |
| UGC0012439 | Clust-UGC0002147 | 14055 | 117 | 1 | 0 | 1 |
| UGC0011944 | Clust-UGC0002147 | 13115 | 74 | 1 | 0 | 1 |
| UGC0008885 | Clust-UGC0002148 | 4858 | 2 | 1 | 0 | 1 |
| UGC0008004 | Clust-UGC0002148 | 7095 | 4 | 1 | 0 | 1 |
| UGC0004554 | Clust-UGC0002148 | 7433 | 8 | 1 | 0 | 1 |
| UGC0002148 | Clust-UGC0002148 | 3193 | 2 | 1 | 0 | 0 |
| UGC0005521 | Clust-UGC0002148 | 1157 | 1 | 0 | 0 | 0 |
| UGC0006871 | Clust-UGC0002148 | 5060 | 2 | 1 | 0 | 1 |
| UGC0008366 | Clust-UGC0002148 | 768 | 1 | 0 | 0 | 0 |
| UGC0010132 | Clust-UGC0002148 | 6933 | 2 | 1 | 1 | 0 |
| UGC0010117 | Clust-UGC0002148 | 3009 | 1 | 0 | 0 | 0 |
| UGC0008240 | Clust-UGC0002148 | 989 | 1 | 0 | 0 | 0 |
| UGC0002367 | Clust-UGC0002148 | 665 | 2 | 1 | 1 | 0 |
| UGC0010216 | Clust-UGC0002148 | 489 | 1 | 0 | 0 | 0 |
| UGC0010499 | Clust-UGC0002148 | 4208 | 1 | 0 | 0 | 0 |
| UGC0010124 | Clust-UGC0002148 | 12558 | 13 | 1 | 1 | 1 |
| UGC0005339 | Clust-UGC0002148 | 907 | 1 | 0 | 0 | 0 |
| UGC0008807 | Clust-UGC0002148 | 2451 | 1 | 0 | 0 | 0 |
| UGC0006556 | Clust-UGC0002151 | 900 | 1 | 0 | 0 | 0 |
| UGC0008272 | Clust-UGC0002151 | 1101 | 1 | 0 | 0 | 0 |
| UGC0008316 | Clust-UGC0002151 | 2090 | 1 | 0 | 0 | 0 |
| UGC0008865 | Clust-UGC0002151 | 3478 | 1 | 0 | 0 | 0 |
| UGC0005371 | Clust-UGC0002151 | 2820 | 2 | 1 | 0 | 1 |
| UGC0007042 | Clust-UGC0002151 | 214 | 1 | 0 | 0 | 0 |
| UGC0010464 | Clust-UGC0002151 | 4085 | 1 | 0 | 0 | 0 |
| UGC0007063 | Clust-UGC0002151 | 906 | 1 | 0 | 0 | 0 |
| UGC0007656 | Clust-UGC0002151 | 6382 | 6 | 1 | 0 | 1 |
| UGC0007001 | Clust-UGC0002151 | 4011 | 1 | 0 | 0 | 0 |
| UGC0006719 | Clust-UGC0002151 | 899 | 1 | 0 | 0 | 0 |
| UGC0007285 | Clust-UGC0002151 | 4707 | 2 | 1 | 0 | 1 |
| UGC0005909 | Clust-UGC0002151 | 3833 | 2 | 1 | 0 | 1 |
| UGC0008511 | Clust-UGC0002151 | 2498 | 1 | 0 | 0 | 0 |
| UGC0007409 | Clust-UGC0002151 | 2530 | 1 | 0 | 0 | 0 |
| UGC0002957 | Clust-UGC0002151 | 1785 | 1 | 0 | 0 | 0 |
| UGC0008533 | Clust-UGC0002151 | 1153 | 1 | 0 | 0 | 0 |
| UGC0009163 | Clust-UGC0002151 | 2312 | 1 | 0 | 0 | 0 |
| UGC0006876 | Clust-UGC0002151 | 1432 | 1 | 0 | 0 | 0 |
| UGC0006979 | Clust-UGC0002151 | 3509 | 3 | 1 | 0 | 1 |
| UGC0006790 | Clust-UGC0002151 | 3299 | 2 | 1 | 0 | 1 |
| UGC0009205 | Clust-UGC0002151 | 5383 | 3 | 1 | 0 | 1 |
| UGC0008530 | Clust-UGC0002151 | 1672 | 1 | 0 | 0 | 0 |
| UGC0008450 | Clust-UGC0002151 | 746 | 1 | 0 | 0 | 0 |
| UGC0008599 | Clust-UGC0002151 | 3681 | 1 | 0 | 0 | 0 |
| UGC0012021 | Clust-UGC0002151 | 5059 | 4 | 1 | 0 | 1 |
| UGC0008380 | Clust-UGC0002151 | 3642 | 1 | 0 | 0 | 0 |
| UGC0002176 | Clust-UGC0002151 | 6633 | 4 | 1 | 0 | 1 |

| | | | | | | |
|---|---|---|---|---|---|---|
| UGC0009162 | Clust-UGC0002151 | 5486 | 7 | 1 | 0 | 1 |
| UGC0012009 | Clust-UGC0002151 | 1140 | 1 | 0 | 0 | 0 |
| UGC0008412 | Clust-UGC0002151 | 1283 | 1 | 0 | 0 | 0 |
| UGC0012275 | Clust-UGC0002151 | 1513 | 1 | 0 | 0 | 0 |
| UGC0008369 | Clust-UGC0002151 | 1093 | 1 | 0 | 0 | 0 |
| UGC0002151 | Clust-UGC0002151 | 8022 | 6 | 1 | 0 | 1 |
| UGC0010115 | Clust-UGC0002151 | 7162 | 4 | 1 | 1 | 1 |
| UGC0007953 | Clust-UGC0002151 | 4214 | 1 | 0 | 0 | 0 |
| UGC0008329 | Clust-UGC0002151 | 1438 | 1 | 0 | 0 | 0 |
| UGC0010589 | Clust-UGC0002151 | 3301 | 1 | 0 | 0 | 0 |
| UGC0007890 | Clust-UGC0002151 | 6957 | 8 | 1 | 0 | 1 |
| UGC0004158 | Clust-UGC0002151 | 4369 | 3 | 1 | 0 | 1 |
| UGC0007938 | Clust-UGC0002151 | 3380 | 4 | 1 | 0 | 1 |
| UGC0008498 | Clust-UGC0002151 | 2658 | 1 | 0 | 0 | 0 |
| UGC0008265 | Clust-UGC0002151 | 2404 | 1 | 0 | 0 | 0 |
| UGC0010458 | Clust-UGC0002151 | 3033 | 1 | 0 | 0 | 0 |
| UGC0007580 | Clust-UGC0002151 | 8262 | 5 | 1 | 0 | 1 |
| UGC0007974 | Clust-UGC0002151 | 2909 | 1 | 0 | 0 | 0 |
| UGC0007258 | Clust-UGC0002151 | 6079 | 3 | 1 | 0 | 1 |
| UGC0008548 | Clust-UGC0002151 | 1031 | 1 | 0 | 0 | 0 |
| UGC0010085 | Clust-UGC0002151 | 7502 | 4 | 1 | 0 | 1 |
| UGC0012814 | Clust-UGC0002151 | 1895 | 1 | 0 | 0 | 0 |
| UGC0011891 | Clust-UGC0002151 | 15061 | 32 | 1 | 0 | 1 |
| UGC0006838 | Clust-UGC0002151 | 2174 | 1 | 0 | 0 | 0 |
| UGC0011644 | Clust-UGC0002151 | 4540 | 3 | 1 | 0 | 1 |
| UGC0007967 | Clust-UGC0002151 | 4390 | 4 | 1 | 0 | 1 |
| UGC0007195 | Clust-UGC0002151 | 646 | 1 | 0 | 0 | 0 |
| UGC0008692 | Clust-UGC0002151 | 1064 | 1 | 0 | 0 | 0 |
| UGC0003277 | Clust-UGC0002151 | 8021 | 7 | 1 | 0 | 1 |
| UGC0010797 | Clust-UGC0002151 | 783 | 2 | 1 | 0 | 0 |
| UGC0002247 | Clust-UGC0002151 | 12858 | 20 | 1 | 0 | 1 |
| UGC0013082 | Clust-UGC0002151 | 2866 | 1 | 0 | 0 | 0 |
| UGC0007470 | Clust-UGC0002151 | 4236 | 2 | 1 | 0 | 1 |
| UGC0012582 | Clust-UGC0002151 | 9283 | 18 | 1 | 0 | 1 |
| UGC0008582 | Clust-UGC0002151 | 1301 | 1 | 0 | 0 | 0 |
| UGC0006840 | Clust-UGC0002151 | 3912 | 2 | 1 | 0 | 1 |
| UGC0006842 | Clust-UGC0002151 | 4058 | 4 | 1 | 1 | 1 |
| UGC0011145 | Clust-UGC0002151 | 1469 | 2 | 1 | 0 | 1 |
| UGC0009374 | Clust-UGC0002151 | 6280 | 4 | 1 | 0 | 1 |
| UGC0012698 | Clust-UGC0002151 | 6457 | 7 | 1 | 0 | 1 |
| UGC0007571 | Clust-UGC0002151 | 7836 | 3 | 1 | 0 | 1 |
| UGC0007743 | Clust-UGC0002151 | 6860 | 5 | 1 | 0 | 1 |
| UGC0006560 | Clust-UGC0002151 | 1874 | 1 | 0 | 0 | 0 |
| UGC0007889 | Clust-UGC0002151 | 5368 | 4 | 1 | 0 | 1 |
| UGC0007558 | Clust-UGC0002151 | 5499 | 2 | 1 | 0 | 1 |
| UGC0005782 | Clust-UGC0002152 | 6605 | 3 | 1 | 0 | 1 |
| UGC0006457 | Clust-UGC0002152 | 7583 | 10 | 1 | 0 | 1 |
| UGC0006452 | Clust-UGC0002152 | 2207 | 1 | 0 | 0 | 0 |
| UGC0011174 | Clust-UGC0002152 | 2987 | 2 | 1 | 0 | 0 |
| UGC0008271 | Clust-UGC0002152 | 1153 | 1 | 0 | 0 | 0 |
| UGC0008486 | Clust-UGC0002152 | 937 | 1 | 0 | 0 | 0 |
| UGC0005992 | Clust-UGC0002152 | 4595 | 1 | 0 | 0 | 0 |
| UGC0004106 | Clust-UGC0002152 | 374 | 1 | 0 | 0 | 0 |
| UGC0008298 | Clust-UGC0002152 | 351 | 1 | 0 | 0 | 0 |
| UGC0006337 | Clust-UGC0002152 | 4073 | 4 | 1 | 0 | 1 |
| UGC0006292 | Clust-UGC0002152 | 4995 | 1 | 0 | 0 | 0 |
| UGC0006075 | Clust-UGC0002152 | 1884 | 1 | 0 | 0 | 0 |
| UGC0005094 | Clust-UGC0002152 | 2962 | 1 | 0 | 0 | 0 |
| UGC0006013 | Clust-UGC0002152 | 3395 | 1 | 0 | 0 | 0 |
| UGC0007025 | Clust-UGC0002152 | 5842 | 2 | 1 | 0 | 1 |
| UGC0012022 | Clust-UGC0002152 | 4288 | 1 | 0 | 0 | 0 |
| UGC0013224 | Clust-UGC0002152 | 349 | 1 | 0 | 0 | 0 |
| UGC0011790 | Clust-UGC0002152 | 1892 | 1 | 0 | 0 | 0 |
| UGC0007404 | Clust-UGC0002152 | 713 | 1 | 0 | 0 | 0 |
| UGC0013246 | Clust-UGC0002152 | 1396 | 1 | 0 | 0 | 0 |
| UGC0013050 | Clust-UGC0002152 | 3993 | 1 | 0 | 0 | 0 |
| UGC0007160 | Clust-UGC0002152 | 4245 | 2 | 1 | 0 | 1 |
| UGC0010481 | Clust-UGC0002152 | 5236 | 4 | 1 | 1 | 1 |
| UGC0006976 | Clust-UGC0002152 | 4276 | 2 | 1 | 1 | 0 |
| UGC0006400 | Clust-UGC0002152 | 4913 | 2 | 1 | 0 | 0 |
| UGC0009433 | Clust-UGC0002152 | 98 | 1 | 0 | 0 | 0 |
| UGC0003599 | Clust-UGC0002152 | 6214 | 1 | 0 | 0 | 0 |
| UGC0005744 | Clust-UGC0002152 | 3172 | 1 | 0 | 0 | 0 |
| UGC0008042 | Clust-UGC0002152 | 1740 | 1 | 0 | 0 | 0 |
| UGC0003387 | Clust-UGC0002152 | 4225 | 2 | 1 | 0 | 1 |
| UGC0004657 | Clust-UGC0002152 | 6073 | 2 | 1 | 0 | 1 |
| UGC0005845 | Clust-UGC0002152 | 6023 | 1 | 0 | 0 | 0 |
| UGC0002152 | Clust-UGC0002152 | 12925 | 5 | 1 | 1 | 1 |
| UGC0005825 | Clust-UGC0002152 | 8902 | 6 | 1 | 0 | 1 |

| | | | | | | |
|---|---|---|---|---|---|---|
| UGC0007753 | Clust-UGC0002152 | 6140 | 1 | 0 | 0 | 0 |
| UGC0006839 | Clust-UGC0002152 | 8396 | 4 | 1 | 0 | 1 |
| UGC0007567 | Clust-UGC0002152 | 5517 | 3 | 1 | 0 | 1 |
| UGC0006303 | Clust-UGC0002152 | 5302 | 2 | 1 | 0 | 1 |
| UGC0009724 | Clust-UGC0002152 | 939 | 1 | 0 | 0 | 0 |
| UGC0004119 | Clust-UGC0002152 | 1220 | 1 | 0 | 0 | 0 |
| UGC0002347 | Clust-UGC0002152 | 532 | 1 | 0 | 0 | 0 |
| UGC0005761 | Clust-UGC0002152 | 6793 | 3 | 1 | 0 | 1 |
| UGC0005946 | Clust-UGC0002152 | 3369 | 1 | 0 | 0 | 0 |
| UGC0011910 | Clust-UGC0002152 | 6867 | 1 | 0 | 0 | 0 |
| UGC0002289 | Clust-UGC0002152 | 3896 | 1 | 0 | 0 | 0 |
| UGC0006414 | Clust-UGC0002152 | 6674 | 1 | 0 | 0 | 0 |
| UGC0006067 | Clust-UGC0002152 | 1091 | 1 | 0 | 0 | 0 |
| UGC0012108 | Clust-UGC0002152 | 889 | 1 | 0 | 0 | 0 |
| UGC0011456 | Clust-UGC0002152 | 3558 | 2 | 1 | 0 | 0 |
| UGC0009765 | Clust-UGC0002152 | 5855 | 2 | 1 | 0 | 1 |
| UGC0006492 | Clust-UGC0002152 | 1625 | 1 | 0 | 0 | 0 |
| UGC0010025 | Clust-UGC0002152 | 2428 | 1 | 0 | 0 | 0 |
| UGC0012875 | Clust-UGC0002152 | 9086 | 3 | 1 | 0 | 0 |
| UGC0006020 | Clust-UGC0002152 | 2082 | 1 | 0 | 0 | 0 |
| UGC0006640 | Clust-UGC0002152 | 1428 | 2 | 1 | 1 | 0 |
| UGC0007376 | Clust-UGC0002152 | 1645 | 1 | 0 | 0 | 0 |
| UGC0011622 | Clust-UGC0002152 | 8004 | 6 | 1 | 0 | 1 |
| UGC0010747 | Clust-UGC0002152 | 197 | 1 | 0 | 0 | 0 |
| UGC0009250 | Clust-UGC0002152 | 10558 | 5 | 1 | 0 | 1 |
| UGC0009253 | Clust-UGC0002152 | 11876 | 8 | 1 | 0 | 1 |
| UGC0012541 | Clust-UGC0002152 | 2880 | 2 | 1 | 0 | 0 |
| UGC0012191 | Clust-UGC0002152 | 5357 | 4 | 1 | 0 | 1 |
| UGC0005579 | Clust-UGC0002152 | 3445 | 3 | 1 | 0 | 1 |
| UGC0006088 | Clust-UGC0002152 | 1364 | 1 | 0 | 0 | 0 |
| UGC0006439 | Clust-UGC0002152 | 4223 | 4 | 1 | 0 | 1 |
| UGC0009988 | Clust-UGC0002152 | 1779 | 1 | 0 | 0 | 0 |
| UGC0002972 | Clust-UGC0002152 | 2293 | 1 | 0 | 0 | 0 |
| UGC0005533 | Clust-UGC0002152 | 1773 | 1 | 0 | 0 | 0 |
| UGC0010914 | Clust-UGC0002152 | 1644 | 1 | 0 | 0 | 0 |
| UGC0005616 | Clust-UGC0002152 | 3630 | 1 | 0 | 0 | 0 |
| UGC0010599 | Clust-UGC0002152 | 2354 | 1 | 0 | 0 | 0 |
| UGC0004146 | Clust-UGC0002152 | 1313 | 1 | 0 | 0 | 0 |
| UGC0006376 | Clust-UGC0002152 | 5631 | 2 | 1 | 0 | 1 |
| UGC0002231 | Clust-UGC0002156 | 10292 | 30 | 1 | 0 | 1 |
| UGC0008383 | Clust-UGC0002156 | 1773 | 3 | 1 | 0 | 0 |
| UGC0002156 | Clust-UGC0002156 | 4824 | 4 | 1 | 0 | 0 |
| UGC0007666 | Clust-UGC0002156 | 8545 | 27 | 1 | 0 | 1 |
| UGC0002224 | Clust-UGC0002156 | 12493 | 72 | 1 | 0 | 1 |
| UGC0012973 | Clust-UGC0002156 | 3466 | 3 | 1 | 0 | 1 |
| UGC0006549 | Clust-UGC0002156 | 2782 | 5 | 1 | 0 | 1 |
| UGC0006527 | Clust-UGC0002157 | 4003 | 3 | 1 | 0 | 1 |
| UGC0008319 | Clust-UGC0002157 | 1725 | 2 | 1 | 0 | 1 |
| UGC0008866 | Clust-UGC0002157 | 3025 | 1 | 0 | 0 | 0 |
| UGC0007047 | Clust-UGC0002157 | 2094 | 3 | 1 | 0 | 1 |
| UGC0003724 | Clust-UGC0002157 | 740 | 1 | 0 | 0 | 0 |
| UGC0012287 | Clust-UGC0002157 | 636 | 2 | 1 | 1 | 0 |
| UGC0008512 | Clust-UGC0002157 | 130 | 1 | 0 | 0 | 0 |
| UGC0007632 | Clust-UGC0002157 | 6110 | 4 | 1 | 0 | 1 |
| UGC0012861 | Clust-UGC0002157 | 14239 | 26 | 1 | 0 | 1 |
| UGC0011886 | Clust-UGC0002157 | 3811 | 1 | 0 | 0 | 0 |
| UGC0005589 | Clust-UGC0002157 | 3598 | 3 | 1 | 0 | 1 |
| UGC0007183 | Clust-UGC0002157 | 2256 | 1 | 0 | 0 | 0 |
| UGC0005938 | Clust-UGC0002157 | 4557 | 6 | 1 | 0 | 1 |
| UGC0009718 | Clust-UGC0002157 | 2165 | 2 | 1 | 0 | 0 |
| UGC0007446 | Clust-UGC0002157 | 5491 | 6 | 1 | 0 | 1 |
| UGC0009458 | Clust-UGC0002157 | 6230 | 7 | 1 | 0 | 1 |
| UGC0012387 | Clust-UGC0002157 | 2263 | 2 | 1 | 0 | 0 |
| UGC0007469 | Clust-UGC0002157 | 6349 | 3 | 1 | 0 | 1 |
| UGC0010156 | Clust-UGC0002157 | 7904 | 5 | 1 | 1 | 1 |
| UGC0007122 | Clust-UGC0002157 | 1516 | 2 | 1 | 0 | 1 |
| UGC0008593 | Clust-UGC0002157 | 5608 | 4 | 1 | 0 | 1 |
| UGC0008950 | Clust-UGC0002157 | 1843 | 1 | 0 | 0 | 0 |
| UGC0005867 | Clust-UGC0002157 | 7482 | 4 | 1 | 0 | 0 |
| UGC0004803 | Clust-UGC0002157 | 10386 | 10 | 1 | 0 | 1 |
| UGC0007528 | Clust-UGC0002157 | 6644 | 7 | 1 | 0 | 1 |
| UGC0002157 | Clust-UGC0002157 | 1660 | 2 | 1 | 0 | 0 |
| UGC0012685 | Clust-UGC0002157 | 5579 | 4 | 1 | 1 | 1 |
| UGC0009123 | Clust-UGC0002157 | 6014 | 3 | 1 | 0 | 1 |
| UGC0006347 | Clust-UGC0002157 | 2294 | 1 | 0 | 0 | 0 |
| UGC0007545 | Clust-UGC0002157 | 7283 | 4 | 1 | 0 | 1 |
| UGC0006559 | Clust-UGC0002157 | 2958 | 1 | 0 | 0 | 0 |
| UGC0002288 | Clust-UGC0002157 | 968 | 1 | 0 | 0 | 0 |
| UGC0007071 | Clust-UGC0002157 | 3246 | 4 | 1 | 1 | 0 |

| | | | | | | |
|---|---|---|---|---|---|---|
| UGC0007253 | Clust-UGC0002157 | 5600 | 4 | 1 | 0 | 1 |
| UGC0006282 | Clust-UGC0002157 | 1184 | 1 | 0 | 0 | 0 |
| UGC0013025 | Clust-UGC0002157 | 4702 | 2 | 1 | 0 | 1 |
| UGC0007359 | Clust-UGC0002157 | 6512 | 3 | 1 | 0 | 1 |
| UGC0006928 | Clust-UGC0002157 | 3793 | 1 | 0 | 0 | 0 |
| UGC0006763 | Clust-UGC0002157 | 153 | 1 | 0 | 0 | 0 |
| UGC0013276 | Clust-UGC0002157 | 1601 | 1 | 0 | 0 | 0 |
| UGC0005029 | Clust-UGC0002157 | 8851 | 5 | 1 | 0 | 1 |
| UGC0009465 | Clust-UGC0002157 | 558 | 1 | 0 | 0 | 0 |
| UGC0010748 | Clust-UGC0002157 | 3261 | 5 | 1 | 0 | 1 |
| UGC0009448 | Clust-UGC0002157 | 521 | 1 | 0 | 0 | 0 |
| UGC0009480 | Clust-UGC0002157 | 2411 | 2 | 1 | 0 | 0 |
| UGC0007334 | Clust-UGC0002157 | 3531 | 4 | 1 | 0 | 1 |
| UGC0009051 | Clust-UGC0002157 | 2024 | 1 | 0 | 0 | 0 |
| UGC0013365 | Clust-UGC0002157 | 5959 | 6 | 1 | 0 | 1 |
| UGC0007130 | Clust-UGC0002157 | 5693 | 1 | 0 | 0 | 0 |
| UGC0009605 | Clust-UGC0002157 | 1378 | 3 | 1 | 0 | 1 |
| UGC0009210 | Clust-UGC0002157 | 5380 | 4 | 1 | 0 | 1 |
| UGC0007551 | Clust-UGC0002157 | 4907 | 4 | 1 | 0 | 1 |
| UGC0002185 | Clust-UGC0002157 | 6397 | 4 | 1 | 0 | 1 |
| UGC0008717 | Clust-UGC0002157 | 1424 | 1 | 0 | 0 | 0 |
| UGC0010896 | Clust-UGC0002157 | 297 | 1 | 0 | 0 | 0 |
| UGC0007289 | Clust-UGC0002157 | 5871 | 3 | 1 | 0 | 1 |
| UGC0007397 | Clust-UGC0002157 | 3641 | 3 | 1 | 0 | 0 |
| UGC0009379 | Clust-UGC0002157 | 5699 | 5 | 1 | 0 | 1 |
| UGC0008378 | Clust-UGC0002157 | 163 | 1 | 0 | 0 | 0 |
| UGC0009338 | Clust-UGC0002157 | 9197 | 14 | 1 | 0 | 1 |
| UGC0007082 | Clust-UGC0002157 | 1713 | 3 | 1 | 0 | 0 |
| UGC0006150 | Clust-UGC0002157 | 2411 | 1 | 0 | 0 | 0 |
| UGC0009413 | Clust-UGC0002158 | 2707 | 3 | 1 | 1 | 1 |
| UGC0008618 | Clust-UGC0002158 | 2845 | 2 | 1 | 0 | 1 |
| UGC0006521 | Clust-UGC0002158 | 1047 | 1 | 0 | 0 | 0 |
| UGC0007264 | Clust-UGC0002158 | 7243 | 4 | 1 | 0 | 1 |
| UGC0007598 | Clust-UGC0002158 | 3043 | 1 | 0 | 0 | 0 |
| UGC0010638 | Clust-UGC0002158 | 1391 | 1 | 0 | 0 | 0 |
| UGC0013179 | Clust-UGC0002158 | 2558 | 2 | 1 | 0 | 1 |
| UGC0007678 | Clust-UGC0002158 | 4004 | 1 | 0 | 0 | 0 |
| UGC0005354 | Clust-UGC0002158 | 1306 | 1 | 0 | 0 | 0 |
| UGC0005355 | Clust-UGC0002158 | 2133 | 1 | 0 | 0 | 0 |
| UGC0005651 | Clust-UGC0002158 | 5028 | 1 | 0 | 0 | 0 |
| UGC0007657 | Clust-UGC0002158 | 11183 | 8 | 1 | 0 | 1 |
| UGC0007658 | Clust-UGC0002158 | 6936 | 3 | 1 | 0 | 1 |
| UGC0007009 | Clust-UGC0002158 | 7155 | 2 | 1 | 1 | 0 |
| UGC0012032 | Clust-UGC0002158 | 1716 | 4 | 1 | 2 | 0 |
| UGC0007028 | Clust-UGC0002158 | 7485 | 2 | 1 | 0 | 1 |
| UGC0012846 | Clust-UGC0002158 | 3900 | 4 | 1 | 2 | 0 |
| UGC0004980 | Clust-UGC0002158 | 9104 | 4 | 1 | 0 | 1 |
| UGC0007189 | Clust-UGC0002158 | 3182 | 1 | 0 | 0 | 0 |
| UGC0010207 | Clust-UGC0002158 | 428 | 1 | 0 | 0 | 0 |
| UGC0005386 | Clust-UGC0002158 | 1705 | 1 | 0 | 0 | 0 |
| UGC0005874 | Clust-UGC0002158 | 4465 | 2 | 1 | 0 | 0 |
| UGC0004943 | Clust-UGC0002158 | 1498 | 1 | 0 | 0 | 0 |
| UGC0004947 | Clust-UGC0002158 | 7849 | 5 | 1 | 0 | 1 |
| UGC0004492 | Clust-UGC0002158 | 6343 | 2 | 1 | 0 | 1 |
| UGC0004493 | Clust-UGC0002158 | 6220 | 1 | 0 | 0 | 0 |
| UGC0005692 | Clust-UGC0002158 | 948 | 1 | 0 | 0 | 0 |
| UGC0007695 | Clust-UGC0002158 | 1221 | 1 | 0 | 0 | 0 |
| UGC0005668 | Clust-UGC0002158 | 286 | 1 | 0 | 0 | 0 |
| UGC0003575 | Clust-UGC0002158 | 9339 | 7 | 1 | 0 | 1 |
| UGC0005719 | Clust-UGC0002158 | 4999 | 1 | 0 | 0 | 0 |
| UGC0002158 | Clust-UGC0002158 | 2441 | 2 | 1 | 0 | 0 |
| UGC0006513 | Clust-UGC0002158 | 1781 | 1 | 0 | 0 | 0 |
| UGC0005123 | Clust-UGC0002158 | 1082 | 1 | 0 | 0 | 0 |
| UGC0005034 | Clust-UGC0002158 | 6148 | 4 | 1 | 0 | 1 |
| UGC0005550 | Clust-UGC0002158 | 1040 | 1 | 0 | 0 | 0 |
| UGC0007772 | Clust-UGC0002158 | 3919 | 5 | 1 | 0 | 1 |
| UGC0004157 | Clust-UGC0002158 | 3350 | 2 | 1 | 0 | 0 |
| UGC0006551 | Clust-UGC0002158 | 1365 | 1 | 0 | 0 | 0 |
| UGC0004828 | Clust-UGC0002158 | 3436 | 1 | 0 | 0 | 0 |
| UGC0004799 | Clust-UGC0002158 | 3117 | 2 | 1 | 0 | 1 |
| UGC0004822 | Clust-UGC0002158 | 7505 | 6 | 1 | 0 | 1 |
| UGC0012624 | Clust-UGC0002158 | 3480 | 3 | 1 | 1 | 0 |
| UGC0007273 | Clust-UGC0002158 | 6657 | 3 | 1 | 0 | 1 |
| UGC0007279 | Clust-UGC0002158 | 7846 | 6 | 1 | 0 | 1 |
| UGC0004112 | Clust-UGC0002158 | 1021 | 1 | 0 | 0 | 0 |
| UGC0005645 | Clust-UGC0002158 | 1090 | 1 | 0 | 0 | 0 |
| UGC0005016 | Clust-UGC0002158 | 6943 | 5 | 1 | 0 | 1 |
| UGC0012839 | Clust-UGC0002158 | 3153 | 3 | 1 | 0 | 0 |
| UGC0012835 | Clust-UGC0002158 | 4986 | 4 | 1 | 1 | 0 |

| | | | | | | |
|---|---|---|---|---|---|---|
| UGC0005345 | Clust-UGC0002158 | 2492 | 2 | 1 | 0 | 0 |
| UGC0005660 | Clust-UGC0002158 | 2754 | 1 | 0 | 0 | 0 |
| UGC0007032 | Clust-UGC0002158 | 4449 | 1 | 0 | 0 | 0 |
| UGC0005046 | Clust-UGC0002158 | 1746 | 1 | 0 | 0 | 0 |
| UGC0005592 | Clust-UGC0002158 | 4417 | 2 | 1 | 0 | 1 |
| UGC0013254 | Clust-UGC0002158 | 476 | 2 | 1 | 1 | 0 |
| UGC0005628 | Clust-UGC0002158 | 5779 | 1 | 0 | 0 | 0 |
| UGC0005629 | Clust-UGC0002158 | 1921 | 1 | 0 | 0 | 0 |
| UGC0012850 | Clust-UGC0002158 | 7532 | 7 | 1 | 0 | 1 |
| UGC0012222 | Clust-UGC0002158 | 6531 | 4 | 1 | 1 | 0 |
| UGC0006506 | Clust-UGC0002158 | 1075 | 1 | 0 | 0 | 0 |
| UGC0004999 | Clust-UGC0002158 | 6407 | 2 | 1 | 0 | 0 |
| UGC0006597 | Clust-UGC0002158 | 1134 | 1 | 0 | 0 | 0 |
| UGC0005001 | Clust-UGC0002158 | 7531 | 7 | 1 | 0 | 1 |
| UGC0005760 | Clust-UGC0002158 | 5741 | 3 | 1 | 0 | 0 |
| UGC0004950 | Clust-UGC0002158 | 475 | 1 | 0 | 0 | 0 |
| UGC0005682 | Clust-UGC0002158 | 1268 | 1 | 0 | 0 | 0 |
| UGC0004562 | Clust-UGC0002158 | 7241 | 3 | 1 | 0 | 1 |
| UGC0006610 | Clust-UGC0002158 | 1127 | 1 | 0 | 0 | 0 |
| UGC0007495 | Clust-UGC0002158 | 3168 | 1 | 0 | 0 | 0 |
| UGC0012245 | Clust-UGC0002158 | 2671 | 1 | 0 | 0 | 0 |
| UGC0004622 | Clust-UGC0002158 | 4674 | 1 | 0 | 0 | 0 |
| UGC0005397 | Clust-UGC0002158 | 2511 | 2 | 1 | 0 | 1 |
| UGC0007740 | Clust-UGC0002158 | 6426 | 5 | 1 | 0 | 1 |
| UGC0012198 | Clust-UGC0002158 | 3960 | 3 | 1 | 0 | 1 |
| UGC0010103 | Clust-UGC0002158 | 6026 | 2 | 1 | 0 | 0 |
| UGC0004149 | Clust-UGC0002158 | 1057 | 1 | 0 | 0 | 0 |
| UGC0007614 | Clust-UGC0002158 | 3651 | 2 | 1 | 0 | 1 |
| UGC0006543 | Clust-UGC0002158 | 1050 | 1 | 0 | 0 | 0 |
| UGC0011199 | Clust-UGC0002158 | 2012 | 3 | 1 | 1 | 0 |
| UGC0005717 | Clust-UGC0002162 | 5726 | 6 | 1 | 0 | 1 |
| UGC0002162 | Clust-UGC0002162 | 8962 | 24 | 1 | 0 | 1 |
| UGC0007206 | Clust-UGC0002164 | 2698 | 2 | 1 | 1 | 0 |
| UGC0011566 | Clust-UGC0002164 | 2368 | 1 | 0 | 0 | 0 |
| UGC0013155 | Clust-UGC0002164 | 1181 | 1 | 0 | 0 | 0 |
| UGC0013152 | Clust-UGC0002164 | 4048 | 1 | 0 | 0 | 0 |
| UGC0009730 | Clust-UGC0002164 | 1881 | 2 | 1 | 0 | 1 |
| UGC0004766 | Clust-UGC0002164 | 2160 | 1 | 0 | 0 | 0 |
| UGC0004817 | Clust-UGC0002164 | 8465 | 3 | 1 | 0 | 1 |
| UGC0004103 | Clust-UGC0002164 | 367 | 1 | 0 | 0 | 0 |
| UGC0003720 | Clust-UGC0002164 | 1485 | 2 | 1 | 1 | 0 |
| UGC0006022 | Clust-UGC0002164 | 285 | 1 | 0 | 0 | 0 |
| UGC0010286 | Clust-UGC0002164 | 4312 | 1 | 0 | 0 | 0 |
| UGC0005350 | Clust-UGC0002164 | 2192 | 1 | 0 | 0 | 0 |
| UGC0003745 | Clust-UGC0002164 | 304 | 1 | 0 | 0 | 0 |
| UGC0003748 | Clust-UGC0002164 | 202 | 1 | 0 | 0 | 0 |
| UGC0011907 | Clust-UGC0002164 | 12121 | 6 | 1 | 0 | 1 |
| UGC0002991 | Clust-UGC0002164 | 1144 | 1 | 0 | 0 | 0 |
| UGC0010260 | Clust-UGC0002164 | 2088 | 1 | 0 | 0 | 0 |
| UGC0012639 | Clust-UGC0002164 | 5881 | 2 | 1 | 0 | 0 |
| UGC0004724 | Clust-UGC0002164 | 2613 | 1 | 0 | 0 | 0 |
| UGC0009548 | Clust-UGC0002164 | 868 | 1 | 0 | 0 | 0 |
| UGC0009174 | Clust-UGC0002164 | 2837 | 1 | 0 | 0 | 0 |
| UGC0006449 | Clust-UGC0002164 | 256 | 1 | 0 | 0 | 0 |
| UGC0003028 | Clust-UGC0002164 | 1730 | 1 | 0 | 0 | 0 |
| UGC0007594 | Clust-UGC0002164 | 3041 | 1 | 0 | 0 | 0 |
| UGC0008882 | Clust-UGC0002164 | 414 | 1 | 0 | 0 | 0 |
| UGC0013019 | Clust-UGC0002164 | 8233 | 2 | 1 | 0 | 1 |
| UGC0013017 | Clust-UGC0002164 | 2552 | 1 | 0 | 0 | 0 |
| UGC0003707 | Clust-UGC0002164 | 910 | 1 | 0 | 0 | 0 |
| UGC0003706 | Clust-UGC0002164 | 90 | 1 | 0 | 0 | 0 |
| UGC0005613 | Clust-UGC0002164 | 430 | 1 | 0 | 0 | 0 |
| UGC0006905 | Clust-UGC0002164 | 2002 | 1 | 0 | 0 | 0 |
| UGC0011885 | Clust-UGC0002164 | 630 | 1 | 0 | 0 | 0 |
| UGC0012250 | Clust-UGC0002164 | 2600 | 3 | 1 | 0 | 0 |
| UGC0011467 | Clust-UGC0002164 | 2285 | 1 | 0 | 0 | 0 |
| UGC0011466 | Clust-UGC0002164 | 2487 | 2 | 1 | 0 | 0 |
| UGC0011465 | Clust-UGC0002164 | 1028 | 3 | 1 | 0 | 0 |
| UGC0012844 | Clust-UGC0002164 | 2322 | 2 | 1 | 0 | 0 |
| UGC0006931 | Clust-UGC0002164 | 1242 | 1 | 0 | 0 | 0 |
| UGC0008515 | Clust-UGC0002164 | 52 | 1 | 0 | 0 | 0 |
| UGC0013244 | Clust-UGC0002164 | 1003 | 2 | 1 | 1 | 0 |
| UGC0008523 | Clust-UGC0002164 | 646 | 1 | 0 | 0 | 0 |
| UGC0009734 | Clust-UGC0002164 | 2348 | 2 | 1 | 0 | 1 |
| UGC0008746 | Clust-UGC0002164 | 93 | 1 | 0 | 0 | 0 |
| UGC0006772 | Clust-UGC0002164 | 4199 | 1 | 0 | 0 | 0 |
| UGC0008263 | Clust-UGC0002164 | 868 | 1 | 0 | 0 | 0 |
| UGC0007592 | Clust-UGC0002164 | 2541 | 1 | 0 | 0 | 0 |
| UGC0009475 | Clust-UGC0002164 | 2938 | 1 | 0 | 0 | 0 |

| | | | | | | |
|---|---|---|---|---|---|---|
| UGC0004942 | Clust-UGC0002164 | 1638 | 1 | 0 | 0 | 0 |
| UGC0004495 | Clust-UGC0002164 | 6324 | 1 | 0 | 0 | 0 |
| UGC0005018 | Clust-UGC0002164 | 5130 | 1 | 0 | 0 | 0 |
| UGC0006096 | Clust-UGC0002164 | 1063 | 1 | 0 | 0 | 0 |
| UGC0012715 | Clust-UGC0002164 | 8043 | 6 | 1 | 0 | 1 |
| UGC0008001 | Clust-UGC0002164 | 1844 | 1 | 0 | 0 | 0 |
| UGC0004557 | Clust-UGC0002164 | 1556 | 1 | 0 | 0 | 0 |
| UGC0009992 | Clust-UGC0002164 | 923 | 1 | 0 | 0 | 0 |
| UGC0005364 | Clust-UGC0002164 | 7016 | 2 | 1 | 0 | 1 |
| UGC0004611 | Clust-UGC0002164 | 4520 | 1 | 0 | 0 | 0 |
| UGC0006616 | Clust-UGC0002164 | 1195 | 1 | 0 | 0 | 0 |
| UGC0008068 | Clust-UGC0002164 | 217 | 1 | 0 | 0 | 0 |
| UGC0008066 | Clust-UGC0002164 | 168 | 1 | 0 | 0 | 0 |
| UGC0004579 | Clust-UGC0002164 | 8454 | 4 | 1 | 0 | 1 |
| UGC0009087 | Clust-UGC0002164 | 1959 | 2 | 1 | 1 | 0 |
| UGC0009081 | Clust-UGC0002164 | 957 | 1 | 0 | 0 | 0 |
| UGC0009144 | Clust-UGC0002164 | 3524 | 1 | 0 | 0 | 0 |
| UGC0010885 | Clust-UGC0002164 | 1778 | 1 | 0 | 0 | 0 |
| UGC0008434 | Clust-UGC0002164 | 1184 | 1 | 0 | 0 | 0 |
| UGC0008608 | Clust-UGC0002164 | 2155 | 1 | 0 | 0 | 0 |
| UGC0009607 | Clust-UGC0002164 | 934 | 1 | 0 | 0 | 0 |
| UGC0007946 | Clust-UGC0002164 | 3120 | 1 | 0 | 0 | 0 |
| UGC0007710 | Clust-UGC0002164 | 5659 | 1 | 0 | 0 | 0 |
| UGC0004807 | Clust-UGC0002164 | 1470 | 2 | 1 | 0 | 1 |
| UGC0008381 | Clust-UGC0002164 | 1589 | 1 | 0 | 0 | 0 |
| UGC0009168 | Clust-UGC0002164 | 3835 | 1 | 0 | 0 | 0 |
| UGC0009598 | Clust-UGC0002164 | 909 | 1 | 0 | 0 | 0 |
| UGC0009591 | Clust-UGC0002164 | 1695 | 2 | 1 | 0 | 0 |
| UGC0008978 | Clust-UGC0002164 | 3079 | 1 | 0 | 0 | 0 |
| UGC0008970 | Clust-UGC0002164 | 136 | 1 | 0 | 0 | 0 |
| UGC0008629 | Clust-UGC0002164 | 3988 | 1 | 0 | 0 | 0 |
| UGC0007738 | Clust-UGC0002164 | 5608 | 1 | 0 | 0 | 0 |
| UGC0010920 | Clust-UGC0002164 | 1003 | 1 | 0 | 0 | 0 |
| UGC0002488 | Clust-UGC0002164 | 223 | 1 | 0 | 0 | 0 |
| UGC0008199 | Clust-UGC0002164 | 3580 | 1 | 0 | 0 | 0 |
| UGC0009342 | Clust-UGC0002164 | 3489 | 1 | 0 | 0 | 0 |
| UGC0009664 | Clust-UGC0002164 | 1046 | 1 | 0 | 0 | 0 |
| UGC0008229 | Clust-UGC0002164 | 244 | 1 | 0 | 0 | 0 |
| UGC0008815 | Clust-UGC0002164 | 2874 | 1 | 0 | 0 | 0 |
| UGC0006367 | Clust-UGC0002164 | 5611 | 1 | 0 | 0 | 0 |
| UGC0003798 | Clust-UGC0002164 | 555 | 2 | 1 | 0 | 0 |
| UGC0004800 | Clust-UGC0002164 | 15211 | 22 | 1 | 0 | 1 |
| UGC0002357 | Clust-UGC0002164 | 4232 | 1 | 0 | 0 | 0 |
| UGC0009304 | Clust-UGC0002164 | 4105 | 2 | 1 | 0 | 1 |
| UGC0009303 | Clust-UGC0002164 | 1309 | 1 | 0 | 0 | 0 |
| UGC0009302 | Clust-UGC0002164 | 262 | 1 | 0 | 0 | 0 |
| UGC0009300 | Clust-UGC0002164 | 1092 | 1 | 0 | 0 | 0 |
| UGC0008494 | Clust-UGC0002164 | 1439 | 1 | 0 | 0 | 0 |
| UGC0007212 | Clust-UGC0002164 | 440 | 1 | 0 | 0 | 0 |
| UGC0004821 | Clust-UGC0002164 | 6006 | 1 | 0 | 0 | 0 |
| UGC0009134 | Clust-UGC0002164 | 2855 | 1 | 0 | 0 | 0 |
| UGC0007665 | Clust-UGC0002164 | 7034 | 2 | 1 | 0 | 1 |
| UGC0013395 | Clust-UGC0002164 | 901 | 1 | 0 | 0 | 0 |
| UGC0006702 | Clust-UGC0002164 | 173 | 1 | 0 | 0 | 0 |
| UGC0009536 | Clust-UGC0002164 | 4485 | 1 | 0 | 0 | 0 |
| UGC0011915 | Clust-UGC0002164 | 3357 | 2 | 1 | 1 | 0 |
| UGC0004757 | Clust-UGC0002164 | 1997 | 1 | 0 | 0 | 0 |
| UGC0005884 | Clust-UGC0002164 | 1673 | 1 | 0 | 0 | 0 |
| UGC0009768 | Clust-UGC0002164 | 619 | 1 | 0 | 0 | 0 |
| UGC0007018 | Clust-UGC0002164 | 2140 | 1 | 0 | 0 | 0 |
| UGC0008474 | Clust-UGC0002164 | 986 | 1 | 0 | 0 | 0 |
| UGC0004735 | Clust-UGC0002164 | 407 | 1 | 0 | 0 | 0 |
| UGC0009575 | Clust-UGC0002164 | 1826 | 2 | 1 | 0 | 1 |
| UGC0003019 | Clust-UGC0002164 | 1416 | 1 | 0 | 0 | 0 |
| UGC0006008 | Clust-UGC0002164 | 2332 | 1 | 0 | 0 | 0 |
| UGC0012788 | Clust-UGC0002164 | 4276 | 1 | 0 | 0 | 0 |
| UGC0012872 | Clust-UGC0002164 | 6054 | 2 | 1 | 0 | 0 |
| UGC0006925 | Clust-UGC0002164 | 1834 | 1 | 0 | 0 | 0 |
| UGC0009031 | Clust-UGC0002164 | 629 | 1 | 0 | 0 | 0 |
| UGC0008175 | Clust-UGC0002164 | 2508 | 1 | 0 | 0 | 0 |
| UGC0004955 | Clust-UGC0002164 | 4473 | 1 | 0 | 0 | 0 |
| UGC0006695 | Clust-UGC0002164 | 278 | 1 | 0 | 0 | 0 |
| UGC0011419 | Clust-UGC0002164 | 2939 | 2 | 1 | 0 | 0 |
| UGC0013068 | Clust-UGC0002164 | 6111 | 1 | 0 | 0 | 0 |
| UGC0008690 | Clust-UGC0002164 | 197 | 1 | 0 | 0 | 0 |
| UGC0004994 | Clust-UGC0002164 | 6639 | 2 | 1 | 0 | 0 |
| UGC0009462 | Clust-UGC0002164 | 1284 | 1 | 0 | 0 | 0 |
| UGC0008038 | Clust-UGC0002164 | 375 | 1 | 0 | 0 | 0 |
| UGC0007782 | Clust-UGC0002164 | 3869 | 1 | 0 | 0 | 0 |

| | | | | | | |
|---|---|---|---|---|---|---|
| UGC0010558 | Clust-UGC0002164 | 2615 | 1 | 0 | 0 | 0 |
| UGC0010559 | Clust-UGC0002164 | 3475 | 2 | 1 | 1 | 0 |
| UGC0009622 | Clust-UGC0002164 | 691 | 1 | 0 | 0 | 0 |
| UGC0012549 | Clust-UGC0002164 | 6225 | 2 | 1 | 0 | 0 |
| UGC0004546 | Clust-UGC0002164 | 2226 | 1 | 0 | 0 | 0 |
| UGC0008905 | Clust-UGC0002164 | 3091 | 1 | 0 | 0 | 0 |
| UGC0013369 | Clust-UGC0002164 | 3141 | 3 | 1 | 0 | 1 |
| UGC0013367 | Clust-UGC0002164 | 487 | 1 | 0 | 0 | 0 |
| UGC0008071 | Clust-UGC0002164 | 1459 | 1 | 0 | 0 | 0 |
| UGC0007554 | Clust-UGC0002164 | 4142 | 1 | 0 | 0 | 0 |
| UGC0007471 | Clust-UGC0002164 | 3703 | 1 | 0 | 0 | 0 |
| UGC0013348 | Clust-UGC0002164 | 2109 | 2 | 1 | 0 | 0 |
| UGC0013347 | Clust-UGC0002164 | 3199 | 4 | 1 | 1 | 1 |
| UGC0012581 | Clust-UGC0002164 | 7606 | 2 | 1 | 0 | 0 |
| UGC0008442 | Clust-UGC0002164 | 390 | 1 | 0 | 0 | 0 |
| UGC0008055 | Clust-UGC0002164 | 442 | 1 | 0 | 0 | 0 |
| UGC0009231 | Clust-UGC0002164 | 890 | 1 | 0 | 0 | 0 |
| UGC0008948 | Clust-UGC0002164 | 334 | 1 | 0 | 0 | 0 |
| UGC0005531 | Clust-UGC0002164 | 713 | 1 | 0 | 0 | 0 |
| UGC0005534 | Clust-UGC0002164 | 483 | 1 | 0 | 0 | 0 |
| UGC0002164 | Clust-UGC0002164 | 4552 | 2 | 1 | 0 | 0 |
| UGC0005390 | Clust-UGC0002164 | 5282 | 1 | 0 | 0 | 0 |
| UGC0005391 | Clust-UGC0002164 | 3621 | 1 | 0 | 0 | 0 |
| UGC0013321 | Clust-UGC0002164 | 3002 | 2 | 1 | 0 | 0 |
| UGC0009171 | Clust-UGC0002164 | 2840 | 1 | 0 | 0 | 0 |
| UGC0008420 | Clust-UGC0002164 | 1365 | 1 | 0 | 0 | 0 |
| UGC0002956 | Clust-UGC0002164 | 881 | 1 | 0 | 0 | 0 |
| UGC0008969 | Clust-UGC0002164 | 1236 | 2 | 1 | 1 | 0 |
| UGC0008392 | Clust-UGC0002164 | 2551 | 1 | 0 | 0 | 0 |
| UGC0009377 | Clust-UGC0002164 | 1621 | 1 | 0 | 0 | 0 |
| UGC0002494 | Clust-UGC0002164 | 111 | 1 | 0 | 0 | 0 |
| UGC0002499 | Clust-UGC0002164 | 2269 | 1 | 0 | 0 | 0 |
| UGC0006861 | Clust-UGC0002164 | 1760 | 1 | 0 | 0 | 0 |
| UGC0005731 | Clust-UGC0002164 | 6161 | 1 | 0 | 0 | 0 |
| UGC0008983 | Clust-UGC0002164 | 1963 | 2 | 1 | 1 | 0 |
| UGC0002553 | Clust-UGC0002164 | 267 | 1 | 0 | 0 | 0 |
| UGC0008984 | Clust-UGC0002164 | 4434 | 1 | 0 | 0 | 0 |
| UGC0008212 | Clust-UGC0002164 | 2217 | 1 | 0 | 0 | 0 |
| UGC0008375 | Clust-UGC0002164 | 155 | 1 | 0 | 0 | 0 |
| UGC0009355 | Clust-UGC0002164 | 2268 | 1 | 0 | 0 | 0 |
| UGC0009357 | Clust-UGC0002164 | 4495 | 2 | 1 | 0 | 1 |
| UGC0005130 | Clust-UGC0002164 | 2167 | 1 | 0 | 0 | 0 |
| UGC0010123 | Clust-UGC0002164 | 5735 | 2 | 1 | 0 | 1 |
| UGC0009655 | Clust-UGC0002164 | 820 | 1 | 0 | 0 | 0 |
| UGC0009650 | Clust-UGC0002164 | 656 | 1 | 0 | 0 | 0 |
| UGC0012193 | Clust-UGC0002164 | 3762 | 1 | 0 | 0 | 0 |
| UGC0008359 | Clust-UGC0002164 | 619 | 1 | 0 | 0 | 0 |
| UGC0008800 | Clust-UGC0002164 | 1807 | 1 | 0 | 0 | 0 |
| UGC0008054 | Clust-UGC0002164 | 380 | 1 | 0 | 0 | 0 |
| UGC0006689 | Clust-UGC0002164 | 5068 | 1 | 0 | 0 | 0 |
| UGC0007087 | Clust-UGC0002164 | 1708 | 1 | 0 | 0 | 0 |
| UGC0007611 | Clust-UGC0002164 | 3819 | 2 | 1 | 0 | 0 |
| UGC0013135 | Clust-UGC0002164 | 425 | 1 | 0 | 0 | 0 |
| UGC0009312 | Clust-UGC0002164 | 1772 | 1 | 0 | 0 | 0 |
| UGC0008622 | Clust-UGC0002164 | 3440 | 1 | 0 | 0 | 0 |
| UGC0006028 | Clust-UGC0002165 | 4057 | 5 | 1 | 0 | 1 |
| UGC0004745 | Clust-UGC0002165 | 415 | 3 | 1 | 0 | 1 |
| UGC0004741 | Clust-UGC0002165 | 502 | 2 | 1 | 0 | 1 |
| UGC0002230 | Clust-UGC0002165 | 4829 | 7 | 1 | 0 | 1 |
| UGC0006097 | Clust-UGC0002165 | 1838 | 1 | 0 | 0 | 0 |
| UGC0006090 | Clust-UGC0002165 | 2145 | 2 | 1 | 0 | 0 |
| UGC0005978 | Clust-UGC0002165 | 5256 | 1 | 0 | 0 | 0 |
| UGC0012279 | Clust-UGC0002165 | 105 | 1 | 0 | 0 | 0 |
| UGC0010171 | Clust-UGC0002165 | 1796 | 3 | 1 | 0 | 1 |
| UGC0011129 | Clust-UGC0002165 | 318 | 1 | 0 | 0 | 0 |
| UGC0011125 | Clust-UGC0002165 | 499 | 1 | 0 | 0 | 0 |
| UGC0010087 | Clust-UGC0002165 | 4170 | 5 | 1 | 0 | 1 |
| UGC0013007 | Clust-UGC0002165 | 4500 | 1 | 0 | 0 | 0 |
| UGC0009444 | Clust-UGC0002165 | 115 | 1 | 0 | 0 | 0 |
| UGC0002538 | Clust-UGC0002165 | 2299 | 3 | 1 | 0 | 1 |
| UGC0002165 | Clust-UGC0002165 | 3357 | 3 | 1 | 0 | 1 |
| UGC0005817 | Clust-UGC0002165 | 4024 | 1 | 0 | 0 | 0 |
| UGC0005893 | Clust-UGC0002165 | 5138 | 7 | 1 | 0 | 1 |
| UGC0004747 | Clust-UGC0002166 | 221 | 1 | 0 | 0 | 0 |
| UGC0009264 | Clust-UGC0002166 | 1946 | 12 | 1 | 0 | 1 |
| UGC0004571 | Clust-UGC0002166 | 5801 | 23 | 1 | 0 | 1 |
| UGC0012310 | Clust-UGC0002166 | 668 | 2 | 1 | 0 | 1 |
| UGC0008610 | Clust-UGC0002166 | 8044 | 31 | 1 | 0 | 1 |
| UGC0002166 | Clust-UGC0002166 | 6219 | 17 | 1 | 0 | 1 |

| | | | | | | |
|---|---|---|---|---|---|---|
| UGC0010894 | Clust-UGC0002166 | 4465 | 13 | 1 | 0 | 1 |
| UGC0009009 | Clust-UGC0002166 | 7220 | 26 | 1 | 1 | 1 |
| UGC0011541 | Clust-UGC0002167 | 2094 | 3 | 1 | 0 | 0 |
| UGC0004108 | Clust-UGC0002167 | 786 | 1 | 0 | 0 | 0 |
| UGC0012134 | Clust-UGC0002167 | 2494 | 7 | 1 | 1 | 1 |
| UGC0003307 | Clust-UGC0002167 | 2167 | 1 | 0 | 0 | 0 |
| UGC0009263 | Clust-UGC0002167 | 1254 | 1 | 0 | 0 | 0 |
| UGC0007501 | Clust-UGC0002167 | 5211 | 3 | 1 | 0 | 1 |
| UGC0013120 | Clust-UGC0002167 | 1573 | 1 | 0 | 0 | 0 |
| UGC0007095 | Clust-UGC0002167 | 262 | 1 | 0 | 0 | 0 |
| UGC0006302 | Clust-UGC0002167 | 5393 | 7 | 1 | 0 | 1 |
| UGC0006413 | Clust-UGC0002167 | 7696 | 21 | 1 | 0 | 1 |
| UGC0013236 | Clust-UGC0002167 | 1235 | 3 | 1 | 2 | 0 |
| UGC0006002 | Clust-UGC0002167 | 6967 | 10 | 1 | 0 | 1 |
| UGC0006411 | Clust-UGC0002167 | 2714 | 2 | 1 | 0 | 1 |
| UGC0008454 | Clust-UGC0002167 | 932 | 1 | 0 | 0 | 0 |
| UGC0003522 | Clust-UGC0002167 | 5660 | 4 | 1 | 0 | 1 |
| UGC0002167 | Clust-UGC0002167 | 3677 | 3 | 1 | 0 | 1 |
| UGC0006392 | Clust-UGC0002167 | 6818 | 6 | 1 | 0 | 1 |
| UGC0005833 | Clust-UGC0002167 | 9058 | 22 | 1 | 0 | 1 |
| UGC0006821 | Clust-UGC0002167 | 2682 | 2 | 1 | 0 | 1 |
| UGC0007941 | Clust-UGC0002167 | 937 | 1 | 0 | 0 | 0 |
| UGC0008312 | Clust-UGC0002172 | 387 | 1 | 0 | 0 | 0 |
| UGC0009190 | Clust-UGC0002172 | 5609 | 6 | 1 | 0 | 1 |
| UGC0008489 | Clust-UGC0002172 | 1409 | 1 | 0 | 0 | 0 |
| UGC0008488 | Clust-UGC0002172 | 731 | 1 | 0 | 0 | 0 |
| UGC0007269 | Clust-UGC0002172 | 8306 | 24 | 1 | 1 | 1 |
| UGC0011464 | Clust-UGC0002172 | 5163 | 9 | 1 | 0 | 1 |
| UGC0010228 | Clust-UGC0002172 | 312 | 2 | 1 | 0 | 0 |
| UGC0008027 | Clust-UGC0002172 | 1455 | 1 | 0 | 0 | 0 |
| UGC0010524 | Clust-UGC0002172 | 3904 | 2 | 1 | 0 | 1 |
| UGC0008564 | Clust-UGC0002172 | 2819 | 2 | 1 | 0 | 1 |
| UGC0002172 | Clust-UGC0002172 | 3813 | 3 | 1 | 0 | 1 |
| UGC0005821 | Clust-UGC0002172 | 3310 | 2 | 1 | 0 | 1 |
| UGC0007757 | Clust-UGC0002172 | 5659 | 10 | 1 | 0 | 1 |
| UGC0008205 | Clust-UGC0002172 | 1725 | 1 | 0 | 0 | 0 |
| UGC0012947 | Clust-UGC0002172 | 681 | 1 | 0 | 0 | 0 |
| UGC0010609 | Clust-UGC0002172 | 1435 | 1 | 0 | 0 | 0 |
| UGC0003778 | Clust-UGC0002172 | 815 | 2 | 1 | 0 | 1 |
| UGC0008177 | Clust-UGC0002172 | 3748 | 3 | 1 | 0 | 1 |
| UGC0009254 | Clust-UGC0002172 | 10454 | 51 | 1 | 0 | 1 |
| UGC0012956 | Clust-UGC0002172 | 3467 | 3 | 1 | 0 | 1 |
| UGC0008708 | Clust-UGC0002172 | 1523 | 1 | 0 | 0 | 0 |
| UGC0003021 | Clust-UGC0002172 | 2449 | 2 | 1 | 0 | 1 |
| UGC0007764 | Clust-UGC0002172 | 2781 | 1 | 0 | 0 | 0 |
| UGC0002334 | Clust-UGC0002174 | 2425 | 1 | 0 | 0 | 0 |
| UGC0003761 | Clust-UGC0002174 | 521 | 1 | 0 | 0 | 0 |
| UGC0008883 | Clust-UGC0002174 | 1843 | 1 | 0 | 0 | 0 |
| UGC0008701 | Clust-UGC0002174 | 334 | 1 | 0 | 0 | 0 |
| UGC0002505 | Clust-UGC0002174 | 3431 | 1 | 0 | 0 | 0 |
| UGC0008931 | Clust-UGC0002174 | 5025 | 1 | 0 | 0 | 0 |
| UGC0002174 | Clust-UGC0002174 | 3713 | 2 | 1 | 0 | 0 |
| UGC0008847 | Clust-UGC0002174 | 1331 | 1 | 0 | 0 | 0 |
| UGC0008367 | Clust-UGC0002174 | 592 | 1 | 0 | 0 | 0 |
| UGC0006168 | Clust-UGC0002174 | 1225 | 1 | 0 | 0 | 0 |
| UGC0008720 | Clust-UGC0002174 | 218 | 1 | 0 | 0 | 0 |
| UGC0005365 | Clust-UGC0002174 | 6364 | 3 | 1 | 0 | 1 |
| UGC0008545 | Clust-UGC0002174 | 734 | 1 | 0 | 0 | 0 |
| UGC0008738 | Clust-UGC0002174 | 807 | 1 | 0 | 0 | 0 |
| UGC0008946 | Clust-UGC0002174 | 2438 | 1 | 0 | 0 | 0 |
| UGC0008732 | Clust-UGC0002174 | 694 | 1 | 0 | 0 | 0 |
| UGC0008750 | Clust-UGC0002174 | 134 | 1 | 0 | 0 | 0 |
| UGC0007170 | Clust-UGC0002174 | 2311 | 1 | 0 | 0 | 0 |
| UGC0008707 | Clust-UGC0002174 | 1567 | 1 | 0 | 0 | 0 |
| UGC0008822 | Clust-UGC0002174 | 2078 | 1 | 0 | 0 | 0 |
| UGC0008563 | Clust-UGC0002174 | 683 | 1 | 0 | 0 | 0 |
| UGC0007681 | Clust-UGC0002174 | 6198 | 1 | 0 | 0 | 0 |
| UGC0007118 | Clust-UGC0002174 | 1036 | 1 | 0 | 0 | 0 |
| UGC0002512 | Clust-UGC0002174 | 3501 | 1 | 0 | 0 | 0 |
| UGC0008718 | Clust-UGC0002174 | 476 | 1 | 0 | 0 | 0 |
| UGC0003247 | Clust-UGC0002174 | 4123 | 1 | 0 | 0 | 0 |
| UGC0008409 | Clust-UGC0002174 | 211 | 1 | 0 | 0 | 0 |
| UGC0008183 | Clust-UGC0002174 | 1560 | 1 | 0 | 0 | 0 |
| UGC0008594 | Clust-UGC0002174 | 5597 | 3 | 1 | 0 | 1 |
| UGC0003299 | Clust-UGC0002174 | 7731 | 2 | 1 | 0 | 1 |
| UGC0008352 | Clust-UGC0002174 | 1188 | 1 | 0 | 0 | 0 |
| UGC0004469 | Clust-UGC0002174 | 3808 | 1 | 0 | 0 | 0 |
| UGC0008251 | Clust-UGC0002174 | 1187 | 1 | 0 | 0 | 0 |
| UGC0002473 | Clust-UGC0002174 | 169 | 1 | 0 | 0 | 0 |

| | | | | | | |
|---|---|---|---|---|---|---|
| UGC0008294 | Clust-UGC0002175 | 2581 | 1 | 0 | 0 | 0 |
| UGC0004722 | Clust-UGC0002175 | 2627 | 4 | 1 | 0 | 1 |
| UGC0013226 | Clust-UGC0002175 | 616 | 2 | 1 | 1 | 0 |
| UGC0004986 | Clust-UGC0002175 | 9215 | 32 | 1 | 0 | 1 |
| UGC0012885 | Clust-UGC0002175 | 2307 | 3 | 1 | 0 | 0 |
| UGC0006592 | Clust-UGC0002175 | 749 | 1 | 0 | 0 | 0 |
| UGC0004653 | Clust-UGC0002175 | 11454 | 74 | 1 | 0 | 1 |
| UGC0002175 | Clust-UGC0002175 | 2452 | 1 | 0 | 0 | 0 |
| UGC0003266 | Clust-UGC0002175 | 908 | 2 | 1 | 0 | 1 |
| UGC0012952 | Clust-UGC0002175 | 1370 | 1 | 0 | 0 | 0 |
| UGC0013360 | Clust-UGC0002175 | 8463 | 34 | 1 | 1 | 1 |
| UGC0008673 | Clust-UGC0002175 | 77 | 1 | 0 | 0 | 0 |
| UGC0007477 | Clust-UGC0002175 | 2465 | 1 | 0 | 0 | 0 |
| UGC0003298 | Clust-UGC0002175 | 7545 | 18 | 1 | 0 | 1 |
| UGC0012699 | Clust-UGC0002175 | 3891 | 4 | 1 | 0 | 1 |
| UGC0007559 | Clust-UGC0002175 | 10227 | 59 | 1 | 0 | 1 |
| UGC0007924 | Clust-UGC0002177 | 3114 | 3 | 1 | 0 | 1 |
| UGC0006528 | Clust-UGC0002177 | 1737 | 4 | 1 | 1 | 1 |
| UGC0013258 | Clust-UGC0002177 | 5351 | 18 | 1 | 1 | 1 |
| UGC0006313 | Clust-UGC0002177 | 6107 | 9 | 1 | 0 | 1 |
| UGC0008623 | Clust-UGC0002177 | 1172 | 1 | 0 | 0 | 0 |
| UGC0007023 | Clust-UGC0002177 | 4051 | 5 | 1 | 0 | 1 |
| UGC0007288 | Clust-UGC0002177 | 3779 | 4 | 1 | 0 | 1 |
| UGC0004704 | Clust-UGC0002177 | 8123 | 18 | 1 | 0 | 1 |
| UGC0013223 | Clust-UGC0002177 | 951 | 1 | 0 | 0 | 0 |
| UGC0004790 | Clust-UGC0002177 | 409 | 1 | 0 | 0 | 0 |
| UGC0011882 | Clust-UGC0002177 | 3455 | 3 | 1 | 0 | 1 |
| UGC0008741 | Clust-UGC0002177 | 1043 | 3 | 1 | 0 | 1 |
| UGC0011795 | Clust-UGC0002177 | 2924 | 9 | 1 | 1 | 1 |
| UGC0011793 | Clust-UGC0002177 | 1334 | 2 | 1 | 0 | 0 |
| UGC0006930 | Clust-UGC0002177 | 2084 | 1 | 0 | 0 | 0 |
| UGC0011631 | Clust-UGC0002177 | 3234 | 7 | 1 | 1 | 1 |
| UGC0004512 | Clust-UGC0002177 | 7253 | 12 | 1 | 0 | 1 |
| UGC0004519 | Clust-UGC0002177 | 3593 | 2 | 1 | 0 | 1 |
| UGC0004518 | Clust-UGC0002177 | 8231 | 28 | 1 | 0 | 1 |
| UGC0008680 | Clust-UGC0002177 | 1617 | 1 | 0 | 0 | 0 |
| UGC0002177 | Clust-UGC0002177 | 6168 | 7 | 1 | 0 | 1 |
| UGC0008025 | Clust-UGC0002177 | 837 | 1 | 0 | 0 | 0 |
| UGC0005954 | Clust-UGC0002177 | 2766 | 2 | 1 | 0 | 1 |
| UGC0006793 | Clust-UGC0002177 | 3926 | 4 | 1 | 1 | 1 |
| UGC0010845 | Clust-UGC0002177 | 3012 | 3 | 1 | 0 | 0 |
| UGC0009201 | Clust-UGC0002177 | 4779 | 5 | 1 | 0 | 1 |
| UGC0006953 | Clust-UGC0002177 | 3475 | 7 | 1 | 0 | 1 |
| UGC0011803 | Clust-UGC0002177 | 3670 | 6 | 1 | 0 | 1 |
| UGC0005695 | Clust-UGC0002177 | 612 | 1 | 0 | 0 | 0 |
| UGC0009220 | Clust-UGC0002177 | 5028 | 6 | 1 | 0 | 1 |
| UGC0011827 | Clust-UGC0002177 | 3147 | 4 | 1 | 0 | 1 |
| UGC0012750 | Clust-UGC0002177 | 2583 | 5 | 1 | 0 | 1 |
| UGC0012754 | Clust-UGC0002177 | 457 | 1 | 0 | 0 | 0 |
| UGC0011845 | Clust-UGC0002177 | 3123 | 2 | 1 | 0 | 1 |
| UGC0009625 | Clust-UGC0002177 | 1651 | 1 | 0 | 0 | 0 |
| UGC0008202 | Clust-UGC0002177 | 4238 | 4 | 1 | 0 | 1 |
| UGC0004478 | Clust-UGC0002177 | 6177 | 16 | 1 | 0 | 1 |
| UGC0008838 | Clust-UGC0002177 | 2477 | 2 | 1 | 0 | 1 |
| UGC0008834 | Clust-UGC0002177 | 1407 | 2 | 1 | 0 | 0 |
| UGC0011532 | Clust-UGC0002177 | 11392 | 71 | 1 | 0 | 1 |
| UGC0005804 | Clust-UGC0002177 | 6104 | 9 | 1 | 0 | 1 |
| UGC0005556 | Clust-UGC0002177 | 226 | 1 | 0 | 0 | 0 |
| UGC0011970 | Clust-UGC0002177 | 11847 | 65 | 1 | 0 | 1 |
| UGC0007239 | Clust-UGC0002177 | 7328 | 22 | 1 | 0 | 1 |
| UGC0008246 | Clust-UGC0002177 | 1149 | 2 | 1 | 0 | 1 |
| UGC0006309 | Clust-UGC0002177 | 5699 | 7 | 1 | 0 | 1 |
| UGC0011959 | Clust-UGC0002177 | 7318 | 23 | 1 | 0 | 1 |
| UGC0010540 | Clust-UGC0002177 | 132 | 1 | 0 | 0 | 0 |
| UGC0008544 | Clust-UGC0002177 | 333 | 1 | 0 | 0 | 0 |
| UGC0007070 | Clust-UGC0002177 | 4337 | 2 | 1 | 0 | 1 |
| UGC0004752 | Clust-UGC0002177 | 2226 | 4 | 1 | 1 | 1 |
| UGC0006491 | Clust-UGC0002177 | 1519 | 2 | 1 | 0 | 1 |
| UGC0011661 | Clust-UGC0002177 | 11020 | 73 | 1 | 0 | 1 |
| UGC0005605 | Clust-UGC0002177 | 3689 | 4 | 1 | 0 | 1 |
| UGC0003549 | Clust-UGC0002177 | 11215 | 72 | 1 | 0 | 1 |
| UGC0003527 | Clust-UGC0002177 | 2280 | 1 | 0 | 0 | 0 |
| UGC0011477 | Clust-UGC0002177 | 7781 | 21 | 1 | 0 | 1 |
| UGC0011643 | Clust-UGC0002177 | 2609 | 6 | 1 | 0 | 1 |
| UGC0004503 | Clust-UGC0002177 | 10413 | 47 | 1 | 0 | 1 |
| UGC0011994 | Clust-UGC0002177 | 581 | 1 | 0 | 0 | 0 |
| UGC0006783 | Clust-UGC0002177 | 7934 | 30 | 1 | 0 | 1 |
| UGC0009628 | Clust-UGC0002177 | 2373 | 2 | 1 | 0 | 1 |
| UGC0004549 | Clust-UGC0002177 | 5601 | 4 | 1 | 0 | 1 |

| | | | | | | |
|---|---|---|---|---|---|---|
| UGC0004545 | Clust-UGC0002177 | 9285 | 30 | 1 | 0 | 1 |
| UGC0004489 | Clust-UGC0002177 | 4745 | 8 | 1 | 0 | 1 |
| UGC0008466 | Clust-UGC0002177 | 1766 | 1 | 0 | 0 | 0 |
| UGC0004569 | Clust-UGC0002177 | 9941 | 40 | 1 | 0 | 1 |
| UGC0007550 | Clust-UGC0002177 | 4952 | 4 | 1 | 0 | 1 |
| UGC0011839 | Clust-UGC0002177 | 6434 | 19 | 1 | 1 | 1 |
| UGC0008715 | Clust-UGC0002177 | 2661 | 3 | 1 | 0 | 1 |
| UGC0012037 | Clust-UGC0002177 | 788 | 1 | 0 | 0 | 0 |
| UGC0007395 | Clust-UGC0002177 | 3643 | 4 | 1 | 0 | 1 |
| UGC0012604 | Clust-UGC0002177 | 2871 | 4 | 1 | 1 | 1 |
| UGC0005870 | Clust-UGC0002177 | 2190 | 2 | 1 | 0 | 1 |
| UGC0005875 | Clust-UGC0002177 | 3185 | 2 | 1 | 1 | 0 |
| UGC0008399 | Clust-UGC0002177 | 770 | 4 | 1 | 1 | 1 |
| UGC0007282 | Clust-UGC0002177 | 95 | 1 | 0 | 0 | 0 |
| UGC0003242 | Clust-UGC0002177 | 1297 | 1 | 0 | 0 | 0 |
| UGC0006804 | Clust-UGC0002177 | 7822 | 25 | 1 | 0 | 1 |
| UGC0008802 | Clust-UGC0002177 | 2666 | 6 | 1 | 0 | 1 |
| UGC0004681 | Clust-UGC0002177 | 15210 | 124 | 1 | 0 | 1 |
| UGC0004683 | Clust-UGC0002177 | 21379 | 213 | 1 | 0 | 1 |
| UGC0006684 | Clust-UGC0002177 | 1091 | 1 | 0 | 0 | 0 |
| UGC0009317 | Clust-UGC0002177 | 5079 | 10 | 1 | 1 | 1 |
| UGC0002237 | Clust-UGC0002179 | 8601 | 12 | 1 | 0 | 1 |
| UGC0002299 | Clust-UGC0002179 | 4897 | 8 | 1 | 0 | 1 |
| UGC0006448 | Clust-UGC0002179 | 1242 | 1 | 0 | 0 | 0 |
| UGC0010152 | Clust-UGC0002179 | 2337 | 2 | 1 | 1 | 0 |
| UGC0010247 | Clust-UGC0002179 | 5310 | 6 | 1 | 2 | 1 |
| UGC0010243 | Clust-UGC0002179 | 3765 | 5 | 1 | 0 | 1 |
| UGC0010155 | Clust-UGC0002179 | 426 | 1 | 0 | 0 | 0 |
| UGC0010229 | Clust-UGC0002179 | 185 | 1 | 0 | 0 | 0 |
| UGC0012677 | Clust-UGC0002179 | 14326 | 40 | 1 | 0 | 1 |
| UGC0010754 | Clust-UGC0002179 | 1878 | 3 | 1 | 0 | 0 |
| UGC0010548 | Clust-UGC0002179 | 4602 | 2 | 1 | 0 | 0 |
| UGC0013093 | Clust-UGC0002179 | 5439 | 6 | 1 | 1 | 1 |
| UGC0003346 | Clust-UGC0002179 | 15935 | 69 | 1 | 2 | 1 |
| UGC0003340 | Clust-UGC0002179 | 14714 | 37 | 1 | 0 | 1 |
| UGC0002250 | Clust-UGC0002179 | 1708 | 2 | 1 | 0 | 0 |
| UGC0004968 | Clust-UGC0002179 | 6068 | 6 | 1 | 0 | 1 |
| UGC0005698 | Clust-UGC0002179 | 5883 | 2 | 1 | 0 | 0 |
| UGC0010076 | Clust-UGC0002179 | 6545 | 7 | 1 | 0 | 1 |
| UGC0010073 | Clust-UGC0002179 | 768 | 1 | 0 | 0 | 0 |
| UGC0002179 | Clust-UGC0002179 | 5185 | 3 | 1 | 0 | 1 |
| UGC0012557 | Clust-UGC0002179 | 10167 | 19 | 1 | 0 | 1 |
| UGC0002277 | Clust-UGC0002179 | 5616 | 5 | 1 | 0 | 1 |
| UGC0006147 | Clust-UGC0002179 | 1989 | 3 | 1 | 0 | 0 |
| UGC0005792 | Clust-UGC0002179 | 9096 | 21 | 1 | 0 | 1 |
| UGC0010947 | Clust-UGC0002179 | 5108 | 4 | 1 | 1 | 0 |
| UGC0003322 | Clust-UGC0002179 | 5209 | 4 | 1 | 0 | 1 |
| UGC0003321 | Clust-UGC0002179 | 693 | 1 | 0 | 0 | 0 |
| UGC0010154 | Clust-UGC0002179 | 14583 | 41 | 1 | 0 | 1 |
| UGC0004171 | Clust-UGC0002179 | 2630 | 6 | 1 | 0 | 1 |
| UGC0004773 | Clust-UGC0002179 | 1928 | 2 | 1 | 0 | 0 |
| UGC0004772 | Clust-UGC0002179 | 1803 | 4 | 1 | 0 | 1 |
| UGC0010080 | Clust-UGC0002179 | 2638 | 1 | 0 | 0 | 0 |
| UGC0012109 | Clust-UGC0002179 | 109 | 1 | 0 | 0 | 0 |
| UGC0012813 | Clust-UGC0002179 | 101 | 1 | 0 | 0 | 0 |
| UGC0009762 | Clust-UGC0002179 | 3318 | 5 | 1 | 0 | 1 |
| UGC0006490 | Clust-UGC0002179 | 3945 | 1 | 0 | 0 | 0 |
| UGC0005066 | Clust-UGC0002179 | 3629 | 2 | 1 | 0 | 1 |
| UGC0010252 | Clust-UGC0002179 | 5098 | 10 | 1 | 0 | 1 |
| UGC0009746 | Clust-UGC0002179 | 2278 | 3 | 1 | 0 | 0 |
| UGC0002226 | Clust-UGC0002179 | 5475 | 4 | 1 | 0 | 1 |
| UGC0008691 | Clust-UGC0002179 | 2569 | 1 | 0 | 0 | 0 |
| UGC0002205 | Clust-UGC0002179 | 4625 | 5 | 1 | 0 | 1 |
| UGC0003334 | Clust-UGC0002179 | 5271 | 7 | 1 | 0 | 0 |
| UGC0003275 | Clust-UGC0002179 | 7660 | 13 | 1 | 0 | 1 |
| UGC0004488 | Clust-UGC0002179 | 6477 | 4 | 1 | 0 | 1 |
| UGC0005688 | Clust-UGC0002179 | 2372 | 3 | 1 | 0 | 1 |
| UGC0010875 | Clust-UGC0002179 | 2785 | 1 | 0 | 0 | 0 |
| UGC0010873 | Clust-UGC0002179 | 3071 | 6 | 1 | 1 | 1 |
| UGC0002266 | Clust-UGC0002179 | 8237 | 16 | 1 | 1 | 1 |
| UGC0008443 | Clust-UGC0002179 | 723 | 1 | 0 | 0 | 0 |
| UGC0009095 | Clust-UGC0002179 | 2690 | 2 | 1 | 0 | 0 |
| UGC0003555 | Clust-UGC0002179 | 6240 | 9 | 1 | 1 | 1 |
| UGC0010930 | Clust-UGC0002179 | 1569 | 1 | 0 | 0 | 0 |
| UGC0003361 | Clust-UGC0002179 | 8193 | 24 | 1 | 2 | 1 |
| UGC0005333 | Clust-UGC0002179 | 4502 | 3 | 1 | 0 | 1 |
| UGC0003258 | Clust-UGC0002179 | 13670 | 37 | 1 | 0 | 1 |
| UGC0008276 | Clust-UGC0002183 | 1087 | 1 | 0 | 0 | 0 |
| UGC0006484 | Clust-UGC0002183 | 2695 | 1 | 0 | 0 | 0 |

| | | | | | | |
|---|---|---|---|---|---|---|
| UGC0006799 | Clust-UGC0002183 | 7439 | 17 | 1 | 0 | 1 |
| UGC0009437 | Clust-UGC0002183 | 4954 | 5 | 1 | 0 | 1 |
| UGC0009647 | Clust-UGC0002183 | 804 | 1 | 0 | 0 | 0 |
| UGC0009354 | Clust-UGC0002183 | 7848 | 26 | 1 | 0 | 1 |
| UGC0007661 | Clust-UGC0002183 | 6384 | 13 | 1 | 0 | 1 |
| UGC0004580 | Clust-UGC0002183 | 5697 | 6 | 1 | 0 | 1 |
| UGC0011450 | Clust-UGC0002183 | 5738 | 7 | 1 | 0 | 1 |
| UGC0009469 | Clust-UGC0002183 | 3171 | 1 | 0 | 0 | 0 |
| UGC0007178 | Clust-UGC0002183 | 2393 | 2 | 1 | 0 | 1 |
| UGC0009070 | Clust-UGC0002183 | 4787 | 4 | 1 | 0 | 1 |
| UGC0002183 | Clust-UGC0002183 | 5483 | 4 | 1 | 0 | 1 |
| UGC0006562 | Clust-UGC0002183 | 3219 | 2 | 1 | 0 | 0 |
| UGC0011193 | Clust-UGC0002183 | 598 | 1 | 0 | 0 | 0 |
| UGC0007202 | Clust-UGC0002184 | 3082 | 3 | 1 | 0 | 1 |
| UGC0004827 | Clust-UGC0002184 | 13556 | 66 | 1 | 0 | 1 |
| UGC0012576 | Clust-UGC0002184 | 8717 | 39 | 1 | 0 | 1 |
| UGC0008641 | Clust-UGC0002184 | 3525 | 10 | 1 | 0 | 1 |
| UGC0008292 | Clust-UGC0002184 | 1816 | 6 | 1 | 0 | 1 |
| UGC0009161 | Clust-UGC0002184 | 5375 | 4 | 1 | 0 | 1 |
| UGC0010921 | Clust-UGC0002184 | 1214 | 1 | 0 | 0 | 0 |
| UGC0007099 | Clust-UGC0002184 | 3015 | 1 | 0 | 0 | 0 |
| UGC0008307 | Clust-UGC0002184 | 1278 | 3 | 1 | 0 | 1 |
| UGC0008262 | Clust-UGC0002184 | 2142 | 6 | 1 | 0 | 1 |
| UGC0007055 | Clust-UGC0002184 | 3077 | 3 | 1 | 0 | 1 |
| UGC0010608 | Clust-UGC0002184 | 3016 | 8 | 1 | 0 | 1 |
| UGC0012898 | Clust-UGC0002184 | 7495 | 17 | 1 | 0 | 1 |
| UGC0012546 | Clust-UGC0002184 | 4112 | 11 | 1 | 0 | 1 |
| UGC0008900 | Clust-UGC0002184 | 4260 | 9 | 1 | 0 | 1 |
| UGC0002184 | Clust-UGC0002184 | 5453 | 10 | 1 | 0 | 1 |
| UGC0005536 | Clust-UGC0002184 | 2487 | 6 | 1 | 0 | 1 |
| UGC0011176 | Clust-UGC0002193 | 6145 | 15 | 1 | 0 | 1 |
| UGC0005378 | Clust-UGC0002193 | 1899 | 1 | 0 | 0 | 0 |
| UGC0012614 | Clust-UGC0002193 | 4510 | 3 | 1 | 0 | 1 |
| UGC0009529 | Clust-UGC0002193 | 5747 | 10 | 1 | 0 | 1 |
| UGC0009776 | Clust-UGC0002193 | 1401 | 2 | 1 | 0 | 0 |
| UGC0005896 | Clust-UGC0002193 | 2722 | 1 | 0 | 0 | 0 |
| UGC0006274 | Clust-UGC0002193 | 2922 | 2 | 1 | 0 | 1 |
| UGC0012033 | Clust-UGC0002193 | 1343 | 1 | 0 | 0 | 0 |
| UGC0004705 | Clust-UGC0002193 | 18758 | 184 | 1 | 0 | 1 |
| UGC0010752 | Clust-UGC0002193 | 1643 | 4 | 1 | 0 | 0 |
| UGC0011987 | Clust-UGC0002193 | 1171 | 1 | 0 | 0 | 0 |
| UGC0008596 | Clust-UGC0002193 | 5314 | 7 | 1 | 0 | 1 |
| UGC0004634 | Clust-UGC0002193 | 4156 | 3 | 1 | 0 | 1 |
| UGC0002193 | Clust-UGC0002193 | 110 | 1 | 0 | 0 | 0 |
| UGC0005762 | Clust-UGC0002193 | 275 | 1 | 0 | 0 | 0 |
| UGC0003388 | Clust-UGC0002193 | 8554 | 32 | 1 | 0 | 1 |
| UGC0005840 | Clust-UGC0002193 | 2462 | 1 | 0 | 0 | 0 |
| UGC0013310 | Clust-UGC0002193 | 6075 | 10 | 1 | 0 | 1 |
| UGC0009640 | Clust-UGC0002193 | 1932 | 4 | 1 | 0 | 1 |
| UGC0008837 | Clust-UGC0002193 | 986 | 1 | 0 | 0 | 0 |
| UGC0006690 | Clust-UGC0002193 | 1877 | 1 | 0 | 0 | 0 |
| UGC0007565 | Clust-UGC0002193 | 6820 | 9 | 1 | 0 | 1 |
| UGC0013127 | Clust-UGC0002193 | 4148 | 5 | 1 | 0 | 1 |
| UGC0008325 | Clust-UGC0002193 | 1899 | 3 | 1 | 0 | 1 |
| UGC0011937 | Clust-UGC0002193 | 4303 | 4 | 1 | 0 | 1 |
| UGC0004771 | Clust-UGC0002193 | 1098 | 1 | 0 | 0 | 0 |
| UGC0006287 | Clust-UGC0002193 | 1772 | 2 | 1 | 0 | 1 |
| UGC0002368 | Clust-UGC0002193 | 542 | 1 | 0 | 0 | 0 |
| UGC0003717 | Clust-UGC0002193 | 258 | 1 | 0 | 0 | 0 |
| UGC0012319 | Clust-UGC0002193 | 3017 | 2 | 1 | 0 | 0 |
| UGC0006720 | Clust-UGC0002193 | 1029 | 1 | 0 | 0 | 0 |
| UGC0009576 | Clust-UGC0002193 | 3795 | 2 | 1 | 0 | 0 |
| UGC0012974 | Clust-UGC0002193 | 5112 | 14 | 1 | 0 | 1 |
| UGC0010874 | Clust-UGC0002193 | 983 | 1 | 0 | 0 | 0 |
| UGC0010305 | Clust-UGC0002193 | 5482 | 3 | 1 | 0 | 1 |
| UGC0011143 | Clust-UGC0002193 | 2801 | 2 | 1 | 0 | 1 |
| UGC0003604 | Clust-UGC0002193 | 7795 | 29 | 1 | 0 | 1 |
| UGC0005856 | Clust-UGC0002193 | 3969 | 2 | 1 | 0 | 1 |
| UGC0010598 | Clust-UGC0002193 | 5266 | 11 | 1 | 1 | 1 |
| UGC0009488 | Clust-UGC0002193 | 1583 | 1 | 0 | 0 | 0 |
| UGC0009508 | Clust-UGC0002194 | 355 | 1 | 0 | 0 | 0 |
| UGC0003722 | Clust-UGC0002194 | 836 | 1 | 0 | 0 | 0 |
| UGC0011791 | Clust-UGC0002194 | 8173 | 5 | 1 | 0 | 1 |
| UGC0009143 | Clust-UGC0002194 | 3286 | 4 | 1 | 0 | 1 |
| UGC0013243 | Clust-UGC0002194 | 1341 | 1 | 0 | 0 | 0 |
| UGC0004514 | Clust-UGC0002194 | 6230 | 3 | 1 | 0 | 1 |
| UGC0005693 | Clust-UGC0002194 | 1103 | 1 | 0 | 0 | 0 |
| UGC0002194 | Clust-UGC0002194 | 390 | 1 | 0 | 0 | 0 |
| UGC0008620 | Clust-UGC0002194 | 3077 | 2 | 1 | 0 | 1 |

| | | | | | | |
|---|---|---|---|---|---|---|
| UGC0009494 | Clust-UGC0002194 | 3489 | 1 | 0 | 0 | 0 |
| UGC0009493 | Clust-UGC0002194 | 1332 | 1 | 0 | 0 | 0 |
| UGC0012142 | Clust-UGC0002194 | 1518 | 1 | 0 | 0 | 0 |
| UGC0008540 | Clust-UGC0002194 | 3467 | 1 | 0 | 0 | 0 |
| UGC0002365 | Clust-UGC0002194 | 4157 | 1 | 0 | 0 | 0 |
| UGC0008347 | Clust-UGC0002194 | 1617 | 2 | 1 | 0 | 1 |
| UGC0009761 | Clust-UGC0002194 | 2883 | 2 | 1 | 0 | 1 |
| UGC0009760 | Clust-UGC0002194 | 683 | 1 | 0 | 0 | 0 |
| UGC0005907 | Clust-UGC0002194 | 2057 | 2 | 1 | 0 | 1 |
| UGC0013042 | Clust-UGC0002194 | 6673 | 5 | 1 | 0 | 1 |
| UGC0009487 | Clust-UGC0002194 | 2708 | 1 | 0 | 0 | 0 |
| UGC0009056 | Clust-UGC0002194 | 4117 | 2 | 1 | 0 | 1 |
| UGC0004618 | Clust-UGC0002194 | 3172 | 1 | 0 | 0 | 0 |
| UGC0012580 | Clust-UGC0002194 | 7179 | 2 | 1 | 0 | 0 |
| UGC0004619 | Clust-UGC0002194 | 7941 | 4 | 1 | 0 | 1 |
| UGC0003554 | Clust-UGC0002194 | 6987 | 3 | 1 | 0 | 1 |
| UGC0009157 | Clust-UGC0002194 | 680 | 1 | 0 | 0 | 0 |
| UGC0013119 | Clust-UGC0002194 | 1004 | 1 | 0 | 0 | 0 |
| UGC0009677 | Clust-UGC0002194 | 2449 | 1 | 0 | 0 | 0 |
| UGC0010955 | Clust-UGC0002194 | 3905 | 4 | 1 | 0 | 1 |
| UGC0005994 | Clust-UGC0002199 | 5398 | 2 | 1 | 0 | 1 |
| UGC0009543 | Clust-UGC0002199 | 3933 | 1 | 0 | 0 | 0 |
| UGC0007428 | Clust-UGC0002199 | 898 | 1 | 0 | 0 | 0 |
| UGC0007308 | Clust-UGC0002199 | 3782 | 3 | 1 | 0 | 1 |
| UGC0002199 | Clust-UGC0002199 | 61 | 1 | 0 | 0 | 0 |
| UGC0009107 | Clust-UGC0002199 | 1042 | 1 | 0 | 0 | 0 |
| UGC0006328 | Clust-UGC0002199 | 3981 | 3 | 1 | 0 | 1 |
| UGC0005981 | Clust-UGC0002199 | 1700 | 2 | 1 | 0 | 1 |
| UGC0006153 | Clust-UGC0002199 | 1640 | 1 | 0 | 0 | 0 |
| UGC0011545 | Clust-UGC0002200 | 8304 | 5 | 1 | 0 | 1 |
| UGC0008845 | Clust-UGC0002200 | 1723 | 1 | 0 | 0 | 0 |
| UGC0008760 | Clust-UGC0002200 | 1340 | 1 | 0 | 0 | 0 |
| UGC0013018 | Clust-UGC0002200 | 1259 | 1 | 0 | 0 | 0 |
| UGC0011468 | Clust-UGC0002200 | 8052 | 12 | 1 | 0 | 1 |
| UGC0012849 | Clust-UGC0002200 | 3300 | 3 | 1 | 0 | 0 |
| UGC0012944 | Clust-UGC0002200 | 8626 | 5 | 1 | 0 | 1 |
| UGC0008935 | Clust-UGC0002200 | 2548 | 2 | 1 | 0 | 0 |
| UGC0002200 | Clust-UGC0002200 | 732 | 1 | 0 | 0 | 0 |
| UGC0013334 | Clust-UGC0002200 | 9111 | 10 | 1 | 0 | 1 |
| UGC0009124 | Clust-UGC0002200 | 3155 | 1 | 0 | 0 | 0 |
| UGC0013147 | Clust-UGC0002200 | 475 | 2 | 1 | 0 | 0 |
| UGC0009305 | Clust-UGC0002200 | 1737 | 1 | 0 | 0 | 0 |
| UGC0008857 | Clust-UGC0002200 | 2543 | 2 | 1 | 0 | 0 |
| UGC0008711 | Clust-UGC0002200 | 163 | 1 | 0 | 0 | 0 |
| UGC0009700 | Clust-UGC0002200 | 1953 | 1 | 0 | 0 | 0 |
| UGC0010491 | Clust-UGC0002200 | 2863 | 1 | 0 | 0 | 0 |
| UGC0008658 | Clust-UGC0002200 | 414 | 2 | 1 | 0 | 0 |
| UGC0008073 | Clust-UGC0002200 | 775 | 1 | 0 | 0 | 0 |
| UGC0008583 | Clust-UGC0002200 | 2020 | 1 | 0 | 0 | 0 |
| UGC0012600 | Clust-UGC0002200 | 3518 | 2 | 1 | 0 | 0 |
| UGC0012268 | Clust-UGC0002200 | 1061 | 2 | 1 | 0 | 0 |
| UGC0005115 | Clust-UGC0002200 | 754 | 2 | 1 | 0 | 1 |
| UGC0013349 | Clust-UGC0002200 | 6416 | 3 | 1 | 1 | 0 |
| UGC0008785 | Clust-UGC0002200 | 2622 | 1 | 0 | 0 | 0 |
| UGC0006167 | Clust-UGC0002202 | 553 | 1 | 0 | 0 | 0 |
| UGC0002202 | Clust-UGC0002202 | 81 | 1 | 0 | 0 | 0 |
| UGC0008481 | Clust-UGC0002203 | 95 | 1 | 0 | 0 | 0 |
| UGC0006026 | Clust-UGC0002203 | 695 | 1 | 0 | 0 | 0 |
| UGC0002992 | Clust-UGC0002203 | 1378 | 1 | 0 | 0 | 0 |
| UGC0008726 | Clust-UGC0002203 | 1232 | 1 | 0 | 0 | 0 |
| UGC0008628 | Clust-UGC0002203 | 4000 | 1 | 0 | 0 | 0 |
| UGC0013380 | Clust-UGC0002203 | 4867 | 4 | 1 | 0 | 1 |
| UGC0006761 | Clust-UGC0002203 | 4229 | 1 | 0 | 0 | 0 |
| UGC0008742 | Clust-UGC0002203 | 3315 | 1 | 0 | 0 | 0 |
| UGC0009067 | Clust-UGC0002203 | 2282 | 1 | 0 | 0 | 0 |
| UGC0008630 | Clust-UGC0002203 | 5910 | 3 | 1 | 0 | 1 |
| UGC0004948 | Clust-UGC0002203 | 9311 | 12 | 1 | 0 | 1 |
| UGC0002528 | Clust-UGC0002203 | 1147 | 1 | 0 | 0 | 0 |
| UGC0004638 | Clust-UGC0002203 | 4163 | 3 | 1 | 1 | 1 |
| UGC0002946 | Clust-UGC0002203 | 2278 | 1 | 0 | 0 | 0 |
| UGC0008403 | Clust-UGC0002203 | 239 | 1 | 0 | 0 | 0 |
| UGC0011841 | Clust-UGC0002203 | 2708 | 1 | 0 | 0 | 0 |
| UGC0006879 | Clust-UGC0002203 | 5633 | 1 | 0 | 0 | 0 |
| UGC0006810 | Clust-UGC0002203 | 7724 | 6 | 1 | 0 | 1 |
| UGC0006100 | Clust-UGC0002203 | 1309 | 1 | 0 | 0 | 0 |
| UGC0004087 | Clust-UGC0002203 | 3604 | 1 | 0 | 0 | 0 |
| UGC0008734 | Clust-UGC0002203 | 896 | 1 | 0 | 0 | 0 |
| UGC0002983 | Clust-UGC0002203 | 2707 | 1 | 0 | 0 | 0 |
| UGC0004759 | Clust-UGC0002203 | 2266 | 1 | 0 | 0 | 0 |

| | | | | | | |
|---|---|---|---|---|---|---|
| UGC0007628 | Clust-UGC0002203 | 4517 | 2 | 1 | 0 | 1 |
| UGC0008719 | Clust-UGC0002203 | 2453 | 1 | 0 | 0 | 0 |
| UGC0008502 | Clust-UGC0002203 | 1152 | 1 | 0 | 0 | 0 |
| UGC0011642 | Clust-UGC0002203 | 3170 | 2 | 1 | 0 | 0 |
| UGC0007997 | Clust-UGC0002203 | 3498 | 1 | 0 | 0 | 0 |
| UGC0010231 | Clust-UGC0002203 | 589 | 1 | 0 | 0 | 0 |
| UGC0003315 | Clust-UGC0002203 | 2305 | 1 | 0 | 0 | 0 |
| UGC0002203 | Clust-UGC0002203 | 3392 | 1 | 0 | 0 | 0 |
| UGC0007789 | Clust-UGC0002203 | 2618 | 1 | 0 | 0 | 0 |
| UGC0009449 | Clust-UGC0002203 | 4167 | 2 | 1 | 0 | 1 |
| UGC0008670 | Clust-UGC0002203 | 106 | 1 | 0 | 0 | 0 |
| UGC0003588 | Clust-UGC0002203 | 11071 | 17 | 1 | 0 | 1 |
| UGC0008476 | Clust-UGC0002203 | 1517 | 1 | 0 | 0 | 0 |
| UGC0003294 | Clust-UGC0002203 | 6068 | 3 | 1 | 0 | 1 |
| UGC0010596 | Clust-UGC0002203 | 2992 | 1 | 0 | 0 | 0 |
| UGC0008808 | Clust-UGC0002203 | 1039 | 1 | 0 | 0 | 0 |
| UGC0010995 | Clust-UGC0002204 | 8778 | 22 | 1 | 0 | 1 |
| UGC0002298 | Clust-UGC0002204 | 5800 | 4 | 1 | 0 | 1 |
| UGC0002297 | Clust-UGC0002204 | 5599 | 4 | 1 | 0 | 1 |
| UGC0004123 | Clust-UGC0002204 | 551 | 1 | 0 | 0 | 0 |
| UGC0012307 | Clust-UGC0002204 | 2449 | 2 | 1 | 0 | 1 |
| UGC0011988 | Clust-UGC0002204 | 4235 | 2 | 1 | 0 | 1 |
| UGC0002218 | Clust-UGC0002204 | 9025 | 20 | 1 | 0 | 1 |
| UGC0010843 | Clust-UGC0002204 | 5522 | 4 | 1 | 0 | 1 |
| UGC0010848 | Clust-UGC0002204 | 1908 | 2 | 1 | 0 | 0 |
| UGC0013097 | Clust-UGC0002204 | 2250 | 1 | 0 | 0 | 0 |
| UGC0012380 | Clust-UGC0002204 | 4594 | 3 | 1 | 0 | 1 |
| UGC0002257 | Clust-UGC0002204 | 2420 | 3 | 1 | 0 | 1 |
| UGC0012773 | Clust-UGC0002204 | 3518 | 2 | 1 | 0 | 1 |
| UGC0006460 | Clust-UGC0002204 | 6747 | 8 | 1 | 0 | 1 |
| UGC0011863 | Clust-UGC0002204 | 5244 | 2 | 1 | 0 | 1 |
| UGC0011869 | Clust-UGC0002204 | 10501 | 48 | 1 | 0 | 1 |
| UGC0006104 | Clust-UGC0002204 | 2211 | 2 | 1 | 0 | 0 |
| UGC0002278 | Clust-UGC0002204 | 6569 | 6 | 1 | 0 | 1 |
| UGC0011971 | Clust-UGC0002204 | 2829 | 1 | 0 | 0 | 0 |
| UGC0011972 | Clust-UGC0002204 | 4072 | 2 | 1 | 0 | 1 |
| UGC0006142 | Clust-UGC0002204 | 2446 | 2 | 1 | 0 | 1 |
| UGC0011160 | Clust-UGC0002204 | 3727 | 4 | 1 | 0 | 1 |
| UGC0012140 | Clust-UGC0002204 | 4450 | 2 | 1 | 0 | 1 |
| UGC0011931 | Clust-UGC0002204 | 10935 | 28 | 1 | 0 | 1 |
| UGC0013214 | Clust-UGC0002204 | 7477 | 9 | 1 | 0 | 1 |
| UGC0012817 | Clust-UGC0002204 | 1671 | 2 | 1 | 0 | 1 |
| UGC0012315 | Clust-UGC0002204 | 3287 | 2 | 1 | 0 | 1 |
| UGC0006005 | Clust-UGC0002204 | 1079 | 1 | 0 | 0 | 0 |
| UGC0012122 | Clust-UGC0002204 | 404 | 2 | 1 | 0 | 0 |
| UGC0013256 | Clust-UGC0002204 | 7282 | 5 | 1 | 0 | 1 |
| UGC0002255 | Clust-UGC0002204 | 11767 | 51 | 1 | 0 | 1 |
| UGC0009257 | Clust-UGC0002204 | 13301 | 103 | 1 | 0 | 1 |
| UGC0002204 | Clust-UGC0002204 | 7018 | 11 | 1 | 0 | 1 |
| UGC0013046 | Clust-UGC0002204 | 9361 | 29 | 1 | 1 | 1 |
| UGC0011819 | Clust-UGC0002204 | 6283 | 7 | 1 | 0 | 1 |
| UGC0002245 | Clust-UGC0002204 | 6493 | 6 | 1 | 0 | 1 |
| UGC0003371 | Clust-UGC0002204 | 1712 | 2 | 1 | 0 | 1 |
| UGC0002265 | Clust-UGC0002204 | 6914 | 8 | 1 | 0 | 1 |
| UGC0002267 | Clust-UGC0002204 | 12594 | 80 | 1 | 0 | 1 |
| UGC0002263 | Clust-UGC0002204 | 10542 | 39 | 1 | 0 | 1 |
| UGC0011142 | Clust-UGC0002204 | 1204 | 1 | 0 | 0 | 0 |
| UGC0011871 | Clust-UGC0002204 | 9264 | 18 | 1 | 0 | 1 |
| UGC0005831 | Clust-UGC0002204 | 3769 | 2 | 1 | 0 | 1 |
| UGC0013110 | Clust-UGC0002204 | 2564 | 2 | 1 | 0 | 1 |
| UGC0010999 | Clust-UGC0002206 | 920 | 1 | 0 | 0 | 0 |
| UGC0004596 | Clust-UGC0002206 | 2467 | 2 | 1 | 0 | 1 |
| UGC0006290 | Clust-UGC0002206 | 1831 | 1 | 0 | 0 | 0 |
| UGC0012026 | Clust-UGC0002206 | 3964 | 2 | 1 | 0 | 1 |
| UGC0004797 | Clust-UGC0002206 | 8250 | 17 | 1 | 0 | 1 |
| UGC0005635 | Clust-UGC0002206 | 5280 | 5 | 1 | 0 | 1 |
| UGC0007365 | Clust-UGC0002206 | 2173 | 1 | 0 | 0 | 0 |
| UGC0013262 | Clust-UGC0002206 | 10327 | 47 | 1 | 1 | 1 |
| UGC0012946 | Clust-UGC0002206 | 1363 | 1 | 0 | 0 | 0 |
| UGC0006124 | Clust-UGC0002206 | 1767 | 1 | 0 | 0 | 0 |
| UGC0002211 | Clust-UGC0002206 | 12884 | 90 | 1 | 0 | 1 |
| UGC0004553 | Clust-UGC0002206 | 7797 | 19 | 1 | 0 | 1 |
| UGC0003345 | Clust-UGC0002206 | 20934 | 192 | 1 | 0 | 1 |
| UGC0006423 | Clust-UGC0002206 | 6548 | 9 | 1 | 0 | 1 |
| UGC0005127 | Clust-UGC0002206 | 4661 | 2 | 1 | 0 | 1 |
| UGC0006343 | Clust-UGC0002206 | 4534 | 2 | 1 | 0 | 1 |
| UGC0006146 | Clust-UGC0002206 | 2967 | 2 | 1 | 0 | 1 |
| UGC0004174 | Clust-UGC0002206 | 3522 | 3 | 1 | 0 | 1 |
| UGC0006988 | Clust-UGC0002206 | 5282 | 9 | 1 | 0 | 1 |

| | | | | | | |
|---|---|---|---|---|---|---|
| UGC0005344 | Clust-UGC0002206 | 5463 | 6 | 1 | 0 | 1 |
| UGC0007627 | Clust-UGC0002206 | 5484 | 6 | 1 | 0 | 1 |
| UGC0012818 | Clust-UGC0002206 | 1818 | 2 | 1 | 0 | 1 |
| UGC0012313 | Clust-UGC0002206 | 2525 | 1 | 0 | 0 | 0 |
| UGC0007299 | Clust-UGC0002206 | 7408 | 19 | 1 | 0 | 1 |
| UGC0002252 | Clust-UGC0002206 | 4876 | 4 | 1 | 0 | 1 |
| UGC0004501 | Clust-UGC0002206 | 9072 | 35 | 1 | 0 | 1 |
| UGC0002221 | Clust-UGC0002206 | 10601 | 34 | 1 | 1 | 1 |
| UGC0003312 | Clust-UGC0002206 | 2044 | 3 | 1 | 0 | 1 |
| UGC0005002 | Clust-UGC0002206 | 7648 | 13 | 1 | 0 | 1 |
| UGC0005949 | Clust-UGC0002206 | 1819 | 2 | 1 | 0 | 1 |
| UGC0002268 | Clust-UGC0002206 | 17864 | 156 | 1 | 0 | 1 |
| UGC0003351 | Clust-UGC0002206 | 12000 | 72 | 1 | 0 | 1 |
| UGC0012643 | Clust-UGC0002206 | 367 | 2 | 1 | 0 | 0 |
| UGC0005745 | Clust-UGC0002206 | 5528 | 6 | 1 | 0 | 1 |
| UGC0006082 | Clust-UGC0002206 | 5577 | 4 | 1 | 0 | 1 |
| UGC0007685 | Clust-UGC0002206 | 6467 | 11 | 1 | 1 | 1 |
| UGC0007688 | Clust-UGC0002206 | 9382 | 42 | 1 | 0 | 1 |
| UGC0010512 | Clust-UGC0002206 | 4960 | 6 | 1 | 0 | 1 |
| UGC0010516 | Clust-UGC0002206 | 592 | 1 | 0 | 0 | 0 |
| UGC0002206 | Clust-UGC0002206 | 590 | 1 | 0 | 0 | 0 |
| UGC0012200 | Clust-UGC0002206 | 11236 | 55 | 1 | 0 | 1 |
| UGC0006113 | Clust-UGC0002206 | 1428 | 1 | 0 | 0 | 0 |
| UGC0012863 | Clust-UGC0002210 | 2460 | 2 | 1 | 0 | 0 |
| UGC0010820 | Clust-UGC0002210 | 1438 | 4 | 1 | 0 | 0 |
| UGC0002210 | Clust-UGC0002210 | 9380 | 21 | 1 | 0 | 1 |
| UGC0005542 | Clust-UGC0002210 | 408 | 1 | 0 | 0 | 0 |
| UGC0012774 | Clust-UGC0002210 | 4654 | 13 | 1 | 0 | 1 |
| UGC0002276 | Clust-UGC0002210 | 6912 | 12 | 1 | 0 | 1 |
| UGC0013041 | Clust-UGC0002210 | 15739 | 121 | 1 | 0 | 1 |
| UGC0002248 | Clust-UGC0002210 | 13311 | 77 | 1 | 0 | 1 |
| UGC0002227 | Clust-UGC0002210 | 12598 | 54 | 1 | 0 | 1 |
| UGC0010978 | Clust-UGC0002210 | 6528 | 10 | 1 | 0 | 1 |
| UGC0004786 | Clust-UGC0002213 | 6718 | 14 | 1 | 0 | 1 |
| UGC0007041 | Clust-UGC0002213 | 1677 | 2 | 1 | 0 | 1 |
| UGC0002337 | Clust-UGC0002213 | 722 | 1 | 0 | 0 | 0 |
| UGC0011650 | Clust-UGC0002213 | 4585 | 7 | 1 | 0 | 1 |
| UGC0003517 | Clust-UGC0002213 | 1053 | 1 | 0 | 0 | 0 |
| UGC0002213 | Clust-UGC0002213 | 15326 | 115 | 1 | 0 | 1 |
| UGC0008005 | Clust-UGC0002213 | 6906 | 17 | 1 | 0 | 1 |
| UGC0002962 | Clust-UGC0002213 | 3859 | 1 | 0 | 0 | 0 |
| UGC0004586 | Clust-UGC0002213 | 6862 | 21 | 1 | 0 | 1 |
| UGC0008856 | Clust-UGC0002213 | 1133 | 2 | 1 | 0 | 1 |
| UGC0004776 | Clust-UGC0002213 | 661 | 1 | 0 | 0 | 0 |
| UGC0007972 | Clust-UGC0002213 | 1011 | 2 | 1 | 0 | 1 |
| UGC0004753 | Clust-UGC0002213 | 336 | 1 | 0 | 0 | 0 |
| UGC0006647 | Clust-UGC0002213 | 4550 | 2 | 1 | 0 | 1 |
| UGC0009422 | Clust-UGC0002213 | 8496 | 25 | 1 | 0 | 1 |
| UGC0003330 | Clust-UGC0002213 | 1431 | 3 | 1 | 0 | 1 |
| UGC0003794 | Clust-UGC0002213 | 721 | 1 | 0 | 0 | 0 |
| UGC0009370 | Clust-UGC0002213 | 2520 | 3 | 1 | 0 | 1 |
| UGC0003732 | Clust-UGC0002213 | 749 | 1 | 0 | 0 | 0 |
| UGC0004780 | Clust-UGC0002214 | 4585 | 1 | 0 | 0 | 0 |
| UGC0002291 | Clust-UGC0002214 | 4420 | 1 | 0 | 0 | 0 |
| UGC0002235 | Clust-UGC0002214 | 9139 | 7 | 1 | 0 | 1 |
| UGC0002214 | Clust-UGC0002214 | 8276 | 6 | 1 | 0 | 1 |
| UGC0011824 | Clust-UGC0002214 | 2899 | 1 | 0 | 0 | 0 |
| UGC0012351 | Clust-UGC0002214 | 5014 | 2 | 1 | 0 | 1 |
| UGC0010884 | Clust-UGC0002214 | 4415 | 3 | 1 | 0 | 1 |
| UGC0003015 | Clust-UGC0002214 | 2362 | 2 | 1 | 1 | 0 |
| UGC0004168 | Clust-UGC0002216 | 5102 | 6 | 1 | 1 | 1 |
| UGC0004161 | Clust-UGC0002216 | 3739 | 4 | 1 | 0 | 1 |
| UGC0002316 | Clust-UGC0002216 | 3099 | 3 | 1 | 0 | 1 |
| UGC0012025 | Clust-UGC0002216 | 2647 | 4 | 1 | 0 | 1 |
| UGC0002233 | Clust-UGC0002216 | 19310 | 168 | 1 | 0 | 1 |
| UGC0011156 | Clust-UGC0002216 | 6069 | 12 | 1 | 0 | 1 |
| UGC0002292 | Clust-UGC0002216 | 7201 | 6 | 1 | 0 | 1 |
| UGC0012809 | Clust-UGC0002216 | 1407 | 4 | 1 | 0 | 1 |
| UGC0012135 | Clust-UGC0002216 | 6810 | 11 | 1 | 0 | 1 |
| UGC0012366 | Clust-UGC0002216 | 11847 | 78 | 1 | 0 | 1 |
| UGC0004532 | Clust-UGC0002216 | 2771 | 2 | 1 | 0 | 1 |
| UGC0002279 | Clust-UGC0002216 | 5686 | 4 | 1 | 0 | 1 |
| UGC0002216 | Clust-UGC0002216 | 10016 | 36 | 1 | 0 | 1 |
| UGC0010840 | Clust-UGC0002216 | 8861 | 20 | 1 | 0 | 1 |
| UGC0013095 | Clust-UGC0002216 | 2925 | 4 | 1 | 0 | 1 |
| UGC0002258 | Clust-UGC0002216 | 14033 | 87 | 1 | 0 | 1 |
| UGC0003380 | Clust-UGC0002216 | 8984 | 18 | 1 | 0 | 1 |
| UGC0012004 | Clust-UGC0002216 | 2207 | 3 | 1 | 0 | 1 |
| UGC0012003 | Clust-UGC0002216 | 4807 | 3 | 1 | 0 | 1 |

| | | | | | | |
|---|---|---|---|---|---|---|
| UGC0005847 | Clust-UGC0002216 | 5314 | 4 | 1 | 0 | 1 |
| UGC0012273 | Clust-UGC0002216 | 2001 | 2 | 1 | 0 | 0 |
| UGC0006341 | Clust-UGC0002216 | 5697 | 5 | 1 | 0 | 1 |
| UGC0003406 | Clust-UGC0002216 | 2103 | 4 | 1 | 0 | 1 |
| UGC0002273 | Clust-UGC0002216 | 6427 | 7 | 1 | 0 | 1 |
| UGC0011976 | Clust-UGC0002216 | 3896 | 6 | 1 | 0 | 1 |
| UGC0006145 | Clust-UGC0002216 | 2099 | 1 | 0 | 0 | 0 |
| UGC0007897 | Clust-UGC0002216 | 9855 | 49 | 1 | 0 | 1 |
| UGC0002301 | Clust-UGC0002216 | 11241 | 55 | 1 | 0 | 1 |
| UGC0011163 | Clust-UGC0002216 | 1714 | 4 | 1 | 0 | 1 |
| UGC0004172 | Clust-UGC0002216 | 8063 | 11 | 1 | 0 | 1 |
| UGC0002264 | Clust-UGC0002216 | 14570 | 92 | 1 | 0 | 1 |
| UGC0012167 | Clust-UGC0002216 | 3250 | 1 | 0 | 0 | 0 |
| UGC0007644 | Clust-UGC0002216 | 6266 | 11 | 1 | 0 | 1 |
| UGC0013089 | Clust-UGC0002216 | 2584 | 2 | 1 | 0 | 0 |
| UGC0004754 | Clust-UGC0002216 | 2788 | 1 | 0 | 0 | 0 |
| UGC0013219 | Clust-UGC0002216 | 4165 | 1 | 0 | 0 | 0 |
| UGC0007603 | Clust-UGC0002216 | 8621 | 20 | 1 | 0 | 1 |
| UGC0002256 | Clust-UGC0002216 | 12591 | 64 | 1 | 0 | 1 |
| UGC0011815 | Clust-UGC0002216 | 4724 | 2 | 1 | 0 | 1 |
| UGC0002223 | Clust-UGC0002216 | 10530 | 36 | 1 | 0 | 1 |
| UGC0002225 | Clust-UGC0002216 | 11874 | 48 | 1 | 0 | 1 |
| UGC0002229 | Clust-UGC0002216 | 1761 | 1 | 0 | 0 | 0 |
| UGC0003316 | Clust-UGC0002216 | 9177 | 17 | 1 | 0 | 1 |
| UGC0009258 | Clust-UGC0002216 | 12952 | 114 | 1 | 0 | 1 |
| UGC0009629 | Clust-UGC0002216 | 2814 | 1 | 0 | 0 | 0 |
| UGC0004540 | Clust-UGC0002216 | 6752 | 16 | 1 | 0 | 1 |
| UGC0004481 | Clust-UGC0002216 | 4126 | 5 | 1 | 1 | 1 |
| UGC0003377 | Clust-UGC0002216 | 5236 | 8 | 1 | 0 | 1 |
| UGC0002970 | Clust-UGC0002216 | 1569 | 3 | 1 | 0 | 1 |
| UGC0012306 | Clust-UGC0002216 | 2853 | 4 | 1 | 0 | 1 |
| UGC0008960 | Clust-UGC0002216 | 722 | 1 | 0 | 0 | 0 |
| UGC0010932 | Clust-UGC0002216 | 6391 | 3 | 1 | 0 | 1 |
| UGC0006139 | Clust-UGC0002216 | 1231 | 2 | 1 | 0 | 0 |
| UGC0006175 | Clust-UGC0002216 | 511 | 1 | 0 | 0 | 0 |
| UGC0011190 | Clust-UGC0002216 | 3443 | 2 | 1 | 0 | 1 |
| UGC0002219 | Clust-UGC0002219 | 12626 | 57 | 1 | 0 | 1 |
| UGC0002232 | Clust-UGC0002232 | 10929 | 29 | 1 | 0 | 1 |
| UGC0011900 | Clust-UGC0002232 | 3928 | 1 | 0 | 0 | 0 |
| UGC0006585 | Clust-UGC0002232 | 3171 | 8 | 1 | 0 | 1 |
| UGC0006327 | Clust-UGC0002232 | 3016 | 1 | 0 | 0 | 0 |
| UGC0009438 | Clust-UGC0002232 | 1287 | 2 | 1 | 0 | 0 |
| UGC0013024 | Clust-UGC0002232 | 1771 | 2 | 1 | 1 | 0 |
| UGC0006459 | Clust-UGC0002232 | 4071 | 3 | 1 | 0 | 1 |
| UGC0011929 | Clust-UGC0002236 | 5690 | 10 | 1 | 1 | 1 |
| UGC0009773 | Clust-UGC0002236 | 2693 | 12 | 1 | 0 | 1 |
| UGC0012139 | Clust-UGC0002236 | 4954 | 15 | 1 | 0 | 1 |
| UGC0011652 | Clust-UGC0002236 | 7090 | 10 | 1 | 0 | 1 |
| UGC0002236 | Clust-UGC0002236 | 7634 | 19 | 1 | 0 | 1 |
| UGC0012412 | Clust-UGC0002236 | 6666 | 6 | 1 | 1 | 1 |
| UGC0002271 | Clust-UGC0002236 | 3519 | 6 | 1 | 0 | 1 |
| UGC0012385 | Clust-UGC0002236 | 11041 | 52 | 1 | 0 | 1 |
| UGC0003384 | Clust-UGC0002236 | 12278 | 48 | 1 | 0 | 1 |
| UGC0012168 | Clust-UGC0002236 | 3696 | 5 | 1 | 0 | 1 |
| UGC0005080 | Clust-UGC0002236 | 3422 | 4 | 1 | 0 | 1 |
| UGC0002254 | Clust-UGC0002236 | 11559 | 70 | 1 | 0 | 1 |
| UGC0011117 | Clust-UGC0002236 | 1678 | 2 | 1 | 0 | 0 |
| UGC0006415 | Clust-UGC0002236 | 3763 | 9 | 1 | 0 | 1 |
| UGC0002241 | Clust-UGC0002236 | 7556 | 16 | 1 | 0 | 1 |
| UGC0003246 | Clust-UGC0002236 | 10278 | 53 | 1 | 0 | 1 |
| UGC0012363 | Clust-UGC0002239 | 6090 | 7 | 1 | 0 | 1 |
| UGC0011658 | Clust-UGC0002239 | 487 | 1 | 0 | 0 | 0 |
| UGC0002239 | Clust-UGC0002239 | 10023 | 35 | 1 | 0 | 1 |
| UGC0011162 | Clust-UGC0002239 | 3648 | 2 | 1 | 0 | 1 |
| UGC0012053 | Clust-UGC0002239 | 3250 | 3 | 1 | 0 | 1 |
| UGC0012424 | Clust-UGC0002239 | 6559 | 13 | 1 | 0 | 1 |
| UGC0002244 | Clust-UGC0002244 | 14548 | 72 | 1 | 0 | 1 |
| UGC0007976 | Clust-UGC0002260 | 2356 | 3 | 1 | 0 | 1 |
| UGC0002260 | Clust-UGC0002260 | 5094 | 4 | 1 | 0 | 1 |
| UGC0003336 | Clust-UGC0002260 | 12341 | 57 | 1 | 0 | 1 |
| UGC0012023 | Clust-UGC0002272 | 2967 | 3 | 1 | 0 | 1 |
| UGC0002272 | Clust-UGC0002272 | 16754 | 129 | 1 | 0 | 1 |
| UGC0006994 | Clust-UGC0002272 | 7299 | 20 | 1 | 0 | 1 |
| UGC0010953 | Clust-UGC0002272 | 9649 | 36 | 1 | 0 | 1 |
| UGC0002282 | Clust-UGC0002282 | 825 | 1 | 0 | 0 | 0 |
| UGC0005370 | Clust-UGC0002283 | 1454 | 1 | 0 | 0 | 0 |
| UGC0007263 | Clust-UGC0002283 | 4574 | 4 | 1 | 0 | 1 |
| UGC0007909 | Clust-UGC0002283 | 3169 | 4 | 1 | 0 | 1 |
| UGC0011152 | Clust-UGC0002283 | 1603 | 2 | 1 | 0 | 0 |

| | | | | | | |
|---|---|---|---|---|---|---|
| UGC0007673 | Clust-UGC0002283 | 3675 | 5 | 1 | 0 | 1 |
| UGC0006336 | Clust-UGC0002283 | 1362 | 2 | 1 | 0 | 1 |
| UGC0007242 | Clust-UGC0002283 | 1739 | 1 | 0 | 0 | 0 |
| UGC0002998 | Clust-UGC0002283 | 2135 | 2 | 1 | 0 | 1 |
| UGC0004122 | Clust-UGC0002283 | 504 | 1 | 0 | 0 | 0 |
| UGC0012153 | Clust-UGC0002283 | 1577 | 1 | 0 | 0 | 0 |
| UGC0008606 | Clust-UGC0002283 | 1314 | 2 | 1 | 0 | 0 |
| UGC0008421 | Clust-UGC0002283 | 1284 | 1 | 0 | 0 | 0 |
| UGC0006773 | Clust-UGC0002283 | 4173 | 2 | 1 | 0 | 1 |
| UGC0005634 | Clust-UGC0002283 | 3494 | 3 | 1 | 0 | 1 |
| UGC0006755 | Clust-UGC0002283 | 4127 | 3 | 1 | 0 | 1 |
| UGC0009431 | Clust-UGC0002283 | 1207 | 1 | 0 | 0 | 0 |
| UGC0009430 | Clust-UGC0002283 | 1770 | 1 | 0 | 0 | 0 |
| UGC0007692 | Clust-UGC0002283 | 2888 | 3 | 1 | 0 | 1 |
| UGC0002503 | Clust-UGC0002283 | 721 | 2 | 1 | 0 | 1 |
| UGC0011822 | Clust-UGC0002283 | 3183 | 3 | 1 | 0 | 1 |
| UGC0007100 | Clust-UGC0002283 | 2338 | 3 | 1 | 0 | 1 |
| UGC0005715 | Clust-UGC0002283 | 6742 | 10 | 1 | 0 | 1 |
| UGC0003338 | Clust-UGC0002283 | 4367 | 4 | 1 | 0 | 1 |
| UGC0002345 | Clust-UGC0002283 | 1546 | 1 | 0 | 0 | 0 |
| UGC0010903 | Clust-UGC0002283 | 2831 | 5 | 1 | 0 | 1 |
| UGC0006832 | Clust-UGC0002283 | 4049 | 3 | 1 | 0 | 1 |
| UGC0006578 | Clust-UGC0002283 | 1858 | 2 | 1 | 0 | 1 |
| UGC0007508 | Clust-UGC0002283 | 4818 | 3 | 1 | 1 | 1 |
| UGC0006301 | Clust-UGC0002283 | 4773 | 7 | 1 | 1 | 1 |
| UGC0006987 | Clust-UGC0002283 | 1259 | 2 | 1 | 0 | 0 |
| UGC0012357 | Clust-UGC0002283 | 3074 | 3 | 1 | 0 | 1 |
| UGC0002349 | Clust-UGC0002283 | 1247 | 1 | 0 | 0 | 0 |
| UGC0002283 | Clust-UGC0002283 | 1077 | 1 | 0 | 0 | 0 |
| UGC0006722 | Clust-UGC0002283 | 2038 | 3 | 1 | 0 | 1 |
| UGC0005604 | Clust-UGC0002283 | 2880 | 5 | 1 | 0 | 1 |
| UGC0013004 | Clust-UGC0002283 | 2406 | 4 | 1 | 0 | 1 |
| UGC0004957 | Clust-UGC0002283 | 3600 | 6 | 1 | 0 | 1 |
| UGC0003335 | Clust-UGC0002283 | 2277 | 3 | 1 | 0 | 1 |
| UGC0007436 | Clust-UGC0002283 | 4680 | 6 | 1 | 0 | 1 |
| UGC0012394 | Clust-UGC0002283 | 2221 | 1 | 0 | 0 | 0 |
| UGC0007455 | Clust-UGC0002283 | 2623 | 3 | 1 | 0 | 1 |
| UGC0007134 | Clust-UGC0002283 | 4189 | 3 | 1 | 0 | 1 |
| UGC0004617 | Clust-UGC0002283 | 7073 | 12 | 1 | 0 | 1 |
| UGC0009159 | Clust-UGC0002283 | 1324 | 4 | 1 | 1 | 1 |
| UGC0006604 | Clust-UGC0002283 | 1637 | 2 | 1 | 0 | 1 |
| UGC0002950 | Clust-UGC0002283 | 786 | 1 | 0 | 0 | 0 |
| UGC0006470 | Clust-UGC0002283 | 631 | 1 | 0 | 0 | 0 |
| UGC0010935 | Clust-UGC0002283 | 1027 | 1 | 0 | 0 | 0 |
| UGC0008184 | Clust-UGC0002283 | 3298 | 3 | 1 | 0 | 1 |
| UGC0009586 | Clust-UGC0002283 | 4891 | 2 | 1 | 0 | 1 |
| UGC0007235 | Clust-UGC0002283 | 3585 | 4 | 1 | 0 | 1 |
| UGC0011520 | Clust-UGC0002283 | 1308 | 1 | 0 | 0 | 0 |
| UGC0008449 | Clust-UGC0002283 | 550 | 2 | 1 | 0 | 1 |
| UGC0012194 | Clust-UGC0002283 | 3225 | 5 | 1 | 0 | 1 |
| UGC0008353 | Clust-UGC0002283 | 578 | 1 | 0 | 0 | 0 |
| UGC0002475 | Clust-UGC0002283 | 75 | 1 | 0 | 0 | 0 |
| UGC0008867 | Clust-UGC0002287 | 2193 | 1 | 0 | 0 | 0 |
| UGC0012752 | Clust-UGC0002287 | 847 | 1 | 0 | 0 | 0 |
| UGC0006319 | Clust-UGC0002287 | 3706 | 4 | 1 | 0 | 1 |
| UGC0011920 | Clust-UGC0002287 | 3927 | 4 | 1 | 0 | 1 |
| UGC0011540 | Clust-UGC0002287 | 5083 | 5 | 1 | 0 | 1 |
| UGC0011489 | Clust-UGC0002287 | 4854 | 3 | 1 | 0 | 1 |
| UGC0007676 | Clust-UGC0002287 | 3289 | 3 | 1 | 0 | 1 |
| UGC0006334 | Clust-UGC0002287 | 244 | 1 | 0 | 0 | 0 |
| UGC0006332 | Clust-UGC0002287 | 3298 | 1 | 0 | 0 | 0 |
| UGC0011909 | Clust-UGC0002287 | 3757 | 1 | 0 | 0 | 0 |
| UGC0009528 | Clust-UGC0002287 | 1020 | 1 | 0 | 0 | 0 |
| UGC0011131 | Clust-UGC0002287 | 1487 | 2 | 1 | 0 | 0 |
| UGC0003763 | Clust-UGC0002287 | 457 | 1 | 0 | 0 | 0 |
| UGC0007004 | Clust-UGC0002287 | 3322 | 1 | 0 | 0 | 0 |
| UGC0006012 | Clust-UGC0002287 | 582 | 1 | 0 | 0 | 0 |
| UGC0012133 | Clust-UGC0002287 | 4733 | 6 | 1 | 1 | 1 |
| UGC0013241 | Clust-UGC0002287 | 2080 | 3 | 1 | 0 | 1 |
| UGC0012790 | Clust-UGC0002287 | 1165 | 1 | 0 | 0 | 0 |
| UGC0009028 | Clust-UGC0002287 | 250 | 1 | 0 | 0 | 0 |
| UGC0009023 | Clust-UGC0002287 | 2706 | 2 | 1 | 1 | 0 |
| UGC0006480 | Clust-UGC0002287 | 545 | 1 | 0 | 0 | 0 |
| UGC0006664 | Clust-UGC0002287 | 1816 | 2 | 1 | 0 | 1 |
| UGC0006667 | Clust-UGC0002287 | 1010 | 4 | 1 | 0 | 0 |
| UGC0005904 | Clust-UGC0002287 | 3565 | 4 | 1 | 0 | 1 |
| UGC0011888 | Clust-UGC0002287 | 13488 | 89 | 1 | 0 | 1 |
| UGC0013075 | Clust-UGC0002287 | 1103 | 1 | 0 | 0 | 0 |
| UGC0006935 | Clust-UGC0002287 | 1850 | 3 | 1 | 0 | 1 |

| | | | | | | |
|---|---|---|---|---|---|---|
| UGC0011408 | Clust-UGC0002287 | 1192 | 3 | 1 | 0 | 1 |
| UGC0009245 | Clust-UGC0002287 | 436 | 1 | 0 | 0 | 0 |
| UGC0013265 | Clust-UGC0002287 | 7964 | 18 | 1 | 1 | 1 |
| UGC0011140 | Clust-UGC0002287 | 2639 | 3 | 1 | 0 | 1 |
| UGC0007169 | Clust-UGC0002287 | 272 | 1 | 0 | 0 | 0 |
| UGC0007309 | Clust-UGC0002287 | 3483 | 1 | 0 | 0 | 0 |
| UGC0011421 | Clust-UGC0002287 | 2318 | 4 | 1 | 0 | 1 |
| UGC0010486 | Clust-UGC0002287 | 1870 | 3 | 1 | 1 | 1 |
| UGC0005777 | Clust-UGC0002287 | 4766 | 3 | 1 | 0 | 1 |
| UGC0006120 | Clust-UGC0002287 | 596 | 1 | 0 | 0 | 0 |
| UGC0007124 | Clust-UGC0002287 | 244 | 1 | 0 | 0 | 0 |
| UGC0012594 | Clust-UGC0002287 | 4449 | 4 | 1 | 0 | 1 |
| UGC0008061 | Clust-UGC0002287 | 1081 | 1 | 0 | 0 | 0 |
| UGC0003365 | Clust-UGC0002287 | 2410 | 1 | 0 | 0 | 0 |
| UGC0006851 | Clust-UGC0002287 | 1597 | 1 | 0 | 0 | 0 |
| UGC0005382 | Clust-UGC0002287 | 1539 | 2 | 1 | 0 | 0 |
| UGC0006874 | Clust-UGC0002287 | 1945 | 1 | 0 | 0 | 0 |
| UGC0011201 | Clust-UGC0002287 | 734 | 1 | 0 | 0 | 0 |
| UGC0007525 | Clust-UGC0002287 | 4783 | 3 | 1 | 0 | 1 |
| UGC0005846 | Clust-UGC0002287 | 2639 | 2 | 1 | 0 | 1 |
| UGC0010924 | Clust-UGC0002287 | 887 | 4 | 1 | 0 | 0 |
| UGC0011861 | Clust-UGC0002287 | 2472 | 1 | 0 | 0 | 0 |
| UGC0007507 | Clust-UGC0002287 | 5677 | 6 | 1 | 0 | 1 |
| UGC0010131 | Clust-UGC0002287 | 8038 | 19 | 1 | 1 | 1 |
| UGC0005827 | Clust-UGC0002287 | 5404 | 5 | 1 | 0 | 1 |
| UGC0013108 | Clust-UGC0002287 | 1490 | 1 | 0 | 0 | 0 |
| UGC0006046 | Clust-UGC0002287 | 467 | 1 | 0 | 0 | 0 |
| UGC0006833 | Clust-UGC0002287 | 3642 | 2 | 1 | 0 | 0 |
| UGC0009495 | Clust-UGC0002287 | 3005 | 2 | 1 | 0 | 1 |
| UGC0007098 | Clust-UGC0002287 | 453 | 1 | 0 | 0 | 0 |
| UGC0004154 | Clust-UGC0002287 | 3586 | 3 | 1 | 0 | 1 |
| UGC0004156 | Clust-UGC0002287 | 3091 | 4 | 1 | 0 | 1 |
| UGC0012431 | Clust-UGC0002287 | 2440 | 1 | 0 | 0 | 0 |
| UGC0010948 | Clust-UGC0002287 | 1689 | 2 | 1 | 0 | 0 |
| UGC0011161 | Clust-UGC0002287 | 1051 | 2 | 1 | 0 | 0 |
| UGC0011165 | Clust-UGC0002287 | 599 | 1 | 0 | 0 | 0 |
| UGC0006306 | Clust-UGC0002287 | 3448 | 2 | 1 | 0 | 1 |
| UGC0011575 | Clust-UGC0002287 | 2982 | 2 | 1 | 0 | 1 |
| UGC0011573 | Clust-UGC0002287 | 3382 | 2 | 1 | 0 | 1 |
| UGC0005945 | Clust-UGC0002287 | 2261 | 1 | 0 | 0 | 0 |
| UGC0007272 | Clust-UGC0002287 | 2155 | 1 | 0 | 0 | 0 |
| UGC0006703 | Clust-UGC0002287 | 687 | 1 | 0 | 0 | 0 |
| UGC0010985 | Clust-UGC0002287 | 3225 | 3 | 1 | 0 | 1 |
| UGC0002287 | Clust-UGC0002287 | 595 | 1 | 0 | 0 | 0 |
| UGC0011452 | Clust-UGC0002287 | 4756 | 2 | 1 | 0 | 1 |
| UGC0005887 | Clust-UGC0002287 | 1685 | 1 | 0 | 0 | 0 |
| UGC0012811 | Clust-UGC0002287 | 2477 | 3 | 1 | 0 | 1 |
| UGC0013029 | Clust-UGC0002287 | 4391 | 2 | 1 | 0 | 1 |
| UGC0011666 | Clust-UGC0002287 | 3865 | 5 | 1 | 0 | 1 |
| UGC0011474 | Clust-UGC0002287 | 4145 | 2 | 1 | 0 | 0 |
| UGC0011786 | Clust-UGC0002287 | 845 | 1 | 0 | 0 | 0 |
| UGC0013001 | Clust-UGC0002287 | 1816 | 2 | 1 | 0 | 1 |
| UGC0006718 | Clust-UGC0002287 | 159 | 1 | 0 | 0 | 0 |
| UGC0006744 | Clust-UGC0002287 | 136 | 1 | 0 | 0 | 0 |
| UGC0013255 | Clust-UGC0002287 | 468 | 1 | 0 | 0 | 0 |
| UGC0007377 | Clust-UGC0002287 | 171 | 1 | 0 | 0 | 0 |
| UGC0011416 | Clust-UGC0002287 | 1985 | 1 | 0 | 0 | 0 |
| UGC0011415 | Clust-UGC0002287 | 7414 | 19 | 1 | 0 | 1 |
| UGC0012854 | Clust-UGC0002287 | 5326 | 8 | 1 | 0 | 1 |
| UGC0002532 | Clust-UGC0002287 | 3675 | 3 | 1 | 0 | 1 |
| UGC0006902 | Clust-UGC0002287 | 1438 | 1 | 0 | 0 | 0 |
| UGC0007317 | Clust-UGC0002287 | 2199 | 2 | 1 | 0 | 1 |
| UGC0011431 | Clust-UGC0002287 | 1300 | 4 | 1 | 0 | 0 |
| UGC0006962 | Clust-UGC0002287 | 1033 | 1 | 0 | 0 | 0 |
| UGC0011602 | Clust-UGC0002287 | 2514 | 1 | 0 | 0 | 0 |
| UGC0012953 | Clust-UGC0002287 | 401 | 1 | 0 | 0 | 0 |
| UGC0007330 | Clust-UGC0002287 | 2179 | 2 | 1 | 0 | 1 |
| UGC0006412 | Clust-UGC0002287 | 4325 | 4 | 1 | 0 | 1 |
| UGC0012978 | Clust-UGC0002287 | 2743 | 2 | 1 | 0 | 1 |
| UGC0012231 | Clust-UGC0002287 | 4785 | 4 | 1 | 0 | 1 |
| UGC0008921 | Clust-UGC0002287 | 1930 | 2 | 1 | 0 | 1 |
| UGC0012915 | Clust-UGC0002287 | 2145 | 3 | 1 | 1 | 0 |
| UGC0007699 | Clust-UGC0002287 | 3579 | 4 | 1 | 0 | 1 |
| UGC0012763 | Clust-UGC0002287 | 1749 | 1 | 0 | 0 | 0 |
| UGC0010870 | Clust-UGC0002287 | 2209 | 1 | 0 | 0 | 0 |
| UGC0002511 | Clust-UGC0002287 | 1774 | 2 | 1 | 0 | 1 |
| UGC0012933 | Clust-UGC0002287 | 2680 | 3 | 1 | 1 | 1 |
| UGC0004626 | Clust-UGC0002287 | 9080 | 22 | 1 | 0 | 1 |
| UGC0009170 | Clust-UGC0002287 | 3409 | 2 | 1 | 0 | 1 |

| | | | | | | |
|---|---|---|---|---|---|---|
| UGC0012303 | Clust-UGC0002287 | 357 | 2 | 1 | 0 | 1 |
| UGC0012266 | Clust-UGC0002287 | 467 | 1 | 0 | 0 | 0 |
| UGC0011680 | Clust-UGC0002287 | 538 | 1 | 0 | 0 | 0 |
| UGC0002321 | Clust-UGC0002287 | 1555 | 2 | 1 | 0 | 1 |
| UGC0009376 | Clust-UGC0002287 | 2479 | 2 | 1 | 0 | 1 |
| UGC0013306 | Clust-UGC0002287 | 1995 | 2 | 1 | 0 | 0 |
| UGC0006866 | Clust-UGC0002287 | 1039 | 1 | 0 | 0 | 0 |
| UGC0006864 | Clust-UGC0002287 | 3284 | 2 | 1 | 0 | 1 |
| UGC0006396 | Clust-UGC0002287 | 5381 | 5 | 1 | 0 | 1 |
| UGC0002555 | Clust-UGC0002287 | 606 | 1 | 0 | 0 | 0 |
| UGC0005850 | Clust-UGC0002287 | 3979 | 3 | 1 | 0 | 1 |
| UGC0005858 | Clust-UGC0002287 | 4268 | 1 | 0 | 0 | 0 |
| UGC0012209 | Clust-UGC0002287 | 2820 | 2 | 1 | 0 | 0 |
| UGC0004667 | Clust-UGC0002287 | 5177 | 4 | 1 | 0 | 1 |
| UGC0003040 | Clust-UGC0002287 | 1272 | 1 | 0 | 0 | 0 |
| UGC0009670 | Clust-UGC0002287 | 1134 | 1 | 0 | 0 | 0 |
| UGC0012432 | Clust-UGC0002287 | 2724 | 3 | 1 | 0 | 1 |
| UGC0002296 | Clust-UGC0002296 | 4287 | 2 | 1 | 0 | 1 |
| UGC0012799 | Clust-UGC0002296 | 2550 | 1 | 0 | 0 | 0 |
| UGC0012435 | Clust-UGC0002296 | 446 | 1 | 0 | 0 | 0 |
| UGC0006060 | Clust-UGC0002296 | 4918 | 2 | 1 | 0 | 1 |
| UGC0007778 | Clust-UGC0002296 | 1525 | 1 | 0 | 0 | 0 |
| UGC0004979 | Clust-UGC0002296 | 5910 | 2 | 1 | 0 | 1 |
| UGC0011835 | Clust-UGC0002296 | 4727 | 4 | 1 | 0 | 1 |
| UGC0010162 | Clust-UGC0002300 | 10730 | 51 | 1 | 0 | 1 |
| UGC0010990 | Clust-UGC0002300 | 2489 | 2 | 1 | 0 | 0 |
| UGC0011130 | Clust-UGC0002300 | 1222 | 1 | 0 | 0 | 0 |
| UGC0005406 | Clust-UGC0002300 | 6009 | 9 | 1 | 0 | 1 |
| UGC0007343 | Clust-UGC0002300 | 5191 | 9 | 1 | 0 | 1 |
| UGC0006938 | Clust-UGC0002300 | 6271 | 11 | 1 | 0 | 1 |
| UGC0005633 | Clust-UGC0002300 | 8491 | 16 | 1 | 0 | 1 |
| UGC0005544 | Clust-UGC0002300 | 1949 | 1 | 0 | 0 | 0 |
| UGC0005562 | Clust-UGC0002300 | 2337 | 1 | 0 | 0 | 0 |
| UGC0007487 | Clust-UGC0002300 | 3107 | 4 | 1 | 0 | 1 |
| UGC0008389 | Clust-UGC0002300 | 1642 | 1 | 0 | 0 | 0 |
| UGC0004656 | Clust-UGC0002300 | 1613 | 1 | 0 | 0 | 0 |
| UGC0004654 | Clust-UGC0002300 | 11167 | 72 | 1 | 1 | 1 |
| UGC0007383 | Clust-UGC0002300 | 752 | 1 | 0 | 0 | 0 |
| UGC0011862 | Clust-UGC0002300 | 5369 | 3 | 1 | 0 | 1 |
| UGC0010134 | Clust-UGC0002300 | 106 | 1 | 0 | 0 | 0 |
| UGC0004844 | Clust-UGC0002300 | 6559 | 15 | 1 | 0 | 1 |
| UGC0002300 | Clust-UGC0002300 | 5873 | 7 | 1 | 0 | 1 |
| UGC0007910 | Clust-UGC0002300 | 6463 | 23 | 1 | 0 | 1 |
| UGC0007915 | Clust-UGC0002300 | 8578 | 62 | 1 | 0 | 1 |
| UGC0005410 | Clust-UGC0002300 | 2831 | 1 | 0 | 0 | 0 |
| UGC0005067 | Clust-UGC0002300 | 2057 | 1 | 0 | 0 | 0 |
| UGC0003333 | Clust-UGC0002300 | 2548 | 3 | 1 | 0 | 1 |
| UGC0005552 | Clust-UGC0002300 | 1545 | 1 | 0 | 0 | 0 |
| UGC0007153 | Clust-UGC0002300 | 4374 | 4 | 1 | 0 | 1 |
| UGC0005765 | Clust-UGC0002300 | 6709 | 11 | 1 | 0 | 1 |
| UGC0006624 | Clust-UGC0002300 | 2936 | 2 | 1 | 0 | 1 |
| UGC0004974 | Clust-UGC0002300 | 1174 | 1 | 0 | 0 | 0 |
| UGC0005539 | Clust-UGC0002300 | 1220 | 2 | 1 | 0 | 1 |
| UGC0004629 | Clust-UGC0002300 | 8524 | 27 | 1 | 0 | 1 |
| UGC0008967 | Clust-UGC0002300 | 4921 | 7 | 1 | 1 | 1 |
| UGC0006867 | Clust-UGC0002300 | 1032 | 1 | 0 | 0 | 0 |
| UGC0007674 | Clust-UGC0002303 | 9268 | 10 | 1 | 0 | 1 |
| UGC0011136 | Clust-UGC0002303 | 1021 | 2 | 1 | 0 | 0 |
| UGC0009470 | Clust-UGC0002303 | 6102 | 5 | 1 | 0 | 1 |
| UGC0006402 | Clust-UGC0002303 | 2015 | 2 | 1 | 0 | 0 |
| UGC0002303 | Clust-UGC0002303 | 9045 | 5 | 1 | 0 | 1 |
| UGC0005985 | Clust-UGC0002303 | 4444 | 2 | 1 | 0 | 0 |
| UGC0004526 | Clust-UGC0002303 | 1236 | 2 | 1 | 1 | 0 |
| UGC0012316 | Clust-UGC0002303 | 3494 | 3 | 1 | 1 | 0 |
| UGC0002304 | Clust-UGC0002304 | 6761 | 5 | 1 | 0 | 1 |
| UGC0002305 | Clust-UGC0002305 | 3963 | 2 | 1 | 0 | 1 |
| UGC0007926 | Clust-UGC0002306 | 4091 | 9 | 1 | 0 | 1 |
| UGC0010996 | Clust-UGC0002306 | 11350 | 72 | 1 | 0 | 1 |
| UGC0010615 | Clust-UGC0002306 | 2960 | 1 | 0 | 0 | 0 |
| UGC0008886 | Clust-UGC0002306 | 2572 | 3 | 1 | 0 | 1 |
| UGC0012960 | Clust-UGC0002306 | 3677 | 5 | 1 | 0 | 1 |
| UGC0007694 | Clust-UGC0002306 | 4134 | 6 | 1 | 0 | 1 |
| UGC0008590 | Clust-UGC0002306 | 1266 | 1 | 0 | 0 | 0 |
| UGC0006859 | Clust-UGC0002306 | 3538 | 4 | 1 | 0 | 1 |
| UGC0008410 | Clust-UGC0002306 | 1012 | 1 | 0 | 0 | 0 |
| UGC0011537 | Clust-UGC0002306 | 1039 | 2 | 1 | 1 | 0 |
| UGC0002306 | Clust-UGC0002306 | 3548 | 4 | 1 | 0 | 1 |
| UGC0009455 | Clust-UGC0002306 | 1331 | 1 | 0 | 0 | 0 |
| UGC0008739 | Clust-UGC0002306 | 944 | 1 | 0 | 0 | 0 |

| | | | | | | |
|---|---|---|---|---|---|---|
| UGC0013232 | Clust-UGC0002306 | 358 | 2 | 1 | 0 | 0 |
| UGC0008552 | Clust-UGC0002306 | 3286 | 4 | 1 | 0 | 1 |
| UGC0009013 | Clust-UGC0002306 | 3851 | 2 | 1 | 0 | 1 |
| UGC0006961 | Clust-UGC0002306 | 4853 | 7 | 1 | 0 | 1 |
| UGC0007780 | Clust-UGC0002306 | 3825 | 4 | 1 | 0 | 1 |
| UGC0011509 | Clust-UGC0002306 | 19016 | 185 | 1 | 0 | 1 |
| UGC0002999 | Clust-UGC0002307 | 1454 | 2 | 1 | 0 | 1 |
| UGC0005658 | Clust-UGC0002307 | 1954 | 1 | 0 | 0 | 0 |
| UGC0012253 | Clust-UGC0002307 | 6902 | 10 | 1 | 0 | 1 |
| UGC0002307 | Clust-UGC0002307 | 2985 | 1 | 0 | 0 | 0 |
| UGC0011492 | Clust-UGC0002307 | 8042 | 19 | 1 | 0 | 1 |
| UGC0005626 | Clust-UGC0002307 | 4644 | 10 | 1 | 0 | 0 |
| UGC0005781 | Clust-UGC0002310 | 7015 | 21 | 1 | 0 | 1 |
| UGC0005789 | Clust-UGC0002310 | 3615 | 3 | 1 | 0 | 1 |
| UGC0002310 | Clust-UGC0002310 | 1366 | 1 | 0 | 0 | 0 |
| UGC0002314 | Clust-UGC0002310 | 5852 | 7 | 1 | 0 | 1 |
| UGC0002319 | Clust-UGC0002310 | 5582 | 4 | 1 | 0 | 1 |
| UGC0006454 | Clust-UGC0002310 | 5335 | 4 | 1 | 0 | 1 |
| UGC0006163 | Clust-UGC0002310 | 1781 | 1 | 0 | 0 | 0 |
| UGC0005995 | Clust-UGC0002310 | 3707 | 4 | 1 | 0 | 1 |
| UGC0012170 | Clust-UGC0002310 | 7374 | 13 | 1 | 0 | 1 |
| UGC0010991 | Clust-UGC0002310 | 1247 | 2 | 1 | 0 | 0 |
| UGC0006059 | Clust-UGC0002310 | 709 | 1 | 0 | 0 | 0 |
| UGC0011157 | Clust-UGC0002310 | 1773 | 1 | 0 | 0 | 0 |
| UGC0011485 | Clust-UGC0002310 | 1512 | 2 | 1 | 0 | 0 |
| UGC0012932 | Clust-UGC0002310 | 8531 | 33 | 1 | 1 | 1 |
| UGC0007677 | Clust-UGC0002310 | 3992 | 2 | 1 | 0 | 1 |
| UGC0004127 | Clust-UGC0002310 | 418 | 1 | 0 | 0 | 0 |
| UGC0006296 | Clust-UGC0002310 | 2653 | 3 | 1 | 0 | 1 |
| UGC0006042 | Clust-UGC0002310 | 1117 | 1 | 0 | 0 | 0 |
| UGC0006123 | Clust-UGC0002310 | 1694 | 2 | 1 | 0 | 1 |
| UGC0013032 | Clust-UGC0002310 | 10049 | 38 | 1 | 1 | 1 |
| UGC0013033 | Clust-UGC0002310 | 5392 | 5 | 1 | 0 | 1 |
| UGC0013034 | Clust-UGC0002310 | 8282 | 25 | 1 | 0 | 1 |
| UGC0005898 | Clust-UGC0002310 | 4835 | 5 | 1 | 0 | 1 |
| UGC0006272 | Clust-UGC0002310 | 5760 | 6 | 1 | 0 | 1 |
| UGC0006447 | Clust-UGC0002310 | 6082 | 6 | 1 | 0 | 1 |
| UGC0006018 | Clust-UGC0002310 | 778 | 1 | 0 | 0 | 0 |
| UGC0010248 | Clust-UGC0002310 | 483 | 2 | 1 | 0 | 1 |
| UGC0009029 | Clust-UGC0002310 | 872 | 1 | 0 | 0 | 0 |
| UGC0012309 | Clust-UGC0002310 | 1735 | 1 | 0 | 0 | 0 |
| UGC0012305 | Clust-UGC0002310 | 1233 | 1 | 0 | 0 | 0 |
| UGC0006488 | Clust-UGC0002310 | 2658 | 2 | 1 | 0 | 1 |
| UGC0006481 | Clust-UGC0002310 | 768 | 1 | 0 | 0 | 0 |
| UGC0008572 | Clust-UGC0002310 | 2233 | 1 | 0 | 0 | 0 |
| UGC0008575 | Clust-UGC0002310 | 6172 | 10 | 1 | 0 | 1 |
| UGC0011657 | Clust-UGC0002310 | 288 | 1 | 0 | 0 | 0 |
| UGC0005902 | Clust-UGC0002310 | 2269 | 1 | 0 | 0 | 0 |
| UGC0005073 | Clust-UGC0002310 | 1409 | 1 | 0 | 0 | 0 |
| UGC0011881 | Clust-UGC0002310 | 1690 | 1 | 0 | 0 | 0 |
| UGC0007612 | Clust-UGC0002310 | 5108 | 6 | 1 | 0 | 1 |
| UGC0006652 | Clust-UGC0002310 | 3597 | 7 | 1 | 0 | 1 |
| UGC0011405 | Clust-UGC0002310 | 2796 | 2 | 1 | 0 | 1 |
| UGC0010759 | Clust-UGC0002310 | 826 | 3 | 1 | 0 | 0 |
| UGC0011985 | Clust-UGC0002310 | 3177 | 1 | 0 | 0 | 0 |
| UGC0009064 | Clust-UGC0002310 | 3080 | 7 | 1 | 0 | 1 |
| UGC0013264 | Clust-UGC0002310 | 903 | 2 | 1 | 0 | 0 |
| UGC0009473 | Clust-UGC0002310 | 2211 | 1 | 0 | 0 | 0 |
| UGC0006473 | Clust-UGC0002310 | 2398 | 2 | 1 | 0 | 1 |
| UGC0006479 | Clust-UGC0002310 | 2624 | 3 | 1 | 0 | 1 |
| UGC0012731 | Clust-UGC0002310 | 2184 | 2 | 1 | 0 | 1 |
| UGC0009049 | Clust-UGC0002310 | 494 | 1 | 0 | 0 | 0 |
| UGC0008525 | Clust-UGC0002310 | 1044 | 2 | 1 | 0 | 1 |
| UGC0008419 | Clust-UGC0002310 | 684 | 1 | 0 | 0 | 0 |
| UGC0012945 | Clust-UGC0002310 | 5005 | 4 | 1 | 0 | 1 |
| UGC0005013 | Clust-UGC0002310 | 12261 | 59 | 1 | 0 | 1 |
| UGC0005548 | Clust-UGC0002310 | 417 | 1 | 0 | 0 | 0 |
| UGC0010846 | Clust-UGC0002310 | 2395 | 2 | 1 | 0 | 0 |
| UGC0006631 | Clust-UGC0002310 | 846 | 1 | 0 | 0 | 0 |
| UGC0009202 | Clust-UGC0002310 | 5573 | 4 | 1 | 0 | 1 |
| UGC0012967 | Clust-UGC0002310 | 5405 | 7 | 1 | 1 | 1 |
| UGC0010527 | Clust-UGC0002310 | 168 | 1 | 0 | 0 | 0 |
| UGC0006324 | Clust-UGC0002310 | 7959 | 19 | 1 | 0 | 1 |
| UGC0010818 | Clust-UGC0002310 | 2564 | 9 | 1 | 0 | 1 |
| UGC0010880 | Clust-UGC0002310 | 1574 | 2 | 1 | 0 | 1 |
| UGC0011595 | Clust-UGC0002310 | 990 | 2 | 1 | 0 | 1 |
| UGC0006929 | Clust-UGC0002310 | 3830 | 6 | 1 | 0 | 1 |
| UGC0005861 | Clust-UGC0002310 | 4703 | 4 | 1 | 0 | 1 |
| UGC0009388 | Clust-UGC0002310 | 7783 | 20 | 1 | 0 | 1 |

| | | | | | | |
|---|---|---|---|---|---|---|
| UGC0009385 | Clust-UGC0002310 | 6916 | 10 | 1 | 0 | 1 |
| UGC0013339 | Clust-UGC0002310 | 2001 | 2 | 1 | 1 | 0 |
| UGC0006387 | Clust-UGC0002310 | 6470 | 11 | 1 | 0 | 1 |
| UGC0007381 | Clust-UGC0002310 | 637 | 1 | 0 | 0 | 0 |
| UGC0011209 | Clust-UGC0002310 | 6251 | 8 | 1 | 0 | 1 |
| UGC0009596 | Clust-UGC0002310 | 996 | 1 | 0 | 0 | 0 |
| UGC0005841 | Clust-UGC0002310 | 8296 | 22 | 1 | 0 | 1 |
| UGC0006788 | Clust-UGC0002310 | 4726 | 3 | 1 | 0 | 1 |
| UGC0009360 | Clust-UGC0002310 | 5628 | 6 | 1 | 0 | 1 |
| UGC0006109 | Clust-UGC0002310 | 2474 | 4 | 1 | 0 | 1 |
| UGC0006103 | Clust-UGC0002310 | 339 | 1 | 0 | 0 | 0 |
| UGC0008999 | Clust-UGC0002310 | 2855 | 2 | 1 | 0 | 0 |
| UGC0013100 | Clust-UGC0002310 | 6284 | 12 | 1 | 0 | 1 |
| UGC0012166 | Clust-UGC0002310 | 3752 | 4 | 1 | 0 | 1 |
| UGC0009344 | Clust-UGC0002310 | 4972 | 6 | 1 | 0 | 1 |
| UGC0009267 | Clust-UGC0002310 | 4053 | 8 | 1 | 0 | 1 |
| UGC0005554 | Clust-UGC0002310 | 1050 | 1 | 0 | 0 | 0 |
| UGC0012187 | Clust-UGC0002310 | 4778 | 7 | 1 | 0 | 1 |
| UGC0012188 | Clust-UGC0002310 | 1676 | 1 | 0 | 0 | 0 |
| UGC0009268 | Clust-UGC0002310 | 7030 | 13 | 1 | 0 | 1 |
| UGC0008226 | Clust-UGC0002310 | 1704 | 1 | 0 | 0 | 0 |
| UGC0010583 | Clust-UGC0002310 | 2167 | 2 | 1 | 0 | 1 |
| UGC0010911 | Clust-UGC0002310 | 255 | 1 | 0 | 0 | 0 |
| UGC0006364 | Clust-UGC0002310 | 1943 | 2 | 1 | 0 | 1 |
| UGC0006149 | Clust-UGC0002310 | 434 | 1 | 0 | 0 | 0 |
| UGC0007541 | Clust-UGC0002310 | 7102 | 6 | 1 | 0 | 1 |
| UGC0011518 | Clust-UGC0002310 | 7397 | 13 | 1 | 0 | 1 |
| UGC0011511 | Clust-UGC0002310 | 3053 | 4 | 1 | 0 | 1 |
| UGC0004153 | Clust-UGC0002310 | 6143 | 9 | 1 | 0 | 1 |
| UGC0005796 | Clust-UGC0002310 | 7450 | 20 | 1 | 0 | 1 |
| UGC0011167 | Clust-UGC0002310 | 1089 | 1 | 0 | 0 | 0 |
| UGC0002354 | Clust-UGC0002310 | 2341 | 1 | 0 | 0 | 0 |
| UGC0009308 | Clust-UGC0002310 | 1752 | 3 | 1 | 0 | 1 |
| UGC0006304 | Clust-UGC0002310 | 4404 | 2 | 1 | 0 | 1 |
| UGC0011577 | Clust-UGC0002310 | 620 | 2 | 1 | 0 | 0 |
| UGC0011572 | Clust-UGC0002310 | 1425 | 1 | 0 | 0 | 0 |
| UGC0010628 | Clust-UGC0002310 | 1479 | 1 | 0 | 0 | 0 |
| UGC0006017 | Clust-UGC0002310 | 2354 | 1 | 0 | 0 | 0 |
| UGC0009516 | Clust-UGC0002310 | 300 | 1 | 0 | 0 | 0 |
| UGC0002322 | Clust-UGC0002310 | 7260 | 7 | 1 | 0 | 1 |
| UGC0012144 | Clust-UGC0002310 | 5093 | 6 | 1 | 0 | 1 |
| UGC0012148 | Clust-UGC0002310 | 2046 | 1 | 0 | 0 | 0 |
| UGC0012116 | Clust-UGC0002310 | 1922 | 4 | 1 | 0 | 1 |
| UGC0005944 | Clust-UGC0002310 | 4076 | 2 | 1 | 0 | 1 |
| UGC0009727 | Clust-UGC0002310 | 1263 | 1 | 0 | 0 | 0 |
| UGC0011933 | Clust-UGC0002310 | 4643 | 6 | 1 | 0 | 1 |
| UGC0005828 | Clust-UGC0002310 | 2875 | 1 | 0 | 0 | 0 |
| UGC0005017 | Clust-UGC0002310 | 3748 | 2 | 1 | 0 | 1 |
| UGC0010989 | Clust-UGC0002310 | 4099 | 4 | 1 | 0 | 1 |
| UGC0006048 | Clust-UGC0002310 | 1289 | 1 | 0 | 0 | 0 |
| UGC0006045 | Clust-UGC0002310 | 655 | 1 | 0 | 0 | 0 |
| UGC0012296 | Clust-UGC0002310 | 5514 | 8 | 1 | 0 | 1 |
| UGC0003777 | Clust-UGC0002310 | 617 | 1 | 0 | 0 | 0 |
| UGC0011919 | Clust-UGC0002310 | 2175 | 2 | 1 | 0 | 1 |
| UGC0011918 | Clust-UGC0002310 | 6440 | 9 | 1 | 0 | 1 |
| UGC0011913 | Clust-UGC0002310 | 1704 | 1 | 0 | 0 | 0 |
| UGC0006416 | Clust-UGC0002310 | 6404 | 6 | 1 | 0 | 1 |
| UGC0004130 | Clust-UGC0002310 | 616 | 2 | 1 | 0 | 1 |
| UGC0006281 | Clust-UGC0002310 | 2990 | 5 | 1 | 0 | 1 |
| UGC0007626 | Clust-UGC0002310 | 8730 | 23 | 1 | 0 | 1 |
| UGC0007624 | Clust-UGC0002310 | 6554 | 7 | 1 | 0 | 1 |
| UGC0009017 | Clust-UGC0002310 | 2032 | 3 | 1 | 1 | 1 |
| UGC0012311 | Clust-UGC0002310 | 2559 | 3 | 1 | 0 | 1 |
| UGC0012317 | Clust-UGC0002310 | 2016 | 2 | 1 | 0 | 1 |
| UGC0006721 | Clust-UGC0002310 | 327 | 1 | 0 | 0 | 0 |
| UGC0011665 | Clust-UGC0002310 | 1513 | 1 | 0 | 0 | 0 |
| UGC0006006 | Clust-UGC0002310 | 3169 | 2 | 1 | 0 | 1 |
| UGC0007609 | Clust-UGC0002310 | 6267 | 8 | 1 | 0 | 1 |
| UGC0010259 | Clust-UGC0002310 | 480 | 2 | 1 | 0 | 1 |
| UGC0013008 | Clust-UGC0002310 | 2068 | 4 | 1 | 1 | 1 |
| UGC0006643 | Clust-UGC0002310 | 143 | 1 | 0 | 0 | 0 |
| UGC0006023 | Clust-UGC0002310 | 2090 | 1 | 0 | 0 | 0 |
| UGC0007379 | Clust-UGC0002310 | 3552 | 3 | 1 | 0 | 1 |
| UGC0010230 | Clust-UGC0002310 | 311 | 2 | 1 | 0 | 1 |
| UGC0009014 | Clust-UGC0002310 | 1939 | 2 | 1 | 0 | 1 |
| UGC0013361 | Clust-UGC0002310 | 7186 | 15 | 1 | 1 | 1 |
| UGC0011621 | Clust-UGC0002310 | 1916 | 3 | 1 | 0 | 0 |
| UGC0007412 | Clust-UGC0002310 | 5188 | 4 | 1 | 0 | 1 |
| UGC0006507 | Clust-UGC0002310 | 85 | 1 | 0 | 0 | 0 |

| | | | | | | |
|---|---|---|---|---|---|---|
| UGC0011483 | Clust-UGC0002310 | 7692 | 13 | 1 | 0 | 1 |
| UGC0005026 | Clust-UGC0002310 | 5245 | 5 | 1 | 0 | 1 |
| UGC0009463 | Clust-UGC0002310 | 3344 | 3 | 1 | 0 | 1 |
| UGC0009467 | Clust-UGC0002310 | 1280 | 3 | 1 | 0 | 1 |
| UGC0006598 | Clust-UGC0002310 | 1962 | 2 | 1 | 0 | 1 |
| UGC0010819 | Clust-UGC0002310 | 1768 | 4 | 1 | 0 | 0 |
| UGC0010215 | Clust-UGC0002310 | 589 | 1 | 0 | 0 | 0 |
| UGC0004524 | Clust-UGC0002310 | 9693 | 17 | 1 | 0 | 1 |
| UGC0006964 | Clust-UGC0002310 | 6004 | 10 | 1 | 0 | 1 |
| UGC0011605 | Clust-UGC0002310 | 3803 | 2 | 1 | 0 | 0 |
| UGC0011600 | Clust-UGC0002310 | 1829 | 1 | 0 | 0 | 0 |
| UGC0012951 | Clust-UGC0002310 | 6720 | 11 | 1 | 1 | 1 |
| UGC0010004 | Clust-UGC0002310 | 1966 | 5 | 1 | 0 | 1 |
| UGC0005335 | Clust-UGC0002310 | 3266 | 3 | 1 | 0 | 1 |
| UGC0005551 | Clust-UGC0002310 | 510 | 1 | 0 | 0 | 0 |
| UGC0009446 | Clust-UGC0002310 | 2091 | 1 | 0 | 0 | 0 |
| UGC0006944 | Clust-UGC0002310 | 6294 | 11 | 1 | 0 | 1 |
| UGC0003414 | Clust-UGC0002310 | 694 | 1 | 0 | 0 | 0 |
| UGC0010240 | Clust-UGC0002310 | 223 | 1 | 0 | 0 | 0 |
| UGC0010799 | Clust-UGC0002310 | 2975 | 8 | 1 | 0 | 1 |
| UGC0011811 | Clust-UGC0002310 | 1887 | 5 | 1 | 0 | 1 |
| UGC0006089 | Clust-UGC0002310 | 301 | 1 | 0 | 0 | 0 |
| UGC0006086 | Clust-UGC0002310 | 2947 | 2 | 1 | 0 | 1 |
| UGC0004568 | Clust-UGC0002310 | 12144 | 54 | 1 | 0 | 1 |
| UGC0004973 | Clust-UGC0002310 | 3708 | 8 | 1 | 1 | 1 |
| UGC0004609 | Clust-UGC0002310 | 9434 | 21 | 1 | 0 | 1 |
| UGC0004603 | Clust-UGC0002310 | 7046 | 10 | 1 | 0 | 1 |
| UGC0012769 | Clust-UGC0002310 | 1172 | 1 | 0 | 0 | 0 |
| UGC0012764 | Clust-UGC0002310 | 1475 | 2 | 1 | 0 | 1 |
| UGC0005603 | Clust-UGC0002310 | 2404 | 2 | 1 | 1 | 0 |
| UGC0006607 | Clust-UGC0002310 | 1190 | 1 | 0 | 0 | 0 |
| UGC0006609 | Clust-UGC0002310 | 483 | 1 | 0 | 0 | 0 |
| UGC0009090 | Clust-UGC0002310 | 598 | 1 | 0 | 0 | 0 |
| UGC0006453 | Clust-UGC0002310 | 1713 | 1 | 0 | 0 | 0 |
| UGC0003398 | Clust-UGC0002310 | 3300 | 2 | 1 | 0 | 1 |
| UGC0011857 | Clust-UGC0002310 | 585 | 1 | 0 | 0 | 0 |
| UGC0009172 | Clust-UGC0002310 | 1799 | 1 | 0 | 0 | 0 |
| UGC0012745 | Clust-UGC0002310 | 1299 | 1 | 0 | 0 | 0 |
| UGC0012744 | Clust-UGC0002310 | 1002 | 1 | 0 | 0 | 0 |
| UGC0012742 | Clust-UGC0002310 | 1419 | 1 | 0 | 0 | 0 |
| UGC0012301 | Clust-UGC0002310 | 1254 | 1 | 0 | 0 | 0 |
| UGC0007390 | Clust-UGC0002310 | 2465 | 3 | 1 | 0 | 1 |
| UGC0007702 | Clust-UGC0002310 | 6223 | 9 | 1 | 0 | 1 |
| UGC0010937 | Clust-UGC0002310 | 2963 | 1 | 0 | 0 | 0 |
| UGC0004640 | Clust-UGC0002310 | 7318 | 12 | 1 | 0 | 1 |
| UGC0011876 | Clust-UGC0002310 | 1196 | 1 | 0 | 0 | 0 |
| UGC0009371 | Clust-UGC0002310 | 2633 | 3 | 1 | 0 | 1 |
| UGC0006393 | Clust-UGC0002310 | 1396 | 1 | 0 | 0 | 0 |
| UGC0006132 | Clust-UGC0002310 | 1300 | 2 | 1 | 0 | 0 |
| UGC0007519 | Clust-UGC0002310 | 6698 | 11 | 1 | 0 | 1 |
| UGC0004098 | Clust-UGC0002310 | 774 | 1 | 0 | 0 | 0 |
| UGC0009353 | Clust-UGC0002310 | 6479 | 14 | 1 | 0 | 1 |
| UGC0011524 | Clust-UGC0002310 | 3646 | 2 | 1 | 0 | 1 |
| UGC0011529 | Clust-UGC0002310 | 13877 | 93 | 1 | 0 | 1 |
| UGC0007579 | Clust-UGC0002310 | 6896 | 4 | 1 | 0 | 1 |
| UGC0009657 | Clust-UGC0002310 | 1746 | 1 | 0 | 0 | 0 |
| UGC0005837 | Clust-UGC0002310 | 5290 | 3 | 1 | 0 | 1 |
| UGC0012199 | Clust-UGC0002310 | 4311 | 2 | 1 | 0 | 1 |
| UGC0010970 | Clust-UGC0002310 | 6142 | 6 | 1 | 0 | 1 |
| UGC0003735 | Clust-UGC0002310 | 545 | 1 | 0 | 0 | 0 |
| UGC0012225 | Clust-UGC0002310 | 2683 | 3 | 1 | 0 | 1 |
| UGC0012991 | Clust-UGC0002310 | 4394 | 5 | 1 | 0 | 1 |
| UGC0006039 | Clust-UGC0002310 | 3648 | 3 | 1 | 0 | 1 |
| UGC0011964 | Clust-UGC0002310 | 4510 | 5 | 1 | 0 | 1 |
| UGC0006170 | Clust-UGC0002310 | 2178 | 3 | 1 | 0 | 1 |
| UGC0011501 | Clust-UGC0002310 | 740 | 1 | 0 | 0 | 0 |
| UGC0007089 | Clust-UGC0002310 | 3374 | 12 | 1 | 0 | 1 |
| UGC0007223 | Clust-UGC0002310 | 4043 | 4 | 1 | 0 | 1 |
| UGC0005813 | Clust-UGC0002310 | 3236 | 1 | 0 | 0 | 0 |
| UGC0012425 | Clust-UGC0002310 | 3304 | 3 | 1 | 0 | 1 |
| UGC0006542 | Clust-UGC0002310 | 2928 | 4 | 1 | 0 | 1 |
| UGC0011198 | Clust-UGC0002310 | 7196 | 18 | 1 | 0 | 1 |
| UGC0002313 | Clust-UGC0002313 | 4159 | 3 | 1 | 0 | 0 |
| UGC0012132 | Clust-UGC0002313 | 1704 | 2 | 1 | 0 | 1 |
| UGC0011446 | Clust-UGC0002313 | 4041 | 4 | 1 | 0 | 0 |
| UGC0007498 | Clust-UGC0002313 | 10264 | 9 | 1 | 0 | 1 |
| UGC0009001 | Clust-UGC0002313 | 140 | 1 | 0 | 0 | 0 |
| UGC0010201 | Clust-UGC0002313 | 505 | 1 | 0 | 0 | 0 |
| UGC0006972 | Clust-UGC0002313 | 1795 | 3 | 1 | 0 | 1 |

| | | | | | | |
|---|---|---|---|---|---|---|
| UGC0012575 | Clust-UGC0002313 | 3294 | 2 | 1 | 0 | 0 |
| UGC0006637 | Clust-UGC0002313 | 1236 | 1 | 0 | 0 | 0 |
| UGC0010862 | Clust-UGC0002313 | 2543 | 2 | 1 | 1 | 0 |
| UGC0005712 | Clust-UGC0002313 | 3879 | 1 | 0 | 0 | 0 |
| UGC0012959 | Clust-UGC0002313 | 5797 | 4 | 1 | 1 | 1 |
| UGC0005803 | Clust-UGC0002313 | 4627 | 1 | 0 | 0 | 0 |
| UGC0005770 | Clust-UGC0002313 | 7245 | 4 | 1 | 0 | 1 |
| UGC0003550 | Clust-UGC0002313 | 10954 | 7 | 1 | 0 | 1 |
| UGC0005972 | Clust-UGC0002313 | 208 | 1 | 0 | 0 | 0 |
| UGC0007076 | Clust-UGC0002313 | 3641 | 3 | 1 | 0 | 1 |
| UGC0010089 | Clust-UGC0002313 | 2644 | 1 | 0 | 0 | 0 |
| UGC0006280 | Clust-UGC0002313 | 2598 | 2 | 1 | 0 | 0 |
| UGC0003713 | Clust-UGC0002313 | 267 | 1 | 0 | 0 | 0 |
| UGC0007450 | Clust-UGC0002313 | 1093 | 1 | 0 | 0 | 0 |
| UGC0004953 | Clust-UGC0002313 | 1912 | 1 | 0 | 0 | 0 |
| UGC0005577 | Clust-UGC0002313 | 3301 | 3 | 1 | 0 | 0 |
| UGC0006885 | Clust-UGC0002313 | 2359 | 2 | 1 | 0 | 1 |
| UGC0005683 | Clust-UGC0002313 | 89 | 1 | 0 | 0 | 0 |
| UGC0009615 | Clust-UGC0002313 | 828 | 2 | 1 | 0 | 0 |
| UGC0004648 | Clust-UGC0002313 | 8724 | 2 | 1 | 0 | 1 |
| UGC0006117 | Clust-UGC0002313 | 879 | 1 | 0 | 0 | 0 |
| UGC0010344 | Clust-UGC0002313 | 124 | 1 | 0 | 0 | 0 |
| UGC0007227 | Clust-UGC0002313 | 6048 | 5 | 1 | 0 | 1 |
| UGC0003781 | Clust-UGC0002313 | 244 | 1 | 0 | 0 | 0 |
| UGC0002317 | Clust-UGC0002317 | 5061 | 2 | 1 | 0 | 1 |
| UGC0007162 | Clust-UGC0002317 | 3233 | 10 | 1 | 0 | 1 |
| UGC0012234 | Clust-UGC0002317 | 7164 | 16 | 1 | 0 | 1 |
| UGC0007057 | Clust-UGC0002317 | 2546 | 1 | 0 | 0 | 0 |
| UGC0011649 | Clust-UGC0002317 | 1845 | 2 | 1 | 0 | 0 |
| UGC0002977 | Clust-UGC0002317 | 2581 | 1 | 0 | 0 | 0 |
| UGC0006553 | Clust-UGC0002318 | 2014 | 1 | 0 | 0 | 0 |
| UGC0002318 | Clust-UGC0002318 | 4423 | 1 | 0 | 0 | 0 |
| UGC0006674 | Clust-UGC0002318 | 6924 | 10 | 1 | 0 | 1 |
| UGC0005919 | Clust-UGC0002318 | 4664 | 4 | 1 | 0 | 1 |
| UGC0004836 | Clust-UGC0002318 | 8650 | 16 | 1 | 0 | 1 |
| UGC0005952 | Clust-UGC0002318 | 1292 | 2 | 1 | 0 | 1 |
| UGC0006442 | Clust-UGC0002318 | 8752 | 12 | 1 | 0 | 1 |
| UGC0006032 | Clust-UGC0002318 | 2548 | 1 | 0 | 0 | 0 |
| UGC0006587 | Clust-UGC0002318 | 251 | 1 | 0 | 0 | 0 |
| UGC0007342 | Clust-UGC0002318 | 5361 | 6 | 1 | 0 | 1 |
| UGC0005913 | Clust-UGC0002318 | 1667 | 2 | 1 | 0 | 1 |
| UGC0009000 | Clust-UGC0002318 | 5501 | 2 | 1 | 0 | 1 |
| UGC0005931 | Clust-UGC0002318 | 5601 | 3 | 1 | 0 | 1 |
| UGC0005939 | Clust-UGC0002318 | 3748 | 2 | 1 | 0 | 1 |
| UGC0006099 | Clust-UGC0002318 | 1859 | 1 | 0 | 0 | 0 |
| UGC0005970 | Clust-UGC0002318 | 4480 | 3 | 1 | 0 | 1 |
| UGC0004610 | Clust-UGC0002318 | 3856 | 3 | 1 | 0 | 1 |
| UGC0013376 | Clust-UGC0002318 | 1520 | 2 | 1 | 0 | 0 |
| UGC0012775 | Clust-UGC0002318 | 2707 | 1 | 0 | 0 | 0 |
| UGC0003572 | Clust-UGC0002318 | 8830 | 12 | 1 | 0 | 1 |
| UGC0012028 | Clust-UGC0002318 | 1404 | 1 | 0 | 0 | 0 |
| UGC0010315 | Clust-UGC0002318 | 2475 | 1 | 0 | 0 | 0 |
| UGC0007716 | Clust-UGC0002318 | 7272 | 13 | 1 | 0 | 1 |
| UGC0005922 | Clust-UGC0002318 | 4168 | 2 | 1 | 0 | 1 |
| UGC0008414 | Clust-UGC0002318 | 2320 | 1 | 0 | 0 | 0 |
| UGC0008621 | Clust-UGC0002318 | 5142 | 2 | 1 | 0 | 1 |
| UGC0002489 | Clust-UGC0002318 | 120 | 1 | 0 | 0 | 0 |
| UGC0012210 | Clust-UGC0002318 | 9272 | 17 | 1 | 0 | 1 |
| UGC0010118 | Clust-UGC0002318 | 14458 | 93 | 1 | 0 | 1 |
| UGC0010172 | Clust-UGC0002318 | 998 | 1 | 0 | 0 | 0 |
| UGC0009689 | Clust-UGC0002318 | 397 | 1 | 0 | 0 | 0 |
| UGC0006300 | Clust-UGC0002318 | 7995 | 13 | 1 | 0 | 1 |
| UGC0008495 | Clust-UGC0002318 | 1050 | 1 | 0 | 0 | 0 |
| UGC0009188 | Clust-UGC0002318 | 1944 | 2 | 1 | 0 | 1 |
| UGC0004581 | Clust-UGC0002318 | 6249 | 6 | 1 | 0 | 1 |
| UGC0009531 | Clust-UGC0002318 | 1204 | 1 | 0 | 0 | 0 |
| UGC0005947 | Clust-UGC0002318 | 1248 | 2 | 1 | 0 | 1 |
| UGC0002363 | Clust-UGC0002318 | 433 | 1 | 0 | 0 | 0 |
| UGC0011455 | Clust-UGC0002318 | 2554 | 1 | 0 | 0 | 0 |
| UGC0013231 | Clust-UGC0002318 | 924 | 1 | 0 | 0 | 0 |
| UGC0009572 | Clust-UGC0002318 | 5892 | 2 | 1 | 0 | 1 |
| UGC0009573 | Clust-UGC0002318 | 2864 | 1 | 0 | 0 | 0 |
| UGC0008716 | Clust-UGC0002318 | 252 | 1 | 0 | 0 | 0 |
| UGC0006401 | Clust-UGC0002318 | 7583 | 8 | 1 | 0 | 1 |
| UGC0007190 | Clust-UGC0002318 | 4702 | 5 | 1 | 0 | 1 |
| UGC0013064 | Clust-UGC0002318 | 2320 | 1 | 0 | 0 | 0 |
| UGC0010490 | Clust-UGC0002318 | 2110 | 1 | 0 | 0 | 0 |
| UGC0006882 | Clust-UGC0002318 | 6026 | 5 | 1 | 0 | 1 |
| UGC0004975 | Clust-UGC0002318 | 1612 | 1 | 0 | 0 | 0 |

| | | | | | | |
|---|---|---|---|---|---|---|
| UGC0007116 | Clust-UGC0002318 | 3787 | 1 | 0 | 0 | 0 |
| UGC0003587 | Clust-UGC0002318 | 11867 | 22 | 1 | 0 | 1 |
| UGC0012586 | Clust-UGC0002318 | 5919 | 7 | 1 | 0 | 1 |
| UGC0006605 | Clust-UGC0002318 | 1392 | 1 | 0 | 0 | 0 |
| UGC0003525 | Clust-UGC0002318 | 3135 | 1 | 0 | 0 | 0 |
| UGC0007535 | Clust-UGC0002318 | 8076 | 11 | 1 | 0 | 1 |
| UGC0006474 | Clust-UGC0002318 | 3403 | 1 | 0 | 0 | 0 |
| UGC0005879 | Clust-UGC0002318 | 2421 | 1 | 0 | 0 | 0 |
| UGC0004645 | Clust-UGC0002318 | 9298 | 21 | 1 | 1 | 1 |
| UGC0004647 | Clust-UGC0002318 | 6975 | 5 | 1 | 1 | 1 |
| UGC0012223 | Clust-UGC0002318 | 4382 | 1 | 0 | 0 | 0 |
| UGC0006827 | Clust-UGC0002318 | 3533 | 1 | 0 | 0 | 0 |
| UGC0008862 | Clust-UGC0002320 | 1019 | 2 | 1 | 0 | 0 |
| UGC0012175 | Clust-UGC0002320 | 5692 | 8 | 1 | 0 | 1 |
| UGC0011675 | Clust-UGC0002320 | 1590 | 2 | 1 | 0 | 0 |
| UGC0009757 | Clust-UGC0002320 | 3799 | 6 | 1 | 0 | 1 |
| UGC0006463 | Clust-UGC0002320 | 1278 | 2 | 1 | 0 | 0 |
| UGC0006815 | Clust-UGC0002320 | 5631 | 5 | 1 | 0 | 1 |
| UGC0008792 | Clust-UGC0002320 | 2003 | 2 | 1 | 0 | 0 |
| UGC0002320 | Clust-UGC0002320 | 5405 | 7 | 1 | 0 | 1 |
| UGC0007918 | Clust-UGC0002320 | 2375 | 5 | 1 | 0 | 1 |
| UGC0008896 | Clust-UGC0002320 | 470 | 1 | 0 | 0 | 0 |
| UGC0005788 | Clust-UGC0002323 | 5707 | 4 | 1 | 0 | 1 |
| UGC0011549 | Clust-UGC0002323 | 7715 | 21 | 1 | 0 | 1 |
| UGC0009667 | Clust-UGC0002323 | 552 | 1 | 0 | 0 | 0 |
| UGC0002323 | Clust-UGC0002323 | 1230 | 1 | 0 | 0 | 0 |
| UGC0013063 | Clust-UGC0002323 | 6665 | 15 | 1 | 0 | 1 |
| UGC0012233 | Clust-UGC0002323 | 4076 | 1 | 0 | 0 | 0 |
| UGC0002325 | Clust-UGC0002325 | 6560 | 6 | 1 | 1 | 1 |
| UGC0006593 | Clust-UGC0002325 | 621 | 1 | 0 | 0 | 0 |
| UGC0003553 | Clust-UGC0002327 | 6405 | 8 | 1 | 2 | 1 |
| UGC0002327 | Clust-UGC0002327 | 1515 | 1 | 0 | 0 | 0 |
| UGC0012832 | Clust-UGC0002327 | 781 | 2 | 1 | 0 | 0 |
| UGC0011172 | Clust-UGC0002330 | 1620 | 3 | 1 | 0 | 1 |
| UGC0002330 | Clust-UGC0002330 | 2599 | 3 | 1 | 1 | 1 |
| UGC0004126 | Clust-UGC0002330 | 1995 | 3 | 1 | 0 | 1 |
| UGC0012118 | Clust-UGC0002330 | 4958 | 5 | 1 | 0 | 1 |
| UGC0009547 | Clust-UGC0002330 | 5753 | 23 | 1 | 0 | 1 |
| UGC0009024 | Clust-UGC0002330 | 11846 | 69 | 1 | 0 | 1 |
| UGC0012300 | Clust-UGC0002330 | 2348 | 2 | 1 | 0 | 1 |
| UGC0007408 | Clust-UGC0002330 | 3728 | 3 | 1 | 0 | 1 |
| UGC0006655 | Clust-UGC0002330 | 10454 | 48 | 1 | 0 | 1 |
| UGC0009240 | Clust-UGC0002330 | 20959 | 192 | 1 | 0 | 1 |
| UGC0009472 | Clust-UGC0002330 | 8095 | 35 | 1 | 0 | 1 |
| UGC0012941 | Clust-UGC0002330 | 10331 | 57 | 1 | 0 | 1 |
| UGC0010014 | Clust-UGC0002330 | 5705 | 22 | 1 | 0 | 1 |
| UGC0009229 | Clust-UGC0002330 | 25812 | 265 | 1 | 0 | 1 |
| UGC0005739 | Clust-UGC0002330 | 6428 | 16 | 1 | 0 | 1 |
| UGC0009418 | Clust-UGC0002330 | 1564 | 3 | 1 | 0 | 1 |
| UGC0005643 | Clust-UGC0002330 | 1744 | 3 | 1 | 0 | 1 |
| UGC0004652 | Clust-UGC0002330 | 15153 | 121 | 1 | 0 | 1 |
| UGC0012277 | Clust-UGC0002330 | 2010 | 3 | 1 | 0 | 1 |
| UGC0009363 | Clust-UGC0002330 | 4529 | 19 | 1 | 0 | 1 |
| UGC0009106 | Clust-UGC0002330 | 1088 | 2 | 1 | 0 | 0 |
| UGC0013106 | Clust-UGC0002330 | 4241 | 7 | 1 | 0 | 1 |
| UGC0009320 | Clust-UGC0002330 | 9093 | 43 | 1 | 0 | 1 |
| UGC0009496 | Clust-UGC0002330 | 7584 | 39 | 1 | 0 | 1 |
| UGC0010151 | Clust-UGC0002330 | 5961 | 30 | 1 | 0 | 1 |
| UGC0008264 | Clust-UGC0002330 | 395 | 1 | 0 | 0 | 0 |
| UGC0006701 | Clust-UGC0002330 | 7168 | 12 | 1 | 0 | 1 |
| UGC0013183 | Clust-UGC0002330 | 5246 | 7 | 1 | 0 | 1 |
| UGC0005343 | Clust-UGC0002330 | 4375 | 10 | 1 | 0 | 1 |
| UGC0004135 | Clust-UGC0002330 | 3827 | 5 | 1 | 0 | 0 |
| UGC0012816 | Clust-UGC0002330 | 2799 | 3 | 1 | 1 | 0 |
| UGC0009767 | Clust-UGC0002330 | 9527 | 55 | 1 | 0 | 1 |
| UGC0006498 | Clust-UGC0002330 | 4019 | 9 | 1 | 0 | 1 |
| UGC0003317 | Clust-UGC0002330 | 10679 | 44 | 1 | 0 | 1 |
| UGC0005027 | Clust-UGC0002330 | 5673 | 18 | 1 | 0 | 1 |
| UGC0006782 | Clust-UGC0002330 | 7603 | 17 | 1 | 0 | 1 |
| UGC0009485 | Clust-UGC0002330 | 6232 | 26 | 1 | 1 | 1 |
| UGC0005598 | Clust-UGC0002330 | 3891 | 5 | 1 | 0 | 1 |
| UGC0012760 | Clust-UGC0002330 | 6228 | 17 | 1 | 0 | 1 |
| UGC0009238 | Clust-UGC0002330 | 17159 | 127 | 1 | 0 | 1 |
| UGC0013320 | Clust-UGC0002330 | 3583 | 3 | 1 | 0 | 1 |
| UGC0009116 | Clust-UGC0002330 | 3570 | 9 | 1 | 0 | 1 |
| UGC0009117 | Clust-UGC0002330 | 3627 | 11 | 1 | 0 | 1 |
| UGC0009588 | Clust-UGC0002330 | 6599 | 25 | 1 | 0 | 1 |
| UGC0009587 | Clust-UGC0002330 | 4755 | 16 | 1 | 0 | 1 |
| UGC0005859 | Clust-UGC0002330 | 5675 | 8 | 1 | 0 | 1 |

| | | | | | | |
|---|---|---|---|---|---|---|
| UGC0009335 | Clust-UGC0002330 | 10207 | 50 | 1 | 0 | 1 |
| UGC0006826 | Clust-UGC0002330 | 7796 | 29 | 1 | 0 | 1 |
| UGC0005810 | Clust-UGC0002330 | 7646 | 18 | 1 | 0 | 1 |
| UGC0012420 | Clust-UGC0002330 | 8318 | 31 | 1 | 0 | 1 |
| UGC0011947 | Clust-UGC0002330 | 3019 | 8 | 1 | 1 | 1 |
| UGC0005914 | Clust-UGC0002331 | 3284 | 2 | 1 | 0 | 1 |
| UGC0002331 | Clust-UGC0002331 | 1637 | 1 | 0 | 0 | 0 |
| UGC0004746 | Clust-UGC0002331 | 293 | 1 | 0 | 0 | 0 |
| UGC0004708 | Clust-UGC0002331 | 18037 | 116 | 1 | 0 | 1 |
| UGC0011632 | Clust-UGC0002331 | 2882 | 5 | 1 | 0 | 1 |
| UGC0006893 | Clust-UGC0002331 | 4152 | 2 | 1 | 0 | 1 |
| UGC0002529 | Clust-UGC0002331 | 2728 | 3 | 1 | 0 | 1 |
| UGC0007697 | Clust-UGC0002331 | 4386 | 4 | 1 | 0 | 1 |
| UGC0012251 | Clust-UGC0002331 | 4550 | 3 | 1 | 0 | 1 |
| UGC0012254 | Clust-UGC0002331 | 1179 | 1 | 0 | 0 | 0 |
| UGC0002487 | Clust-UGC0002331 | 770 | 1 | 0 | 0 | 0 |
| UGC0012214 | Clust-UGC0002331 | 6200 | 5 | 1 | 0 | 1 |
| UGC0002551 | Clust-UGC0002331 | 1343 | 1 | 0 | 0 | 0 |
| UGC0004117 | Clust-UGC0002331 | 5789 | 5 | 1 | 0 | 1 |
| UGC0002343 | Clust-UGC0002331 | 2578 | 2 | 1 | 0 | 1 |
| UGC0006514 | Clust-UGC0002331 | 2023 | 2 | 1 | 0 | 1 |
| UGC0007259 | Clust-UGC0002331 | 6039 | 12 | 1 | 0 | 1 |
| UGC0009550 | Clust-UGC0002331 | 202 | 1 | 0 | 0 | 0 |
| UGC0004738 | Clust-UGC0002331 | 1019 | 2 | 1 | 0 | 1 |
| UGC0009298 | Clust-UGC0002331 | 2147 | 2 | 1 | 0 | 1 |
| UGC0007439 | Clust-UGC0002331 | 4371 | 3 | 1 | 0 | 1 |
| UGC0012987 | Clust-UGC0002331 | 4185 | 11 | 1 | 0 | 1 |
| UGC0004605 | Clust-UGC0002331 | 2702 | 2 | 1 | 0 | 1 |
| UGC0003538 | Clust-UGC0002331 | 1835 | 1 | 0 | 0 | 0 |
| UGC0003586 | Clust-UGC0002331 | 574 | 1 | 0 | 0 | 0 |
| UGC0002510 | Clust-UGC0002331 | 3369 | 6 | 1 | 0 | 1 |
| UGC0007391 | Clust-UGC0002331 | 860 | 1 | 0 | 0 | 0 |
| UGC0013300 | Clust-UGC0002331 | 1421 | 2 | 1 | 1 | 0 |
| UGC0002558 | Clust-UGC0002331 | 142 | 1 | 0 | 0 | 0 |
| UGC0003292 | Clust-UGC0002331 | 4208 | 2 | 1 | 0 | 1 |
| UGC0009313 | Clust-UGC0002331 | 9114 | 31 | 1 | 0 | 1 |
| UGC0002333 | Clust-UGC0002333 | 1370 | 2 | 1 | 0 | 0 |
| UGC0009038 | Clust-UGC0002333 | 788 | 2 | 1 | 0 | 1 |
| UGC0012607 | Clust-UGC0002333 | 3878 | 3 | 1 | 0 | 1 |
| UGC0007705 | Clust-UGC0002333 | 5151 | 7 | 1 | 0 | 1 |
| UGC0002336 | Clust-UGC0002336 | 684 | 1 | 0 | 0 | 0 |
| UGC0009500 | Clust-UGC0002338 | 1501 | 2 | 1 | 0 | 1 |
| UGC0002339 | Clust-UGC0002338 | 2410 | 1 | 0 | 0 | 0 |
| UGC0002338 | Clust-UGC0002338 | 2514 | 1 | 0 | 0 | 0 |
| UGC0007245 | Clust-UGC0002338 | 7603 | 19 | 1 | 1 | 1 |
| UGC0004983 | Clust-UGC0002338 | 4078 | 2 | 1 | 0 | 1 |
| UGC0006594 | Clust-UGC0002338 | 624 | 1 | 0 | 0 | 0 |
| UGC0004558 | Clust-UGC0002338 | 8576 | 20 | 1 | 0 | 1 |
| UGC0007465 | Clust-UGC0002338 | 4326 | 2 | 1 | 0 | 1 |
| UGC0004962 | Clust-UGC0002338 | 7102 | 14 | 1 | 0 | 1 |
| UGC0010191 | Clust-UGC0002338 | 12333 | 36 | 1 | 0 | 1 |
| UGC0002965 | Clust-UGC0002338 | 3767 | 4 | 1 | 0 | 1 |
| UGC0004574 | Clust-UGC0002338 | 3036 | 2 | 1 | 0 | 1 |
| UGC0004572 | Clust-UGC0002338 | 8348 | 15 | 1 | 0 | 1 |
| UGC0005713 | Clust-UGC0002338 | 6770 | 15 | 1 | 0 | 1 |
| UGC0008627 | Clust-UGC0002338 | 4756 | 3 | 1 | 1 | 1 |
| UGC0007733 | Clust-UGC0002338 | 7061 | 16 | 1 | 0 | 1 |
| UGC0003012 | Clust-UGC0002338 | 234 | 2 | 1 | 0 | 1 |
| UGC0004678 | Clust-UGC0002338 | 13866 | 79 | 1 | 0 | 1 |
| UGC0007569 | Clust-UGC0002338 | 1569 | 1 | 0 | 0 | 0 |
| UGC0007952 | Clust-UGC0002338 | 5020 | 2 | 1 | 0 | 1 |
| UGC0010178 | Clust-UGC0002338 | 4535 | 2 | 1 | 0 | 1 |
| UGC0004824 | Clust-UGC0002338 | 9284 | 31 | 1 | 0 | 1 |
| UGC0004588 | Clust-UGC0002338 | 3497 | 1 | 0 | 0 | 0 |
| UGC0005368 | Clust-UGC0002338 | 6601 | 16 | 1 | 0 | 1 |
| UGC0007586 | Clust-UGC0002338 | 5276 | 1 | 0 | 0 | 0 |
| UGC0007276 | Clust-UGC0002338 | 5788 | 3 | 1 | 0 | 1 |
| UGC0006723 | Clust-UGC0002338 | 1564 | 3 | 1 | 0 | 0 |
| UGC0003529 | Clust-UGC0002338 | 4683 | 2 | 1 | 0 | 1 |
| UGC0005332 | Clust-UGC0002338 | 58 | 1 | 0 | 0 | 0 |
| UGC0007531 | Clust-UGC0002338 | 5111 | 4 | 1 | 0 | 1 |
| UGC0005705 | Clust-UGC0002338 | 7831 | 20 | 1 | 0 | 1 |
| UGC0011683 | Clust-UGC0002338 | 6060 | 6 | 1 | 1 | 1 |
| UGC0006564 | Clust-UGC0002338 | 685 | 2 | 1 | 0 | 1 |
| UGC0007226 | Clust-UGC0002338 | 8675 | 25 | 1 | 0 | 1 |
| UGC0004764 | Clust-UGC0002340 | 6031 | 6 | 1 | 0 | 1 |
| UGC0004104 | Clust-UGC0002340 | 353 | 1 | 0 | 0 | 0 |
| UGC0003723 | Clust-UGC0002340 | 98 | 1 | 0 | 0 | 0 |
| UGC0004740 | Clust-UGC0002340 | 1432 | 1 | 0 | 0 | 0 |

| | | | | | | |
|---|---|---|---|---|---|---|
| UGC0003762 | Clust-UGC0002340 | 2137 | 2 | 1 | 0 | 0 |
| UGC0002562 | Clust-UGC0002340 | 836 | 1 | 0 | 0 | 0 |
| UGC0006271 | Clust-UGC0002340 | 4962 | 10 | 1 | 0 | 1 |
| UGC0006445 | Clust-UGC0002340 | 9192 | 41 | 1 | 0 | 1 |
| UGC0008705 | Clust-UGC0002340 | 2658 | 1 | 0 | 0 | 0 |
| UGC0006015 | Clust-UGC0002340 | 3485 | 2 | 1 | 0 | 1 |
| UGC0007407 | Clust-UGC0002340 | 5778 | 13 | 1 | 0 | 1 |
| UGC0006765 | Clust-UGC0002340 | 5180 | 11 | 1 | 0 | 1 |
| UGC0003726 | Clust-UGC0002340 | 1261 | 1 | 0 | 0 | 0 |
| UGC0009061 | Clust-UGC0002340 | 5196 | 6 | 1 | 0 | 1 |
| UGC0012365 | Clust-UGC0002340 | 6090 | 16 | 1 | 0 | 1 |
| UGC0012733 | Clust-UGC0002340 | 1843 | 2 | 1 | 0 | 1 |
| UGC0005975 | Clust-UGC0002340 | 2056 | 1 | 0 | 0 | 0 |
| UGC0009995 | Clust-UGC0002340 | 1389 | 1 | 0 | 0 | 0 |
| UGC0009991 | Clust-UGC0002340 | 1331 | 1 | 0 | 0 | 0 |
| UGC0005565 | Clust-UGC0002340 | 231 | 1 | 0 | 0 | 0 |
| UGC0007274 | Clust-UGC0002340 | 3864 | 3 | 1 | 0 | 1 |
| UGC0011178 | Clust-UGC0002340 | 5629 | 6 | 1 | 0 | 1 |
| UGC0005849 | Clust-UGC0002340 | 9819 | 53 | 1 | 1 | 1 |
| UGC0003568 | Clust-UGC0002340 | 9376 | 47 | 1 | 0 | 1 |
| UGC0007750 | Clust-UGC0002340 | 7218 | 19 | 1 | 0 | 1 |
| UGC0010008 | Clust-UGC0002340 | 1845 | 1 | 0 | 0 | 0 |
| UGC0002340 | Clust-UGC0002340 | 4392 | 3 | 1 | 0 | 1 |
| UGC0011126 | Clust-UGC0002340 | 924 | 1 | 0 | 0 | 0 |
| UGC0012622 | Clust-UGC0002340 | 8597 | 32 | 1 | 0 | 1 |
| UGC0012783 | Clust-UGC0002340 | 2586 | 1 | 0 | 0 | 0 |
| UGC0005684 | Clust-UGC0002340 | 2674 | 1 | 0 | 0 | 0 |
| UGC0007318 | Clust-UGC0002340 | 2498 | 1 | 0 | 0 | 0 |
| UGC0005890 | Clust-UGC0002340 | 5229 | 6 | 1 | 0 | 1 |
| UGC0011013 | Clust-UGC0002340 | 2139 | 1 | 0 | 0 | 0 |
| UGC0006083 | Clust-UGC0002340 | 5798 | 5 | 1 | 0 | 1 |
| UGC0004567 | Clust-UGC0002340 | 3900 | 6 | 1 | 0 | 1 |
| UGC0012938 | Clust-UGC0002340 | 3516 | 4 | 1 | 1 | 1 |
| UGC0010890 | Clust-UGC0002340 | 2315 | 3 | 1 | 0 | 0 |
| UGC0009612 | Clust-UGC0002340 | 2834 | 3 | 1 | 0 | 1 |
| UGC0011873 | Clust-UGC0002340 | 5219 | 4 | 1 | 0 | 1 |
| UGC0012017 | Clust-UGC0002340 | 3479 | 1 | 0 | 0 | 0 |
| UGC0010322 | Clust-UGC0002340 | 1698 | 1 | 0 | 0 | 0 |
| UGC0004094 | Clust-UGC0002340 | 674 | 1 | 0 | 0 | 0 |
| UGC0008985 | Clust-UGC0002340 | 3544 | 2 | 1 | 0 | 1 |
| UGC0012445 | Clust-UGC0002340 | 2814 | 2 | 1 | 0 | 1 |
| UGC0010973 | Clust-UGC0002340 | 4844 | 2 | 1 | 0 | 0 |
| UGC0006829 | Clust-UGC0002340 | 1923 | 1 | 0 | 0 | 0 |
| UGC0011611 | Clust-UGC0002341 | 2360 | 3 | 1 | 0 | 0 |
| UGC0002341 | Clust-UGC0002341 | 1150 | 3 | 1 | 0 | 1 |
| UGC0006858 | Clust-UGC0002342 | 1399 | 1 | 0 | 0 | 0 |
| UGC0009501 | Clust-UGC0002342 | 2903 | 1 | 0 | 0 | 0 |
| UGC0012617 | Clust-UGC0002342 | 8070 | 7 | 1 | 0 | 1 |
| UGC0007650 | Clust-UGC0002342 | 6873 | 4 | 1 | 0 | 1 |
| UGC0007655 | Clust-UGC0002342 | 4003 | 1 | 0 | 0 | 0 |
| UGC0012636 | Clust-UGC0002342 | 364 | 2 | 1 | 0 | 0 |
| UGC0011118 | Clust-UGC0002342 | 2174 | 2 | 1 | 0 | 1 |
| UGC0012793 | Clust-UGC0002342 | 911 | 1 | 0 | 0 | 0 |
| UGC0008687 | Clust-UGC0002342 | 455 | 1 | 0 | 0 | 0 |
| UGC0012710 | Clust-UGC0002342 | 4706 | 2 | 1 | 0 | 0 |
| UGC0002525 | Clust-UGC0002342 | 5554 | 2 | 1 | 0 | 1 |
| UGC0005755 | Clust-UGC0002342 | 833 | 1 | 0 | 0 | 0 |
| UGC0006409 | Clust-UGC0002342 | 7750 | 4 | 1 | 0 | 1 |
| UGC0007481 | Clust-UGC0002342 | 2281 | 1 | 0 | 0 | 0 |
| UGC0007713 | Clust-UGC0002342 | 2527 | 4 | 1 | 0 | 1 |
| UGC0006999 | Clust-UGC0002342 | 2087 | 1 | 0 | 0 | 0 |
| UGC0002342 | Clust-UGC0002342 | 5484 | 3 | 1 | 1 | 1 |
| UGC0004582 | Clust-UGC0002342 | 5083 | 3 | 1 | 0 | 1 |
| UGC0002982 | Clust-UGC0002342 | 3718 | 2 | 1 | 0 | 1 |
| UGC0004116 | Clust-UGC0002342 | 555 | 1 | 0 | 0 | 0 |
| UGC0005014 | Clust-UGC0002342 | 6541 | 4 | 1 | 0 | 1 |
| UGC0005012 | Clust-UGC0002342 | 2257 | 1 | 0 | 0 | 0 |
| UGC0007073 | Clust-UGC0002342 | 3097 | 1 | 0 | 0 | 0 |
| UGC0007017 | Clust-UGC0002342 | 2682 | 2 | 1 | 0 | 1 |
| UGC0013230 | Clust-UGC0002342 | 633 | 1 | 0 | 0 | 0 |
| UGC0007416 | Clust-UGC0002342 | 2402 | 1 | 0 | 0 | 0 |
| UGC0007638 | Clust-UGC0002342 | 5812 | 2 | 1 | 0 | 1 |
| UGC0004560 | Clust-UGC0002342 | 9184 | 3 | 1 | 0 | 1 |
| UGC0007476 | Clust-UGC0002342 | 2854 | 1 | 0 | 0 | 0 |
| UGC0007113 | Clust-UGC0002342 | 1258 | 1 | 0 | 0 | 0 |
| UGC0008058 | Clust-UGC0002342 | 1757 | 1 | 0 | 0 | 0 |
| UGC0007492 | Clust-UGC0002342 | 3439 | 1 | 0 | 0 | 0 |
| UGC0005392 | Clust-UGC0002342 | 4114 | 3 | 1 | 0 | 1 |
| UGC0006648 | Clust-UGC0002342 | 1737 | 2 | 1 | 0 | 1 |

| | | | | | | |
|---|---|---|---|---|---|---|
| UGC0012015 | Clust-UGC0002342 | 744 | 2 | 1 | 0 | 1 |
| UGC0011521 | Clust-UGC0002342 | 7680 | 8 | 1 | 0 | 1 |
| UGC0006561 | Clust-UGC0002342 | 2769 | 6 | 1 | 0 | 1 |
| UGC0005336 | Clust-UGC0002342 | 3366 | 1 | 0 | 0 | 0 |
| UGC0007203 | Clust-UGC0002344 | 612 | 1 | 0 | 0 | 0 |
| UGC0009197 | Clust-UGC0002344 | 3830 | 2 | 1 | 0 | 1 |
| UGC0011543 | Clust-UGC0002344 | 2029 | 3 | 1 | 0 | 0 |
| UGC0007906 | Clust-UGC0002344 | 8744 | 2 | 1 | 0 | 1 |
| UGC0012284 | Clust-UGC0002344 | 867 | 1 | 0 | 0 | 0 |
| UGC0012283 | Clust-UGC0002344 | 352 | 1 | 0 | 0 | 0 |
| UGC0012828 | Clust-UGC0002344 | 4619 | 3 | 1 | 0 | 1 |
| UGC0008843 | Clust-UGC0002344 | 1653 | 1 | 0 | 0 | 0 |
| UGC0007241 | Clust-UGC0002344 | 3870 | 3 | 1 | 0 | 0 |
| UGC0006316 | Clust-UGC0002344 | 6280 | 1 | 0 | 0 | 0 |
| UGC0009525 | Clust-UGC0002344 | 2546 | 2 | 1 | 0 | 1 |
| UGC0006291 | Clust-UGC0002344 | 3472 | 2 | 1 | 0 | 1 |
| UGC0006297 | Clust-UGC0002344 | 3514 | 1 | 0 | 0 | 0 |
| UGC0012156 | Clust-UGC0002344 | 1244 | 3 | 1 | 1 | 0 |
| UGC0011671 | Clust-UGC0002344 | 2201 | 2 | 1 | 0 | 0 |
| UGC0011676 | Clust-UGC0002344 | 4456 | 3 | 1 | 0 | 1 |
| UGC0009175 | Clust-UGC0002344 | 2369 | 2 | 1 | 0 | 1 |
| UGC0012379 | Clust-UGC0002344 | 6005 | 3 | 1 | 0 | 1 |
| UGC0012131 | Clust-UGC0002344 | 1356 | 1 | 0 | 0 | 0 |
| UGC0011448 | Clust-UGC0002344 | 8159 | 4 | 1 | 0 | 1 |
| UGC0012034 | Clust-UGC0002344 | 1642 | 2 | 1 | 1 | 0 |
| UGC0006734 | Clust-UGC0002344 | 270 | 1 | 0 | 0 | 0 |
| UGC0006662 | Clust-UGC0002344 | 4802 | 1 | 0 | 0 | 0 |
| UGC0013222 | Clust-UGC0002344 | 3138 | 2 | 1 | 1 | 0 |
| UGC0005917 | Clust-UGC0002344 | 4343 | 1 | 0 | 0 | 0 |
| UGC0005052 | Clust-UGC0002344 | 2120 | 1 | 0 | 0 | 0 |
| UGC0011402 | Clust-UGC0002344 | 2239 | 1 | 0 | 0 | 0 |
| UGC0010204 | Clust-UGC0002344 | 154 | 1 | 0 | 0 | 0 |
| UGC0009248 | Clust-UGC0002344 | 2798 | 1 | 0 | 0 | 0 |
| UGC0005930 | Clust-UGC0002344 | 3882 | 2 | 1 | 0 | 0 |
| UGC0007425 | Clust-UGC0002344 | 107 | 1 | 0 | 0 | 0 |
| UGC0007423 | Clust-UGC0002344 | 1140 | 1 | 0 | 0 | 0 |
| UGC0006870 | Clust-UGC0002344 | 2008 | 2 | 1 | 0 | 0 |
| UGC0011147 | Clust-UGC0002344 | 1329 | 4 | 1 | 0 | 0 |
| UGC0009471 | Clust-UGC0002344 | 516 | 1 | 0 | 0 | 0 |
| UGC0007163 | Clust-UGC0002344 | 1192 | 1 | 0 | 0 | 0 |
| UGC0012739 | Clust-UGC0002344 | 1067 | 1 | 0 | 0 | 0 |
| UGC0011422 | Clust-UGC0002344 | 1937 | 2 | 1 | 0 | 0 |
| UGC0005955 | Clust-UGC0002344 | 2425 | 1 | 0 | 0 | 0 |
| UGC0006121 | Clust-UGC0002344 | 1771 | 1 | 0 | 0 | 0 |
| UGC0005934 | Clust-UGC0002344 | 3845 | 2 | 1 | 0 | 1 |
| UGC0011009 | Clust-UGC0002344 | 225 | 1 | 0 | 0 | 0 |
| UGC0013092 | Clust-UGC0002344 | 1283 | 1 | 0 | 0 | 0 |
| UGC0006956 | Clust-UGC0002344 | 3413 | 1 | 0 | 0 | 0 |
| UGC0006951 | Clust-UGC0002344 | 2708 | 1 | 0 | 0 | 0 |
| UGC0006406 | Clust-UGC0002344 | 5508 | 4 | 1 | 0 | 0 |
| UGC0010832 | Clust-UGC0002344 | 1272 | 2 | 1 | 0 | 0 |
| UGC0012968 | Clust-UGC0002344 | 1530 | 1 | 0 | 0 | 0 |
| UGC0006320 | Clust-UGC0002344 | 6086 | 2 | 1 | 0 | 1 |
| UGC0010860 | Clust-UGC0002344 | 1471 | 1 | 0 | 0 | 0 |
| UGC0010198 | Clust-UGC0002344 | 2400 | 1 | 0 | 0 | 0 |
| UGC0009085 | Clust-UGC0002344 | 2327 | 1 | 0 | 0 | 0 |
| UGC0006420 | Clust-UGC0002344 | 4172 | 3 | 1 | 0 | 0 |
| UGC0007485 | Clust-UGC0002344 | 3569 | 1 | 0 | 0 | 0 |
| UGC0007104 | Clust-UGC0002344 | 1278 | 1 | 0 | 0 | 0 |
| UGC0009609 | Clust-UGC0002344 | 1166 | 1 | 0 | 0 | 0 |
| UGC0005925 | Clust-UGC0002344 | 3237 | 2 | 1 | 0 | 1 |
| UGC0013337 | Clust-UGC0002344 | 3984 | 6 | 1 | 1 | 1 |
| UGC0012002 | Clust-UGC0002344 | 2005 | 1 | 0 | 0 | 0 |
| UGC0006125 | Clust-UGC0002344 | 3275 | 1 | 0 | 0 | 0 |
| UGC0011204 | Clust-UGC0002344 | 1130 | 1 | 0 | 0 | 0 |
| UGC0008971 | Clust-UGC0002344 | 2941 | 3 | 1 | 1 | 1 |
| UGC0006102 | Clust-UGC0002344 | 2251 | 2 | 1 | 0 | 1 |
| UGC0006041 | Clust-UGC0002344 | 3820 | 2 | 1 | 0 | 1 |
| UGC0005823 | Clust-UGC0002344 | 4264 | 1 | 0 | 0 | 0 |
| UGC0008341 | Clust-UGC0002344 | 968 | 1 | 0 | 0 | 0 |
| UGC0006160 | Clust-UGC0002344 | 1136 | 1 | 0 | 0 | 0 |
| UGC0010111 | Clust-UGC0002344 | 4165 | 1 | 0 | 0 | 0 |
| UGC0007956 | Clust-UGC0002344 | 2452 | 2 | 1 | 0 | 1 |
| UGC0010964 | Clust-UGC0002344 | 4259 | 2 | 1 | 0 | 0 |
| UGC0006576 | Clust-UGC0002344 | 1482 | 1 | 0 | 0 | 0 |
| UGC0011183 | Clust-UGC0002344 | 1957 | 1 | 0 | 0 | 0 |
| UGC0005795 | Clust-UGC0002344 | 5806 | 1 | 0 | 0 | 0 |
| UGC0007936 | Clust-UGC0002344 | 5427 | 1 | 0 | 0 | 0 |
| UGC0010943 | Clust-UGC0002344 | 747 | 1 | 0 | 0 | 0 |

| | | | | | | |
|---|---|---|---|---|---|---|
| UGC0011950 | Clust-UGC0002344 | 2642 | 1 | 0 | 0 | 0 |
| UGC0004173 | Clust-UGC0002344 | 2357 | 1 | 0 | 0 | 0 |
| UGC0006539 | Clust-UGC0002344 | 1125 | 1 | 0 | 0 | 0 |
| UGC0011496 | Clust-UGC0002344 | 545 | 1 | 0 | 0 | 0 |
| UGC0010013 | Clust-UGC0002344 | 2894 | 1 | 0 | 0 | 0 |
| UGC0010546 | Clust-UGC0002344 | 1687 | 1 | 0 | 0 | 0 |
| UGC0008852 | Clust-UGC0002344 | 408 | 1 | 0 | 0 | 0 |
| UGC0005988 | Clust-UGC0002344 | 2167 | 1 | 0 | 0 | 0 |
| UGC0012354 | Clust-UGC0002344 | 3327 | 1 | 0 | 0 | 0 |
| UGC0009450 | Clust-UGC0002344 | 1625 | 1 | 0 | 0 | 0 |
| UGC0012160 | Clust-UGC0002344 | 2320 | 1 | 0 | 0 | 0 |
| UGC0012293 | Clust-UGC0002344 | 1073 | 1 | 0 | 0 | 0 |
| UGC0012292 | Clust-UGC0002344 | 1208 | 2 | 1 | 0 | 1 |
| UGC0007577 | Clust-UGC0002344 | 4381 | 1 | 0 | 0 | 0 |
| UGC0013210 | Clust-UGC0002344 | 264 | 1 | 0 | 0 | 0 |
| UGC0005881 | Clust-UGC0002344 | 2014 | 1 | 0 | 0 | 0 |
| UGC0011454 | Clust-UGC0002344 | 7653 | 3 | 1 | 0 | 1 |
| UGC0012812 | Clust-UGC0002344 | 1770 | 2 | 1 | 0 | 1 |
| UGC0011660 | Clust-UGC0002344 | 4471 | 4 | 1 | 0 | 1 |
| UGC0011663 | Clust-UGC0002344 | 3586 | 5 | 1 | 0 | 0 |
| UGC0004736 | Clust-UGC0002344 | 1251 | 1 | 0 | 0 | 0 |
| UGC0010027 | Clust-UGC0002344 | 947 | 1 | 0 | 0 | 0 |
| UGC0009126 | Clust-UGC0002344 | 5987 | 1 | 0 | 0 | 0 |
| UGC0007358 | Clust-UGC0002344 | 1066 | 1 | 0 | 0 | 0 |
| UGC0012787 | Clust-UGC0002344 | 2885 | 2 | 1 | 0 | 1 |
| UGC0011006 | Clust-UGC0002344 | 3126 | 2 | 1 | 0 | 1 |
| UGC0007033 | Clust-UGC0002344 | 3478 | 3 | 1 | 0 | 0 |
| UGC0005835 | Clust-UGC0002344 | 4112 | 2 | 1 | 0 | 0 |
| UGC0004951 | Clust-UGC0002344 | 2447 | 1 | 0 | 0 | 0 |
| UGC0007961 | Clust-UGC0002344 | 2282 | 1 | 0 | 0 | 0 |
| UGC0007375 | Clust-UGC0002344 | 859 | 1 | 0 | 0 | 0 |
| UGC0012859 | Clust-UGC0002344 | 6767 | 3 | 1 | 0 | 1 |
| UGC0009012 | Clust-UGC0002344 | 6597 | 2 | 1 | 0 | 1 |
| UGC0010575 | Clust-UGC0002344 | 2495 | 1 | 0 | 0 | 0 |
| UGC0013272 | Clust-UGC0002344 | 1396 | 1 | 0 | 0 | 0 |
| UGC0007174 | Clust-UGC0002344 | 4976 | 2 | 1 | 1 | 0 |
| UGC0005897 | Clust-UGC0002344 | 4829 | 2 | 1 | 0 | 1 |
| UGC0007315 | Clust-UGC0002344 | 4798 | 1 | 0 | 0 | 0 |
| UGC0005675 | Clust-UGC0002344 | 3266 | 1 | 0 | 0 | 0 |
| UGC0011436 | Clust-UGC0002344 | 5520 | 4 | 1 | 0 | 0 |
| UGC0010814 | Clust-UGC0002344 | 1381 | 4 | 1 | 0 | 0 |
| UGC0006016 | Clust-UGC0002344 | 3817 | 2 | 1 | 0 | 1 |
| UGC0009447 | Clust-UGC0002344 | 1571 | 1 | 0 | 0 | 0 |
| UGC0009440 | Clust-UGC0002344 | 565 | 1 | 0 | 0 | 0 |
| UGC0012197 | Clust-UGC0002344 | 1392 | 1 | 0 | 0 | 0 |
| UGC0007159 | Clust-UGC0002344 | 2352 | 1 | 0 | 0 | 0 |
| UGC0012542 | Clust-UGC0002344 | 6185 | 5 | 1 | 0 | 1 |
| UGC0010763 | Clust-UGC0002344 | 2222 | 2 | 1 | 0 | 0 |
| UGC0006943 | Clust-UGC0002344 | 1182 | 1 | 0 | 0 | 0 |
| UGC0009058 | Clust-UGC0002344 | 2403 | 1 | 0 | 0 | 0 |
| UGC0006620 | Clust-UGC0002344 | 1802 | 1 | 0 | 0 | 0 |
| UGC0006435 | Clust-UGC0002344 | 6169 | 3 | 1 | 0 | 1 |
| UGC0007473 | Clust-UGC0002344 | 2784 | 1 | 0 | 0 | 0 |
| UGC0009409 | Clust-UGC0002344 | 2899 | 2 | 1 | 0 | 1 |
| UGC0012583 | Clust-UGC0002344 | 3863 | 2 | 1 | 0 | 0 |
| UGC0006606 | Clust-UGC0002344 | 1907 | 1 | 0 | 0 | 0 |
| UGC0002344 | Clust-UGC0002344 | 2226 | 1 | 0 | 0 | 0 |
| UGC0009394 | Clust-UGC0002344 | 3612 | 2 | 1 | 0 | 1 |
| UGC0006844 | Clust-UGC0002344 | 2372 | 1 | 0 | 0 | 0 |
| UGC0010893 | Clust-UGC0002344 | 1597 | 2 | 1 | 0 | 0 |
| UGC0008635 | Clust-UGC0002344 | 4859 | 3 | 1 | 1 | 1 |
| UGC0005873 | Clust-UGC0002344 | 5199 | 1 | 0 | 0 | 0 |
| UGC0005876 | Clust-UGC0002344 | 4544 | 1 | 0 | 0 | 0 |
| UGC0012264 | Clust-UGC0002344 | 706 | 1 | 0 | 0 | 0 |
| UGC0012265 | Clust-UGC0002344 | 669 | 1 | 0 | 0 | 0 |
| UGC0008396 | Clust-UGC0002344 | 1381 | 1 | 0 | 0 | 0 |
| UGC0002502 | Clust-UGC0002344 | 2202 | 1 | 0 | 0 | 0 |
| UGC0006134 | Clust-UGC0002344 | 2403 | 3 | 1 | 0 | 1 |
| UGC0012019 | Clust-UGC0002344 | 407 | 2 | 1 | 1 | 0 |
| UGC0009155 | Clust-UGC0002344 | 1039 | 1 | 0 | 0 | 0 |
| UGC0010915 | Clust-UGC0002344 | 505 | 1 | 0 | 0 | 0 |
| UGC0006421 | Clust-UGC0002344 | 2343 | 3 | 1 | 0 | 0 |
| UGC0006112 | Clust-UGC0002344 | 1743 | 1 | 0 | 0 | 0 |
| UGC0006110 | Clust-UGC0002344 | 1949 | 1 | 0 | 0 | 0 |
| UGC0006114 | Clust-UGC0002344 | 2214 | 1 | 0 | 0 | 0 |
| UGC0010120 | Clust-UGC0002344 | 2314 | 1 | 0 | 0 | 0 |
| UGC0007745 | Clust-UGC0002344 | 6074 | 2 | 1 | 0 | 0 |
| UGC0009489 | Clust-UGC0002344 | 3050 | 1 | 0 | 0 | 0 |
| UGC0006352 | Clust-UGC0002344 | 7251 | 4 | 1 | 0 | 1 |

| | | | | | | |
|---|---|---|---|---|---|---|
| UGC0006682 | Clust-UGC0002344 | 242 | 1 | 0 | 0 | 0 |
| UGC0006174 | Clust-UGC0002344 | 1384 | 1 | 0 | 0 | 0 |
| UGC0010102 | Clust-UGC0002344 | 6604 | 4 | 1 | 0 | 1 |
| UGC0013287 | Clust-UGC0002344 | 3302 | 5 | 1 | 3 | 1 |
| UGC0005818 | Clust-UGC0002344 | 6637 | 4 | 1 | 0 | 1 |
| UGC0010952 | Clust-UGC0002344 | 4242 | 2 | 1 | 0 | 0 |
| UGC0011195 | Clust-UGC0002344 | 8592 | 8 | 1 | 1 | 1 |
| UGC0011646 | Clust-UGC0002344 | 2977 | 2 | 1 | 0 | 0 |
| UGC0012847 | Clust-UGC0002344 | 4772 | 4 | 1 | 0 | 1 |
| UGC0009316 | Clust-UGC0002344 | 1385 | 1 | 0 | 0 | 0 |
| UGC0006378 | Clust-UGC0002344 | 5340 | 6 | 1 | 0 | 1 |
| UGC0011949 | Clust-UGC0002344 | 1440 | 1 | 0 | 0 | 0 |
| UGC0006159 | Clust-UGC0002344 | 664 | 1 | 0 | 0 | 0 |
| UGC0007204 | Clust-UGC0002348 | 5160 | 4 | 1 | 0 | 1 |
| UGC0007205 | Clust-UGC0002348 | 852 | 1 | 0 | 0 | 0 |
| UGC0006557 | Clust-UGC0002348 | 1943 | 1 | 0 | 0 | 0 |
| UGC0006526 | Clust-UGC0002348 | 1889 | 1 | 0 | 0 | 0 |
| UGC0009736 | Clust-UGC0002348 | 3270 | 1 | 0 | 0 | 0 |
| UGC0010635 | Clust-UGC0002348 | 314 | 1 | 0 | 0 | 0 |
| UGC0004767 | Clust-UGC0002348 | 1351 | 2 | 1 | 0 | 1 |
| UGC0006673 | Clust-UGC0002348 | 814 | 2 | 1 | 0 | 1 |
| UGC0009507 | Clust-UGC0002348 | 1784 | 6 | 1 | 0 | 0 |
| UGC0004830 | Clust-UGC0002348 | 12504 | 64 | 1 | 0 | 1 |
| UGC0006509 | Clust-UGC0002348 | 1571 | 2 | 1 | 0 | 1 |
| UGC0012286 | Clust-UGC0002348 | 1728 | 1 | 0 | 0 | 0 |
| UGC0012285 | Clust-UGC0002348 | 768 | 1 | 0 | 0 | 0 |
| UGC0007675 | Clust-UGC0002348 | 5972 | 12 | 1 | 0 | 1 |
| UGC0012825 | Clust-UGC0002348 | 837 | 1 | 0 | 0 | 0 |
| UGC0007061 | Clust-UGC0002348 | 2560 | 2 | 1 | 0 | 0 |
| UGC0008514 | Clust-UGC0002348 | 2387 | 2 | 1 | 0 | 0 |
| UGC0007652 | Clust-UGC0002348 | 4162 | 4 | 1 | 0 | 1 |
| UGC0010096 | Clust-UGC0002348 | 1496 | 1 | 0 | 0 | 0 |
| UGC0004721 | Clust-UGC0002348 | 602 | 1 | 0 | 0 | 0 |
| UGC0009286 | Clust-UGC0002348 | 761 | 1 | 0 | 0 | 0 |
| UGC0008887 | Clust-UGC0002348 | 2057 | 1 | 0 | 0 | 0 |
| UGC0007347 | Clust-UGC0002348 | 4282 | 7 | 1 | 0 | 1 |
| UGC0006512 | Clust-UGC0002348 | 1317 | 1 | 0 | 0 | 0 |
| UGC0006769 | Clust-UGC0002348 | 5775 | 8 | 1 | 0 | 1 |
| UGC0004516 | Clust-UGC0002348 | 9039 | 20 | 1 | 0 | 1 |
| UGC0009063 | Clust-UGC0002348 | 1503 | 3 | 1 | 0 | 1 |
| UGC0007424 | Clust-UGC0002348 | 3274 | 2 | 1 | 0 | 0 |
| UGC0010561 | Clust-UGC0002348 | 1542 | 4 | 1 | 0 | 1 |
| UGC0006947 | Clust-UGC0002348 | 5961 | 5 | 1 | 0 | 1 |
| UGC0006794 | Clust-UGC0002348 | 6805 | 14 | 1 | 1 | 1 |
| UGC0006891 | Clust-UGC0002348 | 3424 | 1 | 0 | 0 | 0 |
| UGC0007144 | Clust-UGC0002348 | 3075 | 1 | 0 | 0 | 0 |
| UGC0009206 | Clust-UGC0002348 | 2120 | 1 | 0 | 0 | 0 |
| UGC0006955 | Clust-UGC0002348 | 3058 | 3 | 1 | 0 | 1 |
| UGC0010521 | Clust-UGC0002348 | 3536 | 6 | 1 | 0 | 1 |
| UGC0006615 | Clust-UGC0002348 | 514 | 1 | 0 | 0 | 0 |
| UGC0009222 | Clust-UGC0002348 | 4154 | 4 | 1 | 0 | 1 |
| UGC0007480 | Clust-UGC0002348 | 3738 | 2 | 1 | 0 | 1 |
| UGC0007270 | Clust-UGC0002348 | 5658 | 6 | 1 | 0 | 1 |
| UGC0006853 | Clust-UGC0002348 | 3108 | 2 | 1 | 0 | 1 |
| UGC0007103 | Clust-UGC0002348 | 2023 | 1 | 0 | 0 | 0 |
| UGC0007102 | Clust-UGC0002348 | 2199 | 1 | 0 | 0 | 0 |
| UGC0002948 | Clust-UGC0002348 | 824 | 1 | 0 | 0 | 0 |
| UGC0005866 | Clust-UGC0002348 | 5025 | 4 | 1 | 1 | 1 |
| UGC0004777 | Clust-UGC0002348 | 191 | 1 | 0 | 0 | 0 |
| UGC0005529 | Clust-UGC0002348 | 2641 | 1 | 0 | 0 | 0 |
| UGC0006658 | Clust-UGC0002348 | 2368 | 1 | 0 | 0 | 0 |
| UGC0007389 | Clust-UGC0002348 | 1499 | 1 | 0 | 0 | 0 |
| UGC0007529 | Clust-UGC0002348 | 2427 | 1 | 0 | 0 | 0 |
| UGC0007521 | Clust-UGC0002348 | 4145 | 2 | 1 | 0 | 1 |
| UGC0006812 | Clust-UGC0002348 | 4017 | 4 | 1 | 0 | 1 |
| UGC0002550 | Clust-UGC0002348 | 1195 | 1 | 0 | 0 | 0 |
| UGC0009346 | Clust-UGC0002348 | 4119 | 7 | 1 | 1 | 0 |
| UGC0007563 | Clust-UGC0002348 | 5947 | 3 | 1 | 0 | 1 |
| UGC0010114 | Clust-UGC0002348 | 2500 | 1 | 0 | 0 | 0 |
| UGC0012982 | Clust-UGC0002348 | 442 | 1 | 0 | 0 | 0 |
| UGC0006574 | Clust-UGC0002348 | 2460 | 2 | 1 | 0 | 1 |
| UGC0007549 | Clust-UGC0002348 | 7114 | 6 | 1 | 0 | 1 |
| UGC0007547 | Clust-UGC0002348 | 7258 | 11 | 1 | 0 | 1 |
| UGC0007542 | Clust-UGC0002348 | 6054 | 3 | 1 | 0 | 1 |
| UGC0007540 | Clust-UGC0002348 | 4630 | 2 | 1 | 0 | 0 |
| UGC0003006 | Clust-UGC0002348 | 409 | 1 | 0 | 0 | 0 |
| UGC0006535 | Clust-UGC0002348 | 1397 | 1 | 0 | 0 | 0 |
| UGC0006534 | Clust-UGC0002348 | 1450 | 1 | 0 | 0 | 0 |
| UGC0007588 | Clust-UGC0002348 | 3129 | 3 | 1 | 0 | 1 |

| | | | | | | |
|---|---|---|---|---|---|---|
| UGC0002348 | Clust-UGC0002348 | 141 | 1 | 0 | 0 | 0 |
| UGC0009451 | Clust-UGC0002348 | 224 | 1 | 0 | 0 | 0 |
| UGC0006948 | Clust-UGC0002348 | 4429 | 4 | 1 | 0 | 1 |
| UGC0003773 | Clust-UGC0002348 | 574 | 2 | 1 | 0 | 0 |
| UGC0007576 | Clust-UGC0002348 | 4644 | 4 | 1 | 0 | 1 |
| UGC0004137 | Clust-UGC0002348 | 60 | 1 | 0 | 0 | 0 |
| UGC0006495 | Clust-UGC0002348 | 1034 | 1 | 0 | 0 | 0 |
| UGC0004730 | Clust-UGC0002348 | 504 | 1 | 0 | 0 | 0 |
| UGC0003038 | Clust-UGC0002348 | 961 | 1 | 0 | 0 | 0 |
| UGC0009032 | Clust-UGC0002348 | 317 | 1 | 0 | 0 | 0 |
| UGC0009037 | Clust-UGC0002348 | 3765 | 11 | 1 | 0 | 1 |
| UGC0007293 | Clust-UGC0002348 | 6292 | 14 | 1 | 0 | 1 |
| UGC0005599 | Clust-UGC0002348 | 2123 | 1 | 0 | 0 | 0 |
| UGC0002531 | Clust-UGC0002348 | 4153 | 2 | 1 | 0 | 0 |
| UGC0010236 | Clust-UGC0002348 | 1051 | 1 | 0 | 0 | 0 |
| UGC0007312 | Clust-UGC0002348 | 6044 | 8 | 1 | 0 | 1 |
| UGC0010212 | Clust-UGC0002348 | 1143 | 2 | 1 | 0 | 0 |
| UGC0008033 | Clust-UGC0002348 | 209 | 1 | 0 | 0 | 0 |
| UGC0006968 | Clust-UGC0002348 | 4531 | 3 | 1 | 0 | 1 |
| UGC0003752 | Clust-UGC0002348 | 145 | 1 | 0 | 0 | 0 |
| UGC0007333 | Clust-UGC0002348 | 403 | 1 | 0 | 0 | 0 |
| UGC0009059 | Clust-UGC0002348 | 2552 | 1 | 0 | 0 | 0 |
| UGC0003357 | Clust-UGC0002348 | 9912 | 31 | 1 | 0 | 1 |
| UGC0006881 | Clust-UGC0002348 | 4786 | 3 | 1 | 1 | 1 |
| UGC0012705 | Clust-UGC0002348 | 7272 | 13 | 1 | 0 | 1 |
| UGC0007557 | Clust-UGC0002348 | 1667 | 2 | 1 | 0 | 0 |
| UGC0010513 | Clust-UGC0002348 | 2987 | 2 | 1 | 0 | 0 |
| UGC0012767 | Clust-UGC0002348 | 2112 | 1 | 0 | 0 | 0 |
| UGC0011588 | Clust-UGC0002348 | 317 | 1 | 0 | 0 | 0 |
| UGC0008457 | Clust-UGC0002348 | 1992 | 2 | 1 | 0 | 1 |
| UGC0007493 | Clust-UGC0002348 | 3120 | 2 | 1 | 0 | 1 |
| UGC0012243 | Clust-UGC0002348 | 6695 | 10 | 1 | 0 | 1 |
| UGC0006996 | Clust-UGC0002348 | 5203 | 3 | 1 | 0 | 1 |
| UGC0003034 | Clust-UGC0002348 | 469 | 1 | 0 | 0 | 0 |
| UGC0011854 | Clust-UGC0002348 | 2248 | 4 | 1 | 0 | 1 |
| UGC0007536 | Clust-UGC0002348 | 3075 | 3 | 1 | 0 | 1 |
| UGC0008968 | Clust-UGC0002348 | 1893 | 2 | 1 | 1 | 0 |
| UGC0006731 | Clust-UGC0002348 | 1271 | 2 | 1 | 0 | 0 |
| UGC0003790 | Clust-UGC0002348 | 272 | 1 | 0 | 0 | 0 |
| UGC0006868 | Clust-UGC0002348 | 3092 | 2 | 1 | 0 | 1 |
| UGC0005119 | Clust-UGC0002348 | 2289 | 1 | 0 | 0 | 0 |
| UGC0003020 | Clust-UGC0002348 | 1460 | 1 | 0 | 0 | 0 |
| UGC0004096 | Clust-UGC0002348 | 121 | 1 | 0 | 0 | 0 |
| UGC0009589 | Clust-UGC0002348 | 1084 | 1 | 0 | 0 | 0 |
| UGC0009636 | Clust-UGC0002348 | 1325 | 1 | 0 | 0 | 0 |
| UGC0010340 | Clust-UGC0002348 | 373 | 2 | 1 | 0 | 1 |
| UGC0009033 | Clust-UGC0002348 | 213 | 1 | 0 | 0 | 0 |
| UGC0006563 | Clust-UGC0002348 | 95 | 1 | 0 | 0 | 0 |
| UGC0006567 | Clust-UGC0002348 | 2283 | 1 | 0 | 0 | 0 |
| UGC0004466 | Clust-UGC0002348 | 6758 | 14 | 1 | 0 | 1 |
| UGC0003279 | Clust-UGC0002348 | 3633 | 2 | 1 | 0 | 1 |
| UGC0011965 | Clust-UGC0002348 | 4109 | 3 | 1 | 0 | 1 |
| UGC0006683 | Clust-UGC0002348 | 1201 | 3 | 1 | 0 | 0 |
| UGC0006680 | Clust-UGC0002348 | 1358 | 1 | 0 | 0 | 0 |
| UGC0007617 | Clust-UGC0002348 | 7350 | 22 | 1 | 0 | 1 |
| UGC0007610 | Clust-UGC0002348 | 6154 | 6 | 1 | 0 | 1 |
| UGC0002478 | Clust-UGC0002348 | 154 | 1 | 0 | 0 | 0 |
| UGC0003518 | Clust-UGC0002361 | 2213 | 1 | 0 | 0 | 0 |
| UGC0010907 | Clust-UGC0002361 | 1241 | 2 | 1 | 0 | 0 |
| UGC0009537 | Clust-UGC0002361 | 3135 | 3 | 1 | 0 | 1 |
| UGC0002361 | Clust-UGC0002361 | 1148 | 1 | 0 | 0 | 0 |
| UGC0008587 | Clust-UGC0002361 | 8016 | 5 | 1 | 0 | 1 |
| UGC0006395 | Clust-UGC0002361 | 9194 | 12 | 1 | 0 | 1 |
| UGC0011526 | Clust-UGC0002361 | 8541 | 2 | 1 | 0 | 1 |
| UGC0008762 | Clust-UGC0002361 | 868 | 1 | 0 | 0 | 0 |
| UGC0010164 | Clust-UGC0002362 | 5238 | 3 | 1 | 0 | 1 |
| UGC0010163 | Clust-UGC0002362 | 7578 | 23 | 1 | 0 | 1 |
| UGC0006552 | Clust-UGC0002362 | 1893 | 2 | 1 | 0 | 1 |
| UGC0004838 | Clust-UGC0002362 | 19173 | 132 | 1 | 1 | 1 |
| UGC0005591 | Clust-UGC0002362 | 3179 | 5 | 1 | 0 | 1 |
| UGC0005372 | Clust-UGC0002362 | 1229 | 1 | 0 | 0 | 0 |
| UGC0004835 | Clust-UGC0002362 | 20575 | 188 | 1 | 1 | 1 |
| UGC0007246 | Clust-UGC0002362 | 8634 | 30 | 1 | 0 | 1 |
| UGC0005077 | Clust-UGC0002362 | 4294 | 6 | 1 | 0 | 1 |
| UGC0005619 | Clust-UGC0002362 | 4287 | 5 | 1 | 0 | 1 |
| UGC0005581 | Clust-UGC0002362 | 2065 | 4 | 1 | 0 | 1 |
| UGC0005583 | Clust-UGC0002362 | 3354 | 2 | 1 | 0 | 1 |
| UGC0005582 | Clust-UGC0002362 | 2354 | 1 | 0 | 0 | 0 |
| UGC0005055 | Clust-UGC0002362 | 2569 | 2 | 1 | 0 | 1 |

| | | | | | | |
|---|---|---|---|---|---|---|
| UGC0005054 | Clust-UGC0002362 | 3812 | 2 | 1 | 0 | 1 |
| UGC0005700 | Clust-UGC0002362 | 1347 | 2 | 1 | 0 | 1 |
| UGC0005549 | Clust-UGC0002362 | 4171 | 2 | 1 | 0 | 1 |
| UGC0005363 | Clust-UGC0002362 | 5832 | 10 | 1 | 0 | 1 |
| UGC0005561 | Clust-UGC0002362 | 5262 | 8 | 1 | 0 | 1 |
| UGC0005690 | Clust-UGC0002362 | 479 | 1 | 0 | 0 | 0 |
| UGC0012593 | Clust-UGC0002362 | 2104 | 3 | 1 | 0 | 1 |
| UGC0005738 | Clust-UGC0002362 | 8437 | 27 | 1 | 0 | 1 |
| UGC0006467 | Clust-UGC0002362 | 2392 | 1 | 0 | 0 | 0 |
| UGC0005711 | Clust-UGC0002362 | 8781 | 31 | 1 | 1 | 1 |
| UGC0005716 | Clust-UGC0002362 | 7107 | 15 | 1 | 0 | 1 |
| UGC0008607 | Clust-UGC0002362 | 241 | 1 | 0 | 0 | 0 |
| UGC0006649 | Clust-UGC0002362 | 725 | 1 | 0 | 0 | 0 |
| UGC0008384 | Clust-UGC0002362 | 289 | 1 | 0 | 0 | 0 |
| UGC0005322 | Clust-UGC0002362 | 5166 | 5 | 1 | 0 | 1 |
| UGC0005326 | Clust-UGC0002362 | 4604 | 5 | 1 | 0 | 1 |
| UGC0008873 | Clust-UGC0002362 | 2654 | 2 | 1 | 0 | 1 |
| UGC0008878 | Clust-UGC0002362 | 4994 | 4 | 1 | 0 | 1 |
| UGC0010159 | Clust-UGC0002362 | 3950 | 6 | 1 | 0 | 1 |
| UGC0007914 | Clust-UGC0002362 | 4961 | 12 | 1 | 0 | 1 |
| UGC0005649 | Clust-UGC0002362 | 4935 | 12 | 1 | 0 | 1 |
| UGC0005661 | Clust-UGC0002362 | 5550 | 5 | 1 | 0 | 1 |
| UGC0002362 | Clust-UGC0002362 | 2378 | 2 | 1 | 0 | 1 |
| UGC0008569 | Clust-UGC0002362 | 3636 | 4 | 1 | 0 | 1 |
| UGC0008179 | Clust-UGC0002362 | 2585 | 1 | 0 | 0 | 0 |
| UGC0005593 | Clust-UGC0002362 | 2740 | 2 | 1 | 0 | 1 |
| UGC0007192 | Clust-UGC0002362 | 8782 | 31 | 1 | 0 | 1 |
| UGC0005685 | Clust-UGC0002362 | 2448 | 2 | 1 | 0 | 1 |
| UGC0007432 | Clust-UGC0002362 | 4079 | 5 | 1 | 0 | 1 |
| UGC0008018 | Clust-UGC0002362 | 7029 | 18 | 1 | 0 | 1 |
| UGC0007452 | Clust-UGC0002362 | 3850 | 1 | 0 | 0 | 0 |
| UGC0005574 | Clust-UGC0002362 | 1549 | 1 | 0 | 0 | 0 |
| UGC0005571 | Clust-UGC0002362 | 2579 | 2 | 1 | 0 | 1 |
| UGC0005515 | Clust-UGC0002362 | 2660 | 4 | 1 | 0 | 1 |
| UGC0007236 | Clust-UGC0002362 | 8212 | 22 | 1 | 0 | 1 |
| UGC0007749 | Clust-UGC0002362 | 8607 | 27 | 1 | 0 | 1 |
| UGC0012673 | Clust-UGC0002366 | 6882 | 17 | 1 | 0 | 1 |
| UGC0009997 | Clust-UGC0002366 | 1429 | 4 | 1 | 0 | 0 |
| UGC0002549 | Clust-UGC0002366 | 435 | 1 | 0 | 0 | 0 |
| UGC0002366 | Clust-UGC0002366 | 2447 | 4 | 1 | 0 | 1 |
| UGC0006558 | Clust-UGC0002372 | 4154 | 4 | 1 | 0 | 1 |
| UGC0007185 | Clust-UGC0002372 | 7590 | 20 | 1 | 0 | 1 |
| UGC0002372 | Clust-UGC0002372 | 2018 | 1 | 0 | 0 | 0 |
| UGC0005752 | Clust-UGC0002372 | 3275 | 7 | 1 | 0 | 1 |
| UGC0010211 | Clust-UGC0002372 | 2037 | 5 | 1 | 0 | 1 |
| UGC0009108 | Clust-UGC0002372 | 1154 | 2 | 1 | 0 | 1 |
| UGC0003770 | Clust-UGC0002372 | 203 | 1 | 0 | 0 | 0 |
| UGC0006305 | Clust-UGC0002372 | 3542 | 1 | 0 | 0 | 0 |
| UGC0007662 | Clust-UGC0002372 | 7222 | 13 | 1 | 0 | 1 |
| UGC0008547 | Clust-UGC0002372 | 985 | 1 | 0 | 0 | 0 |
| UGC0011890 | Clust-UGC0002372 | 7804 | 13 | 1 | 0 | 1 |
| UGC0007199 | Clust-UGC0002372 | 2892 | 5 | 1 | 0 | 1 |
| UGC0006599 | Clust-UGC0002372 | 1615 | 2 | 1 | 0 | 1 |
| UGC0011992 | Clust-UGC0002372 | 1522 | 2 | 1 | 0 | 0 |
| UGC0003793 | Clust-UGC0002372 | 113 | 1 | 0 | 0 | 0 |
| UGC0004669 | Clust-UGC0002372 | 7629 | 35 | 1 | 0 | 1 |
| UGC0005331 | Clust-UGC0002372 | 1646 | 1 | 0 | 0 | 0 |
| UGC0007887 | Clust-UGC0002372 | 3513 | 4 | 1 | 0 | 1 |
| UGC0006152 | Clust-UGC0002372 | 1966 | 2 | 1 | 0 | 1 |
| UGC0005560 | Clust-UGC0002471 | 1995 | 1 | 0 | 0 | 0 |
| UGC0002471 | Clust-UGC0002471 | 111 | 1 | 0 | 0 | 0 |
| UGC0008315 | Clust-UGC0002477 | 1652 | 1 | 0 | 0 | 0 |
| UGC0004128 | Clust-UGC0002477 | 904 | 1 | 0 | 0 | 0 |
| UGC0005655 | Clust-UGC0002477 | 1580 | 2 | 1 | 0 | 1 |
| UGC0006717 | Clust-UGC0002477 | 4125 | 1 | 0 | 0 | 0 |
| UGC0007218 | Clust-UGC0002477 | 3215 | 2 | 1 | 0 | 1 |
| UGC0004707 | Clust-UGC0002477 | 19008 | 29 | 1 | 0 | 1 |
| UGC0004511 | Clust-UGC0002477 | 5448 | 2 | 1 | 0 | 1 |
| UGC0004552 | Clust-UGC0002477 | 1629 | 1 | 0 | 0 | 0 |
| UGC0013094 | Clust-UGC0002477 | 780 | 2 | 1 | 1 | 0 |
| UGC0008643 | Clust-UGC0002477 | 1350 | 1 | 0 | 0 | 0 |
| UGC0003578 | Clust-UGC0002477 | 5127 | 1 | 0 | 0 | 0 |
| UGC0003573 | Clust-UGC0002477 | 5723 | 1 | 0 | 0 | 0 |
| UGC0003577 | Clust-UGC0002477 | 8755 | 1 | 0 | 0 | 0 |
| UGC0012904 | Clust-UGC0002477 | 4141 | 3 | 1 | 0 | 1 |
| UGC0004655 | Clust-UGC0002477 | 10040 | 5 | 1 | 1 | 1 |
| UGC0007735 | Clust-UGC0002477 | 7154 | 2 | 1 | 0 | 1 |
| UGC0004674 | Clust-UGC0002477 | 11265 | 12 | 1 | 0 | 1 |
| UGC0008198 | Clust-UGC0002477 | 3438 | 1 | 0 | 0 | 0 |

| | | | | | | |
|---|---|---|---|---|---|---|
| UGC0007754 | Clust-UGC0002477 | 6219 | 3 | 1 | 0 | 1 |
| UGC0004473 | Clust-UGC0002477 | 4832 | 1 | 0 | 0 | 0 |
| UGC0004694 | Clust-UGC0002477 | 7594 | 3 | 1 | 0 | 1 |
| UGC0012238 | Clust-UGC0002477 | 4112 | 1 | 0 | 0 | 0 |
| UGC0008260 | Clust-UGC0002477 | 748 | 1 | 0 | 0 | 0 |
| UGC0004589 | Clust-UGC0002477 | 4516 | 2 | 1 | 0 | 1 |
| UGC0004779 | Clust-UGC0002477 | 1208 | 1 | 0 | 0 | 0 |
| UGC0004499 | Clust-UGC0002477 | 8101 | 3 | 1 | 1 | 1 |
| UGC0008287 | Clust-UGC0002477 | 843 | 1 | 0 | 0 | 0 |
| UGC0006946 | Clust-UGC0002477 | 3046 | 1 | 0 | 0 | 0 |
| UGC0004133 | Clust-UGC0002477 | 10051 | 5 | 1 | 0 | 1 |
| UGC0004507 | Clust-UGC0002477 | 9787 | 2 | 1 | 0 | 1 |
| UGC0004521 | Clust-UGC0002477 | 1464 | 1 | 0 | 0 | 0 |
| UGC0004522 | Clust-UGC0002477 | 5213 | 2 | 1 | 0 | 1 |
| UGC0004563 | Clust-UGC0002477 | 7268 | 2 | 1 | 0 | 1 |
| UGC0005518 | Clust-UGC0002477 | 491 | 1 | 0 | 0 | 0 |
| UGC0004601 | Clust-UGC0002477 | 2355 | 1 | 0 | 0 | 0 |
| UGC0003581 | Clust-UGC0002477 | 6650 | 2 | 1 | 0 | 1 |
| UGC0002951 | Clust-UGC0002477 | 570 | 1 | 0 | 0 | 0 |
| UGC0007725 | Clust-UGC0002477 | 7873 | 2 | 1 | 0 | 1 |
| UGC0007888 | Clust-UGC0002477 | 4475 | 1 | 0 | 0 | 0 |
| UGC0007761 | Clust-UGC0002477 | 7084 | 1 | 0 | 0 | 0 |
| UGC0011192 | Clust-UGC0002477 | 1564 | 2 | 1 | 0 | 0 |
| UGC0002477 | Clust-UGC0002477 | 242 | 1 | 0 | 0 | 0 |
| UGC0006524 | Clust-UGC0002481 | 699 | 1 | 0 | 0 | 0 |
| UGC0013014 | Clust-UGC0002481 | 3081 | 2 | 1 | 0 | 1 |
| UGC0002481 | Clust-UGC0002481 | 88 | 1 | 0 | 0 | 0 |
| UGC0009127 | Clust-UGC0002481 | 7293 | 14 | 1 | 0 | 1 |
| UGC0007566 | Clust-UGC0002481 | 5074 | 3 | 1 | 0 | 1 |
| UGC0007339 | Clust-UGC0002481 | 757 | 2 | 1 | 0 | 0 |
| UGC0005347 | Clust-UGC0002481 | 2537 | 4 | 1 | 0 | 1 |
| UGC0006350 | Clust-UGC0002481 | 6739 | 17 | 1 | 0 | 1 |
| UGC0006375 | Clust-UGC0002481 | 7491 | 19 | 1 | 0 | 1 |
| UGC0005783 | Clust-UGC0002483 | 7049 | 6 | 1 | 0 | 1 |
| UGC0009698 | Clust-UGC0002483 | 2699 | 2 | 1 | 0 | 0 |
| UGC0010993 | Clust-UGC0002483 | 6931 | 3 | 1 | 0 | 1 |
| UGC0003000 | Clust-UGC0002483 | 1575 | 1 | 0 | 0 | 0 |
| UGC0006074 | Clust-UGC0002483 | 1021 | 1 | 0 | 0 | 0 |
| UGC0006711 | Clust-UGC0002483 | 2447 | 2 | 1 | 0 | 1 |
| UGC0011678 | Clust-UGC0002483 | 3073 | 3 | 1 | 0 | 0 |
| UGC0009549 | Clust-UGC0002483 | 134 | 1 | 0 | 0 | 0 |
| UGC0006277 | Clust-UGC0002483 | 3415 | 1 | 0 | 0 | 0 |
| UGC0011651 | Clust-UGC0002483 | 10209 | 10 | 1 | 1 | 1 |
| UGC0008749 | Clust-UGC0002483 | 1725 | 1 | 0 | 0 | 0 |
| UGC0007181 | Clust-UGC0002483 | 5808 | 2 | 1 | 0 | 0 |
| UGC0010202 | Clust-UGC0002483 | 540 | 1 | 0 | 0 | 0 |
| UGC0010017 | Clust-UGC0002483 | 2988 | 2 | 1 | 0 | 0 |
| UGC0006092 | Clust-UGC0002483 | 3415 | 1 | 0 | 0 | 0 |
| UGC0008916 | Clust-UGC0002483 | 4373 | 2 | 1 | 0 | 0 |
| UGC0012384 | Clust-UGC0002483 | 10068 | 13 | 1 | 0 | 1 |
| UGC0007461 | Clust-UGC0002483 | 7466 | 8 | 1 | 0 | 1 |
| UGC0012966 | Clust-UGC0002483 | 6148 | 4 | 1 | 0 | 1 |
| UGC0002961 | Clust-UGC0002483 | 1390 | 1 | 0 | 0 | 0 |
| UGC0002509 | Clust-UGC0002483 | 3500 | 2 | 1 | 0 | 1 |
| UGC0002547 | Clust-UGC0002483 | 7398 | 4 | 1 | 0 | 1 |
| UGC0009415 | Clust-UGC0002483 | 5790 | 2 | 1 | 0 | 0 |
| UGC0009169 | Clust-UGC0002483 | 5263 | 4 | 1 | 0 | 1 |
| UGC0007386 | Clust-UGC0002483 | 3871 | 3 | 1 | 0 | 1 |
| UGC0006443 | Clust-UGC0002483 | 8571 | 11 | 1 | 0 | 1 |
| UGC0002483 | Clust-UGC0002483 | 3886 | 1 | 0 | 0 | 0 |
| UGC0006813 | Clust-UGC0002483 | 3139 | 1 | 0 | 0 | 0 |
| UGC0009105 | Clust-UGC0002483 | 2866 | 2 | 1 | 0 | 0 |
| UGC0012065 | Clust-UGC0002483 | 1006 | 1 | 0 | 0 | 0 |
| UGC0006368 | Clust-UGC0002483 | 8513 | 8 | 1 | 0 | 1 |
| UGC0007216 | Clust-UGC0002483 | 2075 | 2 | 1 | 0 | 0 |
| UGC0008261 | Clust-UGC0002483 | 2013 | 1 | 0 | 0 | 0 |
| UGC0009130 | Clust-UGC0002483 | 2410 | 2 | 1 | 0 | 0 |
| UGC0003754 | Clust-UGC0002483 | 439 | 1 | 0 | 0 | 0 |
| UGC0002984 | Clust-UGC0002483 | 1684 | 2 | 1 | 0 | 0 |
| UGC0011894 | Clust-UGC0002483 | 8144 | 5 | 1 | 0 | 1 |
| UGC0008710 | Clust-UGC0002483 | 1839 | 2 | 1 | 0 | 0 |
| UGC0007608 | Clust-UGC0002483 | 6029 | 6 | 1 | 0 | 1 |
| UGC0009742 | Clust-UGC0002483 | 1738 | 1 | 0 | 0 | 0 |
| UGC0007039 | Clust-UGC0002483 | 2294 | 2 | 1 | 0 | 1 |
| UGC0012661 | Clust-UGC0002483 | 11766 | 26 | 1 | 0 | 1 |
| UGC0006740 | Clust-UGC0002483 | 504 | 1 | 0 | 0 | 0 |
| UGC0006403 | Clust-UGC0002483 | 6202 | 3 | 1 | 0 | 0 |
| UGC0010813 | Clust-UGC0002483 | 4211 | 5 | 1 | 0 | 1 |
| UGC0009427 | Clust-UGC0002483 | 4140 | 1 | 0 | 0 | 0 |

| | | | | | | |
|---|---|---|---|---|---|---|
| UGC0008920 | Clust-UGC0002483 | 2297 | 2 | 1 | 0 | 0 |
| UGC0006729 | Clust-UGC0002483 | 3436 | 2 | 1 | 0 | 0 |
| UGC0012704 | Clust-UGC0002483 | 5560 | 3 | 1 | 0 | 1 |
| UGC0003374 | Clust-UGC0002483 | 9648 | 15 | 1 | 0 | 1 |
| UGC0012918 | Clust-UGC0002483 | 1227 | 2 | 1 | 0 | 0 |
| UGC0004970 | Clust-UGC0002483 | 9207 | 8 | 1 | 0 | 1 |
| UGC0009151 | Clust-UGC0002483 | 8273 | 9 | 1 | 0 | 1 |
| UGC0012585 | Clust-UGC0002483 | 6746 | 8 | 1 | 0 | 1 |
| UGC0008394 | Clust-UGC0002483 | 3210 | 2 | 1 | 0 | 0 |
| UGC0009635 | Clust-UGC0002483 | 2490 | 2 | 1 | 0 | 0 |
| UGC0007744 | Clust-UGC0002483 | 8958 | 13 | 1 | 0 | 1 |
| UGC0006681 | Clust-UGC0002483 | 1852 | 2 | 1 | 0 | 1 |
| UGC0007085 | Clust-UGC0002483 | 541 | 1 | 0 | 0 | 0 |
| UGC0013130 | Clust-UGC0002483 | 76 | 1 | 0 | 0 | 0 |
| UGC0005597 | Clust-UGC0002485 | 3787 | 4 | 1 | 0 | 1 |
| UGC0008932 | Clust-UGC0002485 | 1024 | 2 | 1 | 1 | 0 |
| UGC0005092 | Clust-UGC0002485 | 556 | 1 | 0 | 0 | 0 |
| UGC0007591 | Clust-UGC0002485 | 4210 | 2 | 1 | 0 | 0 |
| UGC0007287 | Clust-UGC0002485 | 8172 | 6 | 1 | 0 | 1 |
| UGC0006665 | Clust-UGC0002485 | 7866 | 14 | 1 | 0 | 1 |
| UGC0006911 | Clust-UGC0002485 | 1712 | 2 | 1 | 0 | 0 |
| UGC0005030 | Clust-UGC0002485 | 7459 | 6 | 1 | 0 | 1 |
| UGC0004639 | Clust-UGC0002485 | 7871 | 13 | 1 | 1 | 1 |
| UGC0006676 | Clust-UGC0002485 | 3727 | 6 | 1 | 0 | 0 |
| UGC0002485 | Clust-UGC0002485 | 757 | 2 | 1 | 0 | 0 |
| UGC0010119 | Clust-UGC0002485 | 4053 | 3 | 1 | 0 | 0 |
| UGC0010620 | Clust-UGC0002485 | 101 | 1 | 0 | 0 | 0 |
| UGC0010606 | Clust-UGC0002485 | 2392 | 1 | 0 | 0 | 0 |
| UGC0008281 | Clust-UGC0002485 | 875 | 3 | 1 | 0 | 0 |
| UGC0005346 | Clust-UGC0002485 | 2796 | 4 | 1 | 0 | 0 |
| UGC0005060 | Clust-UGC0002485 | 4076 | 6 | 1 | 0 | 1 |
| UGC0005061 | Clust-UGC0002485 | 2352 | 4 | 1 | 1 | 0 |
| UGC0005601 | Clust-UGC0002485 | 2559 | 5 | 1 | 0 | 1 |
| UGC0011898 | Clust-UGC0002485 | 127 | 1 | 0 | 0 | 0 |
| UGC0012858 | Clust-UGC0002485 | 6607 | 5 | 1 | 0 | 0 |
| UGC0004998 | Clust-UGC0002485 | 5446 | 12 | 1 | 0 | 1 |
| UGC0006784 | Clust-UGC0002485 | 4507 | 8 | 1 | 0 | 1 |
| UGC0009482 | Clust-UGC0002485 | 775 | 1 | 0 | 0 | 0 |
| UGC0012588 | Clust-UGC0002485 | 6206 | 15 | 1 | 0 | 1 |
| UGC0005723 | Clust-UGC0002485 | 5914 | 12 | 1 | 0 | 1 |
| UGC0005117 | Clust-UGC0002485 | 2875 | 2 | 1 | 0 | 0 |
| UGC0006642 | Clust-UGC0002485 | 1106 | 1 | 0 | 0 | 0 |
| UGC0004877 | Clust-UGC0002485 | 91 | 1 | 0 | 0 | 0 |
| UGC0008825 | Clust-UGC0002485 | 3739 | 7 | 1 | 0 | 1 |
| UGC0007224 | Clust-UGC0002485 | 8003 | 12 | 1 | 0 | 1 |
| UGC0002508 | Clust-UGC0002486 | 4461 | 2 | 1 | 0 | 1 |
| UGC0002486 | Clust-UGC0002486 | 2313 | 2 | 1 | 0 | 1 |
| UGC0008507 | Clust-UGC0002486 | 2038 | 2 | 1 | 0 | 0 |
| UGC0002492 | Clust-UGC0002492 | 6802 | 15 | 1 | 0 | 1 |
| UGC0003534 | Clust-UGC0002493 | 5136 | 8 | 1 | 0 | 1 |
| UGC0005566 | Clust-UGC0002493 | 1605 | 2 | 1 | 0 | 1 |
| UGC0004829 | Clust-UGC0002493 | 5777 | 3 | 1 | 0 | 1 |
| UGC0002493 | Clust-UGC0002493 | 851 | 1 | 0 | 0 | 0 |
| UGC0002497 | Clust-UGC0002497 | 397 | 1 | 0 | 0 | 0 |
| UGC0006712 | Clust-UGC0002507 | 1663 | 1 | 0 | 0 | 0 |
| UGC0002507 | Clust-UGC0002507 | 558 | 1 | 0 | 0 | 0 |
| UGC0002556 | Clust-UGC0002507 | 476 | 1 | 0 | 0 | 0 |
| UGC0006172 | Clust-UGC0002507 | 1385 | 1 | 0 | 0 | 0 |
| UGC0013282 | Clust-UGC0002507 | 661 | 1 | 0 | 0 | 0 |
| UGC0004839 | Clust-UGC0002514 | 1311 | 1 | 0 | 0 | 0 |
| UGC0012969 | Clust-UGC0002514 | 8031 | 6 | 1 | 0 | 1 |
| UGC0012555 | Clust-UGC0002514 | 3011 | 3 | 1 | 0 | 1 |
| UGC0012183 | Clust-UGC0002514 | 6604 | 7 | 1 | 0 | 1 |
| UGC0004496 | Clust-UGC0002514 | 8703 | 5 | 1 | 1 | 1 |
| UGC0010270 | Clust-UGC0002514 | 1130 | 1 | 0 | 0 | 0 |
| UGC0012874 | Clust-UGC0002514 | 11513 | 9 | 1 | 0 | 1 |
| UGC0005839 | Clust-UGC0002514 | 5407 | 2 | 1 | 0 | 1 |
| UGC0005028 | Clust-UGC0002514 | 2522 | 2 | 1 | 0 | 1 |
| UGC0010835 | Clust-UGC0002514 | 1379 | 1 | 0 | 0 | 0 |
| UGC0010761 | Clust-UGC0002514 | 5928 | 8 | 1 | 0 | 1 |
| UGC0002514 | Clust-UGC0002514 | 351 | 1 | 0 | 0 | 0 |
| UGC0003417 | Clust-UGC0002514 | 766 | 1 | 0 | 0 | 0 |
| UGC0006379 | Clust-UGC0002514 | 6340 | 5 | 1 | 0 | 1 |
| UGC0002515 | Clust-UGC0002515 | 2620 | 2 | 1 | 0 | 1 |
| UGC0006055 | Clust-UGC0002516 | 1979 | 1 | 0 | 0 | 0 |
| UGC0010613 | Clust-UGC0002516 | 1181 | 1 | 0 | 0 | 0 |
| UGC0007651 | Clust-UGC0002516 | 6152 | 12 | 1 | 0 | 1 |
| UGC0012321 | Clust-UGC0002516 | 3305 | 1 | 0 | 0 | 0 |
| UGC0007630 | Clust-UGC0002516 | 5742 | 5 | 1 | 0 | 1 |

| | | | | | | |
|---|---|---|---|---|---|---|
| UGC0007719 | Clust-UGC0002516 | 6327 | 9 | 1 | 0 | 1 |
| UGC0010110 | Clust-UGC0002516 | 690 | 1 | 0 | 0 | 0 |
| UGC0010582 | Clust-UGC0002516 | 3344 | 2 | 1 | 0 | 1 |
| UGC0010624 | Clust-UGC0002516 | 529 | 1 | 0 | 0 | 0 |
| UGC0004809 | Clust-UGC0002516 | 11737 | 79 | 1 | 0 | 1 |
| UGC0007075 | Clust-UGC0002516 | 859 | 1 | 0 | 0 | 0 |
| UGC0005043 | Clust-UGC0002516 | 1273 | 1 | 0 | 0 | 0 |
| UGC0005025 | Clust-UGC0002516 | 7747 | 15 | 1 | 0 | 1 |
| UGC0009099 | Clust-UGC0002516 | 844 | 2 | 1 | 0 | 1 |
| UGC0002516 | Clust-UGC0002516 | 2998 | 2 | 1 | 0 | 1 |
| UGC0013308 | Clust-UGC0002516 | 706 | 1 | 0 | 0 | 0 |
| UGC0005855 | Clust-UGC0002516 | 1248 | 1 | 0 | 0 | 0 |
| UGC0006374 | Clust-UGC0002516 | 6610 | 11 | 1 | 0 | 1 |
| UGC0008439 | Clust-UGC0002517 | 914 | 1 | 0 | 0 | 0 |
| UGC0004737 | Clust-UGC0002517 | 4600 | 2 | 1 | 0 | 0 |
| UGC0002517 | Clust-UGC0002517 | 5744 | 3 | 1 | 0 | 1 |
| UGC0010634 | Clust-UGC0002518 | 1606 | 1 | 0 | 0 | 0 |
| UGC0012635 | Clust-UGC0002518 | 1117 | 3 | 1 | 1 | 0 |
| UGC0008578 | Clust-UGC0002518 | 785 | 1 | 0 | 0 | 0 |
| UGC0008492 | Clust-UGC0002518 | 830 | 1 | 0 | 0 | 0 |
| UGC0008479 | Clust-UGC0002518 | 1041 | 2 | 1 | 0 | 0 |
| UGC0012876 | Clust-UGC0002518 | 8798 | 5 | 1 | 0 | 1 |
| UGC0008178 | Clust-UGC0002518 | 358 | 1 | 0 | 0 | 0 |
| UGC0002518 | Clust-UGC0002518 | 4534 | 1 | 0 | 0 | 0 |
| UGC0002519 | Clust-UGC0002519 | 3751 | 2 | 1 | 0 | 1 |
| UGC0007636 | Clust-UGC0002521 | 3440 | 2 | 1 | 0 | 1 |
| UGC0002521 | Clust-UGC0002521 | 1142 | 1 | 0 | 0 | 0 |
| UGC0008637 | Clust-UGC0002521 | 1665 | 1 | 0 | 0 | 0 |
| UGC0006504 | Clust-UGC0002521 | 1476 | 1 | 0 | 0 | 0 |
| UGC0004633 | Clust-UGC0002521 | 6566 | 5 | 1 | 1 | 1 |
| UGC0004672 | Clust-UGC0002521 | 3353 | 3 | 1 | 0 | 1 |
| UGC0004089 | Clust-UGC0002521 | 586 | 1 | 0 | 0 | 0 |
| UGC0004088 | Clust-UGC0002521 | 430 | 1 | 0 | 0 | 0 |
| UGC0004476 | Clust-UGC0002521 | 4042 | 3 | 1 | 0 | 1 |
| UGC0007054 | Clust-UGC0002521 | 2342 | 2 | 1 | 0 | 1 |
| UGC0002980 | Clust-UGC0002521 | 2954 | 1 | 0 | 0 | 0 |
| UGC0008471 | Clust-UGC0002521 | 842 | 1 | 0 | 0 | 0 |
| UGC0003548 | Clust-UGC0002521 | 3298 | 1 | 0 | 0 | 0 |
| UGC0004504 | Clust-UGC0002521 | 2381 | 1 | 0 | 0 | 0 |
| UGC0004548 | Clust-UGC0002521 | 4772 | 2 | 1 | 1 | 0 |
| UGC0006119 | Clust-UGC0002521 | 1110 | 1 | 0 | 0 | 0 |
| UGC0006299 | Clust-UGC0002533 | 3871 | 8 | 1 | 0 | 1 |
| UGC0005820 | Clust-UGC0002533 | 5549 | 6 | 1 | 1 | 1 |
| UGC0002533 | Clust-UGC0002533 | 7608 | 19 | 1 | 0 | 1 |
| UGC0011684 | Clust-UGC0002533 | 5139 | 6 | 1 | 0 | 1 |
| UGC0004685 | Clust-UGC0002533 | 14470 | 115 | 1 | 0 | 1 |
| UGC0013385 | Clust-UGC0002535 | 4694 | 5 | 1 | 0 | 1 |
| UGC0010071 | Clust-UGC0002535 | 3937 | 5 | 1 | 0 | 1 |
| UGC0002535 | Clust-UGC0002535 | 1402 | 1 | 0 | 0 | 0 |
| UGC0004566 | Clust-UGC0002535 | 177 | 1 | 0 | 0 | 0 |
| UGC0005851 | Clust-UGC0002535 | 2516 | 1 | 0 | 0 | 0 |
| UGC0007597 | Clust-UGC0002536 | 2324 | 2 | 1 | 0 | 0 |
| UGC0006857 | Clust-UGC0002536 | 3620 | 1 | 0 | 0 | 0 |
| UGC0008437 | Clust-UGC0002536 | 773 | 1 | 0 | 0 | 0 |
| UGC0004650 | Clust-UGC0002536 | 5778 | 2 | 1 | 0 | 1 |
| UGC0008201 | Clust-UGC0002536 | 5121 | 2 | 1 | 0 | 1 |
| UGC0008835 | Clust-UGC0002536 | 2819 | 1 | 0 | 0 | 0 |
| UGC0003327 | Clust-UGC0002536 | 2244 | 1 | 0 | 0 | 0 |
| UGC0009535 | Clust-UGC0002536 | 1197 | 1 | 0 | 0 | 0 |
| UGC0011641 | Clust-UGC0002536 | 861 | 2 | 1 | 0 | 0 |
| UGC0009445 | Clust-UGC0002536 | 2463 | 1 | 0 | 0 | 0 |
| UGC0012894 | Clust-UGC0002536 | 10205 | 6 | 1 | 1 | 1 |
| UGC0011017 | Clust-UGC0002536 | 3314 | 3 | 1 | 0 | 0 |
| UGC0002536 | Clust-UGC0002536 | 254 | 1 | 0 | 0 | 0 |
| UGC0003375 | Clust-UGC0002536 | 8938 | 7 | 1 | 0 | 1 |
| UGC0008597 | Clust-UGC0002536 | 3687 | 1 | 0 | 0 | 0 |
| UGC0003250 | Clust-UGC0002536 | 791 | 1 | 0 | 0 | 0 |
| UGC0002537 | Clust-UGC0002537 | 1884 | 1 | 0 | 0 | 0 |
| UGC0008676 | Clust-UGC0002537 | 2190 | 1 | 0 | 0 | 0 |
| UGC0002540 | Clust-UGC0002540 | 5074 | 7 | 1 | 0 | 1 |
| UGC0010203 | Clust-UGC0002542 | 962 | 2 | 1 | 0 | 1 |
| UGC0009479 | Clust-UGC0002542 | 833 | 1 | 0 | 0 | 0 |
| UGC0012886 | Clust-UGC0002542 | 5335 | 6 | 1 | 0 | 1 |
| UGC0011807 | Clust-UGC0002542 | 8532 | 31 | 1 | 0 | 1 |
| UGC0005526 | Clust-UGC0002542 | 837 | 1 | 0 | 0 | 0 |
| UGC0006960 | Clust-UGC0002542 | 3846 | 2 | 1 | 0 | 1 |
| UGC0002542 | Clust-UGC0002542 | 2982 | 2 | 1 | 0 | 1 |
| UGC0004470 | Clust-UGC0002542 | 3657 | 4 | 1 | 0 | 1 |
| UGC0006570 | Clust-UGC0002542 | 1982 | 2 | 1 | 0 | 1 |

| | | | | | | |
|---|---|---|---|---|---|---|
| UGC0008224 | Clust-UGC0002542 | 1135 | 1 | 0 | 0 | 0 |
| UGC0010622 | Clust-UGC0002542 | 2093 | 1 | 0 | 0 | 0 |
| UGC0006785 | Clust-UGC0002542 | 5141 | 3 | 1 | 1 | 1 |
| UGC0012954 | Clust-UGC0002542 | 1429 | 2 | 1 | 1 | 0 |
| UGC0013345 | Clust-UGC0002542 | 5272 | 6 | 1 | 0 | 1 |
| UGC0007729 | Clust-UGC0002542 | 4577 | 4 | 1 | 0 | 1 |
| UGC0003253 | Clust-UGC0002542 | 4097 | 1 | 0 | 0 | 0 |
| UGC0002544 | Clust-UGC0002544 | 895 | 1 | 0 | 0 | 0 |
| UGC0005829 | Clust-UGC0002544 | 3446 | 5 | 1 | 0 | 0 |
| UGC0008619 | Clust-UGC0002552 | 2149 | 5 | 1 | 0 | 1 |
| UGC0010639 | Clust-UGC0002552 | 1321 | 2 | 1 | 0 | 1 |
| UGC0005398 | Clust-UGC0002552 | 2380 | 2 | 1 | 0 | 0 |
| UGC0008703 | Clust-UGC0002552 | 1044 | 2 | 1 | 1 | 0 |
| UGC0010143 | Clust-UGC0002552 | 5171 | 7 | 1 | 0 | 1 |
| UGC0008023 | Clust-UGC0002552 | 2469 | 2 | 1 | 0 | 0 |
| UGC0007698 | Clust-UGC0002552 | 4424 | 3 | 1 | 0 | 0 |
| UGC0008402 | Clust-UGC0002552 | 978 | 2 | 1 | 0 | 0 |
| UGC0007911 | Clust-UGC0002552 | 6048 | 16 | 1 | 0 | 1 |
| UGC0008897 | Clust-UGC0002552 | 2351 | 3 | 1 | 0 | 0 |
| UGC0007996 | Clust-UGC0002552 | 2140 | 3 | 1 | 0 | 1 |
| UGC0010574 | Clust-UGC0002552 | 1694 | 3 | 1 | 0 | 1 |
| UGC0008017 | Clust-UGC0002552 | 56 | 1 | 0 | 0 | 0 |
| UGC0008461 | Clust-UGC0002552 | 1981 | 2 | 1 | 0 | 0 |
| UGC0008589 | Clust-UGC0002552 | 3140 | 6 | 1 | 0 | 1 |
| UGC0008584 | Clust-UGC0002552 | 5160 | 5 | 1 | 0 | 1 |
| UGC0002976 | Clust-UGC0002552 | 1667 | 5 | 1 | 0 | 0 |
| UGC0006391 | Clust-UGC0002552 | 7726 | 17 | 1 | 1 | 1 |
| UGC0008517 | Clust-UGC0002552 | 812 | 2 | 1 | 0 | 0 |
| UGC0002552 | Clust-UGC0002552 | 551 | 1 | 0 | 0 | 0 |
| UGC0008214 | Clust-UGC0002552 | 3916 | 7 | 1 | 0 | 1 |
| UGC0008824 | Clust-UGC0002552 | 2126 | 4 | 1 | 0 | 0 |
| UGC0008827 | Clust-UGC0002552 | 1535 | 2 | 1 | 0 | 0 |
| UGC0003278 | Clust-UGC0002552 | 2531 | 2 | 1 | 0 | 0 |
| UGC0007763 | Clust-UGC0002552 | 1056 | 2 | 1 | 0 | 1 |
| UGC0008335 | Clust-UGC0002552 | 928 | 1 | 0 | 0 | 0 |
| UGC0002557 | Clust-UGC0002557 | 647 | 1 | 0 | 0 | 0 |
| UGC0010461 | Clust-UGC0002560 | 4239 | 12 | 1 | 0 | 1 |
| UGC0002560 | Clust-UGC0002560 | 663 | 1 | 0 | 0 | 0 |
| UGC0005632 | Clust-UGC0002560 | 6311 | 10 | 1 | 0 | 1 |
| UGC0004969 | Clust-UGC0002560 | 6675 | 8 | 1 | 0 | 1 |
| UGC0010738 | Clust-UGC0002560 | 373 | 3 | 1 | 0 | 0 |
| UGC0012062 | Clust-UGC0002560 | 3435 | 2 | 1 | 0 | 1 |
| UGC0011973 | Clust-UGC0002560 | 3661 | 1 | 0 | 0 | 0 |
| UGC0012702 | Clust-UGC0002560 | 4244 | 11 | 1 | 0 | 1 |
| UGC0009987 | Clust-UGC0002560 | 1955 | 4 | 1 | 0 | 1 |
| UGC0009097 | Clust-UGC0002560 | 3529 | 10 | 1 | 0 | 1 |
| UGC0013305 | Clust-UGC0002560 | 2952 | 6 | 1 | 0 | 1 |
| UGC0007209 | Clust-UGC0002564 | 895 | 1 | 0 | 0 | 0 |
| UGC0002564 | Clust-UGC0002564 | 624 | 1 | 0 | 0 | 0 |
| UGC0007634 | Clust-UGC0002564 | 7720 | 24 | 1 | 0 | 1 |
| UGC0006918 | Clust-UGC0002564 | 1622 | 2 | 1 | 0 | 0 |
| UGC0008327 | Clust-UGC0002564 | 906 | 1 | 0 | 0 | 0 |
| UGC0008283 | Clust-UGC0002564 | 702 | 1 | 0 | 0 | 0 |
| UGC0005892 | Clust-UGC0002952 | 6587 | 13 | 1 | 0 | 1 |
| UGC0012921 | Clust-UGC0002952 | 121 | 1 | 0 | 0 | 0 |
| UGC0009666 | Clust-UGC0002952 | 2418 | 2 | 1 | 0 | 1 |
| UGC0006325 | Clust-UGC0002952 | 1652 | 2 | 1 | 0 | 1 |
| UGC0010538 | Clust-UGC0002952 | 1931 | 1 | 0 | 0 | 0 |
| UGC0007112 | Clust-UGC0002952 | 2296 | 1 | 0 | 0 | 0 |
| UGC0002952 | Clust-UGC0002952 | 98 | 1 | 0 | 0 | 0 |
| UGC0002953 | Clust-UGC0002953 | 425 | 1 | 0 | 0 | 0 |
| UGC0008482 | Clust-UGC0002954 | 200 | 1 | 0 | 0 | 0 |
| UGC0003516 | Clust-UGC0002954 | 2269 | 5 | 1 | 0 | 1 |
| UGC0012751 | Clust-UGC0002954 | 2166 | 5 | 1 | 0 | 1 |
| UGC0005126 | Clust-UGC0002954 | 2420 | 7 | 1 | 0 | 1 |
| UGC0012330 | Clust-UGC0002954 | 6456 | 23 | 1 | 0 | 1 |
| UGC0008892 | Clust-UGC0002954 | 2741 | 7 | 1 | 0 | 1 |
| UGC0003018 | Clust-UGC0002954 | 1466 | 4 | 1 | 0 | 1 |
| UGC0012789 | Clust-UGC0002954 | 1047 | 1 | 0 | 0 | 0 |
| UGC0008733 | Clust-UGC0002954 | 872 | 4 | 1 | 0 | 1 |
| UGC0002954 | Clust-UGC0002954 | 3288 | 3 | 1 | 0 | 1 |
| UGC0008393 | Clust-UGC0002954 | 387 | 1 | 0 | 0 | 0 |
| UGC0011610 | Clust-UGC0002958 | 1501 | 3 | 1 | 0 | 0 |
| UGC0008470 | Clust-UGC0002958 | 2484 | 1 | 0 | 0 | 0 |
| UGC0002958 | Clust-UGC0002958 | 968 | 1 | 0 | 0 | 0 |
| UGC0008957 | Clust-UGC0002959 | 223 | 1 | 0 | 0 | 0 |
| UGC0011855 | Clust-UGC0002959 | 1630 | 4 | 1 | 0 | 0 |
| UGC0002959 | Clust-UGC0002959 | 1366 | 2 | 1 | 0 | 0 |
| UGC0005093 | Clust-UGC0002967 | 2959 | 10 | 1 | 0 | 1 |

| | | | | | | |
|---|---|---|---|---|---|---|
| UGC0002967 | Clust-UGC0002967 | 764 | 1 | 0 | 0 | 0 |
| UGC0009146 | Clust-UGC0002967 | 6767 | 17 | 1 | 0 | 1 |
| UGC0005113 | Clust-UGC0002967 | 3153 | 5 | 1 | 0 | 0 |
| UGC0009150 | Clust-UGC0002967 | 1739 | 5 | 1 | 0 | 0 |
| UGC0004831 | Clust-UGC0002968 | 5931 | 13 | 1 | 0 | 1 |
| UGC0008480 | Clust-UGC0002968 | 591 | 1 | 0 | 0 | 0 |
| UGC0006503 | Clust-UGC0002968 | 1481 | 2 | 1 | 0 | 1 |
| UGC0007243 | Clust-UGC0002968 | 3130 | 2 | 1 | 0 | 0 |
| UGC0004742 | Clust-UGC0002968 | 97 | 1 | 0 | 0 | 0 |
| UGC0005091 | Clust-UGC0002968 | 376 | 1 | 0 | 0 | 0 |
| UGC0006845 | Clust-UGC0002968 | 1314 | 2 | 1 | 0 | 1 |
| UGC0012130 | Clust-UGC0002968 | 917 | 1 | 0 | 0 | 0 |
| UGC0006486 | Clust-UGC0002968 | 1826 | 1 | 0 | 0 | 0 |
| UGC0008740 | Clust-UGC0002968 | 395 | 1 | 0 | 0 | 0 |
| UGC0008963 | Clust-UGC0002968 | 369 | 3 | 1 | 0 | 1 |
| UGC0007168 | Clust-UGC0002968 | 2696 | 2 | 1 | 0 | 1 |
| UGC0006795 | Clust-UGC0002968 | 5726 | 7 | 1 | 0 | 1 |
| UGC0010541 | Clust-UGC0002968 | 7292 | 6 | 1 | 0 | 1 |
| UGC0006639 | Clust-UGC0002968 | 481 | 1 | 0 | 0 | 0 |
| UGC0005974 | Clust-UGC0002968 | 51 | 1 | 0 | 0 | 0 |
| UGC0005872 | Clust-UGC0002968 | 5019 | 6 | 1 | 0 | 1 |
| UGC0002968 | Clust-UGC0002968 | 751 | 1 | 0 | 0 | 0 |
| UGC0012755 | Clust-UGC0002968 | 2731 | 2 | 1 | 0 | 1 |
| UGC0011844 | Clust-UGC0002968 | 4987 | 4 | 1 | 0 | 1 |
| UGC0003013 | Clust-UGC0002968 | 2089 | 2 | 1 | 0 | 0 |
| UGC0008810 | Clust-UGC0002968 | 1498 | 6 | 1 | 0 | 1 |
| UGC0012371 | Clust-UGC0002968 | 7940 | 1 | 0 | 0 | 0 |
| UGC0008546 | Clust-UGC0002968 | 825 | 1 | 0 | 0 | 0 |
| UGC0008655 | Clust-UGC0002968 | 533 | 1 | 0 | 0 | 0 |
| UGC0004758 | Clust-UGC0002968 | 324 | 1 | 0 | 0 | 0 |
| UGC0007036 | Clust-UGC0002968 | 188 | 1 | 0 | 0 | 0 |
| UGC0013067 | Clust-UGC0002968 | 859 | 1 | 0 | 0 | 0 |
| UGC0006907 | Clust-UGC0002968 | 712 | 2 | 1 | 0 | 0 |
| UGC0010576 | Clust-UGC0002968 | 552 | 1 | 0 | 0 | 0 |
| UGC0011991 | Clust-UGC0002968 | 1049 | 1 | 0 | 0 | 0 |
| UGC0012955 | Clust-UGC0002968 | 278 | 1 | 0 | 0 | 0 |
| UGC0006726 | Clust-UGC0002968 | 597 | 1 | 0 | 0 | 0 |
| UGC0006359 | Clust-UGC0002968 | 103 | 1 | 0 | 0 | 0 |
| UGC0013342 | Clust-UGC0002968 | 5548 | 5 | 1 | 0 | 1 |
| UGC0012247 | Clust-UGC0002968 | 966 | 4 | 1 | 0 | 1 |
| UGC0003523 | Clust-UGC0002968 | 3012 | 3 | 1 | 0 | 1 |
| UGC0012747 | Clust-UGC0002968 | 838 | 1 | 0 | 0 | 0 |
| UGC0006669 | Clust-UGC0002968 | 1181 | 1 | 0 | 0 | 0 |
| UGC0010302 | Clust-UGC0002968 | 136 | 1 | 0 | 0 | 0 |
| UGC0002974 | Clust-UGC0002974 | 3807 | 2 | 1 | 0 | 1 |
| UGC0004837 | Clust-UGC0002978 | 14385 | 80 | 1 | 1 | 1 |
| UGC0012405 | Clust-UGC0002978 | 4797 | 8 | 1 | 0 | 1 |
| UGC0011170 | Clust-UGC0002978 | 2554 | 6 | 1 | 0 | 0 |
| UGC0008860 | Clust-UGC0002978 | 5084 | 10 | 1 | 0 | 1 |
| UGC0012369 | Clust-UGC0002978 | 4435 | 9 | 1 | 0 | 1 |
| UGC0012174 | Clust-UGC0002978 | 5106 | 7 | 1 | 0 | 1 |
| UGC0012171 | Clust-UGC0002978 | 5132 | 12 | 1 | 0 | 1 |
| UGC0012824 | Clust-UGC0002978 | 845 | 1 | 0 | 0 | 0 |
| UGC0012349 | Clust-UGC0002978 | 2810 | 5 | 1 | 0 | 1 |
| UGC0009526 | Clust-UGC0002978 | 1880 | 6 | 1 | 0 | 0 |
| UGC0010261 | Clust-UGC0002978 | 1638 | 4 | 1 | 0 | 0 |
| UGC0012807 | Clust-UGC0002978 | 2217 | 5 | 1 | 0 | 1 |
| UGC0012320 | Clust-UGC0002978 | 829 | 2 | 1 | 0 | 0 |
| UGC0011119 | Clust-UGC0002978 | 1155 | 2 | 1 | 1 | 0 |
| UGC0003708 | Clust-UGC0002978 | 1067 | 2 | 1 | 0 | 0 |
| UGC0011659 | Clust-UGC0002978 | 10513 | 35 | 1 | 0 | 1 |
| UGC0012256 | Clust-UGC0002978 | 650 | 1 | 0 | 0 | 0 |
| UGC0009737 | Clust-UGC0002978 | 6842 | 14 | 1 | 0 | 1 |
| UGC0010750 | Clust-UGC0002978 | 2981 | 9 | 1 | 0 | 0 |
| UGC0010800 | Clust-UGC0002978 | 2165 | 5 | 1 | 0 | 0 |
| UGC0011403 | Clust-UGC0002978 | 981 | 3 | 1 | 0 | 0 |
| UGC0012146 | Clust-UGC0002978 | 2328 | 6 | 1 | 0 | 1 |
| UGC0011982 | Clust-UGC0002978 | 2989 | 6 | 1 | 0 | 1 |
| UGC0009247 | Clust-UGC0002978 | 10484 | 30 | 1 | 0 | 1 |
| UGC0013270 | Clust-UGC0002978 | 3674 | 9 | 1 | 1 | 1 |
| UGC0010825 | Clust-UGC0002978 | 6349 | 13 | 1 | 0 | 1 |
| UGC0004534 | Clust-UGC0002978 | 3811 | 4 | 1 | 0 | 1 |
| UGC0010012 | Clust-UGC0002978 | 1805 | 3 | 1 | 0 | 0 |
| UGC0010849 | Clust-UGC0002978 | 3846 | 8 | 1 | 0 | 1 |
| UGC0009208 | Clust-UGC0002978 | 5383 | 9 | 1 | 0 | 1 |
| UGC0012382 | Clust-UGC0002978 | 8201 | 21 | 1 | 0 | 1 |
| UGC0012965 | Clust-UGC0002978 | 6231 | 15 | 1 | 1 | 1 |
| UGC0005694 | Clust-UGC0002978 | 353 | 1 | 0 | 0 | 0 |
| UGC0003591 | Clust-UGC0002978 | 4455 | 8 | 1 | 0 | 1 |

| | | | | | | |
|---|---|---|---|---|---|---|
| UGC0003594 | Clust-UGC0002978 | 8538 | 25 | 1 | 0 | 1 |
| UGC0003598 | Clust-UGC0002978 | 13358 | 72 | 1 | 0 | 1 |
| UGC0004575 | Clust-UGC0002978 | 8049 | 23 | 1 | 1 | 1 |
| UGC0009416 | Clust-UGC0002978 | 7538 | 18 | 1 | 0 | 1 |
| UGC0010888 | Clust-UGC0002978 | 4853 | 7 | 1 | 0 | 1 |
| UGC0008954 | Clust-UGC0002978 | 578 | 1 | 0 | 0 | 0 |
| UGC0012928 | Clust-UGC0002978 | 6590 | 14 | 1 | 0 | 1 |
| UGC0004804 | Clust-UGC0002978 | 22444 | 208 | 1 | 0 | 1 |
| UGC0003562 | Clust-UGC0002978 | 6301 | 9 | 1 | 0 | 1 |
| UGC0012000 | Clust-UGC0002978 | 1120 | 2 | 1 | 0 | 0 |
| UGC0011202 | Clust-UGC0002978 | 596 | 1 | 0 | 0 | 0 |
| UGC0009620 | Clust-UGC0002978 | 4643 | 7 | 1 | 0 | 1 |
| UGC0012271 | Clust-UGC0002978 | 3406 | 4 | 1 | 0 | 0 |
| UGC0003749 | Clust-UGC0002978 | 772 | 1 | 0 | 0 | 0 |
| UGC0009365 | Clust-UGC0002978 | 7634 | 15 | 1 | 0 | 1 |
| UGC0008998 | Clust-UGC0002978 | 2961 | 6 | 1 | 0 | 1 |
| UGC0010009 | Clust-UGC0002978 | 2279 | 6 | 1 | 0 | 0 |
| UGC0012048 | Clust-UGC0002978 | 637 | 1 | 0 | 0 | 0 |
| UGC0013296 | Clust-UGC0002978 | 156 | 1 | 0 | 0 | 0 |
| UGC0010965 | Clust-UGC0002978 | 2405 | 4 | 1 | 0 | 1 |
| UGC0010967 | Clust-UGC0002978 | 1848 | 3 | 1 | 0 | 0 |
| UGC0011188 | Clust-UGC0002978 | 2902 | 5 | 1 | 0 | 0 |
| UGC0008790 | Clust-UGC0002978 | 3991 | 8 | 1 | 0 | 1 |
| UGC0012377 | Clust-UGC0002978 | 8156 | 14 | 1 | 0 | 1 |
| UGC0011570 | Clust-UGC0002978 | 1450 | 2 | 1 | 0 | 0 |
| UGC0009518 | Clust-UGC0002978 | 1908 | 5 | 1 | 0 | 1 |
| UGC0011146 | Clust-UGC0002978 | 268 | 1 | 0 | 0 | 0 |
| UGC0009136 | Clust-UGC0002978 | 1339 | 5 | 1 | 0 | 0 |
| UGC0011556 | Clust-UGC0002978 | 4113 | 8 | 1 | 0 | 1 |
| UGC0013109 | Clust-UGC0002978 | 913 | 1 | 0 | 0 | 0 |
| UGC0003567 | Clust-UGC0002978 | 6010 | 15 | 1 | 0 | 1 |
| UGC0007077 | Clust-UGC0002978 | 4205 | 7 | 1 | 0 | 1 |
| UGC0003714 | Clust-UGC0002978 | 998 | 1 | 0 | 0 | 0 |
| UGC0011896 | Clust-UGC0002978 | 5874 | 12 | 1 | 0 | 1 |
| UGC0007606 | Clust-UGC0002978 | 6044 | 9 | 1 | 0 | 1 |
| UGC0011007 | Clust-UGC0002978 | 4392 | 8 | 1 | 0 | 1 |
| UGC0009743 | Clust-UGC0002978 | 4116 | 10 | 1 | 0 | 1 |
| UGC0011412 | Clust-UGC0002978 | 7485 | 18 | 1 | 0 | 1 |
| UGC0011997 | Clust-UGC0002978 | 4230 | 8 | 1 | 0 | 1 |
| UGC0009079 | Clust-UGC0002978 | 3884 | 6 | 1 | 0 | 1 |
| UGC0007435 | Clust-UGC0002978 | 667 | 1 | 0 | 0 | 0 |
| UGC0009441 | Clust-UGC0002978 | 2197 | 4 | 1 | 0 | 0 |
| UGC0012192 | Clust-UGC0002978 | 12841 | 70 | 1 | 0 | 1 |
| UGC0010836 | Clust-UGC0002978 | 503 | 1 | 0 | 0 | 0 |
| UGC0012404 | Clust-UGC0002978 | 3006 | 4 | 1 | 0 | 0 |
| UGC0011012 | Clust-UGC0002978 | 4692 | 5 | 1 | 0 | 0 |
| UGC0009278 | Clust-UGC0002978 | 4395 | 9 | 1 | 0 | 1 |
| UGC0004544 | Clust-UGC0002978 | 8174 | 19 | 1 | 0 | 1 |
| UGC0012391 | Clust-UGC0002978 | 6414 | 9 | 1 | 0 | 1 |
| UGC0010246 | Clust-UGC0002978 | 3035 | 6 | 1 | 0 | 1 |
| UGC0009428 | Clust-UGC0002978 | 3507 | 8 | 1 | 0 | 1 |
| UGC0010850 | Clust-UGC0002978 | 2771 | 7 | 1 | 0 | 1 |
| UGC0005512 | Clust-UGC0002978 | 1156 | 1 | 0 | 0 | 0 |
| UGC0009152 | Clust-UGC0002978 | 538 | 1 | 0 | 0 | 0 |
| UGC0009154 | Clust-UGC0002978 | 5944 | 8 | 1 | 0 | 1 |
| UGC0010879 | Clust-UGC0002978 | 3346 | 6 | 1 | 0 | 1 |
| UGC0002978 | Clust-UGC0002978 | 2482 | 5 | 1 | 0 | 0 |
| UGC0009230 | Clust-UGC0002978 | 14831 | 92 | 1 | 0 | 1 |
| UGC0009239 | Clust-UGC0002978 | 11342 | 55 | 1 | 0 | 1 |
| UGC0005725 | Clust-UGC0002978 | 6103 | 10 | 1 | 0 | 1 |
| UGC0012934 | Clust-UGC0002978 | 3543 | 9 | 1 | 0 | 1 |
| UGC0005535 | Clust-UGC0002978 | 4604 | 8 | 1 | 0 | 1 |
| UGC0009393 | Clust-UGC0002978 | 5118 | 13 | 1 | 0 | 1 |
| UGC0008636 | Clust-UGC0002978 | 5317 | 9 | 1 | 0 | 1 |
| UGC0012261 | Clust-UGC0002978 | 2966 | 6 | 1 | 0 | 1 |
| UGC0012263 | Clust-UGC0002978 | 1986 | 6 | 1 | 0 | 1 |
| UGC0011875 | Clust-UGC0002978 | 3416 | 5 | 1 | 0 | 1 |
| UGC0011878 | Clust-UGC0002978 | 2632 | 5 | 1 | 0 | 0 |
| UGC0011879 | Clust-UGC0002978 | 200 | 1 | 0 | 0 | 0 |
| UGC0009113 | Clust-UGC0002978 | 1325 | 3 | 1 | 0 | 0 |
| UGC0012207 | Clust-UGC0002978 | 7059 | 12 | 1 | 0 | 1 |
| UGC0009139 | Clust-UGC0002978 | 4351 | 10 | 1 | 0 | 1 |
| UGC0009651 | Clust-UGC0002978 | 3505 | 10 | 1 | 0 | 1 |
| UGC0006822 | Clust-UGC0002978 | 5255 | 9 | 1 | 0 | 1 |
| UGC0004686 | Clust-UGC0002978 | 12471 | 68 | 1 | 0 | 1 |
| UGC0007083 | Clust-UGC0002978 | 2712 | 5 | 1 | 0 | 0 |
| UGC0004144 | Clust-UGC0002978 | 3180 | 8 | 1 | 0 | 1 |
| UGC0012433 | Clust-UGC0002978 | 9107 | 29 | 1 | 0 | 1 |
| UGC0007691 | Clust-UGC0002985 | 4739 | 8 | 1 | 0 | 1 |

| | | | | | | |
|---|---|---|---|---|---|---|
| UGC0002985 | Clust-UGC0002985 | 361 | 1 | 0 | 0 | 0 |
| UGC0002988 | Clust-UGC0002985 | 2249 | 1 | 0 | 0 | 0 |
| UGC0007706 | Clust-UGC0002985 | 7255 | 16 | 1 | 0 | 1 |
| UGC0002990 | Clust-UGC0002990 | 1700 | 2 | 1 | 0 | 1 |
| UGC0012551 | Clust-UGC0002990 | 4926 | 3 | 1 | 0 | 0 |
| UGC0002994 | Clust-UGC0002994 | 3305 | 1 | 0 | 0 | 0 |
| UGC0008577 | Clust-UGC0002994 | 595 | 1 | 0 | 0 | 0 |
| UGC0008689 | Clust-UGC0002994 | 1858 | 1 | 0 | 0 | 0 |
| UGC0008356 | Clust-UGC0002994 | 1083 | 1 | 0 | 0 | 0 |
| UGC0010590 | Clust-UGC0002994 | 1315 | 1 | 0 | 0 | 0 |
| UGC0002996 | Clust-UGC0002996 | 1633 | 1 | 0 | 0 | 0 |
| UGC0003542 | Clust-UGC0002996 | 7854 | 3 | 1 | 0 | 1 |
| UGC0008440 | Clust-UGC0002996 | 1205 | 1 | 0 | 0 | 0 |
| UGC0005357 | Clust-UGC0002997 | 3645 | 3 | 1 | 0 | 1 |
| UGC0002997 | Clust-UGC0002997 | 1061 | 1 | 0 | 0 | 0 |
| UGC0010617 | Clust-UGC0002997 | 4197 | 2 | 1 | 0 | 1 |
| UGC0005586 | Clust-UGC0002997 | 4398 | 5 | 1 | 0 | 1 |
| UGC0007712 | Clust-UGC0002997 | 8578 | 31 | 1 | 0 | 1 |
| UGC0005520 | Clust-UGC0002997 | 4654 | 3 | 1 | 0 | 1 |
| UGC0005754 | Clust-UGC0002997 | 5777 | 7 | 1 | 0 | 1 |
| UGC0005006 | Clust-UGC0002997 | 9121 | 30 | 1 | 0 | 1 |
| UGC0010515 | Clust-UGC0002997 | 8395 | 22 | 1 | 0 | 1 |
| UGC0003002 | Clust-UGC0003002 | 3200 | 3 | 1 | 0 | 1 |
| UGC0005996 | Clust-UGC0003003 | 4163 | 3 | 1 | 0 | 1 |
| UGC0005864 | Clust-UGC0003003 | 845 | 1 | 0 | 0 | 0 |
| UGC0012937 | Clust-UGC0003003 | 6154 | 10 | 1 | 1 | 1 |
| UGC0003003 | Clust-UGC0003003 | 1481 | 1 | 0 | 0 | 0 |
| UGC0012971 | Clust-UGC0003003 | 3407 | 3 | 1 | 0 | 1 |
| UGC0012568 | Clust-UGC0003003 | 6321 | 9 | 1 | 0 | 1 |
| UGC0003004 | Clust-UGC0003004 | 868 | 2 | 1 | 0 | 1 |
| UGC0008527 | Clust-UGC0003004 | 1052 | 2 | 1 | 0 | 0 |
| UGC0005366 | Clust-UGC0003004 | 2562 | 3 | 1 | 0 | 0 |
| UGC0012290 | Clust-UGC0003004 | 814 | 1 | 0 | 0 | 0 |
| UGC0006068 | Clust-UGC0003004 | 1124 | 2 | 1 | 0 | 1 |
| UGC0006133 | Clust-UGC0003004 | 1645 | 3 | 1 | 0 | 1 |
| UGC0010126 | Clust-UGC0003004 | 2506 | 4 | 1 | 0 | 1 |
| UGC0008919 | Clust-UGC0003008 | 460 | 1 | 0 | 0 | 0 |
| UGC0003008 | Clust-UGC0003008 | 155 | 1 | 0 | 0 | 0 |
| UGC0013344 | Clust-UGC0003008 | 9467 | 47 | 1 | 0 | 1 |
| UGC0007945 | Clust-UGC0003008 | 3082 | 2 | 1 | 0 | 1 |
| UGC0005097 | Clust-UGC0003011 | 710 | 1 | 0 | 0 | 0 |
| UGC0003011 | Clust-UGC0003011 | 2631 | 4 | 1 | 0 | 0 |
| UGC0007294 | Clust-UGC0003011 | 2342 | 2 | 1 | 0 | 0 |
| UGC0013363 | Clust-UGC0003011 | 2857 | 4 | 1 | 1 | 0 |
| UGC0003017 | Clust-UGC0003017 | 1218 | 1 | 0 | 0 | 0 |
| UGC0008723 | Clust-UGC0003023 | 167 | 1 | 0 | 0 | 0 |
| UGC0003023 | Clust-UGC0003023 | 1612 | 1 | 0 | 0 | 0 |
| UGC0008801 | Clust-UGC0003023 | 3082 | 8 | 1 | 0 | 1 |
| UGC0007659 | Clust-UGC0003025 | 8728 | 6 | 1 | 0 | 1 |
| UGC0012798 | Clust-UGC0003025 | 1171 | 1 | 0 | 0 | 0 |
| UGC0004510 | Clust-UGC0003025 | 6281 | 3 | 1 | 0 | 1 |
| UGC0004538 | Clust-UGC0003025 | 8634 | 3 | 1 | 0 | 1 |
| UGC0005878 | Clust-UGC0003025 | 278 | 1 | 0 | 0 | 0 |
| UGC0007105 | Clust-UGC0003025 | 3023 | 1 | 0 | 0 | 0 |
| UGC0005109 | Clust-UGC0003025 | 3482 | 2 | 1 | 0 | 1 |
| UGC0011490 | Clust-UGC0003025 | 7046 | 1 | 0 | 0 | 0 |
| UGC0004587 | Clust-UGC0003025 | 1752 | 1 | 0 | 0 | 0 |
| UGC0007604 | Clust-UGC0003025 | 6313 | 4 | 1 | 0 | 1 |
| UGC0007371 | Clust-UGC0003025 | 4512 | 1 | 0 | 0 | 0 |
| UGC0003025 | Clust-UGC0003025 | 267 | 1 | 0 | 0 | 0 |
| UGC0003589 | Clust-UGC0003025 | 8547 | 4 | 1 | 0 | 1 |
| UGC0003557 | Clust-UGC0003025 | 4128 | 5 | 1 | 1 | 0 |
| UGC0006429 | Clust-UGC0003025 | 4547 | 3 | 1 | 0 | 1 |
| UGC0003297 | Clust-UGC0003025 | 287 | 1 | 0 | 0 | 0 |
| UGC0006461 | Clust-UGC0003027 | 1692 | 1 | 0 | 0 | 0 |
| UGC0012297 | Clust-UGC0003027 | 1165 | 1 | 0 | 0 | 0 |
| UGC0003039 | Clust-UGC0003027 | 639 | 2 | 1 | 1 | 0 |
| UGC0003027 | Clust-UGC0003027 | 1965 | 1 | 0 | 0 | 0 |
| UGC0003029 | Clust-UGC0003029 | 739 | 1 | 0 | 0 | 0 |
| UGC0003030 | Clust-UGC0003030 | 695 | 1 | 0 | 0 | 0 |
| UGC0007718 | Clust-UGC0003035 | 6254 | 16 | 1 | 0 | 1 |
| UGC0008195 | Clust-UGC0003035 | 1121 | 1 | 0 | 0 | 0 |
| UGC0004668 | Clust-UGC0003035 | 4258 | 2 | 1 | 0 | 0 |
| UGC0003035 | Clust-UGC0003035 | 1074 | 1 | 0 | 0 | 0 |
| UGC0008428 | Clust-UGC0003035 | 1183 | 1 | 0 | 0 | 0 |
| UGC0011925 | Clust-UGC0003036 | 9075 | 4 | 1 | 0 | 1 |
| UGC0009060 | Clust-UGC0003036 | 977 | 1 | 0 | 0 | 0 |
| UGC0003036 | Clust-UGC0003036 | 849 | 1 | 0 | 0 | 0 |
| UGC0003235 | Clust-UGC0003235 | 3204 | 2 | 1 | 0 | 1 |

| | | | | | | |
|---|---|---|---|---|---|---|
| UGC0006362 | Clust-UGC0003235 | 1868 | 1 | 0 | 0 | 0 |
| UGC0006065 | Clust-UGC0003235 | 2075 | 2 | 1 | 0 | 1 |
| UGC0004956 | Clust-UGC0003235 | 5575 | 8 | 1 | 0 | 1 |
| UGC0012762 | Clust-UGC0003235 | 1455 | 1 | 0 | 0 | 0 |
| UGC0012421 | Clust-UGC0003235 | 3725 | 4 | 1 | 0 | 1 |
| UGC0009546 | Clust-UGC0003236 | 4420 | 3 | 1 | 0 | 1 |
| UGC0003236 | Clust-UGC0003236 | 4002 | 3 | 1 | 0 | 1 |
| UGC0004988 | Clust-UGC0003236 | 7026 | 12 | 1 | 0 | 1 |
| UGC0008686 | Clust-UGC0003236 | 2074 | 1 | 0 | 0 | 0 |
| UGC0006872 | Clust-UGC0003236 | 4831 | 3 | 1 | 0 | 1 |
| UGC0010192 | Clust-UGC0003236 | 1170 | 2 | 1 | 0 | 1 |
| UGC0005385 | Clust-UGC0003236 | 823 | 1 | 0 | 0 | 0 |
| UGC0007500 | Clust-UGC0003236 | 4172 | 5 | 1 | 0 | 1 |
| UGC0012212 | Clust-UGC0003236 | 4469 | 6 | 1 | 0 | 1 |
| UGC0009348 | Clust-UGC0003236 | 2356 | 1 | 0 | 0 | 0 |
| UGC0007548 | Clust-UGC0003236 | 4603 | 1 | 0 | 0 | 0 |
| UGC0008798 | Clust-UGC0003236 | 150 | 1 | 0 | 0 | 0 |
| UGC0005414 | Clust-UGC0003236 | 1281 | 1 | 0 | 0 | 0 |
| UGC0007786 | Clust-UGC0003236 | 933 | 1 | 0 | 0 | 0 |
| UGC0008654 | Clust-UGC0003236 | 1789 | 2 | 1 | 0 | 1 |
| UGC0004484 | Clust-UGC0003236 | 4871 | 6 | 1 | 0 | 1 |
| UGC0013341 | Clust-UGC0003236 | 4502 | 6 | 1 | 0 | 1 |
| UGC0006865 | Clust-UGC0003236 | 4298 | 4 | 1 | 0 | 1 |
| UGC0008605 | Clust-UGC0003237 | 6496 | 6 | 1 | 0 | 1 |
| UGC0003237 | Clust-UGC0003237 | 2138 | 3 | 1 | 0 | 0 |
| UGC0009109 | Clust-UGC0003237 | 553 | 1 | 0 | 0 | 0 |
| UGC0009325 | Clust-UGC0003237 | 7056 | 15 | 1 | 0 | 1 |
| UGC0005004 | Clust-UGC0003237 | 6399 | 11 | 1 | 1 | 1 |
| UGC0003239 | Clust-UGC0003239 | 8350 | 31 | 1 | 0 | 1 |
| UGC0006131 | Clust-UGC0003239 | 2703 | 3 | 1 | 0 | 1 |
| UGC0004782 | Clust-UGC0003241 | 1072 | 1 | 0 | 0 | 0 |
| UGC0004749 | Clust-UGC0003241 | 215 | 1 | 0 | 0 | 0 |
| UGC0004729 | Clust-UGC0003241 | 1374 | 1 | 0 | 0 | 0 |
| UGC0004793 | Clust-UGC0003241 | 4258 | 5 | 1 | 0 | 1 |
| UGC0004794 | Clust-UGC0003241 | 3963 | 3 | 1 | 0 | 1 |
| UGC0003241 | Clust-UGC0003241 | 5635 | 12 | 1 | 0 | 1 |
| UGC0004785 | Clust-UGC0003241 | 746 | 1 | 0 | 0 | 0 |
| UGC0007672 | Clust-UGC0003243 | 11210 | 13 | 1 | 0 | 1 |
| UGC0006659 | Clust-UGC0003243 | 3200 | 5 | 1 | 0 | 1 |
| UGC0009749 | Clust-UGC0003243 | 3856 | 3 | 1 | 0 | 0 |
| UGC0006880 | Clust-UGC0003243 | 5593 | 4 | 1 | 0 | 1 |
| UGC0003243 | Clust-UGC0003243 | 7067 | 7 | 1 | 0 | 1 |
| UGC0010168 | Clust-UGC0003248 | 990 | 2 | 1 | 0 | 1 |
| UGC0009738 | Clust-UGC0003248 | 4144 | 1 | 0 | 0 | 0 |
| UGC0004769 | Clust-UGC0003248 | 4351 | 2 | 1 | 0 | 1 |
| UGC0007027 | Clust-UGC0003248 | 10009 | 62 | 1 | 0 | 1 |
| UGC0007402 | Clust-UGC0003248 | 1785 | 1 | 0 | 0 | 0 |
| UGC0004180 | Clust-UGC0003248 | 3903 | 1 | 0 | 0 | 0 |
| UGC0006775 | Clust-UGC0003248 | 9174 | 43 | 1 | 0 | 1 |
| UGC0007488 | Clust-UGC0003248 | 5155 | 3 | 1 | 0 | 1 |
| UGC0006818 | Clust-UGC0003248 | 4821 | 1 | 0 | 0 | 0 |
| UGC0013319 | Clust-UGC0003248 | 512 | 2 | 1 | 0 | 0 |
| UGC0004175 | Clust-UGC0003248 | 90 | 1 | 0 | 0 | 0 |
| UGC0010254 | Clust-UGC0003248 | 389 | 1 | 0 | 0 | 0 |
| UGC0010552 | Clust-UGC0003248 | 4846 | 5 | 1 | 0 | 1 |
| UGC0004485 | Clust-UGC0003248 | 7431 | 11 | 1 | 0 | 1 |
| UGC0005572 | Clust-UGC0003248 | 3374 | 3 | 1 | 0 | 1 |
| UGC0009219 | Clust-UGC0003248 | 15467 | 146 | 1 | 0 | 1 |
| UGC0009985 | Clust-UGC0003248 | 3661 | 3 | 1 | 0 | 1 |
| UGC0009153 | Clust-UGC0003248 | 2932 | 1 | 0 | 0 | 0 |
| UGC0003248 | Clust-UGC0003248 | 4373 | 3 | 1 | 0 | 1 |
| UGC0007080 | Clust-UGC0003248 | 2901 | 2 | 1 | 0 | 1 |
| UGC0010614 | Clust-UGC0003251 | 936 | 1 | 0 | 0 | 0 |
| UGC0006779 | Clust-UGC0003251 | 4734 | 5 | 1 | 0 | 1 |
| UGC0008979 | Clust-UGC0003251 | 1095 | 2 | 1 | 0 | 0 |
| UGC0007582 | Clust-UGC0003251 | 5741 | 5 | 1 | 0 | 1 |
| UGC0008427 | Clust-UGC0003251 | 750 | 2 | 1 | 0 | 1 |
| UGC0003251 | Clust-UGC0003251 | 2658 | 4 | 1 | 0 | 1 |
| UGC0008704 | Clust-UGC0003252 | 296 | 2 | 1 | 0 | 0 |
| UGC0007257 | Clust-UGC0003252 | 2700 | 8 | 1 | 0 | 1 |
| UGC0003252 | Clust-UGC0003252 | 8292 | 31 | 1 | 0 | 1 |
| UGC0003605 | Clust-UGC0003256 | 10661 | 46 | 1 | 0 | 1 |
| UGC0003256 | Clust-UGC0003256 | 4904 | 6 | 1 | 0 | 1 |
| UGC0003259 | Clust-UGC0003259 | 3199 | 3 | 1 | 0 | 1 |
| UGC0007922 | Clust-UGC0003262 | 3486 | 4 | 1 | 0 | 1 |
| UGC0006522 | Clust-UGC0003262 | 1171 | 1 | 0 | 0 | 0 |
| UGC0005998 | Clust-UGC0003262 | 3975 | 2 | 1 | 0 | 0 |
| UGC0011906 | Clust-UGC0003262 | 2574 | 2 | 1 | 0 | 1 |
| UGC0007008 | Clust-UGC0003262 | 4594 | 3 | 1 | 0 | 1 |

| | | | | | | |
|---|---|---|---|---|---|---|
| UGC0011407 | Clust-UGC0003262 | 1773 | 1 | 0 | 0 | 0 |
| UGC0006657 | Clust-UGC0003262 | 627 | 1 | 0 | 0 | 0 |
| UGC0008002 | Clust-UGC0003262 | 1765 | 2 | 1 | 0 | 0 |
| UGC0004559 | Clust-UGC0003262 | 7250 | 11 | 1 | 0 | 1 |
| UGC0011593 | Clust-UGC0003262 | 457 | 1 | 0 | 0 | 0 |
| UGC0003272 | Clust-UGC0003262 | 121 | 1 | 0 | 0 | 0 |
| UGC0012041 | Clust-UGC0003262 | 1819 | 1 | 0 | 0 | 0 |
| UGC0011957 | Clust-UGC0003262 | 2931 | 2 | 1 | 0 | 1 |
| UGC0008499 | Clust-UGC0003262 | 3630 | 3 | 1 | 0 | 1 |
| UGC0009133 | Clust-UGC0003262 | 3587 | 3 | 1 | 0 | 1 |
| UGC0004505 | Clust-UGC0003262 | 4990 | 2 | 1 | 0 | 1 |
| UGC0012722 | Clust-UGC0003262 | 5100 | 5 | 1 | 0 | 1 |
| UGC0004543 | Clust-UGC0003262 | 5766 | 5 | 1 | 0 | 1 |
| UGC0006725 | Clust-UGC0003262 | 953 | 2 | 1 | 0 | 0 |
| UGC0003584 | Clust-UGC0003262 | 10578 | 35 | 1 | 0 | 1 |
| UGC0012039 | Clust-UGC0003262 | 2820 | 3 | 1 | 0 | 1 |
| UGC0012606 | Clust-UGC0003262 | 9343 | 29 | 1 | 0 | 1 |
| UGC0006136 | Clust-UGC0003262 | 976 | 1 | 0 | 0 | 0 |
| UGC0003608 | Clust-UGC0003262 | 8589 | 24 | 1 | 0 | 1 |
| UGC0006111 | Clust-UGC0003262 | 732 | 1 | 0 | 0 | 0 |
| UGC0003262 | Clust-UGC0003262 | 5247 | 1 | 0 | 0 | 0 |
| UGC0006353 | Clust-UGC0003262 | 1912 | 2 | 1 | 0 | 1 |
| UGC0007222 | Clust-UGC0003262 | 5202 | 3 | 1 | 1 | 1 |
| UGC0004147 | Clust-UGC0003262 | 2005 | 1 | 0 | 0 | 0 |
| UGC0006525 | Clust-UGC0003273 | 3924 | 1 | 0 | 0 | 0 |
| UGC0012213 | Clust-UGC0003273 | 9191 | 6 | 1 | 0 | 1 |
| UGC0004479 | Clust-UGC0003273 | 9372 | 3 | 1 | 1 | 1 |
| UGC0003273 | Clust-UGC0003273 | 6804 | 5 | 1 | 0 | 0 |
| UGC0009567 | Clust-UGC0003274 | 232 | 1 | 0 | 0 | 0 |
| UGC0009007 | Clust-UGC0003274 | 3143 | 1 | 0 | 0 | 0 |
| UGC0006762 | Clust-UGC0003274 | 4180 | 3 | 1 | 0 | 1 |
| UGC0003740 | Clust-UGC0003274 | 437 | 1 | 0 | 0 | 0 |
| UGC0011616 | Clust-UGC0003274 | 463 | 1 | 0 | 0 | 0 |
| UGC0011005 | Clust-UGC0003274 | 3780 | 3 | 1 | 0 | 1 |
| UGC0008490 | Clust-UGC0003274 | 1178 | 1 | 0 | 0 | 0 |
| UGC0010023 | Clust-UGC0003274 | 428 | 1 | 0 | 0 | 0 |
| UGC0010550 | Clust-UGC0003274 | 5654 | 4 | 1 | 0 | 1 |
| UGC0003274 | Clust-UGC0003274 | 1322 | 1 | 0 | 0 | 0 |
| UGC0012030 | Clust-UGC0003274 | 2434 | 3 | 1 | 1 | 0 |
| UGC0012992 | Clust-UGC0003274 | 4097 | 3 | 1 | 1 | 1 |
| UGC0003385 | Clust-UGC0003282 | 10861 | 47 | 1 | 0 | 1 |
| UGC0003282 | Clust-UGC0003282 | 4437 | 5 | 1 | 0 | 1 |
| UGC0011892 | Clust-UGC0003282 | 16183 | 114 | 1 | 0 | 1 |
| UGC0006692 | Clust-UGC0003282 | 1571 | 1 | 0 | 0 | 0 |
| UGC0006500 | Clust-UGC0003282 | 2513 | 2 | 1 | 0 | 0 |
| UGC0010166 | Clust-UGC0003285 | 315 | 1 | 0 | 0 | 0 |
| UGC0010992 | Clust-UGC0003285 | 213 | 1 | 0 | 0 | 0 |
| UGC0005403 | Clust-UGC0003285 | 4333 | 1 | 0 | 0 | 0 |
| UGC0005587 | Clust-UGC0003285 | 2541 | 2 | 1 | 0 | 1 |
| UGC0006877 | Clust-UGC0003285 | 2581 | 2 | 1 | 0 | 1 |
| UGC0003285 | Clust-UGC0003285 | 5466 | 5 | 1 | 0 | 1 |
| UGC0007760 | Clust-UGC0003285 | 5402 | 9 | 1 | 0 | 1 |
| UGC0003289 | Clust-UGC0003289 | 1026 | 1 | 0 | 0 | 0 |
| UGC0003296 | Clust-UGC0003296 | 63 | 1 | 0 | 0 | 0 |
| UGC0003302 | Clust-UGC0003302 | 6034 | 34 | 1 | 0 | 1 |
| UGC0008411 | Clust-UGC0003302 | 1319 | 4 | 1 | 0 | 1 |
| UGC0008624 | Clust-UGC0003302 | 4641 | 22 | 1 | 0 | 1 |
| UGC0006816 | Clust-UGC0003302 | 7119 | 31 | 1 | 0 | 1 |
| UGC0007684 | Clust-UGC0003302 | 2702 | 6 | 1 | 0 | 1 |
| UGC0006766 | Clust-UGC0003304 | 3245 | 2 | 1 | 0 | 1 |
| UGC0004550 | Clust-UGC0003304 | 9501 | 30 | 1 | 0 | 1 |
| UGC0003304 | Clust-UGC0003304 | 3061 | 9 | 1 | 0 | 1 |
| UGC0004630 | Clust-UGC0003304 | 9508 | 28 | 1 | 0 | 1 |
| UGC0008977 | Clust-UGC0003304 | 2219 | 2 | 1 | 0 | 1 |
| UGC0006709 | Clust-UGC0003304 | 1205 | 3 | 1 | 0 | 1 |
| UGC0009603 | Clust-UGC0003304 | 485 | 1 | 0 | 0 | 0 |
| UGC0004834 | Clust-UGC0003310 | 7640 | 12 | 1 | 0 | 1 |
| UGC0005779 | Clust-UGC0003310 | 10107 | 24 | 1 | 0 | 1 |
| UGC0009977 | Clust-UGC0003310 | 4157 | 4 | 1 | 0 | 0 |
| UGC0004825 | Clust-UGC0003310 | 21734 | 81 | 1 | 0 | 1 |
| UGC0003310 | Clust-UGC0003310 | 5012 | 15 | 1 | 0 | 1 |
| UGC0010749 | Clust-UGC0003310 | 73 | 1 | 0 | 0 | 0 |
| UGC0010977 | Clust-UGC0003310 | 2144 | 2 | 1 | 0 | 0 |
| UGC0003313 | Clust-UGC0003313 | 7174 | 15 | 1 | 0 | 1 |
| UGC0003320 | Clust-UGC0003320 | 3013 | 1 | 0 | 0 | 0 |
| UGC0003332 | Clust-UGC0003332 | 2477 | 3 | 1 | 0 | 1 |
| UGC0011872 | Clust-UGC0003332 | 8211 | 27 | 1 | 0 | 1 |
| UGC0010482 | Clust-UGC0003339 | 4569 | 1 | 0 | 0 | 0 |
| UGC0003339 | Clust-UGC0003339 | 8093 | 7 | 1 | 0 | 1 |

| | | | | | | |
|---|---|---|---|---|---|---|
| UGC0004169 | Clust-UGC0003344 | 3885 | 12 | 1 | 0 | 1 |
| UGC0003344 | Clust-UGC0003344 | 15357 | 74 | 1 | 0 | 1 |
| UGC0012159 | Clust-UGC0003349 | 2692 | 3 | 1 | 0 | 1 |
| UGC0009179 | Clust-UGC0003349 | 4754 | 6 | 1 | 0 | 1 |
| UGC0006583 | Clust-UGC0003349 | 1175 | 1 | 0 | 0 | 0 |
| UGC0011429 | Clust-UGC0003349 | 3323 | 6 | 1 | 1 | 1 |
| UGC0013077 | Clust-UGC0003349 | 2672 | 1 | 0 | 0 | 0 |
| UGC0009752 | Clust-UGC0003349 | 557 | 1 | 0 | 0 | 0 |
| UGC0004985 | Clust-UGC0003349 | 2524 | 1 | 0 | 0 | 0 |
| UGC0007182 | Clust-UGC0003349 | 4981 | 4 | 1 | 0 | 1 |
| UGC0008966 | Clust-UGC0003349 | 2397 | 1 | 0 | 0 | 0 |
| UGC0005037 | Clust-UGC0003349 | 3152 | 1 | 0 | 0 | 0 |
| UGC0006651 | Clust-UGC0003349 | 117 | 1 | 0 | 0 | 0 |
| UGC0004813 | Clust-UGC0003349 | 18071 | 170 | 1 | 1 | 1 |
| UGC0003349 | Clust-UGC0003349 | 10999 | 58 | 1 | 0 | 1 |
| UGC0008535 | Clust-UGC0003349 | 855 | 1 | 0 | 0 | 0 |
| UGC0003757 | Clust-UGC0003349 | 389 | 1 | 0 | 0 | 0 |
| UGC0004963 | Clust-UGC0003349 | 6503 | 9 | 1 | 0 | 1 |
| UGC0007127 | Clust-UGC0003349 | 3533 | 1 | 0 | 0 | 0 |
| UGC0009084 | Clust-UGC0003349 | 2364 | 1 | 0 | 0 | 0 |
| UGC0009089 | Clust-UGC0003349 | 1062 | 1 | 0 | 0 | 0 |
| UGC0010501 | Clust-UGC0003349 | 2889 | 1 | 0 | 0 | 0 |
| UGC0007271 | Clust-UGC0003349 | 7712 | 23 | 1 | 0 | 1 |
| UGC0009608 | Clust-UGC0003349 | 1125 | 1 | 0 | 0 | 0 |
| UGC0012924 | Clust-UGC0003349 | 6241 | 10 | 1 | 0 | 1 |
| UGC0004805 | Clust-UGC0003349 | 16152 | 142 | 1 | 0 | 1 |
| UGC0005525 | Clust-UGC0003349 | 1678 | 1 | 0 | 0 | 0 |
| UGC0008974 | Clust-UGC0003349 | 1919 | 2 | 1 | 1 | 0 |
| UGC0010926 | Clust-UGC0003349 | 334 | 1 | 0 | 0 | 0 |
| UGC0009368 | Clust-UGC0003349 | 2316 | 1 | 0 | 0 | 0 |
| UGC0010130 | Clust-UGC0003349 | 20874 | 198 | 1 | 0 | 1 |
| UGC0005802 | Clust-UGC0003349 | 4336 | 4 | 1 | 0 | 1 |
| UGC0005980 | Clust-UGC0003349 | 795 | 1 | 0 | 0 | 0 |
| UGC0005664 | Clust-UGC0003349 | 2186 | 1 | 0 | 0 | 0 |
| UGC0011664 | Clust-UGC0003349 | 1925 | 4 | 1 | 0 | 1 |
| UGC0005065 | Clust-UGC0003349 | 1722 | 1 | 0 | 0 | 0 |
| UGC0009036 | Clust-UGC0003349 | 2261 | 3 | 1 | 1 | 1 |
| UGC0005045 | Clust-UGC0003349 | 4082 | 2 | 1 | 0 | 1 |
| UGC0010233 | Clust-UGC0003349 | 74 | 1 | 0 | 0 | 0 |
| UGC0011814 | Clust-UGC0003349 | 2747 | 1 | 0 | 0 | 0 |
| UGC0011609 | Clust-UGC0003349 | 1045 | 2 | 1 | 0 | 0 |
| UGC0007430 | Clust-UGC0003349 | 3123 | 2 | 1 | 1 | 0 |
| UGC0009275 | Clust-UGC0003349 | 580 | 1 | 0 | 0 | 0 |
| UGC0011812 | Clust-UGC0003349 | 2354 | 4 | 1 | 0 | 1 |
| UGC0008463 | Clust-UGC0003349 | 2114 | 1 | 0 | 0 | 0 |
| UGC0003395 | Clust-UGC0003349 | 482 | 1 | 0 | 0 | 0 |
| UGC0007399 | Clust-UGC0003349 | 723 | 1 | 0 | 0 | 0 |
| UGC0007534 | Clust-UGC0003349 | 3819 | 2 | 1 | 0 | 1 |
| UGC0005709 | Clust-UGC0003349 | 6356 | 7 | 1 | 0 | 1 |
| UGC0004095 | Clust-UGC0003349 | 1311 | 1 | 0 | 0 | 0 |
| UGC0009581 | Clust-UGC0003349 | 756 | 1 | 0 | 0 | 0 |
| UGC0009656 | Clust-UGC0003349 | 365 | 1 | 0 | 0 | 0 |
| UGC0010592 | Clust-UGC0003349 | 4594 | 2 | 1 | 0 | 1 |
| UGC0010100 | Clust-UGC0003349 | 3691 | 1 | 0 | 0 | 0 |
| UGC0008334 | Clust-UGC0003349 | 1464 | 2 | 1 | 0 | 1 |
| UGC0006489 | Clust-UGC0003353 | 783 | 1 | 0 | 0 | 0 |
| UGC0003353 | Clust-UGC0003353 | 12374 | 4 | 1 | 0 | 1 |
| UGC0003372 | Clust-UGC0003372 | 1040 | 2 | 1 | 0 | 0 |
| UGC0008585 | Clust-UGC0003376 | 2428 | 1 | 0 | 0 | 0 |
| UGC0003376 | Clust-UGC0003376 | 3186 | 2 | 1 | 0 | 0 |
| UGC0003383 | Clust-UGC0003383 | 8017 | 9 | 1 | 0 | 1 |
| UGC0011981 | Clust-UGC0003386 | 14847 | 48 | 1 | 0 | 1 |
| UGC0007444 | Clust-UGC0003386 | 2210 | 2 | 1 | 0 | 0 |
| UGC0011802 | Clust-UGC0003386 | 1458 | 2 | 1 | 0 | 0 |
| UGC0003386 | Clust-UGC0003386 | 22720 | 67 | 1 | 1 | 1 |
| UGC0012274 | Clust-UGC0003386 | 4259 | 3 | 1 | 0 | 1 |
| UGC0012615 | Clust-UGC0003386 | 6282 | 6 | 1 | 0 | 1 |
| UGC0011935 | Clust-UGC0003386 | 7772 | 3 | 1 | 0 | 1 |
| UGC0012295 | Clust-UGC0003386 | 2306 | 2 | 1 | 1 | 0 |
| UGC0009555 | Clust-UGC0003386 | 672 | 1 | 0 | 0 | 0 |
| UGC0004739 | Clust-UGC0003386 | 670 | 1 | 0 | 0 | 0 |
| UGC0012123 | Clust-UGC0003386 | 339 | 1 | 0 | 0 | 0 |
| UGC0007031 | Clust-UGC0003386 | 5372 | 3 | 1 | 0 | 1 |
| UGC0007410 | Clust-UGC0003386 | 7032 | 5 | 1 | 0 | 1 |
| UGC0006727 | Clust-UGC0003386 | 2045 | 2 | 1 | 0 | 1 |
| UGC0010853 | Clust-UGC0003386 | 1355 | 3 | 1 | 0 | 0 |
| UGC0007511 | Clust-UGC0003386 | 11551 | 8 | 1 | 0 | 1 |
| UGC0004709 | Clust-UGC0003386 | 32590 | 133 | 1 | 0 | 1 |
| UGC0009432 | Clust-UGC0003403 | 513 | 1 | 0 | 0 | 0 |

| | | | | | | |
|---|---|---|---|---|---|---|
| UGC0003403 | Clust-UGC0003403 | 1312 | 1 | 0 | 0 | 0 |
| UGC0005377 | Clust-UGC0003411 | 508 | 1 | 0 | 0 | 0 |
| UGC0007026 | Clust-UGC0003411 | 3330 | 1 | 0 | 0 | 0 |
| UGC0006770 | Clust-UGC0003411 | 4503 | 4 | 1 | 0 | 1 |
| UGC0005543 | Clust-UGC0003411 | 1133 | 1 | 0 | 0 | 0 |
| UGC0009999 | Clust-UGC0003411 | 1209 | 1 | 0 | 0 | 0 |
| UGC0009082 | Clust-UGC0003411 | 345 | 1 | 0 | 0 | 0 |
| UGC0007524 | Clust-UGC0003411 | 2714 | 1 | 0 | 0 | 0 |
| UGC0008975 | Clust-UGC0003411 | 1457 | 1 | 0 | 0 | 0 |
| UGC0008972 | Clust-UGC0003411 | 1133 | 1 | 0 | 0 | 0 |
| UGC0009103 | Clust-UGC0003411 | 641 | 1 | 0 | 0 | 0 |
| UGC0008992 | Clust-UGC0003411 | 3039 | 3 | 1 | 1 | 0 |
| UGC0009646 | Clust-UGC0003411 | 1538 | 1 | 0 | 0 | 0 |
| UGC0006555 | Clust-UGC0003411 | 919 | 1 | 0 | 0 | 0 |
| UGC0007667 | Clust-UGC0003411 | 5053 | 2 | 1 | 0 | 0 |
| UGC0006724 | Clust-UGC0003411 | 1068 | 1 | 0 | 0 | 0 |
| UGC0006591 | Clust-UGC0003411 | 447 | 1 | 0 | 0 | 0 |
| UGC0009076 | Clust-UGC0003411 | 1384 | 3 | 1 | 0 | 1 |
| UGC0010002 | Clust-UGC0003411 | 717 | 1 | 0 | 0 | 0 |
| UGC0009744 | Clust-UGC0003411 | 2265 | 1 | 0 | 0 | 0 |
| UGC0003411 | Clust-UGC0003411 | 2593 | 1 | 0 | 0 | 0 |
| UGC0007686 | Clust-UGC0003411 | 2441 | 2 | 1 | 0 | 1 |
| UGC0006625 | Clust-UGC0003411 | 1121 | 1 | 0 | 0 | 0 |
| UGC0007516 | Clust-UGC0003411 | 3250 | 4 | 1 | 0 | 1 |
| UGC0004097 | Clust-UGC0003411 | 448 | 1 | 0 | 0 | 0 |
| UGC0009401 | Clust-UGC0003411 | 2312 | 2 | 1 | 0 | 0 |
| UGC0007722 | Clust-UGC0003411 | 3378 | 4 | 1 | 0 | 1 |
| UGC0007234 | Clust-UGC0003411 | 3727 | 3 | 1 | 0 | 1 |
| UGC0009658 | Clust-UGC0003411 | 623 | 1 | 0 | 0 | 0 |
| UGC0006688 | Clust-UGC0003411 | 1798 | 1 | 0 | 0 | 0 |
| UGC0004141 | Clust-UGC0003411 | 791 | 1 | 0 | 0 | 0 |
| UGC0003789 | Clust-UGC0003411 | 210 | 1 | 0 | 0 | 0 |
| UGC0003415 | Clust-UGC0003415 | 763 | 2 | 1 | 0 | 0 |
| UGC0008371 | Clust-UGC0003415 | 1259 | 2 | 1 | 0 | 0 |
| UGC0011444 | Clust-UGC0003420 | 2840 | 3 | 1 | 0 | 0 |
| UGC0006917 | Clust-UGC0003420 | 675 | 2 | 1 | 0 | 1 |
| UGC0011592 | Clust-UGC0003420 | 3013 | 3 | 1 | 0 | 1 |
| UGC0011185 | Clust-UGC0003420 | 2918 | 5 | 1 | 1 | 1 |
| UGC0003420 | Clust-UGC0003420 | 2227 | 1 | 0 | 0 | 0 |
| UGC0004585 | Clust-UGC0003420 | 4830 | 9 | 1 | 0 | 1 |
| UGC0004525 | Clust-UGC0003420 | 1069 | 1 | 0 | 0 | 0 |
| UGC0005964 | Clust-UGC0003420 | 1464 | 2 | 1 | 0 | 1 |
| UGC0010128 | Clust-UGC0003420 | 3631 | 1 | 0 | 0 | 0 |
| UGC0004680 | Clust-UGC0003420 | 12238 | 76 | 1 | 0 | 1 |
| UGC0006322 | Clust-UGC0003520 | 1847 | 1 | 0 | 0 | 0 |
| UGC0003520 | Clust-UGC0003520 | 6224 | 4 | 1 | 0 | 1 |
| UGC0006354 | Clust-UGC0003520 | 3779 | 1 | 0 | 0 | 0 |
| UGC0011825 | Clust-UGC0003524 | 631 | 1 | 0 | 0 | 0 |
| UGC0012029 | Clust-UGC0003524 | 2006 | 3 | 1 | 0 | 1 |
| UGC0012681 | Clust-UGC0003524 | 9803 | 46 | 1 | 0 | 1 |
| UGC0013128 | Clust-UGC0003524 | 940 | 2 | 1 | 0 | 1 |
| UGC0013366 | Clust-UGC0003524 | 4365 | 4 | 1 | 0 | 1 |
| UGC0003524 | Clust-UGC0003524 | 2436 | 3 | 1 | 0 | 1 |
| UGC0012999 | Clust-UGC0003524 | 2132 | 4 | 1 | 1 | 1 |
| UGC0003539 | Clust-UGC0003526 | 10244 | 5 | 1 | 0 | 1 |
| UGC0004551 | Clust-UGC0003526 | 10388 | 50 | 1 | 0 | 1 |
| UGC0003526 | Clust-UGC0003526 | 7426 | 4 | 1 | 0 | 1 |
| UGC0011877 | Clust-UGC0003526 | 1671 | 1 | 0 | 0 | 0 |
| UGC0006308 | Clust-UGC0003528 | 3481 | 2 | 1 | 0 | 0 |
| UGC0005511 | Clust-UGC0003528 | 1476 | 1 | 0 | 0 | 0 |
| UGC0003528 | Clust-UGC0003528 | 5383 | 7 | 1 | 1 | 1 |
| UGC0010632 | Clust-UGC0003530 | 110 | 1 | 0 | 0 | 0 |
| UGC0004760 | Clust-UGC0003530 | 1121 | 3 | 1 | 0 | 0 |
| UGC0011180 | Clust-UGC0003530 | 623 | 1 | 0 | 0 | 0 |
| UGC0008308 | Clust-UGC0003530 | 979 | 1 | 0 | 0 | 0 |
| UGC0003530 | Clust-UGC0003530 | 3164 | 1 | 0 | 0 | 0 |
| UGC0003531 | Clust-UGC0003531 | 4157 | 3 | 1 | 0 | 1 |
| UGC0006273 | Clust-UGC0003533 | 2395 | 2 | 1 | 0 | 1 |
| UGC0009100 | Clust-UGC0003533 | 1093 | 1 | 0 | 0 | 0 |
| UGC0003533 | Clust-UGC0003533 | 769 | 1 | 0 | 0 | 0 |
| UGC0005070 | Clust-UGC0003536 | 1988 | 1 | 0 | 0 | 0 |
| UGC0008400 | Clust-UGC0003536 | 1283 | 2 | 1 | 1 | 0 |
| UGC0007770 | Clust-UGC0003536 | 4892 | 2 | 1 | 0 | 1 |
| UGC0007621 | Clust-UGC0003536 | 4180 | 1 | 0 | 0 | 0 |
| UGC0004506 | Clust-UGC0003536 | 11636 | 53 | 1 | 0 | 1 |
| UGC0003536 | Clust-UGC0003536 | 1668 | 1 | 0 | 0 | 0 |
| UGC0007357 | Clust-UGC0003536 | 2516 | 1 | 0 | 0 | 0 |
| UGC0003540 | Clust-UGC0003540 | 4756 | 4 | 1 | 0 | 1 |
| UGC0004604 | Clust-UGC0003540 | 2541 | 2 | 1 | 0 | 1 |

| | | | | | | |
|---|---|---|---|---|---|---|
| UGC0010225 | Clust-UGC0003541 | 1581 | 4 | 1 | 0 | 1 |
| UGC0003541 | Clust-UGC0003541 | 7372 | 14 | 1 | 0 | 1 |
| UGC0008072 | Clust-UGC0003541 | 660 | 2 | 1 | 0 | 1 |
| UGC0004976 | Clust-UGC0003541 | 2441 | 4 | 1 | 0 | 1 |
| UGC0012862 | Clust-UGC0003544 | 2015 | 2 | 1 | 0 | 0 |
| UGC0013013 | Clust-UGC0003544 | 3474 | 1 | 0 | 0 | 0 |
| UGC0006752 | Clust-UGC0003544 | 4409 | 1 | 0 | 0 | 0 |
| UGC0008744 | Clust-UGC0003544 | 1042 | 1 | 0 | 0 | 0 |
| UGC0004536 | Clust-UGC0003544 | 7143 | 4 | 1 | 0 | 1 |
| UGC0004578 | Clust-UGC0003544 | 2768 | 3 | 1 | 0 | 1 |
| UGC0008609 | Clust-UGC0003544 | 4816 | 2 | 1 | 0 | 1 |
| UGC0004676 | Clust-UGC0003544 | 12073 | 18 | 1 | 0 | 1 |
| UGC0008323 | Clust-UGC0003544 | 1009 | 1 | 0 | 0 | 0 |
| UGC0005348 | Clust-UGC0003544 | 3781 | 1 | 0 | 0 | 0 |
| UGC0003544 | Clust-UGC0003544 | 4698 | 3 | 1 | 0 | 1 |
| UGC0008700 | Clust-UGC0003544 | 939 | 1 | 0 | 0 | 0 |
| UGC0006846 | Clust-UGC0003544 | 529 | 1 | 0 | 0 | 0 |
| UGC0003564 | Clust-UGC0003564 | 8078 | 6 | 1 | 1 | 1 |
| UGC0003569 | Clust-UGC0003569 | 194 | 1 | 0 | 0 | 0 |
| UGC0003570 | Clust-UGC0003570 | 2993 | 4 | 1 | 0 | 0 |
| UGC0013206 | Clust-UGC0003576 | 283 | 1 | 0 | 0 | 0 |
| UGC0008282 | Clust-UGC0003576 | 582 | 1 | 0 | 0 | 0 |
| UGC0008694 | Clust-UGC0003576 | 2539 | 3 | 1 | 1 | 1 |
| UGC0008645 | Clust-UGC0003576 | 3395 | 3 | 1 | 0 | 1 |
| UGC0003576 | Clust-UGC0003576 | 11683 | 62 | 1 | 0 | 1 |
| UGC0004092 | Clust-UGC0003576 | 590 | 1 | 0 | 0 | 0 |
| UGC0009659 | Clust-UGC0003576 | 735 | 1 | 0 | 0 | 0 |
| UGC0012422 | Clust-UGC0003576 | 3102 | 4 | 1 | 1 | 1 |
| UGC0006970 | Clust-UGC0003579 | 1789 | 1 | 0 | 0 | 0 |
| UGC0009046 | Clust-UGC0003579 | 432 | 1 | 0 | 0 | 0 |
| UGC0007442 | Clust-UGC0003579 | 7768 | 11 | 1 | 0 | 1 |
| UGC0009457 | Clust-UGC0003579 | 349 | 1 | 0 | 0 | 0 |
| UGC0003579 | Clust-UGC0003579 | 9088 | 4 | 1 | 0 | 1 |
| UGC0007562 | Clust-UGC0003579 | 6196 | 9 | 1 | 0 | 0 |
| UGC0005968 | Clust-UGC0003579 | 336 | 1 | 0 | 0 | 0 |
| UGC0004608 | Clust-UGC0003579 | 3987 | 1 | 0 | 0 | 0 |
| UGC0006456 | Clust-UGC0003579 | 9793 | 10 | 1 | 0 | 1 |
| UGC0006849 | Clust-UGC0003579 | 3541 | 4 | 1 | 0 | 0 |
| UGC0008780 | Clust-UGC0003579 | 1686 | 3 | 1 | 0 | 0 |
| UGC0003580 | Clust-UGC0003580 | 4119 | 4 | 1 | 0 | 1 |
| UGC0006670 | Clust-UGC0003582 | 1580 | 2 | 1 | 0 | 1 |
| UGC0005674 | Clust-UGC0003582 | 1575 | 1 | 0 | 0 | 0 |
| UGC0012136 | Clust-UGC0003582 | 4891 | 6 | 1 | 0 | 1 |
| UGC0006777 | Clust-UGC0003582 | 6210 | 12 | 1 | 0 | 1 |
| UGC0012776 | Clust-UGC0003582 | 2063 | 2 | 1 | 0 | 1 |
| UGC0012923 | Clust-UGC0003582 | 2290 | 4 | 1 | 0 | 1 |
| UGC0007948 | Clust-UGC0003582 | 1853 | 3 | 1 | 0 | 1 |
| UGC0005522 | Clust-UGC0003582 | 1622 | 1 | 0 | 0 | 0 |
| UGC0007739 | Clust-UGC0003582 | 8044 | 34 | 1 | 0 | 1 |
| UGC0012044 | Clust-UGC0003582 | 408 | 1 | 0 | 0 | 0 |
| UGC0008228 | Clust-UGC0003582 | 1420 | 1 | 0 | 0 | 0 |
| UGC0004845 | Clust-UGC0003582 | 13067 | 98 | 1 | 0 | 1 |
| UGC0007214 | Clust-UGC0003582 | 2692 | 5 | 1 | 0 | 1 |
| UGC0005608 | Clust-UGC0003582 | 4232 | 3 | 1 | 0 | 1 |
| UGC0004714 | Clust-UGC0003582 | 12804 | 66 | 1 | 1 | 1 |
| UGC0004990 | Clust-UGC0003582 | 6887 | 17 | 1 | 0 | 1 |
| UGC0004972 | Clust-UGC0003582 | 2770 | 1 | 0 | 0 | 0 |
| UGC0003582 | Clust-UGC0003582 | 4328 | 7 | 1 | 0 | 1 |
| UGC0005394 | Clust-UGC0003582 | 2309 | 4 | 1 | 0 | 1 |
| UGC0007601 | Clust-UGC0003582 | 5340 | 5 | 1 | 0 | 1 |
| UGC0004875 | Clust-UGC0003582 | 51 | 1 | 0 | 0 | 0 |
| UGC0009332 | Clust-UGC0003582 | 4616 | 3 | 1 | 0 | 1 |
| UGC0007225 | Clust-UGC0003582 | 4279 | 5 | 1 | 0 | 1 |
| UGC0008484 | Clust-UGC0003585 | 2642 | 3 | 1 | 0 | 1 |
| UGC0006052 | Clust-UGC0003585 | 193 | 1 | 0 | 0 | 0 |
| UGC0012841 | Clust-UGC0003585 | 760 | 1 | 0 | 0 | 0 |
| UGC0012141 | Clust-UGC0003585 | 2894 | 2 | 1 | 0 | 1 |
| UGC0003585 | Clust-UGC0003585 | 7746 | 14 | 1 | 0 | 1 |
| UGC0012794 | Clust-UGC0003590 | 1200 | 2 | 1 | 0 | 1 |
| UGC0003590 | Clust-UGC0003590 | 8758 | 26 | 1 | 0 | 1 |
| UGC0011177 | Clust-UGC0003595 | 2425 | 3 | 1 | 1 | 0 |
| UGC0005991 | Clust-UGC0003595 | 760 | 1 | 0 | 0 | 0 |
| UGC0005953 | Clust-UGC0003595 | 1905 | 1 | 0 | 0 | 0 |
| UGC0008558 | Clust-UGC0003595 | 1161 | 1 | 0 | 0 | 0 |
| UGC0011441 | Clust-UGC0003595 | 2534 | 2 | 1 | 0 | 0 |
| UGC0012031 | Clust-UGC0003595 | 2104 | 2 | 1 | 1 | 0 |
| UGC0006482 | Clust-UGC0003595 | 1315 | 1 | 0 | 0 | 0 |
| UGC0006584 | Clust-UGC0003595 | 186 | 1 | 0 | 0 | 0 |
| UGC0006586 | Clust-UGC0003595 | 943 | 1 | 0 | 0 | 0 |

| | | | | | | |
|---|---|---|---|---|---|---|
| UGC0010803 | Clust-UGC0003595 | 466 | 4 | 1 | 0 | 0 |
| UGC0007369 | Clust-UGC0003595 | 3190 | 2 | 1 | 0 | 0 |
| UGC0009476 | Clust-UGC0003595 | 6726 | 7 | 1 | 0 | 1 |
| UGC0004531 | Clust-UGC0003595 | 1038 | 1 | 0 | 0 | 0 |
| UGC0006899 | Clust-UGC0003595 | 2707 | 1 | 0 | 0 | 0 |
| UGC0006634 | Clust-UGC0003595 | 634 | 1 | 0 | 0 | 0 |
| UGC0005563 | Clust-UGC0003595 | 1811 | 1 | 0 | 0 | 0 |
| UGC0003595 | Clust-UGC0003595 | 6790 | 6 | 1 | 0 | 1 |
| UGC0009147 | Clust-UGC0003595 | 1192 | 1 | 0 | 0 | 0 |
| UGC0008385 | Clust-UGC0003595 | 584 | 1 | 0 | 0 | 0 |
| UGC0004658 | Clust-UGC0003595 | 3266 | 2 | 1 | 0 | 1 |
| UGC0006992 | Clust-UGC0003595 | 1534 | 1 | 0 | 0 | 0 |
| UGC0008991 | Clust-UGC0003595 | 3018 | 2 | 1 | 0 | 0 |
| UGC0008996 | Clust-UGC0003595 | 2240 | 1 | 0 | 0 | 0 |
| UGC0009648 | Clust-UGC0003595 | 776 | 1 | 0 | 0 | 0 |
| UGC0007958 | Clust-UGC0003595 | 1087 | 1 | 0 | 0 | 0 |
| UGC0009269 | Clust-UGC0003595 | 4938 | 3 | 1 | 0 | 1 |
| UGC0009040 | Clust-UGC0003595 | 954 | 1 | 0 | 0 | 0 |
| UGC0009323 | Clust-UGC0003595 | 6588 | 14 | 1 | 0 | 1 |
| UGC0005798 | Clust-UGC0003595 | 1542 | 1 | 0 | 0 | 0 |
| UGC0010946 | Clust-UGC0003595 | 1230 | 2 | 1 | 0 | 0 |
| UGC0009514 | Clust-UGC0003595 | 3405 | 2 | 1 | 0 | 1 |
| UGC0004131 | Clust-UGC0003595 | 697 | 1 | 0 | 0 | 0 |
| UGC0012810 | Clust-UGC0003595 | 784 | 1 | 0 | 0 | 0 |
| UGC0012120 | Clust-UGC0003595 | 1632 | 1 | 0 | 0 | 0 |
| UGC0013273 | Clust-UGC0003595 | 850 | 1 | 0 | 0 | 0 |
| UGC0009259 | Clust-UGC0003595 | 6359 | 4 | 1 | 1 | 1 |
| UGC0010830 | Clust-UGC0003595 | 1351 | 1 | 0 | 0 | 0 |
| UGC0011818 | Clust-UGC0003595 | 1388 | 1 | 0 | 0 | 0 |
| UGC0013368 | Clust-UGC0003595 | 1697 | 9 | 1 | 0 | 1 |
| UGC0007110 | Clust-UGC0003595 | 1418 | 1 | 0 | 0 | 0 |
| UGC0008988 | Clust-UGC0003595 | 3011 | 4 | 1 | 0 | 1 |
| UGC0012695 | Clust-UGC0003595 | 6107 | 4 | 1 | 0 | 1 |
| UGC0012697 | Clust-UGC0003595 | 4658 | 5 | 1 | 2 | 1 |
| UGC0009654 | Clust-UGC0003595 | 788 | 1 | 0 | 0 | 0 |
| UGC0013131 | Clust-UGC0003595 | 742 | 1 | 0 | 0 | 0 |
| UGC0008945 | Clust-UGC0003595 | 867 | 1 | 0 | 0 | 0 |
| UGC0003784 | Clust-UGC0003595 | 418 | 1 | 0 | 0 | 0 |
| UGC0011941 | Clust-UGC0003595 | 1932 | 1 | 0 | 0 | 0 |
| UGC0003597 | Clust-UGC0003597 | 5314 | 7 | 1 | 0 | 1 |
| UGC0003600 | Clust-UGC0003600 | 10401 | 47 | 1 | 0 | 1 |
| UGC0003601 | Clust-UGC0003601 | 10199 | 8 | 1 | 0 | 1 |
| UGC0006854 | Clust-UGC0003603 | 4525 | 3 | 1 | 0 | 1 |
| UGC0009524 | Clust-UGC0003603 | 525 | 2 | 1 | 0 | 0 |
| UGC0006916 | Clust-UGC0003603 | 668 | 1 | 0 | 0 | 0 |
| UGC0010483 | Clust-UGC0003603 | 3253 | 3 | 1 | 1 | 0 |
| UGC0005869 | Clust-UGC0003603 | 2086 | 2 | 1 | 0 | 0 |
| UGC0011203 | Clust-UGC0003603 | 1438 | 3 | 1 | 0 | 1 |
| UGC0007303 | Clust-UGC0003603 | 3557 | 3 | 1 | 0 | 0 |
| UGC0003603 | Clust-UGC0003603 | 3902 | 3 | 1 | 0 | 0 |
| UGC0007015 | Clust-UGC0003603 | 1413 | 3 | 1 | 0 | 1 |
| UGC0005737 | Clust-UGC0003603 | 5973 | 3 | 1 | 0 | 1 |
| UGC0012013 | Clust-UGC0003603 | 2691 | 2 | 1 | 0 | 0 |
| UGC0004110 | Clust-UGC0003610 | 887 | 1 | 0 | 0 | 0 |
| UGC0003610 | Clust-UGC0003610 | 5483 | 4 | 1 | 0 | 1 |
| UGC0012895 | Clust-UGC0003610 | 1595 | 1 | 0 | 0 | 0 |
| UGC0006869 | Clust-UGC0003610 | 2256 | 2 | 1 | 0 | 0 |
| UGC0003709 | Clust-UGC0003709 | 1554 | 2 | 1 | 0 | 1 |
| UGC0010200 | Clust-UGC0003709 | 400 | 1 | 0 | 0 | 0 |
| UGC0003710 | Clust-UGC0003710 | 727 | 1 | 0 | 0 | 0 |
| UGC0009104 | Clust-UGC0003715 | 4448 | 3 | 1 | 0 | 1 |
| UGC0009185 | Clust-UGC0003715 | 4677 | 6 | 1 | 0 | 1 |
| UGC0003715 | Clust-UGC0003715 | 1223 | 1 | 0 | 0 | 0 |
| UGC0005098 | Clust-UGC0003719 | 661 | 1 | 0 | 0 | 0 |
| UGC0004984 | Clust-UGC0003719 | 5005 | 7 | 1 | 0 | 1 |
| UGC0005637 | Clust-UGC0003719 | 1061 | 1 | 0 | 0 | 0 |
| UGC0010865 | Clust-UGC0003719 | 5644 | 9 | 1 | 0 | 1 |
| UGC0006382 | Clust-UGC0003719 | 3588 | 2 | 1 | 0 | 1 |
| UGC0009624 | Clust-UGC0003719 | 609 | 2 | 1 | 0 | 1 |
| UGC0007305 | Clust-UGC0003719 | 5117 | 7 | 1 | 1 | 1 |
| UGC0008348 | Clust-UGC0003719 | 384 | 1 | 0 | 0 | 0 |
| UGC0010603 | Clust-UGC0003719 | 979 | 2 | 1 | 0 | 1 |
| UGC0004801 | Clust-UGC0003719 | 11189 | 69 | 1 | 0 | 1 |
| UGC0004111 | Clust-UGC0003719 | 445 | 1 | 0 | 0 | 0 |
| UGC0008286 | Clust-UGC0003719 | 1098 | 1 | 0 | 0 | 0 |
| UGC0007250 | Clust-UGC0003719 | 4342 | 7 | 1 | 0 | 1 |
| UGC0012628 | Clust-UGC0003719 | 5319 | 6 | 1 | 0 | 1 |
| UGC0003719 | Clust-UGC0003719 | 410 | 1 | 0 | 0 | 0 |
| UGC0008012 | Clust-UGC0003719 | 252 | 1 | 0 | 0 | 0 |

| | | | | | | |
|---|---|---|---|---|---|---|
| UGC0008074 | Clust-UGC0003719 | 235 | 1 | 0 | 0 | 0 |
| UGC0012766 | Clust-UGC0003719 | 2348 | 3 | 1 | 0 | 1 |
| UGC0013324 | Clust-UGC0003719 | 944 | 2 | 1 | 0 | 0 |
| UGC0005706 | Clust-UGC0003719 | 5505 | 8 | 1 | 0 | 1 |
| UGC0008213 | Clust-UGC0003719 | 53 | 1 | 0 | 0 | 0 |
| UGC0006809 | Clust-UGC0003719 | 2498 | 3 | 1 | 0 | 1 |
| UGC0003741 | Clust-UGC0003741 | 529 | 1 | 0 | 0 | 0 |
| UGC0007959 | Clust-UGC0003741 | 3330 | 7 | 1 | 0 | 1 |
| UGC0009402 | Clust-UGC0003741 | 8660 | 30 | 1 | 0 | 1 |
| UGC0009562 | Clust-UGC0003743 | 1844 | 2 | 1 | 0 | 1 |
| UGC0003743 | Clust-UGC0003743 | 435 | 1 | 0 | 0 | 0 |
| UGC0004132 | Clust-UGC0003743 | 1049 | 3 | 1 | 0 | 1 |
| UGC0004666 | Clust-UGC0003743 | 9895 | 45 | 1 | 0 | 1 |
| UGC0007746 | Clust-UGC0003743 | 8620 | 41 | 1 | 0 | 1 |
| UGC0012845 | Clust-UGC0003747 | 3423 | 6 | 1 | 0 | 1 |
| UGC0003747 | Clust-UGC0003747 | 791 | 1 | 0 | 0 | 0 |
| UGC0010169 | Clust-UGC0003756 | 1524 | 2 | 1 | 0 | 0 |
| UGC0005376 | Clust-UGC0003756 | 4279 | 7 | 1 | 0 | 1 |
| UGC0005912 | Clust-UGC0003756 | 3978 | 2 | 1 | 0 | 0 |
| UGC0010844 | Clust-UGC0003756 | 2558 | 3 | 1 | 0 | 0 |
| UGC0009384 | Clust-UGC0003756 | 8434 | 13 | 1 | 0 | 1 |
| UGC0005644 | Clust-UGC0003756 | 1214 | 1 | 0 | 0 | 0 |
| UGC0013129 | Clust-UGC0003756 | 292 | 1 | 0 | 0 | 0 |
| UGC0011787 | Clust-UGC0003756 | 4365 | 3 | 1 | 0 | 0 |
| UGC0003756 | Clust-UGC0003756 | 1148 | 2 | 1 | 0 | 0 |
| UGC0008053 | Clust-UGC0003756 | 1619 | 2 | 1 | 0 | 0 |
| UGC0010182 | Clust-UGC0003756 | 1855 | 2 | 1 | 0 | 0 |
| UGC0009585 | Clust-UGC0003756 | 6115 | 7 | 1 | 1 | 1 |
| UGC0006803 | Clust-UGC0003756 | 3813 | 4 | 1 | 0 | 0 |
| UGC0009674 | Clust-UGC0003756 | 2285 | 4 | 1 | 1 | 0 |
| UGC0006767 | Clust-UGC0003759 | 5484 | 1 | 0 | 0 | 0 |
| UGC0003759 | Clust-UGC0003759 | 303 | 1 | 0 | 0 | 0 |
| UGC0006923 | Clust-UGC0003759 | 682 | 1 | 0 | 0 | 0 |
| UGC0003769 | Clust-UGC0003769 | 274 | 1 | 0 | 0 | 0 |
| UGC0007921 | Clust-UGC0003776 | 12853 | 97 | 1 | 1 | 1 |
| UGC0008448 | Clust-UGC0003776 | 337 | 1 | 0 | 0 | 0 |
| UGC0003776 | Clust-UGC0003776 | 1525 | 2 | 1 | 0 | 1 |
| UGC0008562 | Clust-UGC0003776 | 2644 | 2 | 1 | 0 | 1 |
| UGC0008254 | Clust-UGC0003776 | 371 | 1 | 0 | 0 | 0 |
| UGC0009459 | Clust-UGC0003779 | 2617 | 1 | 0 | 0 | 0 |
| UGC0003779 | Clust-UGC0003779 | 417 | 1 | 0 | 0 | 0 |
| UGC0007727 | Clust-UGC0003779 | 5374 | 3 | 1 | 0 | 1 |
| UGC0009563 | Clust-UGC0003787 | 127 | 1 | 0 | 0 | 0 |
| UGC0005362 | Clust-UGC0003787 | 1354 | 1 | 0 | 0 | 0 |
| UGC0009590 | Clust-UGC0003787 | 2322 | 2 | 1 | 0 | 1 |
| UGC0009101 | Clust-UGC0003787 | 989 | 2 | 1 | 0 | 1 |
| UGC0009669 | Clust-UGC0003787 | 1490 | 1 | 0 | 0 | 0 |
| UGC0009497 | Clust-UGC0003787 | 4775 | 3 | 1 | 0 | 1 |
| UGC0009729 | Clust-UGC0003787 | 1973 | 1 | 0 | 0 | 0 |
| UGC0006077 | Clust-UGC0003787 | 1406 | 1 | 0 | 0 | 0 |
| UGC0009705 | Clust-UGC0003787 | 3149 | 4 | 1 | 0 | 1 |
| UGC0008614 | Clust-UGC0003787 | 2579 | 1 | 0 | 0 | 0 |
| UGC0009632 | Clust-UGC0003787 | 452 | 1 | 0 | 0 | 0 |
| UGC0006808 | Clust-UGC0003787 | 2665 | 1 | 0 | 0 | 0 |
| UGC0003787 | Clust-UGC0003787 | 439 | 1 | 0 | 0 | 0 |
| UGC0012249 | Clust-UGC0003796 | 3811 | 11 | 1 | 0 | 1 |
| UGC0004941 | Clust-UGC0003796 | 2048 | 1 | 0 | 0 | 0 |
| UGC0006895 | Clust-UGC0003796 | 1634 | 2 | 1 | 0 | 0 |
| UGC0012963 | Clust-UGC0003796 | 5494 | 9 | 1 | 0 | 1 |
| UGC0004480 | Clust-UGC0003796 | 5293 | 5 | 1 | 0 | 1 |
| UGC0005728 | Clust-UGC0003796 | 6852 | 13 | 1 | 0 | 1 |
| UGC0007537 | Clust-UGC0003796 | 6340 | 14 | 1 | 0 | 1 |
| UGC0003796 | Clust-UGC0003796 | 504 | 1 | 0 | 0 | 0 |
| UGC0008571 | Clust-UGC0003797 | 5147 | 5 | 1 | 0 | 1 |
| UGC0003797 | Clust-UGC0003797 | 417 | 1 | 0 | 0 | 0 |
| UGC0012998 | Clust-UGC0003797 | 2574 | 1 | 0 | 0 | 0 |
| UGC0004090 | Clust-UGC0004090 | 159 | 1 | 0 | 0 | 0 |
| UGC0006056 | Clust-UGC0004093 | 1008 | 1 | 0 | 0 | 0 |
| UGC0010895 | Clust-UGC0004093 | 3233 | 3 | 1 | 0 | 1 |
| UGC0004093 | Clust-UGC0004093 | 583 | 1 | 0 | 0 | 0 |
| UGC0004105 | Clust-UGC0004105 | 141 | 1 | 0 | 0 | 0 |
| UGC0010094 | Clust-UGC0004105 | 10428 | 36 | 1 | 0 | 1 |
| UGC0006408 | Clust-UGC0004105 | 4891 | 7 | 1 | 0 | 1 |
| UGC0011821 | Clust-UGC0004105 | 1426 | 2 | 1 | 1 | 0 |
| UGC0005104 | Clust-UGC0004105 | 1341 | 2 | 1 | 0 | 0 |
| UGC0010453 | Clust-UGC0004105 | 3424 | 6 | 1 | 0 | 1 |
| UGC0010272 | Clust-UGC0004105 | 4936 | 3 | 1 | 0 | 1 |
| UGC0004107 | Clust-UGC0004107 | 206 | 1 | 0 | 0 | 0 |
| UGC0009784 | Clust-UGC0004107 | 2415 | 2 | 1 | 0 | 1 |

| | | | | | | |
|---|---|---|---|---|---|---|
| UGC0007384 | Clust-UGC0004107 | 1533 | 2 | 1 | 0 | 1 |
| UGC0009490 | Clust-UGC0004107 | 4519 | 2 | 1 | 0 | 1 |
| UGC0006748 | Clust-UGC0004107 | 6407 | 6 | 1 | 0 | 1 |
| UGC0006735 | Clust-UGC0004107 | 427 | 1 | 0 | 0 | 0 |
| UGC0006825 | Clust-UGC0004107 | 7230 | 15 | 1 | 0 | 1 |
| UGC0009026 | Clust-UGC0004118 | 109 | 1 | 0 | 0 | 0 |
| UGC0004118 | Clust-UGC0004118 | 1155 | 1 | 0 | 0 | 0 |
| UGC0011120 | Clust-UGC0004118 | 1274 | 1 | 0 | 0 | 0 |
| UGC0006329 | Clust-UGC0004136 | 3309 | 3 | 1 | 0 | 1 |
| UGC0004136 | Clust-UGC0004136 | 1654 | 3 | 1 | 0 | 1 |
| UGC0010997 | Clust-UGC0004140 | 5794 | 11 | 1 | 1 | 1 |
| UGC0007491 | Clust-UGC0004140 | 6069 | 7 | 1 | 0 | 1 |
| UGC0011154 | Clust-UGC0004140 | 2870 | 5 | 1 | 0 | 1 |
| UGC0007679 | Clust-UGC0004140 | 4839 | 4 | 1 | 0 | 1 |
| UGC0012342 | Clust-UGC0004140 | 315 | 1 | 0 | 0 | 0 |
| UGC0007283 | Clust-UGC0004140 | 6282 | 9 | 1 | 0 | 1 |
| UGC0007349 | Clust-UGC0004140 | 3983 | 6 | 1 | 0 | 1 |
| UGC0006890 | Clust-UGC0004140 | 5393 | 5 | 1 | 0 | 1 |
| UGC0006407 | Clust-UGC0004140 | 9909 | 46 | 1 | 0 | 1 |
| UGC0011865 | Clust-UGC0004140 | 76 | 1 | 0 | 0 | 0 |
| UGC0011953 | Clust-UGC0004140 | 8214 | 19 | 1 | 0 | 1 |
| UGC0012147 | Clust-UGC0004140 | 4659 | 8 | 1 | 0 | 1 |
| UGC0009182 | Clust-UGC0004140 | 7928 | 28 | 1 | 0 | 1 |
| UGC0012353 | Clust-UGC0004140 | 2633 | 3 | 1 | 0 | 1 |
| UGC0005941 | Clust-UGC0004140 | 6521 | 12 | 1 | 0 | 1 |
| UGC0005665 | Clust-UGC0004140 | 3136 | 3 | 1 | 0 | 1 |
| UGC0008478 | Clust-UGC0004140 | 827 | 2 | 1 | 0 | 0 |
| UGC0013065 | Clust-UGC0004140 | 9783 | 41 | 1 | 0 | 1 |
| UGC0011019 | Clust-UGC0004140 | 4901 | 2 | 1 | 0 | 1 |
| UGC0010181 | Clust-UGC0004140 | 1516 | 1 | 0 | 0 | 0 |
| UGC0011210 | Clust-UGC0004140 | 601 | 1 | 0 | 0 | 0 |
| UGC0006472 | Clust-UGC0004140 | 3006 | 2 | 1 | 0 | 1 |
| UGC0012990 | Clust-UGC0004140 | 9479 | 25 | 1 | 0 | 1 |
| UGC0009030 | Clust-UGC0004140 | 1195 | 1 | 0 | 0 | 0 |
| UGC0004140 | Clust-UGC0004140 | 1405 | 4 | 1 | 1 | 1 |
| UGC0013289 | Clust-UGC0004140 | 420 | 1 | 0 | 0 | 0 |
| UGC0013284 | Clust-UGC0004140 | 1427 | 1 | 0 | 0 | 0 |
| UGC0010956 | Clust-UGC0004140 | 4527 | 5 | 1 | 0 | 1 |
| UGC0009717 | Clust-UGC0004142 | 1751 | 2 | 1 | 0 | 0 |
| UGC0011461 | Clust-UGC0004142 | 12747 | 20 | 1 | 0 | 1 |
| UGC0010019 | Clust-UGC0004142 | 2357 | 1 | 0 | 0 | 0 |
| UGC0012211 | Clust-UGC0004142 | 1719 | 1 | 0 | 0 | 0 |
| UGC0004142 | Clust-UGC0004142 | 559 | 1 | 0 | 0 | 0 |
| UGC0005911 | Clust-UGC0004143 | 3112 | 2 | 1 | 0 | 1 |
| UGC0004946 | Clust-UGC0004143 | 4984 | 10 | 1 | 0 | 1 |
| UGC0007489 | Clust-UGC0004143 | 3488 | 3 | 1 | 0 | 1 |
| UGC0013331 | Clust-UGC0004143 | 505 | 2 | 1 | 0 | 0 |
| UGC0008344 | Clust-UGC0004143 | 731 | 2 | 1 | 0 | 1 |
| UGC0009637 | Clust-UGC0004143 | 1001 | 2 | 1 | 0 | 1 |
| UGC0006700 | Clust-UGC0004143 | 213 | 1 | 0 | 0 | 0 |
| UGC0005044 | Clust-UGC0004143 | 2429 | 1 | 0 | 0 | 0 |
| UGC0007171 | Clust-UGC0004143 | 900 | 3 | 1 | 0 | 1 |
| UGC0008016 | Clust-UGC0004143 | 1333 | 1 | 0 | 0 | 0 |
| UGC0005704 | Clust-UGC0004143 | 3872 | 5 | 1 | 0 | 1 |
| UGC0004143 | Clust-UGC0004143 | 556 | 2 | 1 | 0 | 1 |
| UGC0009196 | Clust-UGC0004145 | 4093 | 2 | 1 | 0 | 1 |
| UGC0010227 | Clust-UGC0004145 | 488 | 1 | 0 | 0 | 0 |
| UGC0009209 | Clust-UGC0004145 | 6972 | 2 | 1 | 0 | 1 |
| UGC0008064 | Clust-UGC0004145 | 314 | 1 | 0 | 0 | 0 |
| UGC0009083 | Clust-UGC0004145 | 966 | 1 | 0 | 0 | 0 |
| UGC0007647 | Clust-UGC0004145 | 4597 | 1 | 0 | 0 | 0 |
| UGC0009764 | Clust-UGC0004145 | 5193 | 2 | 1 | 0 | 1 |
| UGC0006644 | Clust-UGC0004145 | 2166 | 2 | 1 | 0 | 1 |
| UGC0004145 | Clust-UGC0004145 | 1431 | 1 | 0 | 0 | 0 |
| UGC0004151 | Clust-UGC0004151 | 6385 | 11 | 1 | 0 | 1 |
| UGC0012280 | Clust-UGC0004152 | 816 | 1 | 0 | 0 | 0 |
| UGC0009716 | Clust-UGC0004152 | 1408 | 2 | 1 | 0 | 1 |
| UGC0011418 | Clust-UGC0004152 | 6748 | 7 | 1 | 0 | 1 |
| UGC0012400 | Clust-UGC0004152 | 5601 | 4 | 1 | 0 | 1 |
| UGC0012800 | Clust-UGC0004152 | 1628 | 1 | 0 | 0 | 0 |
| UGC0007616 | Clust-UGC0004152 | 4513 | 3 | 1 | 0 | 1 |
| UGC0013073 | Clust-UGC0004152 | 6171 | 12 | 1 | 0 | 1 |
| UGC0013058 | Clust-UGC0004152 | 17643 | 187 | 1 | 1 | 1 |
| UGC0012943 | Clust-UGC0004152 | 2942 | 2 | 1 | 1 | 0 |
| UGC0007448 | Clust-UGC0004152 | 2600 | 1 | 0 | 0 | 0 |
| UGC0006632 | Clust-UGC0004152 | 1191 | 1 | 0 | 0 | 0 |
| UGC0013096 | Clust-UGC0004152 | 803 | 1 | 0 | 0 | 0 |
| UGC0012389 | Clust-UGC0004152 | 2438 | 2 | 1 | 0 | 1 |
| UGC0011596 | Clust-UGC0004152 | 1616 | 1 | 0 | 0 | 0 |

| | | | | | | |
|---|---|---|---|---|---|---|
| UGC0007730 | Clust-UGC0004152 | 2535 | 1 | 0 | 0 | 0 |
| UGC0011127 | Clust-UGC0004152 | 4369 | 4 | 1 | 0 | 1 |
| UGC0004471 | Clust-UGC0004152 | 8252 | 15 | 1 | 0 | 1 |
| UGC0006169 | Clust-UGC0004152 | 1278 | 2 | 1 | 0 | 1 |
| UGC0010968 | Clust-UGC0004152 | 7990 | 19 | 1 | 0 | 1 |
| UGC0011184 | Clust-UGC0004152 | 3244 | 4 | 1 | 0 | 1 |
| UGC0004152 | Clust-UGC0004152 | 2334 | 1 | 0 | 0 | 0 |
| UGC0004584 | Clust-UGC0004152 | 8950 | 32 | 1 | 0 | 1 |
| UGC0011939 | Clust-UGC0004152 | 3440 | 3 | 1 | 1 | 1 |
| UGC0012299 | Clust-UGC0004152 | 1874 | 2 | 1 | 0 | 1 |
| UGC0005064 | Clust-UGC0004152 | 3419 | 2 | 1 | 0 | 1 |
| UGC0009577 | Clust-UGC0004152 | 5746 | 3 | 1 | 0 | 1 |
| UGC0013271 | Clust-UGC0004152 | 743 | 1 | 0 | 0 | 0 |
| UGC0004528 | Clust-UGC0004152 | 5182 | 3 | 1 | 0 | 1 |
| UGC0013045 | Clust-UGC0004152 | 5844 | 7 | 1 | 2 | 1 |
| UGC0010837 | Clust-UGC0004152 | 749 | 1 | 0 | 0 | 0 |
| UGC0010852 | Clust-UGC0004152 | 3698 | 3 | 1 | 0 | 1 |
| UGC0012985 | Clust-UGC0004152 | 2824 | 2 | 1 | 0 | 1 |
| UGC0005670 | Clust-UGC0004152 | 5227 | 6 | 1 | 0 | 1 |
| UGC0012761 | Clust-UGC0004152 | 1679 | 1 | 0 | 0 | 0 |
| UGC0008615 | Clust-UGC0004152 | 3309 | 3 | 1 | 0 | 1 |
| UGC0011870 | Clust-UGC0004152 | 5030 | 3 | 1 | 0 | 1 |
| UGC0009582 | Clust-UGC0004152 | 5675 | 5 | 1 | 0 | 1 |
| UGC0005832 | Clust-UGC0004152 | 5066 | 3 | 1 | 0 | 1 |
| UGC0006173 | Clust-UGC0004152 | 2198 | 2 | 1 | 0 | 1 |
| UGC0012055 | Clust-UGC0004152 | 1166 | 1 | 0 | 0 | 0 |
| UGC0006546 | Clust-UGC0004152 | 912 | 1 | 0 | 0 | 0 |
| UGC0004163 | Clust-UGC0004163 | 2786 | 1 | 0 | 0 | 0 |
| UGC0007248 | Clust-UGC0004163 | 9608 | 46 | 1 | 0 | 1 |
| UGC0012611 | Clust-UGC0004163 | 15549 | 137 | 1 | 0 | 1 |
| UGC0005404 | Clust-UGC0004163 | 241 | 1 | 0 | 0 | 0 |
| UGC0007188 | Clust-UGC0004163 | 2528 | 3 | 1 | 0 | 1 |
| UGC0006835 | Clust-UGC0004163 | 5500 | 14 | 1 | 0 | 1 |
| UGC0007056 | Clust-UGC0004163 | 2343 | 2 | 1 | 0 | 1 |
| UGC0006645 | Clust-UGC0004163 | 2820 | 1 | 0 | 0 | 0 |
| UGC0007158 | Clust-UGC0004163 | 7275 | 18 | 1 | 0 | 1 |
| UGC0004167 | Clust-UGC0004167 | 2339 | 2 | 1 | 0 | 0 |
| UGC0006275 | Clust-UGC0004176 | 4689 | 1 | 0 | 0 | 0 |
| UGC0009631 | Clust-UGC0004176 | 209 | 1 | 0 | 0 | 0 |
| UGC0004692 | Clust-UGC0004176 | 19095 | 24 | 1 | 0 | 1 |
| UGC0004176 | Clust-UGC0004176 | 1466 | 1 | 0 | 0 | 0 |
| UGC0007052 | Clust-UGC0004176 | 304 | 1 | 0 | 0 | 0 |
| UGC0008695 | Clust-UGC0004176 | 776 | 1 | 0 | 0 | 0 |
| UGC0007115 | Clust-UGC0004176 | 695 | 1 | 0 | 0 | 0 |
| UGC0008456 | Clust-UGC0004176 | 680 | 1 | 0 | 0 | 0 |
| UGC0004490 | Clust-UGC0004490 | 11567 | 4 | 1 | 0 | 1 |
| UGC0004508 | Clust-UGC0004508 | 4964 | 10 | 1 | 1 | 1 |
| UGC0004509 | Clust-UGC0004509 | 8372 | 26 | 1 | 0 | 1 |
| UGC0006063 | Clust-UGC0004513 | 1223 | 1 | 0 | 0 | 0 |
| UGC0008768 | Clust-UGC0004513 | 926 | 1 | 0 | 0 | 0 |
| UGC0004513 | Clust-UGC0004513 | 5314 | 6 | 1 | 0 | 1 |
| UGC0005865 | Clust-UGC0004513 | 6052 | 12 | 1 | 0 | 1 |
| UGC0007933 | Clust-UGC0004513 | 1155 | 2 | 1 | 0 | 0 |
| UGC0004515 | Clust-UGC0004515 | 5636 | 6 | 1 | 0 | 1 |
| UGC0008872 | Clust-UGC0004515 | 1734 | 2 | 1 | 0 | 1 |
| UGC0008851 | Clust-UGC0004515 | 1710 | 1 | 0 | 0 | 0 |
| UGC0004995 | Clust-UGC0004515 | 4651 | 6 | 1 | 0 | 1 |
| UGC0004517 | Clust-UGC0004517 | 2617 | 1 | 0 | 0 | 0 |
| UGC0008349 | Clust-UGC0004517 | 71 | 1 | 0 | 0 | 0 |
| UGC0006151 | Clust-UGC0004517 | 1178 | 2 | 1 | 1 | 0 |
| UGC0004527 | Clust-UGC0004527 | 4487 | 11 | 1 | 0 | 1 |
| UGC0007590 | Clust-UGC0004533 | 4999 | 7 | 1 | 1 | 0 |
| UGC0004599 | Clust-UGC0004533 | 2860 | 4 | 1 | 0 | 1 |
| UGC0005657 | Clust-UGC0004533 | 1681 | 3 | 1 | 0 | 1 |
| UGC0011440 | Clust-UGC0004533 | 2286 | 5 | 1 | 0 | 0 |
| UGC0008727 | Clust-UGC0004533 | 1126 | 2 | 1 | 0 | 0 |
| UGC0004533 | Clust-UGC0004533 | 1532 | 2 | 1 | 0 | 1 |
| UGC0004949 | Clust-UGC0004533 | 1667 | 1 | 0 | 0 | 0 |
| UGC0004961 | Clust-UGC0004533 | 6735 | 12 | 1 | 0 | 1 |
| UGC0005714 | Clust-UGC0004533 | 9078 | 33 | 1 | 0 | 1 |
| UGC0005107 | Clust-UGC0004533 | 1732 | 1 | 0 | 0 | 0 |
| UGC0006342 | Clust-UGC0004533 | 6619 | 8 | 1 | 0 | 1 |
| UGC0012989 | Clust-UGC0004533 | 5384 | 9 | 1 | 0 | 1 |
| UGC0012143 | Clust-UGC0004533 | 2345 | 2 | 1 | 1 | 0 |
| UGC0009559 | Clust-UGC0004533 | 4296 | 7 | 1 | 0 | 1 |
| UGC0008774 | Clust-UGC0004533 | 1315 | 3 | 1 | 0 | 0 |
| UGC0009025 | Clust-UGC0004533 | 3646 | 4 | 1 | 0 | 0 |
| UGC0004539 | Clust-UGC0004539 | 1474 | 1 | 0 | 0 | 0 |
| UGC0008556 | Clust-UGC0004539 | 1865 | 1 | 0 | 0 | 0 |

| | | | | | | |
|---|---|---|---|---|---|---|
| UGC0007619 | Clust-UGC0004539 | 3217 | 1 | 0 | 0 | 0 |
| UGC0007618 | Clust-UGC0004539 | 2080 | 2 | 1 | 0 | 1 |
| UGC0006856 | Clust-UGC0004539 | 4442 | 2 | 1 | 1 | 0 |
| UGC0008200 | Clust-UGC0004539 | 2164 | 1 | 0 | 0 | 0 |
| UGC0008221 | Clust-UGC0004539 | 571 | 1 | 0 | 0 | 0 |
| UGC0013257 | Clust-UGC0004541 | 2185 | 1 | 0 | 0 | 0 |
| UGC0004541 | Clust-UGC0004541 | 1429 | 1 | 0 | 0 | 0 |
| UGC0011420 | Clust-UGC0004547 | 3082 | 2 | 1 | 0 | 0 |
| UGC0011864 | Clust-UGC0004547 | 2531 | 1 | 0 | 0 | 0 |
| UGC0004547 | Clust-UGC0004547 | 6131 | 2 | 1 | 0 | 1 |
| UGC0007329 | Clust-UGC0004556 | 967 | 1 | 0 | 0 | 0 |
| UGC0004556 | Clust-UGC0004556 | 8263 | 9 | 1 | 0 | 1 |
| UGC0010193 | Clust-UGC0004556 | 1726 | 6 | 1 | 0 | 0 |
| UGC0004570 | Clust-UGC0004570 | 5848 | 5 | 1 | 0 | 1 |
| UGC0004590 | Clust-UGC0004590 | 7404 | 19 | 1 | 0 | 1 |
| UGC0008176 | Clust-UGC0004590 | 2156 | 2 | 1 | 0 | 1 |
| UGC0004592 | Clust-UGC0004592 | 5929 | 3 | 1 | 0 | 0 |
| UGC0005351 | Clust-UGC0004592 | 2895 | 4 | 1 | 0 | 1 |
| UGC0008576 | Clust-UGC0004592 | 1833 | 3 | 1 | 0 | 0 |
| UGC0006476 | Clust-UGC0004592 | 2030 | 1 | 0 | 0 | 0 |
| UGC0011599 | Clust-UGC0004592 | 937 | 1 | 0 | 0 | 0 |
| UGC0005383 | Clust-UGC0004592 | 3054 | 1 | 0 | 0 | 0 |
| UGC0011495 | Clust-UGC0004592 | 4442 | 2 | 1 | 0 | 1 |
| UGC0008714 | Clust-UGC0004592 | 426 | 1 | 0 | 0 | 0 |
| UGC0007615 | Clust-UGC0004592 | 2320 | 2 | 1 | 0 | 1 |
| UGC0012936 | Clust-UGC0004595 | 8861 | 39 | 1 | 0 | 1 |
| UGC0004595 | Clust-UGC0004595 | 2434 | 3 | 1 | 0 | 1 |
| UGC0005353 | Clust-UGC0004595 | 473 | 1 | 0 | 0 | 0 |
| UGC0005400 | Clust-UGC0004595 | 2254 | 2 | 1 | 0 | 1 |
| UGC0005612 | Clust-UGC0004595 | 5387 | 5 | 1 | 0 | 1 |
| UGC0006617 | Clust-UGC0004595 | 1210 | 1 | 0 | 0 | 0 |
| UGC0012855 | Clust-UGC0004595 | 3755 | 7 | 1 | 1 | 1 |
| UGC0005672 | Clust-UGC0004595 | 3460 | 3 | 1 | 0 | 1 |
| UGC0004621 | Clust-UGC0004595 | 6210 | 4 | 1 | 0 | 1 |
| UGC0013325 | Clust-UGC0004595 | 3379 | 5 | 1 | 1 | 1 |
| UGC0005703 | Clust-UGC0004595 | 15501 | 134 | 1 | 0 | 1 |
| UGC0005854 | Clust-UGC0004595 | 1704 | 2 | 1 | 0 | 1 |
| UGC0006355 | Clust-UGC0004595 | 5261 | 5 | 1 | 1 | 1 |
| UGC0004720 | Clust-UGC0004606 | 6491 | 13 | 1 | 0 | 1 |
| UGC0007079 | Clust-UGC0004606 | 3056 | 3 | 1 | 1 | 1 |
| UGC0004606 | Clust-UGC0004606 | 5478 | 13 | 1 | 1 | 1 |
| UGC0009141 | Clust-UGC0004613 | 688 | 1 | 0 | 0 | 0 |
| UGC0004613 | Clust-UGC0004613 | 1963 | 2 | 1 | 0 | 1 |
| UGC0005324 | Clust-UGC0004613 | 2353 | 3 | 1 | 0 | 1 |
| UGC0009356 | Clust-UGC0004613 | 4798 | 6 | 1 | 0 | 1 |
| UGC0004614 | Clust-UGC0004614 | 8598 | 26 | 1 | 0 | 1 |
| UGC0004615 | Clust-UGC0004615 | 4972 | 7 | 1 | 0 | 1 |
| UGC0004679 | Clust-UGC0004624 | 14082 | 88 | 1 | 0 | 1 |
| UGC0007090 | Clust-UGC0004624 | 2488 | 3 | 1 | 0 | 1 |
| UGC0004624 | Clust-UGC0004624 | 8032 | 23 | 1 | 0 | 1 |
| UGC0004628 | Clust-UGC0004628 | 4582 | 3 | 1 | 0 | 1 |
| UGC0004632 | Clust-UGC0004632 | 6386 | 2 | 1 | 0 | 1 |
| UGC0011594 | Clust-UGC0004632 | 1929 | 3 | 1 | 0 | 0 |
| UGC0011206 | Clust-UGC0004632 | 7610 | 3 | 1 | 0 | 1 |
| UGC0007078 | Clust-UGC0004632 | 620 | 2 | 1 | 0 | 0 |
| UGC0010578 | Clust-UGC0004632 | 2339 | 2 | 1 | 0 | 1 |
| UGC0005118 | Clust-UGC0004632 | 2458 | 1 | 0 | 0 | 0 |
| UGC0004635 | Clust-UGC0004635 | 5817 | 9 | 1 | 0 | 1 |
| UGC0004636 | Clust-UGC0004636 | 5672 | 5 | 1 | 0 | 1 |
| UGC0006530 | Clust-UGC0004636 | 2205 | 3 | 1 | 0 | 1 |
| UGC0007019 | Clust-UGC0004636 | 1332 | 1 | 0 | 0 | 0 |
| UGC0006070 | Clust-UGC0004637 | 171 | 1 | 0 | 0 | 0 |
| UGC0004637 | Clust-UGC0004637 | 5267 | 3 | 1 | 0 | 1 |
| UGC0011539 | Clust-UGC0004637 | 1444 | 2 | 1 | 0 | 0 |
| UGC0005031 | Clust-UGC0004641 | 5223 | 7 | 1 | 0 | 1 |
| UGC0012609 | Clust-UGC0004641 | 8085 | 18 | 1 | 0 | 1 |
| UGC0011000 | Clust-UGC0004641 | 2947 | 2 | 1 | 0 | 1 |
| UGC0004641 | Clust-UGC0004641 | 11223 | 55 | 1 | 0 | 1 |
| UGC0008660 | Clust-UGC0004641 | 2434 | 2 | 1 | 0 | 0 |
| UGC0007300 | Clust-UGC0004642 | 1178 | 3 | 1 | 0 | 0 |
| UGC0004642 | Clust-UGC0004642 | 7376 | 19 | 1 | 0 | 1 |
| UGC0004643 | Clust-UGC0004643 | 7157 | 4 | 1 | 0 | 1 |
| UGC0005389 | Clust-UGC0004644 | 2117 | 5 | 1 | 0 | 1 |
| UGC0004806 | Clust-UGC0004644 | 11599 | 54 | 1 | 0 | 1 |
| UGC0010292 | Clust-UGC0004644 | 243 | 1 | 0 | 0 | 0 |
| UGC0005620 | Clust-UGC0004644 | 4376 | 11 | 1 | 0 | 1 |
| UGC0004644 | Clust-UGC0004644 | 7503 | 21 | 1 | 0 | 1 |
| UGC0005334 | Clust-UGC0004644 | 3109 | 4 | 1 | 0 | 1 |
| UGC0005786 | Clust-UGC0004646 | 7076 | 3 | 1 | 0 | 1 |

| | | | | | | |
|---|---|---|---|---|---|---|
| UGC0004646 | Clust-UGC0004646 | 7722 | 6 | 1 | 0 | 1 |
| UGC0007320 | Clust-UGC0004646 | 4139 | 2 | 1 | 0 | 1 |
| UGC0007468 | Clust-UGC0004646 | 2998 | 1 | 0 | 0 | 0 |
| UGC0007126 | Clust-UGC0004646 | 6654 | 2 | 1 | 0 | 1 |
| UGC0006019 | Clust-UGC0004646 | 3392 | 2 | 1 | 0 | 1 |
| UGC0005676 | Clust-UGC0004646 | 1517 | 1 | 0 | 0 | 0 |
| UGC0007983 | Clust-UGC0004662 | 919 | 1 | 0 | 0 | 0 |
| UGC0004662 | Clust-UGC0004662 | 2358 | 1 | 0 | 0 | 0 |
| UGC0010983 | Clust-UGC0004662 | 3546 | 2 | 1 | 0 | 1 |
| UGC0008928 | Clust-UGC0004662 | 2857 | 2 | 1 | 0 | 1 |
| UGC0005019 | Clust-UGC0004670 | 920 | 1 | 0 | 0 | 0 |
| UGC0004670 | Clust-UGC0004670 | 7921 | 32 | 1 | 0 | 1 |
| UGC0005686 | Clust-UGC0004670 | 3551 | 1 | 0 | 0 | 0 |
| UGC0005395 | Clust-UGC0004670 | 3136 | 3 | 1 | 0 | 1 |
| UGC0009521 | Clust-UGC0004671 | 294 | 2 | 1 | 0 | 1 |
| UGC0004671 | Clust-UGC0004671 | 6081 | 8 | 1 | 0 | 1 |
| UGC0008174 | Clust-UGC0004671 | 3585 | 2 | 1 | 0 | 1 |
| UGC0012437 | Clust-UGC0004684 | 4126 | 2 | 1 | 0 | 1 |
| UGC0007648 | Clust-UGC0004684 | 4933 | 5 | 1 | 0 | 1 |
| UGC0004684 | Clust-UGC0004684 | 7587 | 8 | 1 | 0 | 1 |
| UGC0004691 | Clust-UGC0004691 | 12347 | 80 | 1 | 0 | 1 |
| UGC0004711 | Clust-UGC0004711 | 4387 | 3 | 1 | 0 | 1 |
| UGC0013274 | Clust-UGC0004711 | 860 | 4 | 1 | 1 | 1 |
| UGC0004728 | Clust-UGC0004728 | 2496 | 6 | 1 | 0 | 1 |
| UGC0006441 | Clust-UGC0004728 | 374 | 1 | 0 | 0 | 0 |
| UGC0005413 | Clust-UGC0004728 | 1350 | 2 | 1 | 0 | 0 |
| UGC0011132 | Clust-UGC0004755 | 5506 | 10 | 1 | 0 | 1 |
| UGC0004755 | Clust-UGC0004755 | 446 | 1 | 0 | 0 | 0 |
| UGC0004761 | Clust-UGC0004761 | 222 | 1 | 0 | 0 | 0 |
| UGC0004768 | Clust-UGC0004768 | 1007 | 3 | 1 | 0 | 0 |
| UGC0004784 | Clust-UGC0004784 | 4675 | 2 | 1 | 0 | 1 |
| UGC0012386 | Clust-UGC0004792 | 7514 | 9 | 1 | 0 | 1 |
| UGC0004792 | Clust-UGC0004792 | 335 | 1 | 0 | 0 | 0 |
| UGC0010897 | Clust-UGC0004792 | 1922 | 5 | 1 | 0 | 1 |
| UGC0004810 | Clust-UGC0004810 | 18546 | 176 | 1 | 0 | 1 |
| UGC0004812 | Clust-UGC0004812 | 10462 | 44 | 1 | 0 | 1 |
| UGC0010092 | Clust-UGC0004812 | 3285 | 3 | 1 | 0 | 1 |
| UGC0006516 | Clust-UGC0004812 | 2869 | 1 | 0 | 0 | 0 |
| UGC0007361 | Clust-UGC0004812 | 1754 | 2 | 1 | 0 | 1 |
| UGC0009068 | Clust-UGC0004812 | 2531 | 3 | 1 | 0 | 1 |
| UGC0008681 | Clust-UGC0004812 | 443 | 1 | 0 | 0 | 0 |
| UGC0009592 | Clust-UGC0004812 | 2307 | 3 | 1 | 0 | 1 |
| UGC0007731 | Clust-UGC0004812 | 4247 | 4 | 1 | 0 | 1 |
| UGC0008194 | Clust-UGC0004812 | 4697 | 6 | 1 | 1 | 1 |
| UGC0006749 | Clust-UGC0004812 | 5487 | 6 | 1 | 0 | 1 |
| UGC0007373 | Clust-UGC0004812 | 2877 | 1 | 0 | 0 | 0 |
| UGC0006883 | Clust-UGC0004812 | 3134 | 3 | 1 | 0 | 1 |
| UGC0007139 | Clust-UGC0004812 | 3085 | 3 | 1 | 0 | 1 |
| UGC0006162 | Clust-UGC0004814 | 2800 | 1 | 0 | 0 | 0 |
| UGC0009565 | Clust-UGC0004814 | 2416 | 2 | 1 | 0 | 1 |
| UGC0009006 | Clust-UGC0004814 | 1553 | 1 | 0 | 0 | 0 |
| UGC0004814 | Clust-UGC0004814 | 17386 | 25 | 1 | 0 | 1 |
| UGC0008285 | Clust-UGC0004814 | 925 | 1 | 0 | 0 | 0 |
| UGC0006741 | Clust-UGC0004814 | 414 | 1 | 0 | 0 | 0 |
| UGC0010597 | Clust-UGC0004814 | 2436 | 1 | 0 | 0 | 0 |
| UGC0004819 | Clust-UGC0004819 | 14547 | 93 | 1 | 0 | 1 |
| UGC0012185 | Clust-UGC0004819 | 4305 | 3 | 1 | 0 | 1 |
| UGC0004820 | Clust-UGC0004820 | 7213 | 17 | 1 | 0 | 1 |
| UGC0012240 | Clust-UGC0004820 | 6063 | 12 | 1 | 1 | 1 |
| UGC0004833 | Clust-UGC0004833 | 14802 | 115 | 1 | 0 | 1 |
| UGC0008600 | Clust-UGC0004833 | 7706 | 23 | 1 | 0 | 1 |
| UGC0007755 | Clust-UGC0004833 | 6814 | 20 | 1 | 0 | 1 |
| UGC0004843 | Clust-UGC0004843 | 9033 | 17 | 1 | 0 | 1 |
| UGC0004959 | Clust-UGC0004959 | 5671 | 4 | 1 | 0 | 1 |
| UGC0009978 | Clust-UGC0004960 | 504 | 1 | 0 | 0 | 0 |
| UGC0008747 | Clust-UGC0004960 | 380 | 2 | 1 | 0 | 0 |
| UGC0009166 | Clust-UGC0004960 | 1122 | 2 | 1 | 0 | 0 |
| UGC0010631 | Clust-UGC0004960 | 1686 | 1 | 0 | 0 | 0 |
| UGC0010489 | Clust-UGC0004960 | 2621 | 1 | 0 | 0 | 0 |
| UGC0004960 | Clust-UGC0004960 | 3485 | 1 | 0 | 0 | 0 |
| UGC0007128 | Clust-UGC0004960 | 379 | 1 | 0 | 0 | 0 |
| UGC0007690 | Clust-UGC0004960 | 3612 | 2 | 1 | 0 | 0 |
| UGC0006878 | Clust-UGC0004960 | 3712 | 1 | 0 | 0 | 0 |
| UGC0006697 | Clust-UGC0004960 | 777 | 1 | 0 | 0 | 0 |
| UGC0011513 | Clust-UGC0004960 | 21831 | 46 | 1 | 0 | 1 |
| UGC0005942 | Clust-UGC0004960 | 788 | 1 | 0 | 0 | 0 |
| UGC0007313 | Clust-UGC0004960 | 6155 | 1 | 0 | 0 | 0 |
| UGC0013044 | Clust-UGC0004960 | 7292 | 9 | 1 | 1 | 1 |
| UGC0007154 | Clust-UGC0004960 | 2619 | 2 | 1 | 0 | 1 |

| | | | | | | |
|---|---|---|---|---|---|---|
| UGC0010531 | Clust-UGC0004960 | 4139 | 5 | 1 | 0 | 1 |
| UGC0005530 | Clust-UGC0004960 | 998 | 1 | 0 | 0 | 0 |
| UGC0004965 | Clust-UGC0004965 | 1827 | 6 | 1 | 0 | 1 |
| UGC0005124 | Clust-UGC0004965 | 1171 | 1 | 0 | 0 | 0 |
| UGC0004966 | Clust-UGC0004966 | 2554 | 3 | 1 | 0 | 1 |
| UGC0010503 | Clust-UGC0004966 | 2403 | 2 | 1 | 0 | 1 |
| UGC0008388 | Clust-UGC0004966 | 958 | 1 | 0 | 0 | 0 |
| UGC0005976 | Clust-UGC0004967 | 1910 | 1 | 0 | 0 | 0 |
| UGC0009611 | Clust-UGC0004967 | 532 | 1 | 0 | 0 | 0 |
| UGC0008006 | Clust-UGC0004967 | 799 | 1 | 0 | 0 | 0 |
| UGC0004967 | Clust-UGC0004967 | 671 | 1 | 0 | 0 | 0 |
| UGC0006315 | Clust-UGC0004967 | 2457 | 2 | 1 | 0 | 1 |
| UGC0007715 | Clust-UGC0004967 | 5155 | 4 | 1 | 0 | 1 |
| UGC0008543 | Clust-UGC0004967 | 268 | 1 | 0 | 0 | 0 |
| UGC0012640 | Clust-UGC0004967 | 5512 | 8 | 1 | 0 | 1 |
| UGC0008653 | Clust-UGC0004967 | 599 | 1 | 0 | 0 | 0 |
| UGC0006736 | Clust-UGC0004967 | 239 | 1 | 0 | 0 | 0 |
| UGC0008189 | Clust-UGC0004967 | 1732 | 2 | 1 | 0 | 1 |
| UGC0008373 | Clust-UGC0004967 | 197 | 1 | 0 | 0 | 0 |
| UGC0004971 | Clust-UGC0004971 | 6110 | 7 | 1 | 0 | 1 |
| UGC0004981 | Clust-UGC0004981 | 6875 | 13 | 1 | 0 | 1 |
| UGC0005569 | Clust-UGC0004981 | 3822 | 1 | 0 | 0 | 0 |
| UGC0010072 | Clust-UGC0004981 | 1670 | 1 | 0 | 0 | 0 |
| UGC0008364 | Clust-UGC0004981 | 988 | 1 | 0 | 0 | 0 |
| UGC0007172 | Clust-UGC0004987 | 4536 | 2 | 1 | 0 | 1 |
| UGC0004987 | Clust-UGC0004987 | 9777 | 32 | 1 | 0 | 1 |
| UGC0011482 | Clust-UGC0004992 | 4009 | 2 | 1 | 0 | 1 |
| UGC0005412 | Clust-UGC0004992 | 1342 | 1 | 0 | 0 | 0 |
| UGC0004992 | Clust-UGC0004992 | 4302 | 2 | 1 | 0 | 0 |
| UGC0006410 | Clust-UGC0004992 | 926 | 1 | 0 | 0 | 0 |
| UGC0006431 | Clust-UGC0004992 | 5759 | 3 | 1 | 0 | 0 |
| UGC0010108 | Clust-UGC0004992 | 10159 | 4 | 1 | 0 | 1 |
| UGC0005053 | Clust-UGC0005000 | 2037 | 2 | 1 | 0 | 0 |
| UGC0006126 | Clust-UGC0005000 | 421 | 1 | 0 | 0 | 0 |
| UGC0005000 | Clust-UGC0005000 | 7234 | 13 | 1 | 0 | 1 |
| UGC0010253 | Clust-UGC0005000 | 1138 | 2 | 1 | 0 | 1 |
| UGC0005005 | Clust-UGC0005005 | 7248 | 3 | 1 | 0 | 1 |
| UGC0005007 | Clust-UGC0005007 | 9686 | 40 | 1 | 0 | 1 |
| UGC0010139 | Clust-UGC0005010 | 6872 | 6 | 1 | 0 | 1 |
| UGC0005010 | Clust-UGC0005010 | 9875 | 32 | 1 | 0 | 1 |
| UGC0010121 | Clust-UGC0005010 | 2405 | 1 | 0 | 0 | 0 |
| UGC0005020 | Clust-UGC0005020 | 9787 | 30 | 1 | 0 | 1 |
| UGC0007265 | Clust-UGC0005022 | 401 | 1 | 0 | 0 | 0 |
| UGC0006058 | Clust-UGC0005022 | 321 | 1 | 0 | 0 | 0 |
| UGC0007000 | Clust-UGC0005022 | 4763 | 5 | 1 | 0 | 1 |
| UGC0007020 | Clust-UGC0005022 | 4338 | 7 | 1 | 1 | 1 |
| UGC0006750 | Clust-UGC0005022 | 6021 | 6 | 1 | 0 | 1 |
| UGC0005546 | Clust-UGC0005022 | 1231 | 1 | 0 | 0 | 0 |
| UGC0007736 | Clust-UGC0005022 | 5077 | 4 | 1 | 0 | 0 |
| UGC0007505 | Clust-UGC0005022 | 4813 | 5 | 1 | 0 | 1 |
| UGC0005667 | Clust-UGC0005022 | 2419 | 3 | 1 | 0 | 0 |
| UGC0007350 | Clust-UGC0005022 | 1039 | 2 | 1 | 0 | 0 |
| UGC0007131 | Clust-UGC0005022 | 179 | 1 | 0 | 0 | 0 |
| UGC0005022 | Clust-UGC0005022 | 6668 | 6 | 1 | 0 | 1 |
| UGC0007496 | Clust-UGC0005022 | 1408 | 2 | 1 | 1 | 0 |
| UGC0007728 | Clust-UGC0005022 | 2102 | 1 | 0 | 0 | 0 |
| UGC0007231 | Clust-UGC0005022 | 3290 | 2 | 1 | 0 | 0 |
| UGC0007741 | Clust-UGC0005022 | 6622 | 10 | 1 | 0 | 1 |
| UGC0010088 | Clust-UGC0005024 | 3639 | 5 | 1 | 0 | 1 |
| UGC0005024 | Clust-UGC0005024 | 5981 | 9 | 1 | 0 | 1 |
| UGC0008554 | Clust-UGC0005035 | 2865 | 3 | 1 | 0 | 1 |
| UGC0005035 | Clust-UGC0005035 | 2928 | 2 | 1 | 0 | 1 |
| UGC0006464 | Clust-UGC0005035 | 865 | 1 | 0 | 0 | 0 |
| UGC0006101 | Clust-UGC0005035 | 1770 | 2 | 1 | 0 | 1 |
| UGC0010116 | Clust-UGC0005035 | 208 | 1 | 0 | 0 | 0 |
| UGC0008794 | Clust-UGC0005035 | 3693 | 3 | 1 | 0 | 1 |
| UGC0009236 | Clust-UGC0005035 | 1652 | 3 | 1 | 0 | 1 |
| UGC0009617 | Clust-UGC0005035 | 988 | 1 | 0 | 0 | 0 |
| UGC0005071 | Clust-UGC0005071 | 3061 | 6 | 1 | 0 | 1 |
| UGC0005074 | Clust-UGC0005074 | 3347 | 1 | 0 | 0 | 0 |
| UGC0013260 | Clust-UGC0005074 | 4991 | 9 | 1 | 1 | 1 |
| UGC0006800 | Clust-UGC0005074 | 4650 | 6 | 1 | 0 | 1 |
| UGC0012177 | Clust-UGC0005075 | 4555 | 4 | 1 | 0 | 1 |
| UGC0009778 | Clust-UGC0005075 | 1274 | 1 | 0 | 0 | 0 |
| UGC0005075 | Clust-UGC0005075 | 1401 | 1 | 0 | 0 | 0 |
| UGC0009041 | Clust-UGC0005075 | 1762 | 2 | 1 | 0 | 1 |
| UGC0007711 | Clust-UGC0005075 | 5082 | 5 | 1 | 0 | 1 |
| UGC0006388 | Clust-UGC0005075 | 5775 | 10 | 1 | 0 | 1 |
| UGC0005076 | Clust-UGC0005076 | 3162 | 1 | 0 | 0 | 0 |

| | | | | | | |
|---|---|---|---|---|---|---|
| UGC0005784 | Clust-UGC0005087 | 5655 | 2 | 1 | 0 | 1 |
| UGC0005999 | Clust-UGC0005087 | 3158 | 1 | 0 | 0 | 0 |
| UGC0005408 | Clust-UGC0005087 | 5763 | 7 | 1 | 0 | 1 |
| UGC0011445 | Clust-UGC0005087 | 7312 | 17 | 1 | 0 | 1 |
| UGC0008320 | Clust-UGC0005087 | 1065 | 1 | 0 | 0 | 0 |
| UGC0005965 | Clust-UGC0005087 | 216 | 1 | 0 | 0 | 0 |
| UGC0005087 | Clust-UGC0005087 | 2527 | 1 | 0 | 0 | 0 |
| UGC0013115 | Clust-UGC0005087 | 6368 | 6 | 1 | 0 | 1 |
| UGC0005100 | Clust-UGC0005100 | 2304 | 1 | 0 | 0 | 0 |
| UGC0007977 | Clust-UGC0005100 | 824 | 1 | 0 | 0 | 0 |
| UGC0012856 | Clust-UGC0005100 | 167 | 1 | 0 | 0 | 0 |
| UGC0008904 | Clust-UGC0005100 | 1254 | 2 | 1 | 0 | 1 |
| UGC0008234 | Clust-UGC0005100 | 503 | 1 | 0 | 0 | 0 |
| UGC0005630 | Clust-UGC0005102 | 1434 | 1 | 0 | 0 | 0 |
| UGC0005102 | Clust-UGC0005102 | 2751 | 2 | 1 | 0 | 1 |
| UGC0008688 | Clust-UGC0005122 | 222 | 1 | 0 | 0 | 0 |
| UGC0012712 | Clust-UGC0005122 | 4987 | 13 | 1 | 1 | 1 |
| UGC0008044 | Clust-UGC0005122 | 50 | 1 | 0 | 0 | 0 |
| UGC0005122 | Clust-UGC0005122 | 2609 | 2 | 1 | 0 | 1 |
| UGC0008779 | Clust-UGC0005122 | 4490 | 5 | 1 | 0 | 1 |
| UGC0012852 | Clust-UGC0005122 | 288 | 1 | 0 | 0 | 0 |
| UGC0009052 | Clust-UGC0005122 | 755 | 1 | 0 | 0 | 0 |
| UGC0012396 | Clust-UGC0005122 | 2680 | 6 | 1 | 0 | 1 |
| UGC0009211 | Clust-UGC0005122 | 4608 | 5 | 1 | 0 | 1 |
| UGC0008423 | Clust-UGC0005122 | 847 | 1 | 0 | 0 | 0 |
| UGC0008235 | Clust-UGC0005122 | 627 | 1 | 0 | 0 | 0 |
| UGC0010093 | Clust-UGC0005321 | 2446 | 1 | 0 | 0 | 0 |
| UGC0010224 | Clust-UGC0005321 | 2548 | 2 | 1 | 1 | 0 |
| UGC0005940 | Clust-UGC0005321 | 5139 | 1 | 0 | 0 | 0 |
| UGC0005321 | Clust-UGC0005321 | 638 | 1 | 0 | 0 | 0 |
| UGC0007960 | Clust-UGC0005321 | 6267 | 14 | 1 | 0 | 1 |
| UGC0009336 | Clust-UGC0005321 | 10106 | 58 | 1 | 0 | 1 |
| UGC0005679 | Clust-UGC0005323 | 2161 | 2 | 1 | 0 | 0 |
| UGC0009564 | Clust-UGC0005323 | 3715 | 2 | 1 | 0 | 0 |
| UGC0005323 | Clust-UGC0005323 | 1448 | 2 | 1 | 0 | 0 |
| UGC0005528 | Clust-UGC0005340 | 1895 | 1 | 0 | 0 | 0 |
| UGC0005340 | Clust-UGC0005340 | 2785 | 1 | 0 | 0 | 0 |
| UGC0011486 | Clust-UGC0005342 | 4117 | 7 | 1 | 0 | 1 |
| UGC0012288 | Clust-UGC0005342 | 2528 | 1 | 0 | 0 | 0 |
| UGC0006511 | Clust-UGC0005342 | 1829 | 1 | 0 | 0 | 0 |
| UGC0007145 | Clust-UGC0005342 | 481 | 1 | 0 | 0 | 0 |
| UGC0008500 | Clust-UGC0005342 | 1699 | 1 | 0 | 0 | 0 |
| UGC0013358 | Clust-UGC0005342 | 5583 | 9 | 1 | 0 | 1 |
| UGC0006993 | Clust-UGC0005342 | 3114 | 2 | 1 | 0 | 1 |
| UGC0007975 | Clust-UGC0005342 | 1421 | 2 | 1 | 0 | 1 |
| UGC0007277 | Clust-UGC0005342 | 3926 | 2 | 1 | 0 | 1 |
| UGC0006707 | Clust-UGC0005342 | 1734 | 3 | 1 | 0 | 0 |
| UGC0005342 | Clust-UGC0005342 | 1166 | 1 | 0 | 0 | 0 |
| UGC0006900 | Clust-UGC0005342 | 2618 | 1 | 0 | 0 | 0 |
| UGC0008735 | Clust-UGC0005342 | 1018 | 1 | 0 | 0 | 0 |
| UGC0007995 | Clust-UGC0005342 | 362 | 1 | 0 | 0 | 0 |
| UGC0010556 | Clust-UGC0005342 | 1965 | 1 | 0 | 0 | 0 |
| UGC0008050 | Clust-UGC0005342 | 1953 | 4 | 1 | 0 | 1 |
| UGC0010127 | Clust-UGC0005342 | 2856 | 3 | 1 | 0 | 1 |
| UGC0005358 | Clust-UGC0005358 | 5383 | 7 | 1 | 0 | 1 |
| UGC0005933 | Clust-UGC0005358 | 1960 | 2 | 1 | 0 | 0 |
| UGC0007584 | Clust-UGC0005358 | 4542 | 6 | 1 | 0 | 1 |
| UGC0007327 | Clust-UGC0005360 | 824 | 1 | 0 | 0 | 0 |
| UGC0005360 | Clust-UGC0005360 | 2202 | 1 | 0 | 0 | 0 |
| UGC0005369 | Clust-UGC0005369 | 2420 | 4 | 1 | 0 | 1 |
| UGC0005379 | Clust-UGC0005379 | 2717 | 2 | 1 | 0 | 1 |
| UGC0005730 | Clust-UGC0005379 | 6269 | 15 | 1 | 0 | 1 |
| UGC0005381 | Clust-UGC0005381 | 2934 | 3 | 1 | 0 | 1 |
| UGC0008245 | Clust-UGC0005381 | 1749 | 2 | 1 | 0 | 0 |
| UGC0005384 | Clust-UGC0005384 | 2454 | 3 | 1 | 0 | 1 |
| UGC0013084 | Clust-UGC0005384 | 7617 | 29 | 1 | 0 | 1 |
| UGC0005388 | Clust-UGC0005388 | 478 | 1 | 0 | 0 | 0 |
| UGC0006694 | Clust-UGC0005388 | 1083 | 1 | 0 | 0 | 0 |
| UGC0008487 | Clust-UGC0005402 | 592 | 1 | 0 | 0 | 0 |
| UGC0005652 | Clust-UGC0005402 | 2100 | 1 | 0 | 0 | 0 |
| UGC0005402 | Clust-UGC0005402 | 2636 | 2 | 1 | 0 | 1 |
| UGC0008306 | Clust-UGC0005402 | 591 | 1 | 0 | 0 | 0 |
| UGC0005517 | Clust-UGC0005517 | 1272 | 1 | 0 | 0 | 0 |
| UGC0007654 | Clust-UGC0005524 | 10907 | 70 | 1 | 0 | 1 |
| UGC0005524 | Clust-UGC0005524 | 1433 | 1 | 0 | 0 | 0 |
| UGC0010149 | Clust-UGC0005573 | 274 | 1 | 0 | 0 | 0 |
| UGC0010619 | Clust-UGC0005573 | 1804 | 1 | 0 | 0 | 0 |
| UGC0008269 | Clust-UGC0005573 | 982 | 1 | 0 | 0 | 0 |
| UGC0013043 | Clust-UGC0005573 | 3103 | 2 | 1 | 1 | 0 |

| | | | | | | |
|---|---|---|---|---|---|---|
| UGC0005573 | Clust-UGC0005573 | 1931 | 1 | 0 | 0 | 0 |
| UGC0012229 | Clust-UGC0005573 | 4195 | 2 | 1 | 0 | 1 |
| UGC0005580 | Clust-UGC0005580 | 4880 | 4 | 1 | 0 | 0 |
| UGC0005594 | Clust-UGC0005594 | 1755 | 1 | 0 | 0 | 0 |
| UGC0005957 | Clust-UGC0005594 | 561 | 1 | 0 | 0 | 0 |
| UGC0009574 | Clust-UGC0005609 | 6643 | 21 | 1 | 0 | 1 |
| UGC0005609 | Clust-UGC0005609 | 6266 | 23 | 1 | 0 | 1 |
| UGC0005894 | Clust-UGC0005610 | 81 | 1 | 0 | 0 | 0 |
| UGC0005610 | Clust-UGC0005610 | 1729 | 1 | 0 | 0 | 0 |
| UGC0006475 | Clust-UGC0005610 | 2025 | 1 | 0 | 0 | 0 |
| UGC0013283 | Clust-UGC0005610 | 1033 | 1 | 0 | 0 | 0 |
| UGC0010485 | Clust-UGC0005623 | 2962 | 4 | 1 | 0 | 1 |
| UGC0010179 | Clust-UGC0005623 | 1341 | 2 | 1 | 1 | 0 |
| UGC0005623 | Clust-UGC0005623 | 2848 | 2 | 1 | 0 | 0 |
| UGC0005625 | Clust-UGC0005625 | 151 | 1 | 0 | 0 | 0 |
| UGC0012688 | Clust-UGC0005627 | 5427 | 8 | 1 | 1 | 1 |
| UGC0005627 | Clust-UGC0005627 | 2778 | 2 | 1 | 0 | 1 |
| UGC0005640 | Clust-UGC0005640 | 2315 | 1 | 0 | 0 | 0 |
| UGC0005646 | Clust-UGC0005646 | 985 | 1 | 0 | 0 | 0 |
| UGC0010256 | Clust-UGC0005646 | 187 | 1 | 0 | 0 | 0 |
| UGC0007985 | Clust-UGC0005647 | 73 | 1 | 0 | 0 | 0 |
| UGC0007184 | Clust-UGC0005647 | 3372 | 2 | 1 | 0 | 1 |
| UGC0007387 | Clust-UGC0005647 | 4537 | 3 | 1 | 0 | 1 |
| UGC0005647 | Clust-UGC0005647 | 3121 | 1 | 0 | 0 | 0 |
| UGC0007256 | Clust-UGC0005647 | 5857 | 9 | 1 | 0 | 1 |
| UGC0008565 | Clust-UGC0005647 | 2316 | 2 | 1 | 0 | 1 |
| UGC0007150 | Clust-UGC0005647 | 4573 | 4 | 1 | 0 | 1 |
| UGC0008662 | Clust-UGC0005647 | 415 | 1 | 0 | 0 | 0 |
| UGC0009419 | Clust-UGC0005648 | 1202 | 1 | 0 | 0 | 0 |
| UGC0005648 | Clust-UGC0005648 | 1086 | 2 | 1 | 1 | 0 |
| UGC0005650 | Clust-UGC0005650 | 1826 | 2 | 1 | 0 | 1 |
| UGC0005666 | Clust-UGC0005666 | 2360 | 8 | 1 | 0 | 1 |
| UGC0005671 | Clust-UGC0005671 | 2020 | 1 | 0 | 0 | 0 |
| UGC0010530 | Clust-UGC0005671 | 1629 | 1 | 0 | 0 | 0 |
| UGC0008293 | Clust-UGC0005673 | 2893 | 1 | 0 | 0 | 0 |
| UGC0005673 | Clust-UGC0005673 | 2609 | 1 | 0 | 0 | 0 |
| UGC0005681 | Clust-UGC0005681 | 1611 | 2 | 1 | 0 | 1 |
| UGC0012659 | Clust-UGC0005681 | 1063 | 2 | 1 | 0 | 0 |
| UGC0006548 | Clust-UGC0005681 | 2060 | 1 | 0 | 0 | 0 |
| UGC0005721 | Clust-UGC0005721 | 4739 | 3 | 1 | 0 | 1 |
| UGC0005990 | Clust-UGC0005726 | 2778 | 2 | 1 | 0 | 1 |
| UGC0005726 | Clust-UGC0005726 | 359 | 1 | 0 | 0 | 0 |
| UGC0005727 | Clust-UGC0005727 | 6908 | 19 | 1 | 0 | 1 |
| UGC0005732 | Clust-UGC0005732 | 9187 | 40 | 1 | 0 | 1 |
| UGC0005743 | Clust-UGC0005743 | 7566 | 21 | 1 | 0 | 1 |
| UGC0005747 | Clust-UGC0005747 | 5389 | 5 | 1 | 0 | 1 |
| UGC0006455 | Clust-UGC0005801 | 5253 | 2 | 1 | 0 | 1 |
| UGC0006072 | Clust-UGC0005801 | 2442 | 1 | 0 | 0 | 0 |
| UGC0012325 | Clust-UGC0005801 | 2243 | 1 | 0 | 0 | 0 |
| UGC0005899 | Clust-UGC0005801 | 225 | 1 | 0 | 0 | 0 |
| UGC0006444 | Clust-UGC0005801 | 1461 | 1 | 0 | 0 | 0 |
| UGC0005905 | Clust-UGC0005801 | 4255 | 2 | 1 | 0 | 1 |
| UGC0011636 | Clust-UGC0005801 | 2413 | 2 | 1 | 0 | 0 |
| UGC0006913 | Clust-UGC0005801 | 1930 | 1 | 0 | 0 | 0 |
| UGC0009065 | Clust-UGC0005801 | 2245 | 1 | 0 | 0 | 0 |
| UGC0009474 | Clust-UGC0005801 | 3373 | 1 | 0 | 0 | 0 |
| UGC0012738 | Clust-UGC0005801 | 151 | 1 | 0 | 0 | 0 |
| UGC0011591 | Clust-UGC0005801 | 925 | 2 | 1 | 0 | 0 |
| UGC0009602 | Clust-UGC0005801 | 1321 | 1 | 0 | 0 | 0 |
| UGC0012270 | Clust-UGC0005801 | 1826 | 2 | 1 | 0 | 1 |
| UGC0006333 | Clust-UGC0005801 | 5605 | 1 | 0 | 0 | 0 |
| UGC0012046 | Clust-UGC0005801 | 203 | 1 | 0 | 0 | 0 |
| UGC0005801 | Clust-UGC0005801 | 4201 | 2 | 1 | 1 | 0 |
| UGC0012180 | Clust-UGC0005801 | 1116 | 1 | 0 | 0 | 0 |
| UGC0010941 | Clust-UGC0005801 | 4616 | 3 | 1 | 1 | 1 |
| UGC0006010 | Clust-UGC0005801 | 2563 | 1 | 0 | 0 | 0 |
| UGC0009725 | Clust-UGC0005801 | 1455 | 1 | 0 | 0 | 0 |
| UGC0007585 | Clust-UGC0005801 | 1533 | 1 | 0 | 0 | 0 |
| UGC0006446 | Clust-UGC0005801 | 6387 | 2 | 1 | 0 | 1 |
| UGC0012169 | Clust-UGC0005801 | 2317 | 1 | 0 | 0 | 0 |
| UGC0011669 | Clust-UGC0005801 | 1708 | 1 | 0 | 0 | 0 |
| UGC0011472 | Clust-UGC0005801 | 2067 | 2 | 1 | 0 | 0 |
| UGC0006595 | Clust-UGC0005801 | 405 | 1 | 0 | 0 | 0 |
| UGC0010817 | Clust-UGC0005801 | 1870 | 5 | 1 | 0 | 0 |
| UGC0009423 | Clust-UGC0005801 | 2364 | 2 | 1 | 0 | 0 |
| UGC0012997 | Clust-UGC0005801 | 2648 | 1 | 0 | 0 | 0 |
| UGC0011968 | Clust-UGC0005801 | 2990 | 2 | 1 | 0 | 1 |
| UGC0011197 | Clust-UGC0005801 | 3843 | 5 | 1 | 0 | 1 |
| UGC0005806 | Clust-UGC0005806 | 1513 | 1 | 0 | 0 | 0 |

| | | | | | | |
|---|---|---|---|---|---|---|
| UGC0010507 | Clust-UGC0005807 | 2902 | 3 | 1 | 0 | 0 |
| UGC0005807 | Clust-UGC0005807 | 1114 | 1 | 0 | 0 | 0 |
| UGC0012137 | Clust-UGC0005812 | 5537 | 8 | 1 | 0 | 1 |
| UGC0007325 | Clust-UGC0005812 | 1672 | 1 | 0 | 0 | 0 |
| UGC0009228 | Clust-UGC0005812 | 8729 | 45 | 1 | 0 | 1 |
| UGC0007091 | Clust-UGC0005812 | 544 | 1 | 0 | 0 | 0 |
| UGC0010457 | Clust-UGC0005812 | 5178 | 6 | 1 | 0 | 1 |
| UGC0009408 | Clust-UGC0005812 | 4935 | 9 | 1 | 0 | 1 |
| UGC0005812 | Clust-UGC0005812 | 4624 | 12 | 1 | 0 | 1 |
| UGC0012737 | Clust-UGC0005815 | 1823 | 2 | 1 | 0 | 1 |
| UGC0012717 | Clust-UGC0005815 | 8156 | 33 | 1 | 0 | 1 |
| UGC0012226 | Clust-UGC0005815 | 6224 | 11 | 1 | 0 | 1 |
| UGC0005815 | Clust-UGC0005815 | 474 | 1 | 0 | 0 | 0 |
| UGC0013386 | Clust-UGC0005816 | 165 | 1 | 0 | 0 | 0 |
| UGC0009281 | Clust-UGC0005816 | 405 | 1 | 0 | 0 | 0 |
| UGC0008973 | Clust-UGC0005816 | 1071 | 1 | 0 | 0 | 0 |
| UGC0006036 | Clust-UGC0005816 | 1509 | 2 | 1 | 0 | 1 |
| UGC0009183 | Clust-UGC0005816 | 2011 | 4 | 1 | 0 | 1 |
| UGC0009533 | Clust-UGC0005816 | 498 | 2 | 1 | 1 | 0 |
| UGC0006283 | Clust-UGC0005816 | 1225 | 2 | 1 | 0 | 1 |
| UGC0006050 | Clust-UGC0005816 | 2317 | 2 | 1 | 0 | 1 |
| UGC0007994 | Clust-UGC0005816 | 63 | 1 | 0 | 0 | 0 |
| UGC0011018 | Clust-UGC0005816 | 3221 | 4 | 1 | 0 | 1 |
| UGC0009057 | Clust-UGC0005816 | 632 | 1 | 0 | 0 | 0 |
| UGC0011810 | Clust-UGC0005816 | 1017 | 1 | 0 | 0 | 0 |
| UGC0012242 | Clust-UGC0005816 | 3972 | 5 | 1 | 0 | 1 |
| UGC0010976 | Clust-UGC0005816 | 2393 | 2 | 1 | 0 | 1 |
| UGC0012228 | Clust-UGC0005816 | 11803 | 78 | 1 | 1 | 1 |
| UGC0005816 | Clust-UGC0005816 | 3063 | 4 | 1 | 0 | 1 |
| UGC0006377 | Clust-UGC0005816 | 3816 | 4 | 1 | 0 | 1 |
| UGC0005822 | Clust-UGC0005822 | 1875 | 1 | 0 | 0 | 0 |
| UGC0007593 | Clust-UGC0005824 | 4620 | 6 | 1 | 1 | 1 |
| UGC0006834 | Clust-UGC0005824 | 6724 | 22 | 1 | 1 | 1 |
| UGC0005824 | Clust-UGC0005824 | 6015 | 17 | 1 | 0 | 1 |
| UGC0013268 | Clust-UGC0005853 | 12963 | 91 | 1 | 0 | 1 |
| UGC0012743 | Clust-UGC0005853 | 1696 | 3 | 1 | 0 | 1 |
| UGC0005853 | Clust-UGC0005853 | 3431 | 4 | 1 | 0 | 1 |
| UGC0010125 | Clust-UGC0005853 | 10993 | 29 | 1 | 1 | 1 |
| UGC0009503 | Clust-UGC0005863 | 2851 | 2 | 1 | 0 | 0 |
| UGC0006035 | Clust-UGC0005863 | 4256 | 4 | 1 | 0 | 1 |
| UGC0013057 | Clust-UGC0005863 | 10283 | 15 | 1 | 0 | 1 |
| UGC0005863 | Clust-UGC0005863 | 1593 | 1 | 0 | 0 | 0 |
| UGC0011559 | Clust-UGC0005863 | 5100 | 5 | 1 | 0 | 1 |
| UGC0011608 | Clust-UGC0005863 | 3538 | 3 | 1 | 0 | 1 |
| UGC0012957 | Clust-UGC0005863 | 304 | 2 | 1 | 1 | 0 |
| UGC0011587 | Clust-UGC0005863 | 761 | 1 | 0 | 0 | 0 |
| UGC0006390 | Clust-UGC0005863 | 6422 | 6 | 1 | 0 | 1 |
| UGC0006137 | Clust-UGC0005863 | 711 | 1 | 0 | 0 | 0 |
| UGC0011008 | Clust-UGC0005882 | 6078 | 8 | 1 | 0 | 1 |
| UGC0005882 | Clust-UGC0005882 | 2945 | 3 | 1 | 0 | 1 |
| UGC0008612 | Clust-UGC0005882 | 3410 | 6 | 1 | 0 | 1 |
| UGC0007762 | Clust-UGC0005882 | 6514 | 12 | 1 | 0 | 1 |
| UGC0008350 | Clust-UGC0005882 | 2173 | 3 | 1 | 0 | 0 |
| UGC0013138 | Clust-UGC0005882 | 1406 | 2 | 1 | 0 | 1 |
| UGC0007447 | Clust-UGC0005886 | 1619 | 1 | 0 | 0 | 0 |
| UGC0011849 | Clust-UGC0005886 | 2696 | 1 | 0 | 0 | 0 |
| UGC0005886 | Clust-UGC0005886 | 1606 | 1 | 0 | 0 | 0 |
| UGC0013192 | Clust-UGC0005900 | 1654 | 3 | 1 | 0 | 1 |
| UGC0005900 | Clust-UGC0005900 | 2788 | 4 | 1 | 0 | 1 |
| UGC0008947 | Clust-UGC0005900 | 1616 | 2 | 1 | 0 | 1 |
| UGC0006285 | Clust-UGC0005900 | 6039 | 5 | 1 | 0 | 1 |
| UGC0006330 | Clust-UGC0005908 | 4921 | 7 | 1 | 0 | 1 |
| UGC0009772 | Clust-UGC0005908 | 3897 | 6 | 1 | 0 | 1 |
| UGC0005908 | Clust-UGC0005908 | 2403 | 4 | 1 | 0 | 1 |
| UGC0007482 | Clust-UGC0005908 | 3444 | 5 | 1 | 0 | 1 |
| UGC0012757 | Clust-UGC0005908 | 2644 | 4 | 1 | 0 | 1 |
| UGC0005928 | Clust-UGC0005908 | 2203 | 4 | 1 | 0 | 1 |
| UGC0006656 | Clust-UGC0005908 | 6854 | 14 | 1 | 0 | 1 |
| UGC0013297 | Clust-UGC0005908 | 2992 | 4 | 1 | 0 | 1 |
| UGC0009071 | Clust-UGC0005908 | 1399 | 3 | 1 | 0 | 1 |
| UGC0006995 | Clust-UGC0005908 | 4755 | 7 | 1 | 0 | 1 |
| UGC0011782 | Clust-UGC0005920 | 992 | 1 | 0 | 0 | 0 |
| UGC0005920 | Clust-UGC0005920 | 3319 | 7 | 1 | 0 | 1 |
| UGC0009378 | Clust-UGC0005920 | 4586 | 8 | 1 | 0 | 1 |
| UGC0009119 | Clust-UGC0005920 | 601 | 1 | 0 | 0 | 0 |
| UGC0013168 | Clust-UGC0005948 | 517 | 1 | 0 | 0 | 0 |
| UGC0005948 | Clust-UGC0005948 | 1750 | 1 | 0 | 0 | 0 |
| UGC0011799 | Clust-UGC0005960 | 1096 | 3 | 1 | 0 | 0 |
| UGC0006910 | Clust-UGC0005960 | 2049 | 1 | 0 | 0 | 0 |

| | | | | | | |
|---|---|---|---|---|---|---|
| UGC0005960 | Clust-UGC0005960 | 2543 | 2 | 1 | 0 | 1 |
| UGC0007553 | Clust-UGC0005960 | 4835 | 1 | 0 | 0 | 0 |
| UGC0005982 | Clust-UGC0005982 | 83 | 1 | 0 | 0 | 0 |
| UGC0005983 | Clust-UGC0005983 | 786 | 1 | 0 | 0 | 0 |
| UGC0012727 | Clust-UGC0005983 | 3825 | 3 | 1 | 0 | 0 |
| UGC0013037 | Clust-UGC0005984 | 5580 | 10 | 1 | 0 | 1 |
| UGC0011148 | Clust-UGC0005984 | 1470 | 4 | 1 | 0 | 1 |
| UGC0007660 | Clust-UGC0005984 | 6563 | 10 | 1 | 0 | 1 |
| UGC0005984 | Clust-UGC0005984 | 1328 | 1 | 0 | 0 | 0 |
| UGC0012165 | Clust-UGC0005984 | 3590 | 3 | 1 | 0 | 1 |
| UGC0007252 | Clust-UGC0005984 | 2016 | 1 | 0 | 0 | 0 |
| UGC0006418 | Clust-UGC0005984 | 155 | 1 | 0 | 0 | 0 |
| UGC0013217 | Clust-UGC0005984 | 1390 | 1 | 0 | 0 | 0 |
| UGC0006969 | Clust-UGC0005984 | 5552 | 6 | 1 | 0 | 1 |
| UGC0006394 | Clust-UGC0005984 | 2306 | 2 | 1 | 0 | 1 |
| UGC0005993 | Clust-UGC0005993 | 5838 | 7 | 1 | 0 | 1 |
| UGC0012713 | Clust-UGC0005993 | 10528 | 58 | 1 | 1 | 1 |
| UGC0006009 | Clust-UGC0006009 | 2779 | 1 | 0 | 0 | 0 |
| UGC0006014 | Clust-UGC0006014 | 4330 | 5 | 1 | 0 | 1 |
| UGC0012756 | Clust-UGC0006021 | 2633 | 8 | 1 | 0 | 1 |
| UGC0012827 | Clust-UGC0006021 | 5992 | 16 | 1 | 0 | 1 |
| UGC0012900 | Clust-UGC0006021 | 8682 | 35 | 1 | 0 | 1 |
| UGC0006021 | Clust-UGC0006021 | 943 | 1 | 0 | 0 | 0 |
| UGC0006487 | Clust-UGC0006029 | 1380 | 1 | 0 | 0 | 0 |
| UGC0013051 | Clust-UGC0006029 | 3026 | 3 | 1 | 1 | 0 |
| UGC0006029 | Clust-UGC0006029 | 504 | 1 | 0 | 0 | 0 |
| UGC0006780 | Clust-UGC0006029 | 447 | 1 | 0 | 0 | 0 |
| UGC0010312 | Clust-UGC0006043 | 1152 | 1 | 0 | 0 | 0 |
| UGC0006043 | Clust-UGC0006043 | 2820 | 1 | 0 | 0 | 0 |
| UGC0010958 | Clust-UGC0006043 | 743 | 2 | 1 | 0 | 0 |
| UGC0012759 | Clust-UGC0006047 | 1058 | 1 | 0 | 0 | 0 |
| UGC0012001 | Clust-UGC0006047 | 1132 | 1 | 0 | 0 | 0 |
| UGC0010332 | Clust-UGC0006047 | 132 | 1 | 0 | 0 | 0 |
| UGC0006047 | Clust-UGC0006047 | 614 | 1 | 0 | 0 | 0 |
| UGC0012162 | Clust-UGC0006047 | 496 | 1 | 0 | 0 | 0 |
| UGC0007640 | Clust-UGC0006047 | 5123 | 3 | 1 | 0 | 1 |
| UGC0010534 | Clust-UGC0006047 | 6475 | 12 | 1 | 0 | 1 |
| UGC0008337 | Clust-UGC0006047 | 526 | 1 | 0 | 0 | 0 |
| UGC0006062 | Clust-UGC0006062 | 319 | 1 | 0 | 0 | 0 |
| UGC0009047 | Clust-UGC0006062 | 253 | 1 | 0 | 0 | 0 |
| UGC0006071 | Clust-UGC0006071 | 1107 | 1 | 0 | 0 | 0 |
| UGC0006079 | Clust-UGC0006079 | 1346 | 1 | 0 | 0 | 0 |
| UGC0006094 | Clust-UGC0006094 | 1472 | 1 | 0 | 0 | 0 |
| UGC0009412 | Clust-UGC0006095 | 4932 | 4 | 1 | 0 | 1 |
| UGC0011175 | Clust-UGC0006095 | 1035 | 2 | 1 | 1 | 0 |
| UGC0011153 | Clust-UGC0006095 | 760 | 1 | 0 | 0 | 0 |
| UGC0012117 | Clust-UGC0006095 | 2411 | 4 | 1 | 0 | 1 |
| UGC0006471 | Clust-UGC0006095 | 1540 | 1 | 0 | 0 | 0 |
| UGC0006095 | Clust-UGC0006095 | 1246 | 1 | 0 | 0 | 0 |
| UGC0010925 | Clust-UGC0006095 | 1720 | 1 | 0 | 0 | 0 |
| UGC0012931 | Clust-UGC0006095 | 3647 | 7 | 1 | 0 | 1 |
| UGC0012940 | Clust-UGC0006095 | 2987 | 4 | 1 | 0 | 1 |
| UGC0012163 | Clust-UGC0006095 | 2470 | 3 | 1 | 0 | 1 |
| UGC0011128 | Clust-UGC0006095 | 1887 | 1 | 0 | 0 | 0 |
| UGC0013279 | Clust-UGC0006095 | 427 | 1 | 0 | 0 | 0 |
| UGC0007431 | Clust-UGC0006095 | 2773 | 4 | 1 | 0 | 1 |
| UGC0012972 | Clust-UGC0006095 | 3754 | 7 | 1 | 0 | 1 |
| UGC0007494 | Clust-UGC0006095 | 1141 | 1 | 0 | 0 | 0 |
| UGC0006732 | Clust-UGC0006095 | 3762 | 3 | 1 | 0 | 0 |
| UGC0007514 | Clust-UGC0006095 | 6311 | 12 | 1 | 0 | 1 |
| UGC0009634 | Clust-UGC0006095 | 3127 | 4 | 1 | 0 | 1 |
| UGC0007574 | Clust-UGC0006095 | 6294 | 9 | 1 | 0 | 1 |
| UGC0012996 | Clust-UGC0006095 | 3682 | 8 | 1 | 0 | 1 |
| UGC0006108 | Clust-UGC0006108 | 1426 | 2 | 1 | 0 | 1 |
| UGC0006128 | Clust-UGC0006128 | 749 | 1 | 0 | 0 | 0 |
| UGC0006129 | Clust-UGC0006129 | 1078 | 1 | 0 | 0 | 0 |
| UGC0006141 | Clust-UGC0006141 | 2399 | 1 | 0 | 0 | 0 |
| UGC0006166 | Clust-UGC0006166 | 415 | 2 | 1 | 1 | 0 |
| UGC0006294 | Clust-UGC0006294 | 3497 | 4 | 1 | 0 | 1 |
| UGC0006295 | Clust-UGC0006295 | 7930 | 20 | 1 | 0 | 1 |
| UGC0006298 | Clust-UGC0006298 | 1812 | 3 | 1 | 1 | 1 |
| UGC0006321 | Clust-UGC0006321 | 1517 | 2 | 1 | 0 | 1 |
| UGC0011568 | Clust-UGC0006323 | 1993 | 1 | 0 | 0 | 0 |
| UGC0013313 | Clust-UGC0006323 | 3454 | 4 | 1 | 0 | 1 |
| UGC0006365 | Clust-UGC0006323 | 2732 | 2 | 1 | 0 | 1 |
| UGC0006323 | Clust-UGC0006323 | 4119 | 3 | 1 | 0 | 1 |
| UGC0013060 | Clust-UGC0006323 | 1367 | 1 | 0 | 0 | 0 |
| UGC0006477 | Clust-UGC0006323 | 3983 | 1 | 0 | 0 | 0 |
| UGC0007704 | Clust-UGC0006323 | 5487 | 4 | 1 | 0 | 1 |

| | | | | | | |
|---|---|---|---|---|---|---|
| UGC0008358 | Clust-UGC0006323 | 472 | 1 | 0 | 0 | 0 |
| UGC0012157 | Clust-UGC0006340 | 5726 | 2 | 1 | 0 | 0 |
| UGC0006462 | Clust-UGC0006340 | 3689 | 3 | 1 | 0 | 1 |
| UGC0006340 | Clust-UGC0006340 | 4658 | 2 | 1 | 0 | 1 |
| UGC0010815 | Clust-UGC0006340 | 975 | 4 | 1 | 0 | 0 |
| UGC0006344 | Clust-UGC0006344 | 5979 | 10 | 1 | 0 | 1 |
| UGC0010078 | Clust-UGC0006346 | 5466 | 7 | 1 | 1 | 1 |
| UGC0007957 | Clust-UGC0006346 | 702 | 1 | 0 | 0 | 0 |
| UGC0006346 | Clust-UGC0006346 | 4221 | 3 | 1 | 1 | 1 |
| UGC0007121 | Clust-UGC0006351 | 2084 | 5 | 1 | 0 | 1 |
| UGC0006351 | Clust-UGC0006351 | 4450 | 9 | 1 | 0 | 1 |
| UGC0006366 | Clust-UGC0006366 | 5012 | 1 | 0 | 0 | 0 |
| UGC0006493 | Clust-UGC0006370 | 1634 | 2 | 1 | 0 | 1 |
| UGC0009979 | Clust-UGC0006370 | 1821 | 2 | 1 | 0 | 1 |
| UGC0006370 | Clust-UGC0006370 | 7048 | 15 | 1 | 0 | 1 |
| UGC0012592 | Clust-UGC0006381 | 2017 | 2 | 1 | 0 | 0 |
| UGC0006381 | Clust-UGC0006381 | 4688 | 2 | 1 | 0 | 1 |
| UGC0012882 | Clust-UGC0006385 | 7322 | 20 | 1 | 1 | 1 |
| UGC0006385 | Clust-UGC0006385 | 4786 | 9 | 1 | 0 | 1 |
| UGC0010554 | Clust-UGC0006385 | 3847 | 2 | 1 | 0 | 0 |
| UGC0006389 | Clust-UGC0006389 | 6143 | 18 | 1 | 0 | 1 |
| UGC0006404 | Clust-UGC0006404 | 8029 | 4 | 1 | 0 | 1 |
| UGC0006425 | Clust-UGC0006425 | 6628 | 15 | 1 | 0 | 1 |
| UGC0006428 | Clust-UGC0006428 | 3430 | 3 | 1 | 0 | 1 |
| UGC0006451 | Clust-UGC0006451 | 7629 | 19 | 1 | 0 | 1 |
| UGC0007208 | Clust-UGC0006483 | 1554 | 1 | 0 | 0 | 0 |
| UGC0007284 | Clust-UGC0006483 | 6558 | 3 | 1 | 0 | 1 |
| UGC0006483 | Clust-UGC0006483 | 2433 | 1 | 0 | 0 | 0 |
| UGC0007346 | Clust-UGC0006483 | 3320 | 2 | 1 | 1 | 0 |
| UGC0006936 | Clust-UGC0006483 | 6023 | 6 | 1 | 1 | 1 |
| UGC0007215 | Clust-UGC0006483 | 2357 | 1 | 0 | 0 | 0 |
| UGC0007338 | Clust-UGC0006483 | 2372 | 1 | 0 | 0 | 0 |
| UGC0006884 | Clust-UGC0006483 | 6833 | 2 | 1 | 0 | 1 |
| UGC0006889 | Clust-UGC0006483 | 4841 | 4 | 1 | 0 | 1 |
| UGC0007398 | Clust-UGC0006483 | 1796 | 1 | 0 | 0 | 0 |
| UGC0006485 | Clust-UGC0006485 | 1199 | 1 | 0 | 0 | 0 |
| UGC0009710 | Clust-UGC0006497 | 2957 | 4 | 1 | 0 | 1 |
| UGC0007166 | Clust-UGC0006497 | 5422 | 7 | 1 | 1 | 1 |
| UGC0009261 | Clust-UGC0006497 | 13291 | 90 | 1 | 0 | 1 |
| UGC0007467 | Clust-UGC0006497 | 748 | 1 | 0 | 0 | 0 |
| UGC0008322 | Clust-UGC0006497 | 53 | 1 | 0 | 0 | 0 |
| UGC0009181 | Clust-UGC0006497 | 4835 | 6 | 1 | 0 | 1 |
| UGC0006497 | Clust-UGC0006497 | 564 | 1 | 0 | 0 | 0 |
| UGC0009234 | Clust-UGC0006497 | 9606 | 41 | 1 | 0 | 1 |
| UGC0006515 | Clust-UGC0006515 | 609 | 1 | 0 | 0 | 0 |
| UGC0006532 | Clust-UGC0006532 | 5207 | 3 | 1 | 0 | 1 |
| UGC0007726 | Clust-UGC0006532 | 8151 | 14 | 1 | 0 | 1 |
| UGC0011628 | Clust-UGC0006532 | 2276 | 4 | 1 | 0 | 1 |
| UGC0008706 | Clust-UGC0006532 | 448 | 1 | 0 | 0 | 0 |
| UGC0012577 | Clust-UGC0006533 | 7118 | 21 | 1 | 0 | 1 |
| UGC0006533 | Clust-UGC0006533 | 1397 | 1 | 0 | 0 | 0 |
| UGC0013002 | Clust-UGC0006533 | 3853 | 4 | 1 | 0 | 1 |
| UGC0008683 | Clust-UGC0006547 | 679 | 1 | 0 | 0 | 0 |
| UGC0006547 | Clust-UGC0006547 | 899 | 1 | 0 | 0 | 0 |
| UGC0011443 | Clust-UGC0006565 | 5114 | 4 | 1 | 0 | 0 |
| UGC0009484 | Clust-UGC0006565 | 4923 | 10 | 1 | 0 | 1 |
| UGC0006565 | Clust-UGC0006565 | 3005 | 2 | 1 | 0 | 0 |
| UGC0010195 | Clust-UGC0006566 | 5464 | 2 | 1 | 0 | 1 |
| UGC0006566 | Clust-UGC0006566 | 2188 | 3 | 1 | 0 | 1 |
| UGC0007200 | Clust-UGC0006612 | 878 | 1 | 0 | 0 | 0 |
| UGC0008311 | Clust-UGC0006612 | 826 | 1 | 0 | 0 | 0 |
| UGC0013266 | Clust-UGC0006612 | 346 | 3 | 1 | 2 | 0 |
| UGC0006612 | Clust-UGC0006612 | 1228 | 1 | 0 | 0 | 0 |
| UGC0012929 | Clust-UGC0006612 | 3404 | 2 | 1 | 0 | 1 |
| UGC0011846 | Clust-UGC0006612 | 6312 | 5 | 1 | 0 | 1 |
| UGC0011534 | Clust-UGC0006612 | 1646 | 1 | 0 | 0 | 0 |
| UGC0012410 | Clust-UGC0006612 | 3499 | 4 | 1 | 0 | 1 |
| UGC0011606 | Clust-UGC0006612 | 1926 | 7 | 1 | 0 | 1 |
| UGC0007742 | Clust-UGC0006612 | 5211 | 6 | 1 | 0 | 1 |
| UGC0006653 | Clust-UGC0006653 | 4112 | 4 | 1 | 0 | 1 |
| UGC0007261 | Clust-UGC0006654 | 6524 | 1 | 0 | 0 | 0 |
| UGC0006654 | Clust-UGC0006654 | 3093 | 1 | 0 | 0 | 0 |
| UGC0007589 | Clust-UGC0006654 | 458 | 1 | 0 | 0 | 0 |
| UGC0006661 | Clust-UGC0006661 | 1982 | 1 | 0 | 0 | 0 |
| UGC0010090 | Clust-UGC0006672 | 1399 | 2 | 1 | 0 | 1 |
| UGC0006672 | Clust-UGC0006672 | 1864 | 4 | 1 | 0 | 1 |
| UGC0008387 | Clust-UGC0006672 | 361 | 1 | 0 | 0 | 0 |
| UGC0008284 | Clust-UGC0006672 | 631 | 2 | 1 | 0 | 1 |
| UGC0008855 | Clust-UGC0006672 | 811 | 1 | 0 | 0 | 0 |

| | | | | | | |
|---|---|---|---|---|---|---|
| UGC0011670 | Clust-UGC0006678 | 6248 | 6 | 1 | 0 | 1 |
| UGC0006678 | Clust-UGC0006678 | 2589 | 1 | 0 | 0 | 0 |
| UGC0006687 | Clust-UGC0006687 | 2960 | 4 | 1 | 0 | 1 |
| UGC0007506 | Clust-UGC0006704 | 9354 | 6 | 1 | 0 | 1 |
| UGC0009706 | Clust-UGC0006704 | 1447 | 1 | 0 | 0 | 0 |
| UGC0006704 | Clust-UGC0006704 | 7637 | 7 | 1 | 0 | 1 |
| UGC0006738 | Clust-UGC0006704 | 656 | 2 | 1 | 0 | 0 |
| UGC0006733 | Clust-UGC0006733 | 3891 | 1 | 0 | 0 | 0 |
| UGC0008861 | Clust-UGC0006754 | 832 | 1 | 0 | 0 | 0 |
| UGC0006754 | Clust-UGC0006754 | 982 | 1 | 0 | 0 | 0 |
| UGC0012125 | Clust-UGC0006754 | 4762 | 3 | 1 | 0 | 0 |
| UGC0009395 | Clust-UGC0006754 | 3702 | 2 | 1 | 0 | 0 |
| UGC0006756 | Clust-UGC0006756 | 3691 | 5 | 1 | 0 | 1 |
| UGC0006792 | Clust-UGC0006792 | 4283 | 4 | 1 | 0 | 1 |
| UGC0006798 | Clust-UGC0006798 | 3931 | 1 | 0 | 0 | 0 |
| UGC0006805 | Clust-UGC0006805 | 4572 | 4 | 1 | 0 | 1 |
| UGC0006814 | Clust-UGC0006814 | 8112 | 24 | 1 | 0 | 1 |
| UGC0010902 | Clust-UGC0006814 | 5167 | 6 | 1 | 1 | 1 |
| UGC0013146 | Clust-UGC0006836 | 757 | 2 | 1 | 0 | 0 |
| UGC0006836 | Clust-UGC0006836 | 5595 | 10 | 1 | 0 | 1 |
| UGC0012547 | Clust-UGC0006836 | 3807 | 8 | 1 | 0 | 1 |
| UGC0006977 | Clust-UGC0006841 | 4351 | 5 | 1 | 0 | 1 |
| UGC0010330 | Clust-UGC0006841 | 205 | 2 | 1 | 0 | 1 |
| UGC0008553 | Clust-UGC0006841 | 3990 | 2 | 1 | 0 | 1 |
| UGC0006841 | Clust-UGC0006841 | 3570 | 1 | 0 | 0 | 0 |
| UGC0006892 | Clust-UGC0006863 | 2908 | 4 | 1 | 0 | 0 |
| UGC0006863 | Clust-UGC0006863 | 3688 | 2 | 1 | 0 | 0 |
| UGC0007459 | Clust-UGC0006906 | 1113 | 1 | 0 | 0 | 0 |
| UGC0006906 | Clust-UGC0006906 | 1611 | 2 | 1 | 0 | 1 |
| UGC0009215 | Clust-UGC0006906 | 1682 | 2 | 1 | 0 | 1 |
| UGC0010161 | Clust-UGC0006920 | 4711 | 4 | 1 | 0 | 1 |
| UGC0010500 | Clust-UGC0006920 | 2430 | 2 | 1 | 0 | 1 |
| UGC0012006 | Clust-UGC0006920 | 3272 | 4 | 1 | 0 | 1 |
| UGC0010175 | Clust-UGC0006920 | 517 | 2 | 1 | 0 | 1 |
| UGC0006920 | Clust-UGC0006920 | 1510 | 1 | 0 | 0 | 0 |
| UGC0007599 | Clust-UGC0006921 | 3007 | 1 | 0 | 0 | 0 |
| UGC0007059 | Clust-UGC0006921 | 422 | 1 | 0 | 0 | 0 |
| UGC0006921 | Clust-UGC0006921 | 1631 | 2 | 1 | 1 | 0 |
| UGC0007230 | Clust-UGC0006921 | 4446 | 2 | 1 | 0 | 1 |
| UGC0009676 | Clust-UGC0006921 | 1334 | 1 | 0 | 0 | 0 |
| UGC0010142 | Clust-UGC0006927 | 3512 | 2 | 1 | 0 | 1 |
| UGC0006927 | Clust-UGC0006927 | 3399 | 6 | 1 | 0 | 1 |
| UGC0010465 | Clust-UGC0006932 | 6883 | 10 | 1 | 0 | 1 |
| UGC0006932 | Clust-UGC0006932 | 2263 | 2 | 1 | 0 | 1 |
| UGC0006934 | Clust-UGC0006934 | 1203 | 1 | 0 | 0 | 0 |
| UGC0007670 | Clust-UGC0006940 | 7572 | 4 | 1 | 0 | 1 |
| UGC0006940 | Clust-UGC0006940 | 5562 | 2 | 1 | 0 | 1 |
| UGC0006967 | Clust-UGC0006967 | 6387 | 13 | 1 | 0 | 1 |
| UGC0006975 | Clust-UGC0006971 | 5964 | 5 | 1 | 0 | 1 |
| UGC0006971 | Clust-UGC0006971 | 6877 | 7 | 1 | 0 | 1 |
| UGC0008324 | Clust-UGC0006971 | 1240 | 2 | 1 | 0 | 1 |
| UGC0008297 | Clust-UGC0006980 | 736 | 1 | 0 | 0 | 0 |
| UGC0006980 | Clust-UGC0006980 | 6519 | 11 | 1 | 0 | 1 |
| UGC0007003 | Clust-UGC0007003 | 2579 | 2 | 1 | 0 | 1 |
| UGC0007034 | Clust-UGC0007034 | 3065 | 2 | 1 | 0 | 0 |
| UGC0007060 | Clust-UGC0007060 | 114 | 1 | 0 | 0 | 0 |
| UGC0009362 | Clust-UGC0007060 | 1381 | 1 | 0 | 0 | 0 |
| UGC0009369 | Clust-UGC0007060 | 4096 | 3 | 1 | 0 | 1 |
| UGC0010535 | Clust-UGC0007060 | 659 | 3 | 1 | 2 | 0 |
| UGC0007062 | Clust-UGC0007062 | 1480 | 1 | 0 | 0 | 0 |
| UGC0007068 | Clust-UGC0007068 | 1739 | 3 | 1 | 1 | 1 |
| UGC0012258 | Clust-UGC0007068 | 232 | 1 | 0 | 0 | 0 |
| UGC0011840 | Clust-UGC0007068 | 2757 | 4 | 1 | 0 | 0 |
| UGC0008793 | Clust-UGC0007068 | 2248 | 6 | 1 | 0 | 1 |
| UGC0007737 | Clust-UGC0007074 | 4791 | 3 | 1 | 0 | 1 |
| UGC0007074 | Clust-UGC0007074 | 3429 | 2 | 1 | 0 | 1 |
| UGC0007092 | Clust-UGC0007092 | 1048 | 1 | 0 | 0 | 0 |
| UGC0007093 | Clust-UGC0007093 | 501 | 1 | 0 | 0 | 0 |
| UGC0007106 | Clust-UGC0007106 | 1190 | 2 | 1 | 0 | 0 |
| UGC0007120 | Clust-UGC0007120 | 2427 | 2 | 1 | 0 | 1 |
| UGC0009769 | Clust-UGC0007120 | 3779 | 2 | 1 | 0 | 1 |
| UGC0009255 | Clust-UGC0007120 | 15585 | 150 | 1 | 0 | 1 |
| UGC0010637 | Clust-UGC0007125 | 725 | 2 | 1 | 0 | 0 |
| UGC0007596 | Clust-UGC0007125 | 6317 | 4 | 1 | 0 | 1 |
| UGC0007125 | Clust-UGC0007125 | 761 | 2 | 1 | 0 | 0 |
| UGC0010239 | Clust-UGC0007125 | 774 | 1 | 0 | 0 | 0 |
| UGC0007147 | Clust-UGC0007147 | 771 | 1 | 0 | 0 | 0 |
| UGC0007155 | Clust-UGC0007155 | 3164 | 1 | 0 | 0 | 0 |
| UGC0010493 | Clust-UGC0007155 | 5348 | 4 | 1 | 1 | 1 |

| | | | | | | |
|---|---|---|---|---|---|---|
| UGC0007156 | Clust-UGC0007156 | 3753 | 4 | 1 | 0 | 1 |
| UGC0010205 | Clust-UGC0007157 | 804 | 4 | 1 | 1 | 0 |
| UGC0008196 | Clust-UGC0007157 | 886 | 1 | 0 | 0 | 0 |
| UGC0007157 | Clust-UGC0007157 | 2087 | 3 | 1 | 0 | 1 |
| UGC0008010 | Clust-UGC0007157 | 3794 | 5 | 1 | 0 | 1 |
| UGC0008425 | Clust-UGC0007157 | 96 | 1 | 0 | 0 | 0 |
| UGC0011960 | Clust-UGC0007157 | 1705 | 2 | 1 | 0 | 1 |
| UGC0007164 | Clust-UGC0007164 | 1287 | 2 | 1 | 0 | 0 |
| UGC0007165 | Clust-UGC0007165 | 121 | 1 | 0 | 0 | 0 |
| UGC0011411 | Clust-UGC0007165 | 6628 | 6 | 1 | 0 | 1 |
| UGC0007186 | Clust-UGC0007186 | 1724 | 1 | 0 | 0 | 0 |
| UGC0008925 | Clust-UGC0007186 | 5191 | 5 | 1 | 0 | 1 |
| UGC0013247 | Clust-UGC0007191 | 1723 | 2 | 1 | 0 | 1 |
| UGC0007191 | Clust-UGC0007191 | 4148 | 9 | 1 | 0 | 1 |
| UGC0012393 | Clust-UGC0007191 | 7679 | 22 | 1 | 0 | 1 |
| UGC0007213 | Clust-UGC0007213 | 1498 | 2 | 1 | 0 | 1 |
| UGC0007229 | Clust-UGC0007229 | 5708 | 11 | 1 | 0 | 1 |
| UGC0007247 | Clust-UGC0007247 | 7377 | 14 | 1 | 0 | 1 |
| UGC0011653 | Clust-UGC0007247 | 4851 | 4 | 1 | 0 | 1 |
| UGC0011983 | Clust-UGC0007278 | 2925 | 2 | 1 | 0 | 0 |
| UGC0007278 | Clust-UGC0007278 | 5765 | 5 | 1 | 0 | 1 |
| UGC0007454 | Clust-UGC0007278 | 2705 | 1 | 0 | 0 | 0 |
| UGC0007515 | Clust-UGC0007278 | 3795 | 2 | 1 | 0 | 0 |
| UGC0007292 | Clust-UGC0007292 | 5478 | 3 | 1 | 0 | 1 |
| UGC0012920 | Clust-UGC0007298 | 4244 | 7 | 1 | 0 | 1 |
| UGC0007298 | Clust-UGC0007298 | 7937 | 28 | 1 | 0 | 1 |
| UGC0007336 | Clust-UGC0007336 | 3970 | 13 | 1 | 0 | 1 |
| UGC0007360 | Clust-UGC0007360 | 3292 | 1 | 0 | 0 | 0 |
| UGC0007374 | Clust-UGC0007374 | 2807 | 1 | 0 | 0 | 0 |
| UGC0013194 | Clust-UGC0007406 | 423 | 1 | 0 | 0 | 0 |
| UGC0007406 | Clust-UGC0007406 | 1509 | 2 | 1 | 0 | 1 |
| UGC0010319 | Clust-UGC0007406 | 739 | 1 | 0 | 0 | 0 |
| UGC0008567 | Clust-UGC0007406 | 3610 | 2 | 1 | 0 | 1 |
| UGC0007990 | Clust-UGC0007406 | 832 | 1 | 0 | 0 | 0 |
| UGC0012565 | Clust-UGC0007406 | 4798 | 2 | 1 | 0 | 0 |
| UGC0010517 | Clust-UGC0007406 | 5654 | 7 | 1 | 0 | 1 |
| UGC0013107 | Clust-UGC0007415 | 1804 | 2 | 1 | 0 | 1 |
| UGC0011459 | Clust-UGC0007415 | 8837 | 38 | 1 | 0 | 1 |
| UGC0007415 | Clust-UGC0007415 | 4624 | 3 | 1 | 0 | 1 |
| UGC0008804 | Clust-UGC0007415 | 3063 | 2 | 1 | 0 | 0 |
| UGC0010794 | Clust-UGC0007418 | 6561 | 12 | 1 | 1 | 1 |
| UGC0007418 | Clust-UGC0007418 | 5425 | 5 | 1 | 0 | 1 |
| UGC0013111 | Clust-UGC0007418 | 729 | 1 | 0 | 0 | 0 |
| UGC0007421 | Clust-UGC0007421 | 2868 | 1 | 0 | 0 | 0 |
| UGC0008291 | Clust-UGC0007421 | 1397 | 1 | 0 | 0 | 0 |
| UGC0007893 | Clust-UGC0007421 | 5087 | 1 | 0 | 0 | 0 |
| UGC0012666 | Clust-UGC0007421 | 4600 | 3 | 1 | 0 | 1 |
| UGC0008906 | Clust-UGC0007421 | 2821 | 2 | 1 | 0 | 1 |
| UGC0007441 | Clust-UGC0007441 | 1073 | 2 | 1 | 0 | 1 |
| UGC0011884 | Clust-UGC0007456 | 3635 | 2 | 1 | 0 | 0 |
| UGC0007456 | Clust-UGC0007456 | 1328 | 1 | 0 | 0 | 0 |
| UGC0011428 | Clust-UGC0007464 | 9785 | 42 | 1 | 0 | 1 |
| UGC0007464 | Clust-UGC0007464 | 8768 | 33 | 1 | 0 | 1 |
| UGC0011200 | Clust-UGC0007464 | 771 | 1 | 0 | 0 | 0 |
| UGC0009129 | Clust-UGC0007464 | 7185 | 18 | 1 | 0 | 1 |
| UGC0011938 | Clust-UGC0007464 | 9136 | 36 | 1 | 0 | 1 |
| UGC0007613 | Clust-UGC0007464 | 5099 | 10 | 1 | 0 | 1 |
| UGC0007475 | Clust-UGC0007475 | 2480 | 1 | 0 | 0 | 0 |
| UGC0007484 | Clust-UGC0007484 | 2183 | 3 | 1 | 0 | 1 |
| UGC0007509 | Clust-UGC0007509 | 7108 | 20 | 1 | 0 | 1 |
| UGC0007512 | Clust-UGC0007512 | 5663 | 10 | 1 | 0 | 1 |
| UGC0007523 | Clust-UGC0007523 | 736 | 1 | 0 | 0 | 0 |
| UGC0010180 | Clust-UGC0007532 | 4123 | 2 | 1 | 0 | 1 |
| UGC0007532 | Clust-UGC0007532 | 4898 | 5 | 1 | 1 | 1 |
| UGC0012962 | Clust-UGC0007543 | 11278 | 81 | 1 | 0 | 1 |
| UGC0007543 | Clust-UGC0007543 | 4947 | 3 | 1 | 0 | 1 |
| UGC0007988 | Clust-UGC0007544 | 1448 | 1 | 0 | 0 | 0 |
| UGC0007544 | Clust-UGC0007544 | 2616 | 4 | 1 | 0 | 1 |
| UGC0007622 | Clust-UGC0007622 | 3172 | 2 | 1 | 0 | 0 |
| UGC0007631 | Clust-UGC0007631 | 6028 | 12 | 1 | 0 | 1 |
| UGC0009294 | Clust-UGC0007631 | 755 | 2 | 1 | 0 | 0 |
| UGC0007642 | Clust-UGC0007642 | 5903 | 4 | 1 | 0 | 1 |
| UGC0007643 | Clust-UGC0007643 | 5201 | 8 | 1 | 0 | 1 |
| UGC0009125 | Clust-UGC0007689 | 3720 | 6 | 1 | 0 | 1 |
| UGC0007689 | Clust-UGC0007689 | 8137 | 29 | 1 | 0 | 1 |
| UGC0008239 | Clust-UGC0007689 | 2332 | 2 | 1 | 0 | 1 |
| UGC0007701 | Clust-UGC0007701 | 6212 | 10 | 1 | 0 | 1 |
| UGC0007734 | Clust-UGC0007734 | 8919 | 26 | 1 | 0 | 1 |
| UGC0007752 | Clust-UGC0007752 | 3851 | 2 | 1 | 0 | 1 |

| | | | | | | |
|---|---|---|---|---|---|---|
| UGC0011565 | Clust-UGC0007767 | 7063 | 16 | 1 | 0 | 1 |
| UGC0012891 | Clust-UGC0007767 | 6757 | 15 | 1 | 0 | 1 |
| UGC0007767 | Clust-UGC0007767 | 1496 | 2 | 1 | 0 | 1 |
| UGC0007775 | Clust-UGC0007775 | 1041 | 1 | 0 | 0 | 0 |
| UGC0008754 | Clust-UGC0007775 | 408 | 1 | 0 | 0 | 0 |
| UGC0009212 | Clust-UGC0007775 | 1522 | 1 | 0 | 0 | 0 |
| UGC0010101 | Clust-UGC0007775 | 9859 | 28 | 1 | 0 | 1 |
| UGC0007779 | Clust-UGC0007779 | 512 | 1 | 0 | 0 | 0 |
| UGC0007904 | Clust-UGC0007904 | 7381 | 4 | 1 | 0 | 1 |
| UGC0011831 | Clust-UGC0007904 | 1355 | 1 | 0 | 0 | 0 |
| UGC0012994 | Clust-UGC0007904 | 4556 | 4 | 1 | 0 | 1 |
| UGC0007917 | Clust-UGC0007917 | 5964 | 18 | 1 | 1 | 1 |
| UGC0007923 | Clust-UGC0007923 | 1700 | 1 | 0 | 0 | 0 |
| UGC0007931 | Clust-UGC0007931 | 1960 | 5 | 1 | 1 | 1 |
| UGC0007935 | Clust-UGC0007935 | 6291 | 10 | 1 | 0 | 1 |
| UGC0008328 | Clust-UGC0007962 | 969 | 2 | 1 | 0 | 0 |
| UGC0007962 | Clust-UGC0007962 | 4006 | 5 | 1 | 0 | 1 |
| UGC0007964 | Clust-UGC0007964 | 692 | 2 | 1 | 0 | 0 |
| UGC0008014 | Clust-UGC0008014 | 8925 | 43 | 1 | 0 | 1 |
| UGC0008021 | Clust-UGC0008021 | 1201 | 1 | 0 | 0 | 0 |
| UGC0008028 | Clust-UGC0008028 | 1317 | 1 | 0 | 0 | 0 |
| UGC0008035 | Clust-UGC0008035 | 3004 | 3 | 1 | 0 | 1 |
| UGC0011014 | Clust-UGC0008035 | 2756 | 3 | 1 | 0 | 1 |
| UGC0012975 | Clust-UGC0008035 | 4910 | 6 | 1 | 0 | 1 |
| UGC0009156 | Clust-UGC0008035 | 377 | 1 | 0 | 0 | 0 |
| UGC0008067 | Clust-UGC0008067 | 919 | 1 | 0 | 0 | 0 |
| UGC0008602 | Clust-UGC0008067 | 1849 | 2 | 1 | 0 | 1 |
| UGC0011463 | Clust-UGC0008075 | 3300 | 6 | 1 | 0 | 1 |
| UGC0009436 | Clust-UGC0008075 | 3184 | 3 | 1 | 0 | 1 |
| UGC0009512 | Clust-UGC0008075 | 115 | 1 | 0 | 0 | 0 |
| UGC0008075 | Clust-UGC0008075 | 3087 | 1 | 0 | 0 | 0 |
| UGC0008203 | Clust-UGC0008203 | 2178 | 1 | 0 | 0 | 0 |
| UGC0008204 | Clust-UGC0008204 | 2363 | 1 | 0 | 0 | 0 |
| UGC0008219 | Clust-UGC0008219 | 1905 | 6 | 1 | 0 | 0 |
| UGC0008244 | Clust-UGC0008244 | 228 | 1 | 0 | 0 | 0 |
| UGC0008446 | Clust-UGC0008257 | 987 | 1 | 0 | 0 | 0 |
| UGC0008257 | Clust-UGC0008257 | 4399 | 8 | 1 | 0 | 1 |
| UGC0008258 | Clust-UGC0008258 | 298 | 1 | 0 | 0 | 0 |
| UGC0008279 | Clust-UGC0008279 | 432 | 1 | 0 | 0 | 0 |
| UGC0008303 | Clust-UGC0008303 | 1187 | 1 | 0 | 0 | 0 |
| UGC0008317 | Clust-UGC0008317 | 1452 | 1 | 0 | 0 | 0 |
| UGC0008357 | Clust-UGC0008357 | 728 | 1 | 0 | 0 | 0 |
| UGC0008376 | Clust-UGC0008376 | 1419 | 3 | 1 | 2 | 0 |
| UGC0008415 | Clust-UGC0008415 | 871 | 1 | 0 | 0 | 0 |
| UGC0008451 | Clust-UGC0008451 | 375 | 1 | 0 | 0 | 0 |
| UGC0008455 | Clust-UGC0008455 | 2233 | 1 | 0 | 0 | 0 |
| UGC0008468 | Clust-UGC0008468 | 464 | 1 | 0 | 0 | 0 |
| UGC0009593 | Clust-UGC0008532 | 5703 | 4 | 1 | 0 | 1 |
| UGC0008532 | Clust-UGC0008532 | 1556 | 1 | 0 | 0 | 0 |
| UGC0008536 | Clust-UGC0008536 | 107 | 1 | 0 | 0 | 0 |
| UGC0008537 | Clust-UGC0008537 | 2023 | 4 | 1 | 0 | 0 |
| UGC0012550 | Clust-UGC0008537 | 10170 | 19 | 1 | 0 | 1 |
| UGC0008542 | Clust-UGC0008542 | 732 | 1 | 0 | 0 | 0 |
| UGC0008561 | Clust-UGC0008561 | 5526 | 8 | 1 | 0 | 1 |
| UGC0008566 | Clust-UGC0008566 | 1775 | 6 | 1 | 0 | 1 |
| UGC0011508 | Clust-UGC0008566 | 9912 | 35 | 1 | 0 | 1 |
| UGC0008598 | Clust-UGC0008598 | 1364 | 4 | 1 | 0 | 1 |
| UGC0008625 | Clust-UGC0008625 | 5119 | 2 | 1 | 0 | 1 |
| UGC0008677 | Clust-UGC0008677 | 426 | 1 | 0 | 0 | 0 |
| UGC0008722 | Clust-UGC0008722 | 797 | 1 | 0 | 0 | 0 |
| UGC0010186 | Clust-UGC0008722 | 760 | 1 | 0 | 0 | 0 |
| UGC0008725 | Clust-UGC0008725 | 1812 | 1 | 0 | 0 | 0 |
| UGC0008737 | Clust-UGC0008737 | 1207 | 1 | 0 | 0 | 0 |
| UGC0008773 | Clust-UGC0008773 | 1467 | 1 | 0 | 0 | 0 |
| UGC0010610 | Clust-UGC0008773 | 1973 | 1 | 0 | 0 | 0 |
| UGC0008811 | Clust-UGC0008811 | 183 | 1 | 0 | 0 | 0 |
| UGC0008820 | Clust-UGC0008820 | 4447 | 5 | 1 | 0 | 1 |
| UGC0010104 | Clust-UGC0008820 | 4511 | 3 | 1 | 0 | 1 |
| UGC0008840 | Clust-UGC0008840 | 2533 | 1 | 0 | 0 | 0 |
| UGC0010135 | Clust-UGC0008840 | 2732 | 2 | 1 | 0 | 1 |
| UGC0008848 | Clust-UGC0008848 | 402 | 1 | 0 | 0 | 0 |
| UGC0008858 | Clust-UGC0008858 | 2853 | 9 | 1 | 0 | 1 |
| UGC0008902 | Clust-UGC0008902 | 1514 | 1 | 0 | 0 | 0 |
| UGC0008918 | Clust-UGC0008918 | 2546 | 2 | 1 | 0 | 1 |
| UGC0011414 | Clust-UGC0008918 | 4022 | 2 | 1 | 0 | 0 |
| UGC0008976 | Clust-UGC0008976 | 8755 | 24 | 1 | 0 | 1 |
| UGC0010910 | Clust-UGC0008976 | 9485 | 23 | 1 | 0 | 1 |
| UGC0012892 | Clust-UGC0008982 | 7603 | 15 | 1 | 0 | 1 |
| UGC0008982 | Clust-UGC0008982 | 2389 | 2 | 1 | 0 | 1 |

| | | | | | | |
|---|---|---|---|---|---|---|
| UGC0009078 | Clust-UGC0009078 | 4224 | 3 | 1 | 0 | 1 |
| UGC0009088 | Clust-UGC0009088 | 1290 | 1 | 0 | 0 | 0 |
| UGC0011667 | Clust-UGC0009088 | 5465 | 8 | 1 | 0 | 1 |
| UGC0009112 | Clust-UGC0009112 | 2782 | 1 | 0 | 0 | 0 |
| UGC0009714 | Clust-UGC0009180 | 2777 | 6 | 1 | 0 | 1 |
| UGC0010075 | Clust-UGC0009180 | 3404 | 2 | 1 | 0 | 1 |
| UGC0009180 | Clust-UGC0009180 | 5628 | 17 | 1 | 0 | 1 |
| UGC0012302 | Clust-UGC0009217 | 1311 | 1 | 0 | 0 | 0 |
| UGC0012252 | Clust-UGC0009217 | 4012 | 6 | 1 | 0 | 1 |
| UGC0011409 | Clust-UGC0009217 | 549 | 3 | 1 | 0 | 0 |
| UGC0012836 | Clust-UGC0009217 | 6611 | 10 | 1 | 2 | 1 |
| UGC0009217 | Clust-UGC0009217 | 7365 | 11 | 1 | 0 | 0 |
| UGC0009282 | Clust-UGC0009282 | 4305 | 5 | 1 | 0 | 1 |
| UGC0009310 | Clust-UGC0009310 | 2573 | 5 | 1 | 0 | 1 |
| UGC0011169 | Clust-UGC0009331 | 1523 | 2 | 1 | 0 | 0 |
| UGC0009331 | Clust-UGC0009331 | 3993 | 5 | 1 | 0 | 1 |
| UGC0009340 | Clust-UGC0009340 | 12383 | 90 | 1 | 0 | 1 |
| UGC0009387 | Clust-UGC0009387 | 4152 | 7 | 1 | 0 | 1 |
| UGC0009561 | Clust-UGC0009390 | 4685 | 1 | 0 | 0 | 0 |
| UGC0009390 | Clust-UGC0009390 | 2904 | 1 | 0 | 0 | 0 |
| UGC0010001 | Clust-UGC0009404 | 3221 | 3 | 1 | 0 | 1 |
| UGC0009404 | Clust-UGC0009404 | 5026 | 7 | 1 | 0 | 1 |
| UGC0009429 | Clust-UGC0009429 | 2630 | 2 | 1 | 0 | 1 |
| UGC0009439 | Clust-UGC0009439 | 7839 | 28 | 1 | 0 | 1 |
| UGC0009464 | Clust-UGC0009464 | 5415 | 7 | 1 | 0 | 1 |
| UGC0013112 | Clust-UGC0009464 | 529 | 2 | 1 | 1 | 0 |
| UGC0009660 | Clust-UGC0009510 | 1148 | 3 | 1 | 0 | 1 |
| UGC0009510 | Clust-UGC0009510 | 3318 | 5 | 1 | 0 | 1 |
| UGC0010595 | Clust-UGC0009510 | 3248 | 1 | 0 | 0 | 0 |
| UGC0009668 | Clust-UGC0009668 | 989 | 2 | 1 | 0 | 0 |
| UGC0009712 | Clust-UGC0009712 | 1960 | 3 | 1 | 0 | 0 |
| UGC0009733 | Clust-UGC0009733 | 1868 | 2 | 1 | 0 | 1 |
| UGC0009989 | Clust-UGC0009989 | 2468 | 2 | 1 | 0 | 0 |
| UGC0013388 | Clust-UGC0010028 | 3509 | 9 | 1 | 1 | 1 |
| UGC0010028 | Clust-UGC0010028 | 1652 | 2 | 1 | 0 | 1 |
| UGC0010081 | Clust-UGC0010081 | 2001 | 5 | 1 | 0 | 1 |
| UGC0010627 | Clust-UGC0010106 | 804 | 1 | 0 | 0 | 0 |
| UGC0010106 | Clust-UGC0010106 | 4677 | 2 | 1 | 0 | 1 |
| UGC0010109 | Clust-UGC0010109 | 7915 | 16 | 1 | 0 | 1 |
| UGC0010133 | Clust-UGC0010133 | 2661 | 3 | 1 | 0 | 1 |
| UGC0010241 | Clust-UGC0010241 | 110 | 1 | 0 | 0 | 0 |
| UGC0010310 | Clust-UGC0010310 | 1195 | 1 | 0 | 0 | 0 |
| UGC0010454 | Clust-UGC0010454 | 5608 | 3 | 1 | 0 | 1 |
| UGC0010460 | Clust-UGC0010460 | 5509 | 7 | 1 | 0 | 1 |
| UGC0010463 | Clust-UGC0010463 | 447 | 1 | 0 | 0 | 0 |
| UGC0013195 | Clust-UGC0010463 | 1712 | 1 | 0 | 0 | 0 |
| UGC0011576 | Clust-UGC0010463 | 335 | 1 | 0 | 0 | 0 |
| UGC0010466 | Clust-UGC0010466 | 1210 | 1 | 0 | 0 | 0 |
| UGC0010497 | Clust-UGC0010497 | 3444 | 4 | 1 | 0 | 1 |
| UGC0010502 | Clust-UGC0010502 | 2267 | 1 | 0 | 0 | 0 |
| UGC0013377 | Clust-UGC0010514 | 4024 | 2 | 1 | 0 | 0 |
| UGC0010514 | Clust-UGC0010514 | 685 | 1 | 0 | 0 | 0 |
| UGC0010553 | Clust-UGC0010553 | 2638 | 1 | 0 | 0 | 0 |
| UGC0010557 | Clust-UGC0010557 | 2628 | 7 | 1 | 1 | 1 |
| UGC0010584 | Clust-UGC0010584 | 4060 | 2 | 1 | 0 | 1 |
| UGC0010604 | Clust-UGC0010604 | 483 | 1 | 0 | 0 | 0 |
| UGC0011958 | Clust-UGC0010605 | 7537 | 15 | 1 | 0 | 1 |
| UGC0010605 | Clust-UGC0010605 | 2487 | 6 | 1 | 0 | 1 |
| UGC0010630 | Clust-UGC0010630 | 571 | 1 | 0 | 0 | 0 |
| UGC0010877 | Clust-UGC0010877 | 166 | 1 | 0 | 0 | 0 |
| UGC0010917 | Clust-UGC0010917 | 1591 | 1 | 0 | 0 | 0 |
| UGC0013066 | Clust-UGC0010938 | 13107 | 101 | 1 | 0 | 1 |
| UGC0010938 | Clust-UGC0010938 | 3532 | 2 | 1 | 0 | 0 |
| UGC0010980 | Clust-UGC0010980 | 474 | 1 | 0 | 0 | 0 |
| UGC0011470 | Clust-UGC0011470 | 7661 | 23 | 1 | 0 | 1 |
| UGC0011500 | Clust-UGC0011500 | 173 | 1 | 0 | 0 | 0 |
| UGC0011505 | Clust-UGC0011505 | 4793 | 9 | 1 | 0 | 1 |
| UGC0011555 | Clust-UGC0011555 | 1896 | 2 | 1 | 0 | 0 |
| UGC0012392 | Clust-UGC0011555 | 4410 | 4 | 1 | 0 | 1 |
| UGC0011928 | Clust-UGC0011557 | 3599 | 3 | 1 | 0 | 1 |
| UGC0012178 | Clust-UGC0011557 | 2121 | 1 | 0 | 0 | 0 |
| UGC0011557 | Clust-UGC0011557 | 6908 | 14 | 1 | 1 | 1 |
| UGC0012407 | Clust-UGC0011557 | 2740 | 1 | 0 | 0 | 0 |
| UGC0011574 | Clust-UGC0011574 | 3235 | 6 | 1 | 0 | 1 |
| UGC0012158 | Clust-UGC0011598 | 3453 | 5 | 1 | 1 | 1 |
| UGC0011598 | Clust-UGC0011598 | 1394 | 1 | 0 | 0 | 0 |
| UGC0011601 | Clust-UGC0011601 | 122 | 1 | 0 | 0 | 0 |
| UGC0011645 | Clust-UGC0011645 | 5999 | 5 | 1 | 0 | 1 |
| UGC0011672 | Clust-UGC0011672 | 2196 | 4 | 1 | 0 | 1 |

| | | | | | | |
|---|---|---|---|---|---|---|
| UGC0011833 | Clust-UGC0011833 | 7014 | 16 | 1 | 0 | 1 |
| UGC0011887 | Clust-UGC0011887 | 3095 | 5 | 1 | 0 | 1 |
| UGC0011923 | Clust-UGC0011923 | 4679 | 3 | 1 | 0 | 1 |
| UGC0011927 | Clust-UGC0011927 | 2229 | 2 | 1 | 0 | 1 |
| UGC0012402 | Clust-UGC0011948 | 5551 | 13 | 1 | 0 | 1 |
| UGC0011948 | Clust-UGC0011948 | 6880 | 7 | 1 | 0 | 1 |
| UGC0012616 | Clust-UGC0011954 | 6026 | 10 | 1 | 0 | 1 |
| UGC0011954 | Clust-UGC0011954 | 3301 | 1 | 0 | 0 | 0 |
| UGC0012917 | Clust-UGC0011954 | 2721 | 2 | 1 | 0 | 0 |
| UGC0011969 | Clust-UGC0011969 | 5522 | 2 | 1 | 1 | 0 |
| UGC0012007 | Clust-UGC0012007 | 2514 | 2 | 1 | 0 | 1 |
| UGC0012038 | Clust-UGC0012038 | 3519 | 2 | 1 | 0 | 1 |
| UGC0012088 | Clust-UGC0012088 | 805 | 5 | 1 | 1 | 0 |
| UGC0012150 | Clust-UGC0012150 | 976 | 1 | 0 | 0 | 0 |
| UGC0012151 | Clust-UGC0012151 | 2708 | 6 | 1 | 0 | 1 |
| UGC0012164 | Clust-UGC0012164 | 3187 | 3 | 1 | 0 | 1 |
| UGC0012216 | Clust-UGC0012216 | 1679 | 1 | 0 | 0 | 0 |
| UGC0012239 | Clust-UGC0012239 | 7759 | 8 | 1 | 0 | 1 |
| UGC0012244 | Clust-UGC0012244 | 7271 | 21 | 1 | 0 | 1 |
| UGC0012347 | Clust-UGC0012347 | 815 | 4 | 1 | 0 | 1 |
| UGC0012362 | Clust-UGC0012362 | 8676 | 33 | 1 | 0 | 1 |
| UGC0012638 | Clust-UGC0012638 | 3591 | 4 | 1 | 0 | 1 |
| UGC0012654 | Clust-UGC0012654 | 6914 | 20 | 1 | 0 | 1 |
| UGC0012690 | Clust-UGC0012690 | 9131 | 41 | 1 | 0 | 1 |
| UGC0012693 | Clust-UGC0012693 | 7025 | 13 | 1 | 0 | 1 |
| UGC0012721 | Clust-UGC0012721 | 6169 | 4 | 1 | 0 | 1 |
| UGC0012728 | Clust-UGC0012728 | 2329 | 2 | 1 | 0 | 0 |
| UGC0012758 | Clust-UGC0012758 | 3723 | 3 | 1 | 0 | 1 |
| UGC0012803 | Clust-UGC0012803 | 2811 | 1 | 0 | 0 | 0 |
| UGC0012819 | Clust-UGC0012819 | 3105 | 1 | 0 | 0 | 0 |
| UGC0012830 | Clust-UGC0012830 | 1678 | 4 | 1 | 0 | 0 |
| UGC0012848 | Clust-UGC0012848 | 3556 | 8 | 1 | 1 | 1 |
| UGC0012877 | Clust-UGC0012877 | 7505 | 17 | 1 | 0 | 1 |
| UGC0012887 | Clust-UGC0012887 | 8649 | 24 | 1 | 0 | 1 |
| UGC0012899 | Clust-UGC0012899 | 7015 | 16 | 1 | 0 | 1 |
| UGC0012906 | Clust-UGC0012906 | 3446 | 4 | 1 | 0 | 1 |
| UGC0012908 | Clust-UGC0012908 | 719 | 1 | 0 | 0 | 0 |
| UGC0012942 | Clust-UGC0012942 | 4328 | 3 | 1 | 0 | 0 |
| UGC0012979 | Clust-UGC0012979 | 3234 | 3 | 1 | 0 | 1 |
| UGC0013010 | Clust-UGC0013010 | 3255 | 2 | 1 | 0 | 0 |
| UGC0013012 | Clust-UGC0013012 | 1460 | 1 | 0 | 0 | 0 |
| UGC0013102 | Clust-UGC0013102 | 6990 | 19 | 1 | 0 | 1 |
| UGC0013143 | Clust-UGC0013143 | 161 | 1 | 0 | 0 | 0 |
| UGC0013162 | Clust-UGC0013162 | 632 | 1 | 0 | 0 | 0 |
| UGC0013185 | Clust-UGC0013185 | 784 | 2 | 1 | 0 | 1 |
| UGC0013250 | Clust-UGC0013250 | 3171 | 9 | 1 | 0 | 1 |
| UGC0013304 | Clust-UGC0013304 | 2688 | 3 | 1 | 0 | 1 |
| UGC0013315 | Clust-UGC0013315 | 5782 | 19 | 1 | 0 | 1 |

| Additional File 5: Specimens with >3% divergence from Illumina barcodes for final consolidated barcode sets. | | | | |
|---|---|---|---|---|
| **Specimen** | **Minion full** | **Minion 313** | **Illumina 313** | **Matches to trace sequences in Illumina data (>10x coverage) in the same dataset** |
| UGC0012088 | Identical to UGC0011906 | Identical to UGC0011906 | singleton (>9%) | matches to secondary signal at 99.042% in Illumina (count of 33) |
| UGC0010204 | Identical to UGC0005689 | Identical to UGC0005689 | Identical to UGC0011195 | no match |
| UGC0012864 | Identical to UGC0012864 | Identical to UGC0012864 | Identical to UGC0009545 | no match |
| UGC0012285 | Identical to UGC0008675 | Identical to UGC0008675 | Identical UGC0012286 | no match |
| UGC0013329 | Identical to UGC0013328 | Identical to UGC0013328 | Identical to UGC0002114 | matches to secondary signal at 99.681% in Illumina (count of 452) |
| UGC0011199 | Identical to UGC0008501 | Identical to UGC0008501 | Identical to UGC0005660 | matches to secondary signal at 99.361% in Illumina (count of 15) |
| UGC0010207 | Identical to UGC0005689 | Identical to UGC0005689 | Identical to UGC0005873 | no match |
| UGC0010179 | Identical to UGC0010482 | Identical to UGC0010482 | Identical to UGC0010485 | no match |
| UGC0007496 | Identical to UGC0012958 | Identical to UGC0012958 | Identical to UGC0006750 | matches to secondary signal at 99.681% in Illumina (count of 59) |
| UGC0013325 | Identical to UGC0002270 | Identical to UGC0002270 | Identical to UGC0005703 | matches to secondary signal at 99.681% in Illumina (count of 44) |
| UGC0010274 | Identical to UGC0007271 | Identical to UGC0007271 | Identical to UGC0009165 | matches to secondary signal at 99.681% in Illumina (count of 558) |
| UGC0011155 | Identical to UGC0007243 | Identical to UGC0007243 | Identical to UGC0006386 | matches to secondary signal at 99.361% in Illumina (count of 66) |
| UGC0003372 | singleton (>9%) | singleton (>9%) | singleton to 7.4% | no match |
| UGC0006133 | Identical to UGC0006068 | Identical to UGC0006068 | Identical to UGC0008527 | matches to secondary signal at 99.042% in Illumina (count of 15) |
| UGC0010126 | <0.5% away from UGC003004 | <0.3 % away from UGC003004 | Identical to UGC0008527 | matches to secondary signal at 99.042% in Illumina (count of 42) |
| UGC0003004 | Identical to UGC0008527 | Identical to UGC0008527 | Identical to UGC0008527 | no match |
| UGC0008527 | Identical to UGC0003004 | Identical to UGC0003004 | Identical to UGC0003004 | no match |
| UGC0006068 | Identical to UGC0006133 | Identical to UGC0006133 | Identical to UGC0012290 | no match |
| UGC0004742 | In 0.6% cluster with UGC0012371 | In 0.6% cluster with UGC0012371 | In 0.3% cluster with UGC0005091 | no match |
| UGC0010485 | Singleton (3.4%) | Singleton (3.4%) | Idencal to UGC0010485 | matches to secondary signal at 99.681% in Illumina (count of 252) |
| UGC0012855 | Identical to UGC0012835 | Identical to UGC0012835 | Identical to UGC0006355 | matches to secondary signal at 99.042% in Illumina (count of 82) |
| UGC0012846 | Identical to UGC0012855 | Identical to UGC0012855 | Identical to UGC0012245 | matches to secondary signal at 99.681% in Illumina (count of 37) |
| UGC0012835 | Identical to UGC0012855 | Identical to UGC0012855 | Identical to UGC0012245 | matches to secondary signal at 99.681% in Illumina (count of 73) |
| UGC0012624 | Identical to UGC0012855 | Identical to UGC0012855 | Identical to UGC0012245 | matches to secondary signal at 99.681% in Illumina (count of 24) |
| UGC0013225 | Identical to UGC0012368 | Identical to UGC0012368 | Identical to UGC0004621 | matches to secondary signal at 100% in Illumina (count of 43) |
